# The molecular structure of plant sporopollenin

**DOI:** 10.1101/415612

**Authors:** Fu-Shuang Li, Pyae Phyo, Joseph Jacobowitz, Mei Hong, Jing-Ke Weng

## Abstract

Sporopollenin is a ubiquitous and extremely chemically inert biopolymer that constitutes the outer wall of all land-plant spores and pollen grains. Sporopollenin protects the vulnerable plant gametes against a wide range of environmental assaults, and is considered as a prerequisite for the migration of early plants onto land. Despite its importance, the chemical structure of plant sporopollenin has remained elusive. Using a newly developed thioacidolysis degradative method together with state-of-the-art solid-state NMR techniques, we determined the detailed molecular structure of pine sporopollenin. We show that pine sporopollenin is primarily composed of aliphatic-polyketide-derived polyvinyl alcohol units and 7-*O*-*p*-coumaroylated C16 aliphatic units, crosslinked through a distinctive m-dioxane moiety featuring an acetal. Naringenin was also identified as a minor component of pine sporopollenin. This discovery answers the long-standing question about the chemical makeup of plant sporopollenin, laying the foundation for future investigations of sporopollenin biosynthesis and for design of new biomimetic polymers with desirable inert properties.

---

Early land plants arose approximately 450 million years ago. To cope with a multitude of abiotic and biotic stresses associated with the terrestrial environments, plants evolved the ability to synthesize numerous specialized biopolymers, including lignin, condensed tannin, cutin, suberin, natural rubber, and sporopollenin. Each of these biopolymers possesses a unique set of physicochemical properties pertinent to their biological functions^1^. Whereas the chemical structures of most plant biopolymers have been elucidated, sporopollenin stands as one of the last and least-understood biopolymers regarding its structural composition^2^. While enabling plant spores and pollen to travel long distances and survive hostile conditions, the extreme chemical inertness of sporopollenin has presented a major technical challenge to its full structure elucidation.

Using relatively harsh chemical degradation methods and whole-polymer spectroscopic methods, previous studies have suggested that sporopollenin contains hydroxylated aliphatic and aromatic units (Supplementary Note 1). Extensive genetic studies of plant mutants defective in pollen exine formation have also identified a series of metabolic enzymes that are highly conserved among land plants and apparently involved in sporopollenin biosynthesis^2^. Many of these enzymes exhibit *in vitro* biochemical specificities that suggest their functions in fatty acid and polyketide metabolism^2^. However, the exact monomer composition and inter-monomer linkages of sporopollenin are unknown. Consequently, the precise sporopollenin biosynthetic pathway has not been established, even though many of the participating enzymes have been identified.

In this study, we interrogated sporopollenin polymer structure by exploring new chemical and biophysical methodologies. Unlike most of the other plant biopolymers, sporopollenin is challenging to obtain in large quantities from model plants that are commonly used for lab research. We therefore chose to study sporopollenin of the pitch pine *Pinus rigida*, from which kilogram quantities of pollen can be collected for analysis (Fig. 1A). Inspired by the recent adoption of high-energy ball milling to increase lignin solubility in organic solvents^3^, we incorporated ball milling in an improved protocol^4^ for preparing pine sporopollenin (Figs. 1A, B). Ball milling of whole pollen grains produced a fine powder, which was hydrolyzed enzymatically by cellulase cocktails, and washed excessively with a series of solvents with increasing polarity. The remaining insoluble residue was lyophilized to yield the total pine sporopollenin.

**Fig. 1.**
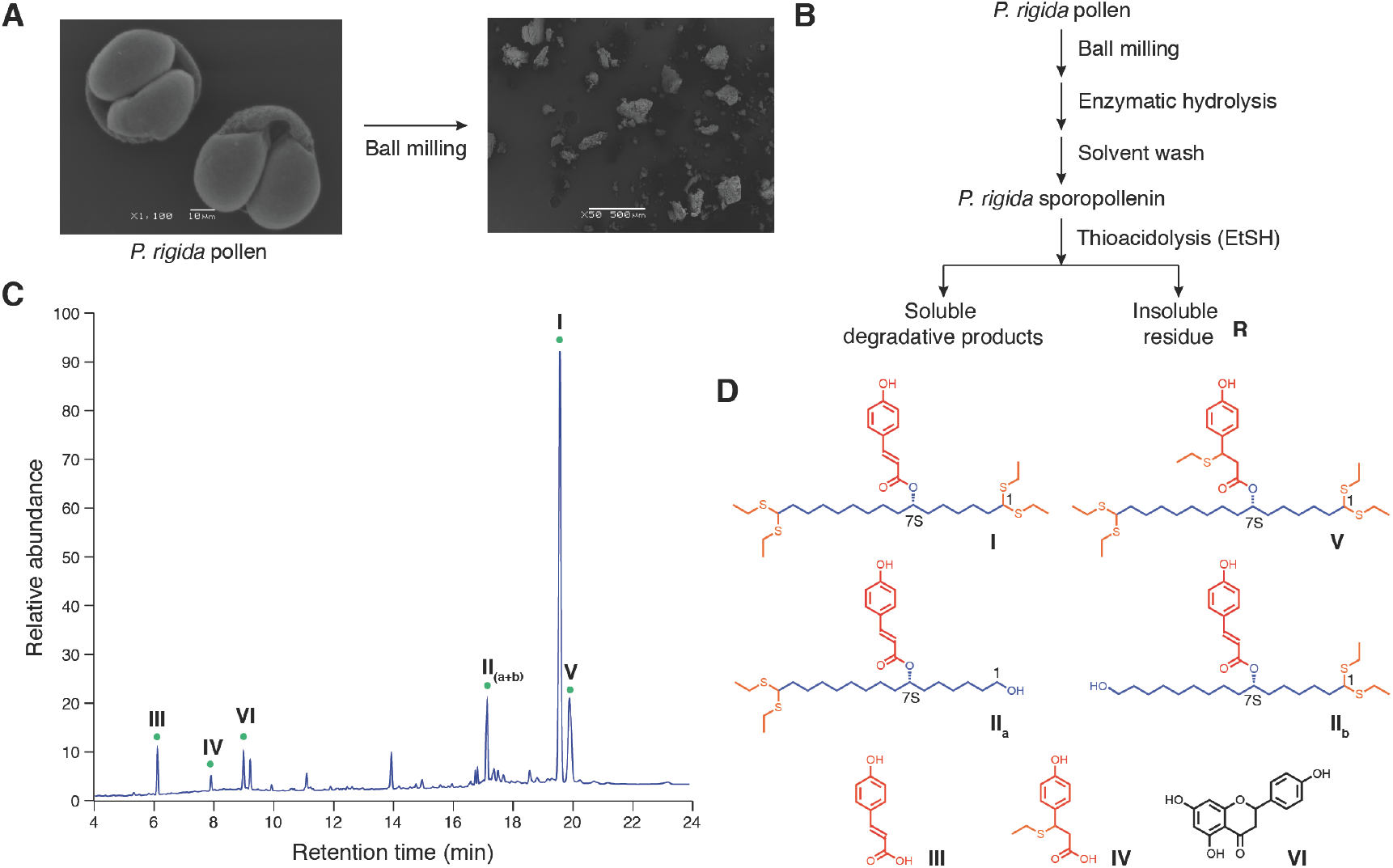
Pine sporopollenin preparation and degradative analysis by thioacidolysis. **A**, Electron micrographs of *P. rigida* pollen grains and the fine powder derived from them after ball milling. **B**, The optimized protocol for pine sporopollenin preparation and thioacidolysis degradation using ethyl mercaptan (EtSH). **C**, Thioacidolysis of pine sporopollenin gave three major (**I**, **II_a_**, **II_b_**) and several minor (**III**-**VI**) UV-detectable degradative products under HPLC-UV-MS analysis. **D**, Structural elucidation of the major thioacidolysis products of pine sporopollenin. The Roman numerals correspond to HPLC peaks as shown in **C**.

Structure elucidation of polymers benefits from the development of suitable enzymatic and chemical degradation methods that can provide information about polymer fragments, from which the whole molecular structure can be deduced. To establish a degradative method for sporopollenin, we surveyed a variety of chemical reagents known to depolymerize other biopolymers (Supplementary Note 2). From this survey, several thioacidolysis reagents^5^, namely benzyl mercaptan, n-dodecyl mercaptan and ethyl mercaptan, were found to partially degrade pine sporopollenin under acidic conditions and to yield a number of small-molecular-weight compounds readily detectable by high-performance liquid chromatography coupled in tandem with UV detector and mass spectrometer (HPLC-UV-MS) (Supplementary Figs. 5, 40, 41). We further optimized an ethyl-mercaptan-based thioacidolysis protocol, and observed that 55±3% (w/w) of the starting pine sporopollenin can be degraded under this condition. HPLC-UV-MS analysis of the soluble fraction of the reaction products gave three major and several minor UV-detectable degradation products (Figs. 1B, C and Supplementary Fig. 6). These compounds were purified chromatographically, and their structures were determined by a combination of MS/MS and ^1^H, ^13^C, HSQC, HMBC NMR analyses (Fig. 1D and Supplementary Figs. 11, 12, 20-23).

Compound **I** is the most abundant thioacidolysis product from the soluble fraction, and features a *p*-coumaroylated 7-OH-C16 aliphatic core flanked by two diethylsulfanyl groups on both ends (Fig. 1D). Compounds **II_a_** and **II_b_** are chromatographically inseparable and were therefore co-purified as one fraction. Upon detailed NMR analysis of the mixture (Supplementary Fig. 12), **II_a_** and **II_b_** were identified to be related to **I**, with either the 1- or 16-diethylsulfanyl group of **I** substituted by a free hydroxy group (Fig. 1D). We also identified *p*-coumaric acid (**III**) and naringenin (**VI**) as minor thioacidolysis products, suggesting that they exist as covalently linked structural units in pine sporopollenin, although **III** could be solely derived from **I** and **II_(a+b)_** through extensive acid hydrolysis (Figs. 1C, D and Supplementary Fig. 20). Compounds **IV** and **V** are additional minor products, which were verified to be derived from **III** and **I** respectively through nucleophilic addition of an ethylsulfanyl group to the *p*-coumaroyl olefin during the thioacidolysis reaction (Figs. 1C, D and Supplementary Figs. 21, 22). Based on peak area integration of the HPLC-UV chromatogram, the molar ratio of **I** to **II_(a+b)_** is estimated to be approximately 6∼7:1 (Supplementary Fig. 6).

To probe the origin of various free hydroxy groups observed in **I** and **II_(a+b)_** prior to thioacidolysis, we prepared methylated and acetylated total pine sporopollenin samples, respectively, to methylate or acetylate any free hydroxy and carboxy groups that may be present in the sporopollenin polymer. Subsequent thioacidolysis experiments starting with methylated or acetylated sporopollenin produced the corresponding methylated or acetylated **I** and **II_(a+b)_** respectively (Fig. 2A and Supplementary Figs. 1, 2, 7, 8, 13-19), suggesting that the *p*-OH group of the *p*-coumaroyl moiety of **I** and **II_(a+b)_** and the ω-OH group of **II_(a+b)_** exist as free hydroxy groups not engaged in inter-unit coupling in sporopollenin polymer.

**Fig. 2.**
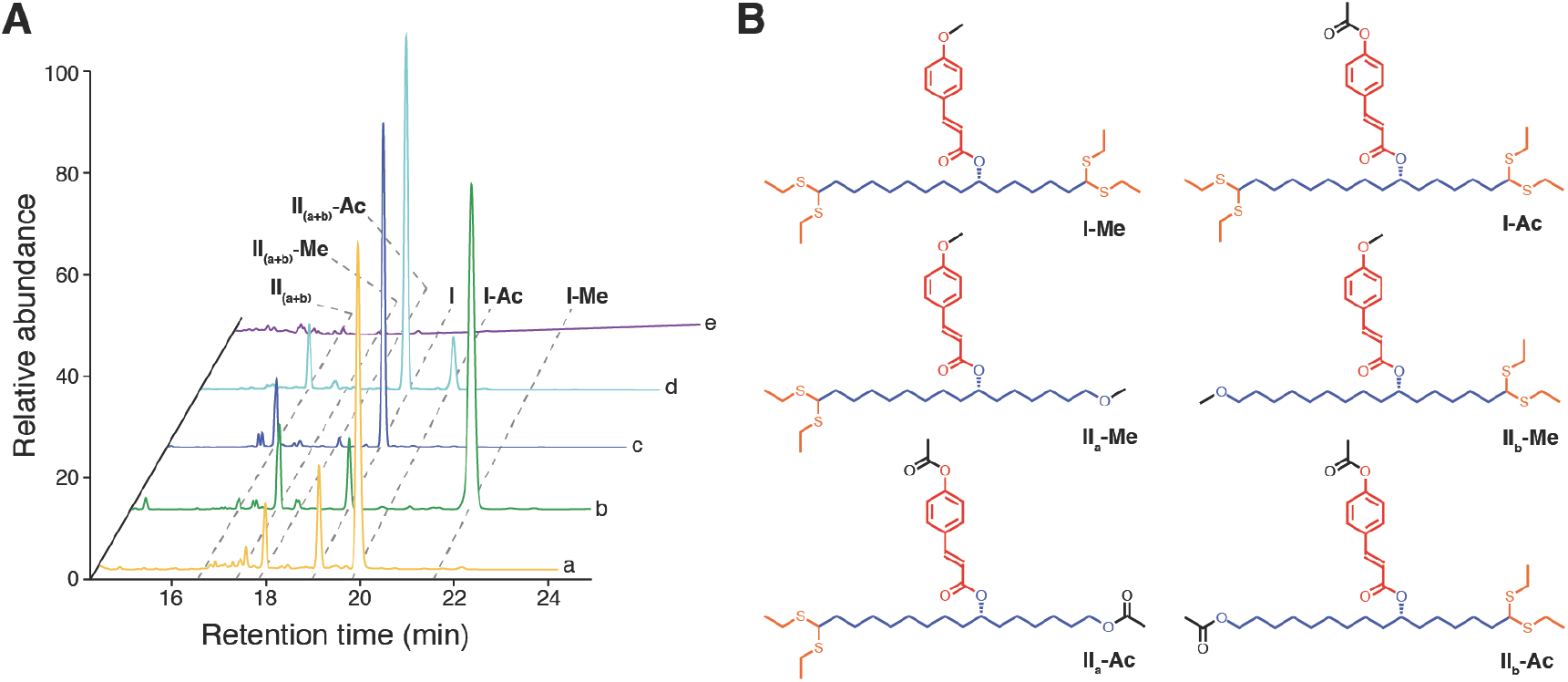
Structural characterization of pine sporopollenin by whole-polymer derivatization followed by thioacidolysis. **A**, HPLC-UV chromatograms of thioacidolysis products after various pretreatments. The five traces are: (a), Acetylated pine sporopollenin; (b), Methylated pine sporopollenin; (c), Untreated pine sporopollenin; (d), Acetylated pine sporopollenin subsequently treated with NaBH_4_, and (e), Acetylated pine sporopollenin subsequently treated with TMDS/Cu(OTf)_2_. **B**, Structural elucidation of thioacidolysis products from methylated or acetylated whole sporopollenin polymer. These results show that the *p*-OH group of the *p*-coumaroyl moiety of **I** and **II_(a+b)_** as well as the ω-OH of **II_(a+b)_** exist as free hydroxy groups not engaged in interunit coupling in sporopollenin polymer.

The *p*-coumaroyl moiety of **I** and **II_(a+b)_** is linked to the C16 aliphatic core through a chiral carbon at the C_7_ position. Chiral HPLC analysis of **I** suggests a predominant enantiomeric configuration (>97%) (Supplementary Fig. 33). However, the very similar alkyl neighborhoods flanking each side of this chiral center makes it difficult to determine the absolute configuration of this chiral center using the conventional Mosher’s method^6,7^. Moreover, circular dichroism (CD) measurement of **I** gave only baseline spectrum (Supplementary Fig. 42), while the oily nature of **I** also precludes structural elucidation by conventional X-ray crystallography. We therefore turned to a pair of recently reported chiral labeling reagents, (1*R*,2*R*)-(-)-2-(2,3-anthracenedicarboximide) cyclohexane carboxylic acid and (1*S*,2*S*)-(+)-2-(2,3-anthracenedicarboximide) cyclohexane carboxylic acid^8,9^, and used them as modified Mosher’s reagents capable of engaging in long-distance spatial interactions. By reacting **I_a_** (de-esterified **I,** Supplementary Figs. S34 and S35) with these two chiral labeling reagents, we generated two corresponding diastereomers (**I_a_-*RR*** and **I_a_-*SS***) (Supplementary Figs. 32, 36, 37). Comparison of their ^1^H-NMR spectra led us to unequivocally assign the absolute configuration of C_7_ to be *S* (Supplementary Fig. 38).

Thioacidolysis of *β*-*O*-4-linked lignin using ethyl mercaptan results in monoethylsulfanyl substitutions at *α, β* and *γ* positions of the cleaved monolignols via a well-characterized ether-cleavage mechanism^5^ (Supplementary Fig. 43). However, the diethylsulfanyl moiety observed at either or both ends of the main sporopollenin thioacidolysis products indicates a different linkage prior to cleavage. Previous studies of thioacidolysis chemistry using a panel of model compounds showed that reacting aldehyde or acetal with ethyl mercaptan gives diethylsulfanyl derivatives^5^. We then tested whether the diethylsulfanyl moiety of the sporopollenin thioacidolysis products could be derived from aldehyde and/or acetal functional groups present in sporopollenin polymer. We first performed sporopollenin pretreatment with sodium borohydride (NaBH_4_), a reagent that reduces aldehydes to the corresponding alcohols and therefore renders them non-reactive to ethyl mercaptan. However, subsequent thioacidolysis was not impacted by NaBH_4_ pretreatment (Fig. 2A), suggesting that the diethylsulfanyl moiety is unlikely derived from free aldehyde. We then employed a newly reported reagent, TMDS/Cu(OTf)_2_, which was shown to selectively cleave acetals to yield the corresponding ethers^10,11^ (Supplementary Fig. 44). Interestingly, sporopollenin pretreated with TMDS/Cu(OTf)_2_ becomes completely inert to ethyl mercaptan degradation (Fig. 2A), consistent with the hypothesis that sporopollenin contains prevalent thioacidolysis-liable acetals, which are converted to ethers by TMDS/Cu(OTf)_2_ pretreatment. To further validate this hypothesis, we conducted thioacidolysis experiments on two acetal-containing model compounds, 1,1-dimethoxydodecane (**XIII**) and polyvinyl butyral (PVB). Both experiments produced degradation products featuring diethylsulfanyl moieties exactly as predicted, reminiscent of those derived from sporopollenin thioacidolysis (Supplementary Figs. 4, 30, 31).

**Fig. 4.**
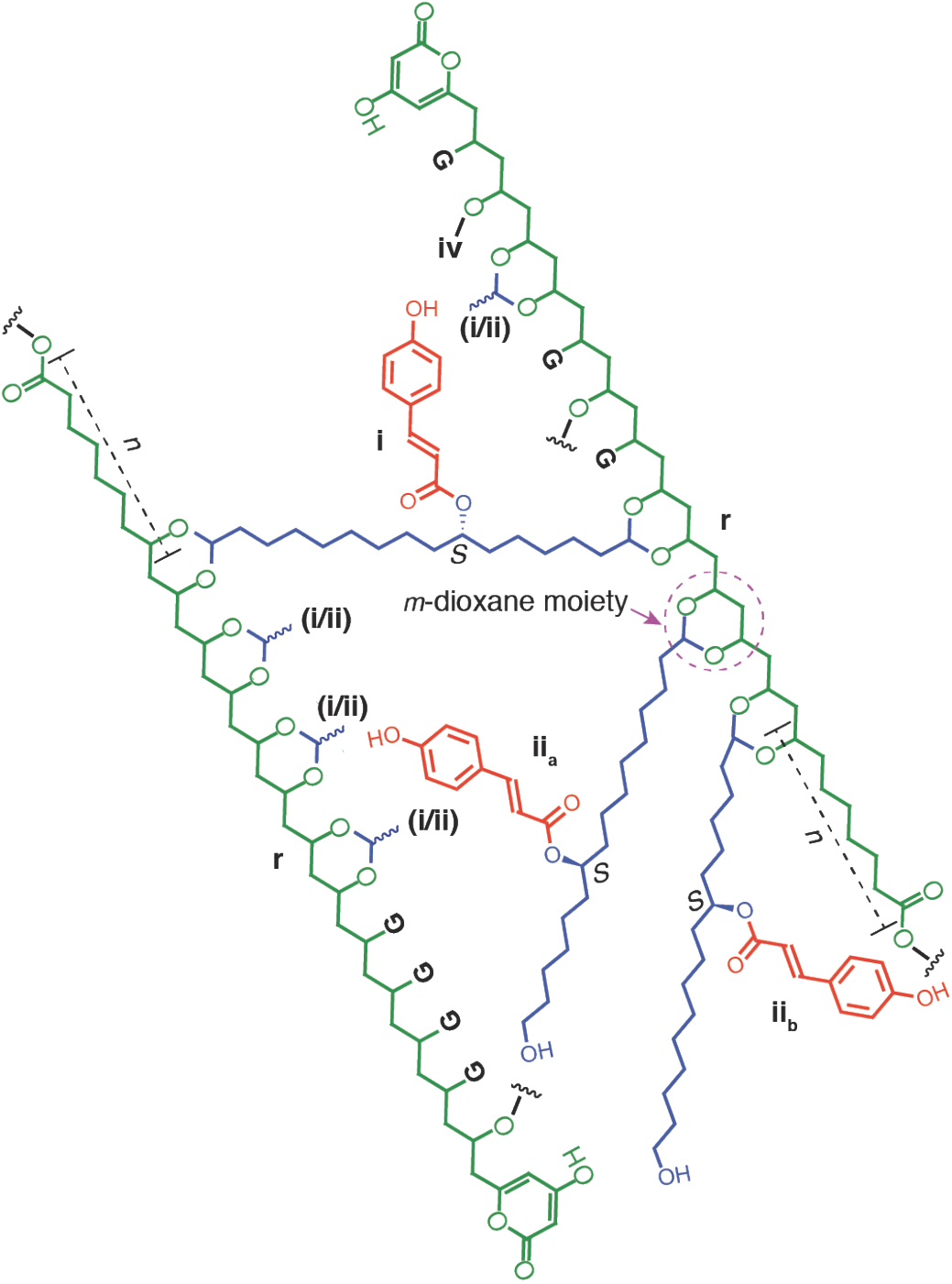
Structural model of pine sporopollenin highlighting the major units and linkages elucidated through this study. In this “averaged” representation of pine sporopollenin, each polyketide-derived PVA-like unit (r) contains 38 carbons and is flanked at two ends by an *ɑ*-pyrone and an ester, respectively. To reflect the “averaged” SSMNR quantitation of different carbon features, r is drawn as derived from a caprylic acid precursor (*n*=8) with 15 cycles of polyketide chain extension, although in a heterogeneous sporopollenin polymer, *n* could also be other even numbers representing other fatty acids employed as polyketide precursors for sporopollenin biosynthesis. The length of the PVA portion of r may also vary in a heterogenous sporopollenin polymer. A fraction of the *ɑ*-pyrone moiety may be substituted by an ester moiety, as *ɑ*-pyrone can be linearized through hydrolysis to give β-keto acid, capable of engaging in ester formation. Each r unit is drawn to form four *m*-dioxane moieties featuring an acetal to crosslink with the 7-*O*-*p*-coumaroylated C16 aliphatic units (i/ii). The ratio of i:ii is approximately 6∼7:1. Glycerol-like moieties (G) and small amount of naringenin (vi) are ether-linked to the hydroxy groups of r. Although the interunit crosslinking is drawn in two dimensions, higher-dimensional cross-linking is likely the case in natural sporopollenin polymer.

Thioacidolysis of pine sporopollenin consistently yields a residue (**R**), accounting for 45±3% (w/w) of the starting sporopollenin material. Like sporopollenin, **R** is recalcitrant to chemical modification, degradation, and dissolution by a wide variety of reagents (Supplementary Note 3). Nonetheless, refluxing **R** with acetic anhydride over 72 hours yielded acetylated **R** (**R-Ac**), which could be dissolved in DMSO at appreciable concentration. We surmise that thioacidolysis cleavage of sporopollenin yields free hydroxy groups in **R**, which are acylated by acetic anhydride to produce **R-Ac** with higher solubility in DMSO than **R**, allowing a set of ^1^H, HSQC and HMBC spectra to be recorded and analyzed (Supplementary Fig. 39). **R-Ac** resembles polyvinyl acetate (PVAc), and is composed of polyhydroxylated aliphatic units interlinked through ester bonds, although the exact length of each individual polyhydroxylated aliphatic unit of **R** could not be determined due to its non-degradable nature.

Magic-angle-spinning (MAS) solid-state NMR (SSNMR) spectroscopy is well suited to the structure determination of insoluble biopolymers such as plant cell walls^12– 15^, sporopollenin^16^, and other biomaterials^17,18^. We measured the ^13^C NMR spectra of pine sporopollenin using the multi-cross-polarization (CP) experiment^19^, which not only enhances the sensitivity of ^13^C in natural abundance (1.1%) but also gives quantitative intensities that reflect the relative number of different types of carbons (Supplementary Fig. 45 and Table S1). At a high magnetic field of 18.8 Tesla, where spectral resolution is optimal, pine sporopollenin exhibits a 103-ppm ^13^C signal indicative of acetal carbons (Fig. 3A) and no intensities in the 200-210-ppm range that is expected for aldehydes, supporting the hypothesis that **I** and **II_(a+b)_** are derived from thioacidolysis cleavage of acetal linkages but not aldehydes. SSNMR spectrum of **R** (Supplementary Fig. 46) shows that thioacidolysis suppresses the 103-ppm acetal ^13^C peak, further supporting acetal as the major thioacidolysis-labile linkage in sporopollenin. Consistently, NaBH_4_ treatment has no impact on the SSNMR spectrum, while TMDS/Cu(OTf)_2_ treatment greatly reduces the 103-ppm peak intensity, confirming acetal as a major inter-unit linkage in pine sporopollenin.

**Fig. 3.**
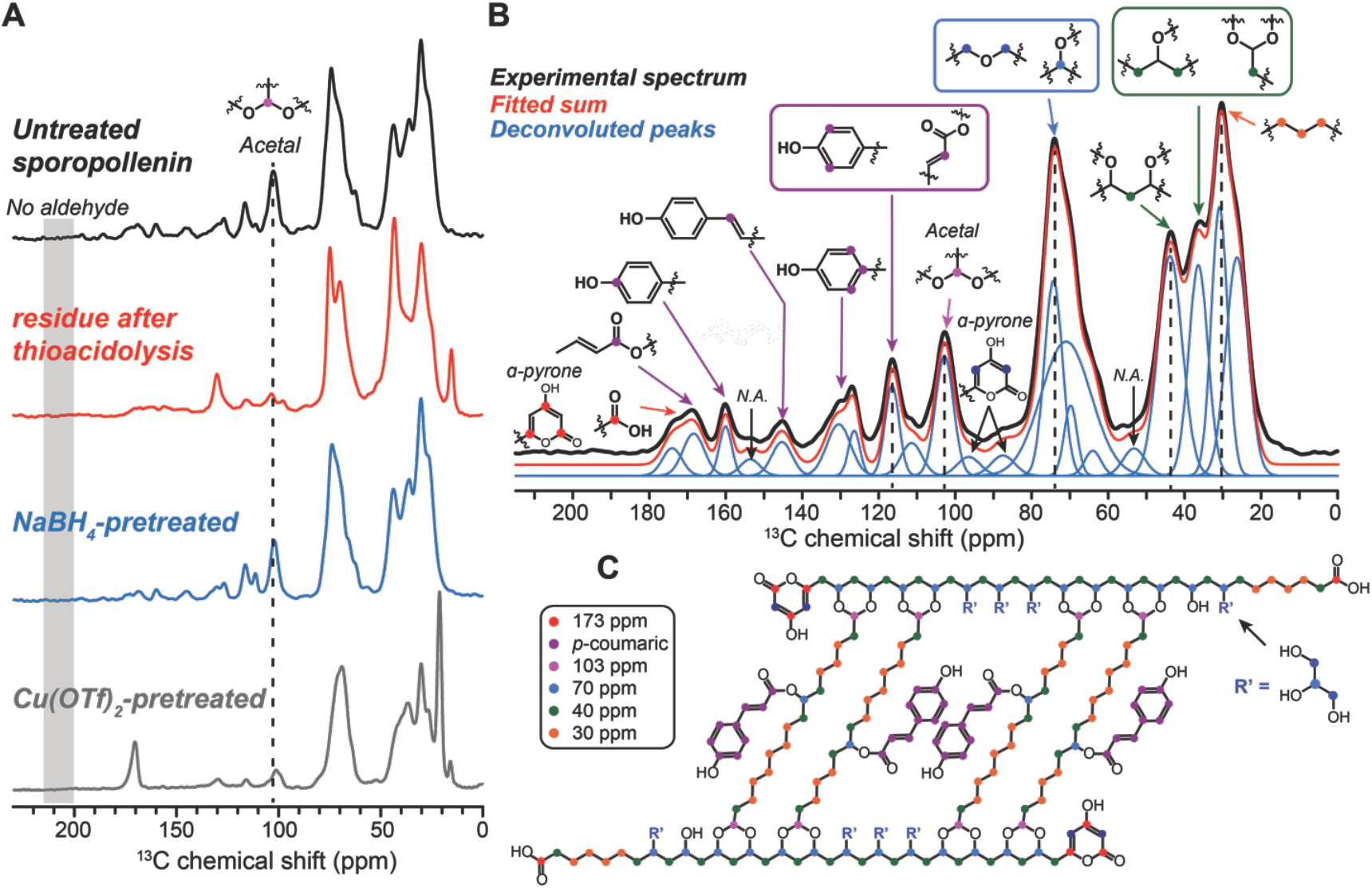
^13^C solid-state NMR spectra of untreated and treated pine sporopollenin for structure determination. **A**, ^13^C multi-CP spectra of untreated sporopollenin, residue (**R**) after thioacidolysis, residue after NaBH_4_ treatment, and residue after TMDS/Cu(OTf)_2_ treatment. All spectra except for **R** were measured on an 800 MHz NMR spectrometer under 14 kHz MAS. **B**, Multi-CP ^13^C spectrum of untreated sporopollenin measured on the 400 MHz NMR under 13 kHz MAS. Spectral deconvolution yielded the relative number of carbons for each assigned peak. **C**, Average chemical structure of pine sporopollenin based on the quantitative ^13^C NMR spectra.

Since carbonyl and aromatic carbons have large chemical shift anisotropies, which spread their intensities over multiple sidebands at high magnetic fields, we measured the ^13^C spectrum of the untreated pine sporopollenin at 9.4 Tesla and 13-kHz MAS to obtain more quantitative intensities for these carbons. The spectrum (Fig. 3B) shows two carboxyl signals at 173 and 168 ppm, six *p*-coumaric acid peaks between 160 and 110 ppm (Table S2), a 103-ppm acetal peak, a strong 74-ppm peak of oxygen-bearing carbons, and three strong aliphatic peaks between 44 and 30 ppm, respectively. The carboxyl signals can be assigned to ester linkages in the major thioacidolysis degradative products **I** and **II_(a+b)_**, respectively (Fig. 1D). Two partially resolved signals at 87 and 96 ppm are assigned to *α*-pyrone, which have been reported to be products of the conserved type III polyketide synthases involved in sporopollenin biosynthesis (e.g. LAP5 and LAP6 in Arabidopsis)^20,21^. To determine whether the 103-ppm, 87-ppm and 60-75 ppm ^13^C peaks result from polysaccharides, we conducted a modified phenol–sulfuric acid reaction^22^ and found no polysaccharide degradative products released (Supplementary Fig. 10). To verify the ^13^C peak assignment, we conducted additional spectral-editing experiments that selectively detect methyl and quaternary ^13^C signals^23^ or CH_2_ signals (Supplementary Fig. 47)^24^. The former confirmed the assignment of the 173-ppm, 168-ppm, 160-ppm, and 126-ppm signals to quaternary carbons in *p*-coumaric acid and *α*-pyrone, while the latter showed that a significant fraction of the 75-26 ppm intensities and the 44-36 ppm intensites are from CH_2_ groups. We deconvoluted the whole pine sporopollenin spectrum to determine the relative number of carbons for the different functional groups. Assuming the resolved 160-ppm peak to represent two carbons from two *p*-coumaric acids, we obtained 98 carbons per repeat unit, among which 18 carbons result from the two *p*-coumaric acid moieties, 5 are acetal carbons, 5 are ester and pyrone carbons, 26 are oxygen-bearing carbons, and 42 are aliphatic carbons (Tables S2 and S3). The remaining carbons for the 153-ppm and 53-ppm chemical shifts are unassigned.

To probe the structure of residue **R** after thioacidolysis, we measured the quantitative ^13^C multi-CP SSNMR spectrum of **R** (Supplementary Fig. 46). The spectrum shows much reduced *p*-coumaroyl ^13^C intensities and the appearance of a new 131-ppm peak attributable to alkenes. Below 80 ppm, the spectrum contains three dominant peaks at 70, 40, and 30 ppm. This pattern resembles the polyvinyl alcohol (PVA) spectrum, which also exhibits dominant signals at 70 and 40 ppm. The 30-ppm peak and the 103-ppm acetal peak suggest that half of the acetal-linked C16 backbones (**i** and **ii)** are retained in **R** but without the *p*-coumaroyl sidechain. The presence of the 88-ppm, 99-ppm, and 160-175 ppm peaks indicate that the *ɑ*-pyrone moiety also remains after thioacidolysis. The two methyl carbons may result from the substitution of glycerol-like moieties by two diethylsulfanyl groups. The deconvoluted spectrum of **R** indicates 71 carbons per repeat unit (Tables S4 and S5), among which 35 are aliphatic, 19 are oxygen-bearing, 6 are ester and pyrone carbons, 4 are olefinic, and 2 are acetals. The remaining five carbons were not assigned (Tables S4).

These degradation product analyses and quantitative ^13^C SSNMR spectra (Supplementary Fig. 48) indicate an average pine sporopollenin structure that consists of two fatty acid-derived PVA-like units (**r**), each flanked at two ends by an *ɑ*-pyrone moiety and an ester group (Fig. 3C and 4). The two PVA-like units are crosslinked by a series of 7-*O*-*p*-coumaroylated C16 aliphatic units (**i**/**ii**) through a distinctive *m*-dioxane moiety featuring an acetal. A fraction (∼15%) of these C16 aliphatic units (**ii_a/b_**) are crosslinked only on one end, with the other end existing as free hydroxy group (Supplementary Fig. 6). The PVA-like units are likely linked through ether bonds with glycerol-like moieties (**G**), and small amount of naringenin (**vi**). The average structure is depicted with an approximate two-fold symmetry and a total of 200 carbons (Fig. 4), but the true structure of natural sporopollenin polymer is likely heterogeneous and involves higher-dimensional inter-unit crosslinking.

Our study indicates that pine sporopollenin is crosslinked through two major types of linkages: ester and acetal. Whereas ester is relatively stable under the acidic condition, acetal can withstand strong alkaline condition^25^. The involvement of both linkages likely accounts for the superior chemical inertness of sporopollenin relative to other plant biopolymers that are crosslinked predominantly via only one type of linkages, e.g. cutin (ester-linked), lignin (ether-linked), cellulose (glycosidic bond-linked) and synthetic polymers such as PVB (acetal-linked). The interweaving hydrophobic and hydrophilic elements stemming from various monomer units also contribute to the poor solubility of sporopollenin in most solvents. From an evolutionary perspective, the identification of *p*-coumaroyl and naringenin moieties as integral components of sporopollenin not only explains sporopollenin’s essential function of protecting gametes against damaging UV light, but also suggests that phenylpropanoid and flavonoid metabolism might have first emerged as part of the evolutionary development towards constructing sporopollenin in early land plants^26^. In addition, the monomer composition and linkage heterogeneity in sporopollenin may render it resistant to enzymatic degradation by microbes and animals, which consequently contributed to the massive accumulation of plant spores from lycophytes during the Carboniferous period and the formation of maceral sporinite as a major component of many coals^27^.

Determining the chemical structure of sporopollenin paves the way towards understanding plant sporopollenin biosynthesis. For example, the identification of structural motifs **i** and **ii** corroborates that CYP703A2 and CYP704B1, two P450s required for pollen exine formation in Arabidopsis and possessing in vitro fatty acid in-chain and ω-hydroxylase activities respectively^28,29^, are involved in installing key hydroxyl groups on C16 fatty acid precursors of **i** and **ii**. In particular, the chiral-specific configuration of the 7-substituted C16 aliphatic units further suggests that the in-chain hydroxylation reaction catalyzed by CYP703A2 likely occurs on a bent C16 substrate with disparate functional groups at each end, presenting only one predominant conformation when entering the CYP703A2 active site. Moreover, the ester-linked *p*- coumaroyl moiety predicts the involvement of an hydroxycinnamoyl transferase yet to be identified. On the other hand, the PVA-like units (**r**) are most likely extended from fatty acid precursors through a type III polyketide biosynthetic mechanism unique to plants^30^. A number of sporopollenin biosynthetic enzymes, including an acyl-CoA ligase, ACOS5 ^31^, type III polyketide synthases LAP5 and LAP6^20^, and two NAD(P)H-dependent reductases, DRL1 and DRL2^21^, previously identified through genetic studies, likely catalyze sequential steps to produce PVA-like monomers of **r**. From an application perspective, knowledge gained from this study on the detailed structure and the biosynthetic logic of sporopollenin natural polymer will also inspire new biomimetic polymer design.

## Acknowledgement

This work was supported by the Pew Scholar Program in the Biomedical Sciences (J.K.W.) and the Searle Scholars Program (J.K.W.). The solid-state NMR part of this work (by P.P. and M.H.) was supported by the Center for Lignocellulose Structure and Formation, an Energy Frontier Research Center funded by the U.S. Department of Energy, Office of Science, Basic Energy Sciences under Award # DE-SC0001090.

## Author Contributions

F.S.L. and J.K.W. designed the research. F.S.L. developed thioacidolysis and pretreatment methods for studying sporopollenin, and carried out structural elucidation of all degradative products. P.P. performed SSNMR experiments. J.J. performed EM imaging. F.S.L, P.P., M.H. and J.K.W. interpreted the results and wrote the paper.

**Supplementary Information for The molecular structure of plant sporopollenin**

## Supplementary Notes

### Supplementary Note 1

#### Prior knowledge regarding the chemical composition of plant sporopollenin

Various attempts have been made to chemically degrade sporopollenin from different species in order to obtain monomers/oligomers. Unlike nucleic acids, proteins or polysaccharides, which contain uniform inter-unit linkages susceptible to enzymatic or chemical hydrolysis, sporopollenin is enriched with multiple, unknown carbon-carbon and ether/ester cross-links which partially explain its extremely resistant characteristics. Conventional techniques can only determine limited information of chemical composition while harsh protocols (ozonolysis, potash fusion, nitrobenzene oxidation, aluminium iodide attack, etc.) destroy linkage information and may produce artefacts. Shaw reported that *P. silvestris* sporopollenin gave 7-hydroxyhexadecanedioic acid and *p*-coumaric acid by alkaline hydrolysis^1^. The *p*-coumaric acid was also found as the main degradative product in Osthoff^2^ and Wehling’s^3^ work when *P. rigida* sporopollenin was saponified and treated with aluminium iodide, respectively. In tracing experiments, the aromatic systems were quite well defined in sporopollenin structure. Gubatz^4^ discovered that the whole carbon skeleton of phenylalanine was incorporated at substantial rates into the sporopollenin of *Tulipa* and he assumed that the phenylpropane unit, such as *p*-coumaric acid, formed the actual monomer. Oleic and stearic acids were also observed to be significantly incorporated into sporopollenin, implicating aliphatic and fatty acid metabolism^5,6^. Hot 2-aminoethanol^7^ was claimed to “dissolve” sporopollenin of *Typha Angustifolia* which made whole-polymer spectroscopic analysis possible, but its poor solubility allowed only ^1^H NMR with restriction to the aromatic region. Moreover, piperidine^8^ turned out to be a more suitable solvent for sporopollenin than hot 2-aminoethanol, as it allowed for higher solubility for 2D-NMR investigations. Unfortunately, the HETCOR and COSY spectra of silylated and acetylated *Typha* samples only evidenced the aliphatic polyhydroxy monomers as well as phenolics substituted by -OH groups. Altogether, these experiments yielded no information for the linkages between different units.

### Supplementary Note 2

#### Chemical degradation of pine sporopollenin by various reagents

Before thioacidolysis, various chemical degradative methods were tested to break down pine sporopollenin. After vigorous stirring with 70% nitric acid for 24 hours, *P. rigida* sporopollenin dissolved/degraded and gave a series of aromatic nitro compounds (**VII-X**) which revealed *p*-coumaric acid as a possible main aromatic unit in *P. rigida* sporopollenin. By stirring *P. rigida* sporopollenin with methyl iodide in dry acetone^9^, methylated *P. rigida* sporopollenin produced the corresponding methylated degradative products (**XI-XII**) during nitric acid degradation, this observation undoubtedly suggested that free *p*-OH groups occurred on *p*-coumaroyl units in *P. rigida* sporopollenin (Supplementary Fig. 3, 9, 24-29). These results only provided limited structural information. Several degradative methods previously developed for analyzing plant polymers (ethyl mercaptan/n-dodecyl mercaptan thioacidolysis for lignin^10,11^, benzyl mercaptan thioacidolysis for tannins^12^ and methanolic base hydrolysis for cutins ^13^) were tested, none of them reacted with *P. rigida* sporopollenin to a significant degree except ethyl mercaptan-based thioacidolysis. Ethyl mercaptan-based thioacidolysis degraded about 55±3% of *P. rigida* sporopollenin and HPLC-UV-MS analysis of the soluble fraction of the reaction gave three major and several minor UV-detectable degradation products (Figs. 1B, C and Supplementary Fig. 6).

### Supplementary Note 3

#### Methods been used for studying chemical structure of R

Many efforts (acetylation by acetic chloride, silylation by BSTFA, Hydrolysis by potassium hydroxide, etc) had been devoted to modify/degrade **R**. None of them worked well.

## Methods

### Chemicals and instruments

All chemicals were purchased from Sigma-Aldrich, unless otherwise specified. *P. rigida* pollen was collected in Cape Cod, MA, United States. Solvents for liquid chromatography high-resolution mass spectrometry were Optima^®^ LC-MS grade (Fisher Scientific) or LiChrosolv^®^ LC-MS grade (Millipore). High-resolution mass spectrometry analysis was performed on a Thermo ESI-Q-Exactive Orbitrap MS coupled to a Thermo Ultimate 3000 UHPLC system. Low-resolution mass spectrometry analysis was done on a Thermo ESI-QQQ MS coupled to a Thermo Ultimate 3000 UHPLC system. Prep-HPLC was performed on a Shimadzu Preparative HPLC with LC-20AP pump and SPD-20A UV-VIS detector. Solution NMR spectra were recorded on a Bruker AVANCE-400 NMR spectrometer with a Spectro Spin superconducting magnet in the Massachusetts Institute of Technology, Department of Chemistry Instrumentation Facility (MIT DCIF) and Oxford 600 MHz Magnet, Bruker AVANCE II Console system, equipped with 5mm Prodigy HCN TXI cold probe in the Harvard Medical School, East Quad NMR facility. SSNMR spectra were measured on an 800 MHz (^1^H Larmor frequency) Bruker AVANCE II spectrometer at 18.8 T using a 3.2 mm MAS probe and a 400 MHz (9.4 T) AVANCE III spectrometer using a 4 mm probe. Typical radiofrequency field strengths were 50-62.5 kHz for ^13^C and 50-80 kHz for ^1^H. All ^13^C chemical shifts were externally referenced to the adamantine CH_2_ peak at 38.48 ppm on the tetramethylsilane (TMS) scale. Circular Dichroism (CD) spectrum was performed on a JASCO Model J-1500 Circular Dichroism Spectrometer equipped with a multi-cuvette holder. Ball milling was done on a high energy Retsch PM 100 Ball Mill.

### *P. rigida* sporopollenin preparation

The ball milling technique^14^ combined with a modified conventional protocol ^7^ was used for *P. rigida* sporopollenin preparation. Briefly, *P. rigida* pollen was ball milled to a super fine powder using a high energy Retsch PM 100 Ball Mill and enzymatically hydrolyzed at 37 °C for 16 hours using a mixture of Macerozym R10 and Cellulase Onozuka R10 (Yakult, Nishinomiya, Japan) (1% of each in 0.1 M sodium acetate buffer, pH 4.5). The resulting material was further extracted exhaustively using a series of solvents of increasing polarity, including hexane, chloroform, ethyl acetate, acetone, methanol, and water. After extraction, the remaining material was lyophilized to obtain *P. rigida* sporopollenin, which accounts for ∼ 33% of the starting pollen in dry weight.

### Acetylation and methylation of *P. rigida* sporopollenin

Acetylation of *P. rigida* sporopollenin. According to the method of Lu^14^, *P. rigida* sporopollenin (0.1 g) was suspended in dimethyl sulfoxide (DMSO, 10 ml) and N-methylimidazole (NMI, 5 ml) was added. The solution was stirred for 2 hours, excess acetic anhydride (3 ml) was added, and the mixture was stirred for 2 hours. The brown solution was transferred into cold water (500 ml), and the mixture was allowed to stand overnight in a 4 °C cold room. The precipitate was recovered by filtration and washed with water (50 mL). The final product was then lyophilized.

*P. rigida* sporopollenin methylation. According to the method of Kettley^9^, *P. rigida* sporopollenin (0.1 g) was stirred with acetone (10 mL) and anhydrous K_2_CO_3_ (0.68 g, more than 3 equivalents relative to the loading of sporopollenin hydroxyls) at room temperature for 1 hour. Methyl iodide (0.5 mL, more than 3 equivalents) was then added and the mixture was refluxed under nitrogen for 24 hours. The mixture was cooled and the sporopollenin was collected by filtration and washed with acetone (3 × 20 mL), water (3 × 20 mL), methanol (MeOH, 3 × 20 mL) and dichloromethane (DCM, 3 × 20 mL). The final product was then lyophilized.

### Thioacidolysis of untreated/acetylated/methylated *P. rigida* sporopollenin and model compound 1, 1-dimethoxydodecane, polyvinyl butyral

According to the method of Rolando^10^, untreated/acetylated/methylated *P. rigida* sporopollenin (0.1 g) or model compound 1, 1-dimethoxydodecane (20 mg)/polyvinyl butyral (0.1 g) was reacted in a 100 mL solution containing ethyl mercaptan (10%, v/v) and 0.2 M boron trifluoride etherate (BF_3_OEt_2_) in dioxane at 100 °C for 4 hours with stirring in a Teflon-screw cap flask under an atmosphere of nitrogen. After the reaction mixture cooled, adjusted its pH to 3-4 by 0.4 M NaHCO_3_ then extracted with CHCl_3_ (3 × 100 mL). The combined organic extracts were dried over anhydrous Na_2_SO_4_, evaporated under reduced pressure and analyzed by HPLC-UV-MS [Thermo ESI-QQQ MS coupled to a Thermo Ultimate 3000 UHPLC system, Phenomenex Kinetex^®^ 2.6 μm C_18_ reverse phase 100 Å 150 x 3 mm LC column, LC gradient: solvent A-water (0.1% formic acid), solvent B-acetonitrile (0.1% formic acid), 0-1 minutes 5% B, 1-15 minutes 5-80% B, 15-25 minutes 95% B, 25-30 minutes 5% B, 0.7 mL/min). The residue after thioacidolysis was collected by filtration, washed by water and dried by lyophilization. Thioacidolysis consistently solubilized about 55±3% of *P. rigida* sporopollenin by dry weight.

### Acetylation of *P. rigida* sporopollenin thioacidolysis residue

According to a modified method of Kettley^9^, *P. rigida* sporopollenin thioacidolysis residue (10 mg) and acetic anhydride (2 mL) were refluxed with stirring for 72 hours, The cooled acetylated residue was then recovered by filtration and washed with water (3 × 30 mL). The final product was then lyophilized.

### Untreated/methylated *P. rigida* sporopollenin reacted with 70% nitric acid

Untreated or methylated *P. rigida* sporopollenin (0.1 g) vigorous stirred with 100 mL 70% nitric acid in room temperature for 24h. The reaction mixture was then slowly adjusted (in 0 °C) its pH to 7 by 10 M NaOH, then extracted with ethyl acetate (3 × 100 mL). The combined organic extracts were dried over anhydrous Na_2_SO_4_, evaporated under reduced pressure and analyzed by HPLC-UV-MS [Thermo ESI-QQQ MS coupled to a Thermo Ultimate 3000 UHPLC system, Phenomenex Kinetex^®^ 2.6 μm C_18_ reverse phase 100 Å 150 x 3 mm LC column, LC gradient: solvent A-water (0.1% formic acid), solvent B-acetonitrile (0.1% formic acid), 0-1 minutes 5% B, 1-15 minutes 5-80% B, 15-25 minutes 95% B, 25-30 minutes 5% B, 0.7 mL/min). The degradative products were separated by Sephadex LH20 eluted with CHCl_3_/MeOH (1:1). Pure compounds were dried *in vacuo*, subjected to NMR analysis in DMSO-*d*_*6*_.

### *P. rigida* sporopollenin treated with modified phenol–sulfuric acid reaction

According to a modified method of Rasouli^15^, *P. rigida* sporopollenin or cellulose (0.1 g) vigorous stirred with 100 mL 80% sulfuric acid in 100 °C for 30 minutes. The reaction mixture pH was then slowly adjusted (at 0°C) to 7 by addition of 10 M NaOH, then extracted with ethyl acetate (3 × 100 mL). The combined organic extracts were dried over anhydrous Na_2_SO_4_, evaporated under reduced pressure and analyzed by HPLC-UV-MS [Thermo ESI-QQQ MS coupled to a Thermo Ultimate 3000 UHPLC system, Phenomenex Kinetex^®^ 2.6 μm C_18_ reverse phase 100 Å 150 x 3 mm LC column, LC gradient: solvent A-water (0.1% formic acid), solvent B-acetonitrile (0.1% formic acid), 0-1 minutes 5% B, 1-15 minutes 5-80% B, 15-25 minutes 95% B, 25-30 minutes 5% B, 0.7 mL/min).

### *P. rigida* sporopollenin treated with TMDS/Cu(OTf)_2_

According to a modified method of Zhang^16^, Cu(OTf)_2_ (74 mg) and 1,1,3,3-tetramethyldisiloxane (TMDS; 1 mL) were added to a suspension of *P. rigida* sporopollenin (0.1 g) in CHCl_3_ (15 mL) at room temperature. After stirring overnight, the mixture was diluted with CHCl_3_, washed with HCl aq. and water. The residue (PM_Cu) was then recovered by filtration and lyophilization. In the subsequent thioacidolysis treatment of PM_Cu, compounds **I**, **II** were not detected in the HPLC-UV-MS chromatogram and it produced an almost undegraded residue (PM_Cu_Thio_Residue). If less Cu(OTf)_2_ (7.4 mg) and TMDS (0.1 mL) were added in the TMDS/Cu(OTf)_2_ reaction, the corresponding residue PM_Cu_Less produced compound **I**, **II** in its HPLC-UV-MS chromatogram in the subsequent thioacidolysis treatment. These results suggested that TMDS/Cu(OTf)_2_ converted acetal to ether interlinkages and resulted in a modified sporopollenin that was unsusceptible to thioacidolysis.

### Purification and characterization of thioacidolysis degradative products from *P. rigida* sporopollenin

The combined organic extracts after thioacidolysis treatment of untreated/acetylated/methylated *P. rigida* sporopollenin were separated by Sephadex LH20 eluted with CHCl_3_/MeOH (1:1). Fractions were collected and analyzed by HPLC-UV-MS [Thermo ESI-QQQ MS coupled to a Thermo Ultimate 3000 UHPLC system, Phenomenex Kinetex^®^ 2.6 μm C_18_ reverse phase 100 Å 150 x 3 mm LC column, LC gradient: solvent A-water (0.1% formic acid), solvent B-acetonitrile (0.1% formic acid), 0-1 minutes 5% B, 1-15 minutes 5-80% B, 15-25 minutes 95% B, 25-30 minutes 5% B, 0.7 mL/min). Further purification was performed on Prep-HPLC (Kinetex 5μ C_18_ 100A, 150 × 21.2 mm column, solvent A-water, solvent B-acetonitrile, 0-30 minutes 40-80% B, 30-35 minutes 80-95% B, 35-65 minutes 95% B, 10 mL/min). Pure compounds were dried *in vacuo* and subjected to NMR analysis in CDCl_3_.

Absolute configuration identification of compound **I** Potassium hydroxide hydrolysis of compound **I**. Compound **I** (10 mg) was refluxed in 1M KOH for 1h. After the reaction mixture cooled, its pH was adjusted to 7 by addition of 1 M HCl and then extracted with CHCl_3_ (3 × 10 mL). The combined organic extracts were dried over anhydrous Na_2_SO_4_ and then evaporated under reduced pressure at 40 °C. Further purification was performed on Prep-HPLC (Kinetex 5μ C_18_ 100A, 150 × 21.2 mm column, solvent A-water, solvent B-acetonitrile, 0-30 minutes 40-80% B, 30-35 minutes 80-95% B, 35-65 minutes 95% B, 10 mL/min). Pure compound **I_a_** was dried *in vacuo*, subjected to NMR analysis in CDCl_3_. Compound **I** was analyzed by chiral HPLC-UV-MS [Thermo ESI-QQQ MS coupled to a Thermo Ultimate 3000 UHPLC system, Lux 3 μ Cellulose-4, 250 × 4.6 mm chiral column, LC gradient: solvent A-water (0.1% formic acid), solvent B-acetonitrile (0.1% formic acid), 0-30 minutes 50-95% B, 30-40 minutes 95% B, 40-42 minutes 50% B, 0.7 mL/min).

Derivatization of compound **I_a_** with (1R,2R)-(-)-2-(2,3-anthracenedicarboximide) cyclohexane carboxylic acids and (1S,2S)-(+)-2-(2,3-anthracenedicarboximide) cyclohexane carboxylic acids. According to a modified method of Ohrui^17^, Compound **I_a_** was esterified with (1R,2R)-(-)-2-(2,3-anthracenedicarboximide) cyclohexane carboxylic acids or (1S,2S)-(+)-2-(2,3-anthracenedicarboximide) cyclohexane carboxylic acids by using 1-ethyl-3- (3-dimethylaminopropyl) carbodiimide hydrochloride (EDC) in the presence of 4-(dimethylamino)pyridine (DMAP) in a mixture of toluene and acetonitrile (1:1, v/v). The resultant mixture was stirred at 40 °C for overnight. The reaction mixture was analyzed by HPLC-UV-MS [Thermo ESI-QQQ MS coupled to a Thermo Ultimate 3000 UHPLC system, Phenomenex Kinetex^®^ 2.6 μm C_18_ reverse phase 100 Å 150 x 3 mm LC column, LC gradient: solvent A-water (0.1% formic acid), solvent B-acetonitrile (0.1% formic acid), 0-1 minutes 5% B, 1-15 minutes 5-80% B, 15-25 minutes 95% B, 25-30 minutes 98% B, 30-35 minutes 5% B, 0.7 mL/min). Further purification was performed on Prep-HPLC (Kinetex 5μ C_18_ 100A, 150 × 21.2 mm column, solvent A-water, solvent B-acetonitrile, 0-30 minutes 40-80% B, 30-35 minutes 80-95% B, 35-65 minutes 95% B, 10 mL/min). Pure **I_a_-*RR*** and **I_a_-*SS*** were dried *in vacuo*, subjected to NMR analysis in CDCl_3_.

### Solid-state NMR experiments

To obtain quantitative ^13^C spectra that reflect the relative numbers of chemically distinct carbons in sporopollenin, we conducted the multiple cross-polarization (multi-CP) experiment ^18^. Compared to direct polarization (DP) ^13^C experiments with long recycle delays, which is the traditional method of quantitative ^13^C NMR, multi-CP achieves two-fold higher sensitivity by repeated polarization transfer from protons. Moreover, multi-CP gives uniform intensities for carbons with different numbers of attached protons by repolarizing the ^1^H magnetization during the ^1^H spin-lattice relaxation periods (t_z_) that alternate with the CP periods. The CP contact times and the t_z_ periods were empirically optimized using the model tripeptide, formyl-MLF, by comparing the multi-CP spectrum with the quantitative DP spectrum measured with a 35 s recycle delay. For the untreated and treated sporopollenin samples, we used five cycles of CP periods where each cycle has a contact time of 330 μs and a t_z_ period of 0.4 s to 1 s. ^13^C multi-CP spectra with gated ^1^H decoupling^19,20^ were measured on the 400 MHz NMR spectrometer under 13 kHz MAS at 296 K. The gate period was 36 μs before and after the 180° pulse, which suppresses the protonated ^13^C signals due to their strong ^13^C-^1^H dipolar couplings. For the untreated pine sporopollenin sample, 7190 scans were acquired for the gated multi-CP spectrum while 10496 scans were acquired for the regular multi-CP spectrum. For the sample after thioacidolysis treatment, 8192 scans were acquired for the gated multi-CP spectrum and 6400 scans were acquired for the regular multi-CP spectrum. The CH_2_ selection experiment^21^ was conducted on the 800 MHz spectrometer under 6 kHz MAS at 296 K. A short CP contact time of 50 μs was used to suppress the signals of methyl and non-protonated carbons. During the CH_2_ selection period, the ^1^H 0° and 90° pulses were constructed from two phase-cycled 45° pulses with an rf field strength of 73.5 kHz, while the ^13^C 101° pulse had an rf field strength of 67.5 kHz. The z-filter, which was designed to further suppress the methyl ^13^C signals by fast T_1_ relaxation, was set to 0.1 μs, since it was found to not affect the spectra of natural-abundance L-Leucine, which was used to optimize the experiment. For leucine, 512 scans were averaged for the CH_2_ selected single-CP spectrum while 8 scans were acquired for the non-selective single-CP spectrum. For the untreated sporopollenin sample, 68,608 scans were acquired for the CH_2_-selected single-CP spectrum and 6144 scans were acquired for the non-selective single-CP spectrum.

### Spectral assignment and deconvolution

^13^C chemical shifts in the MAS NMR spectra were assigned using ChemDraw Professional predictions and literature chemical shifts^20,22,23.^ The multi-CP spectra were deconvoluted into Gaussian peaks using the DMFit software^24^ to extract the relative intensity of each peak. The model used for deconvolution was “Gaus/Lor”, and adjustable parameters were chemical shift, peak amplitude, width, and type of line shape. Carbonyl and aromatic carbons with large chemical shift anisotropies (CSAs) lost part of their intensities to spinning sidebands even at 9.4 T (with a ^13^C Larmor frequency of 100 MHz) under an MAS frequency of 13 kHz. We quantitatively accounted for the fraction of intensities lost to the sidebands by simulating the CSA sideband patterns for these groups using the “CSA mas” model in DMFit. The inputs for the simulations include the MAS frequency, the magnetic field strength, and chemical shift anisotropy parameter and asymmetry parameter, which are well known. This information is included in tables S1 and S3 as centerband to total intensity ratios, and is necessary for determining the accurate carbon counts. Finally, in determining the carbon counts, we also took into account imperfections in the multi-CP experiment, which is reflected by the slightly lower intensities of carbonyl and aromatic carbons (20-25%) relative to the quantitative direct polarization intensities measured on the model compound, formyl-MLF (Tables S1 and S3).

## Supplementary Figures

**Fig. S1.**
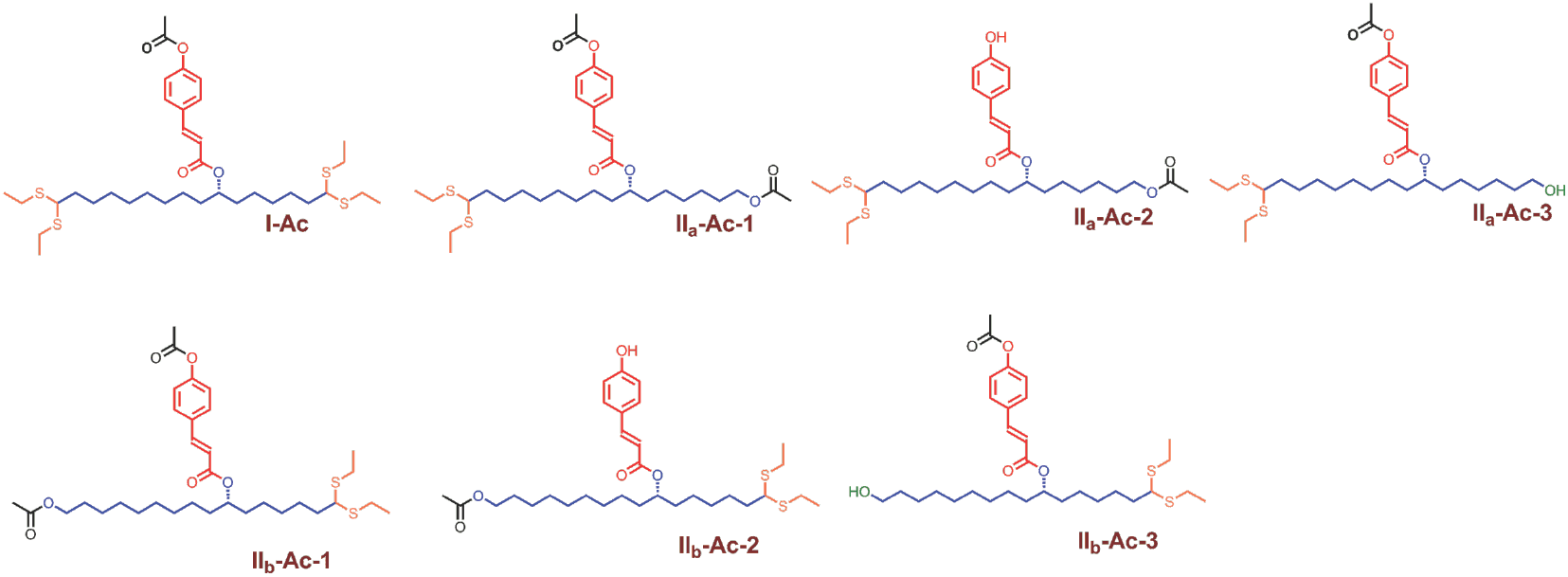
Degradative products of acetylated *P. rigida* sporopollenin thioacidolysis.

**Fig. S2.**
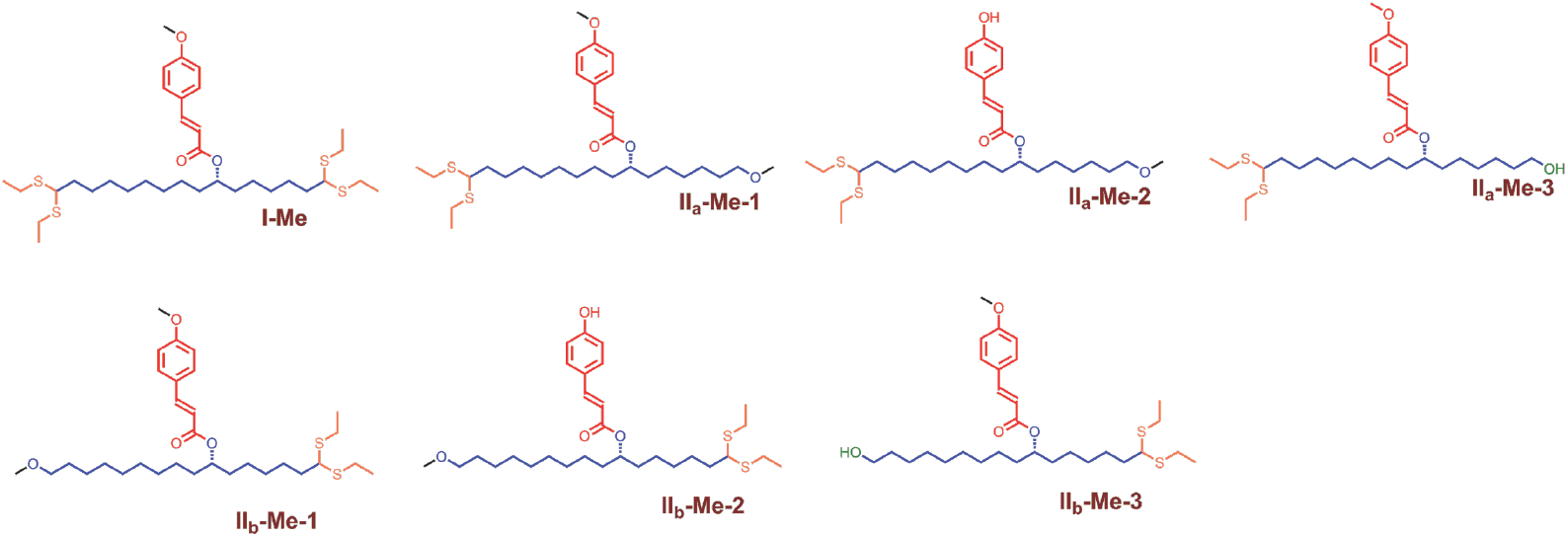
Degradative products of methylated *P. rigida* sporopollenin thioacidolysis.

**Fig. S3.**
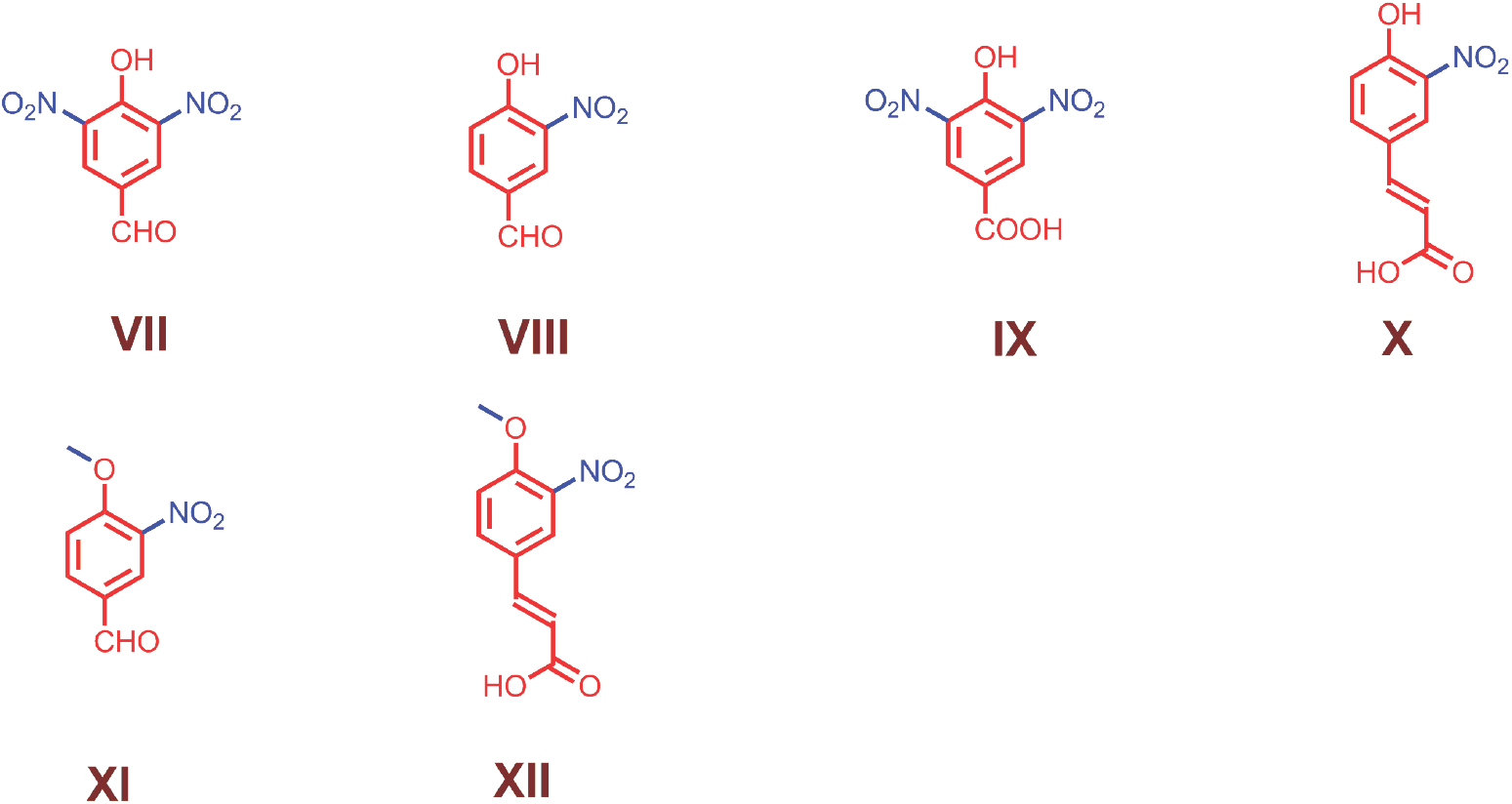
Degradative products of *P. rigida* sporopollenin and methylated *P. rigida* sporopollenin treated with nitric acid.

**Fig. S4.**
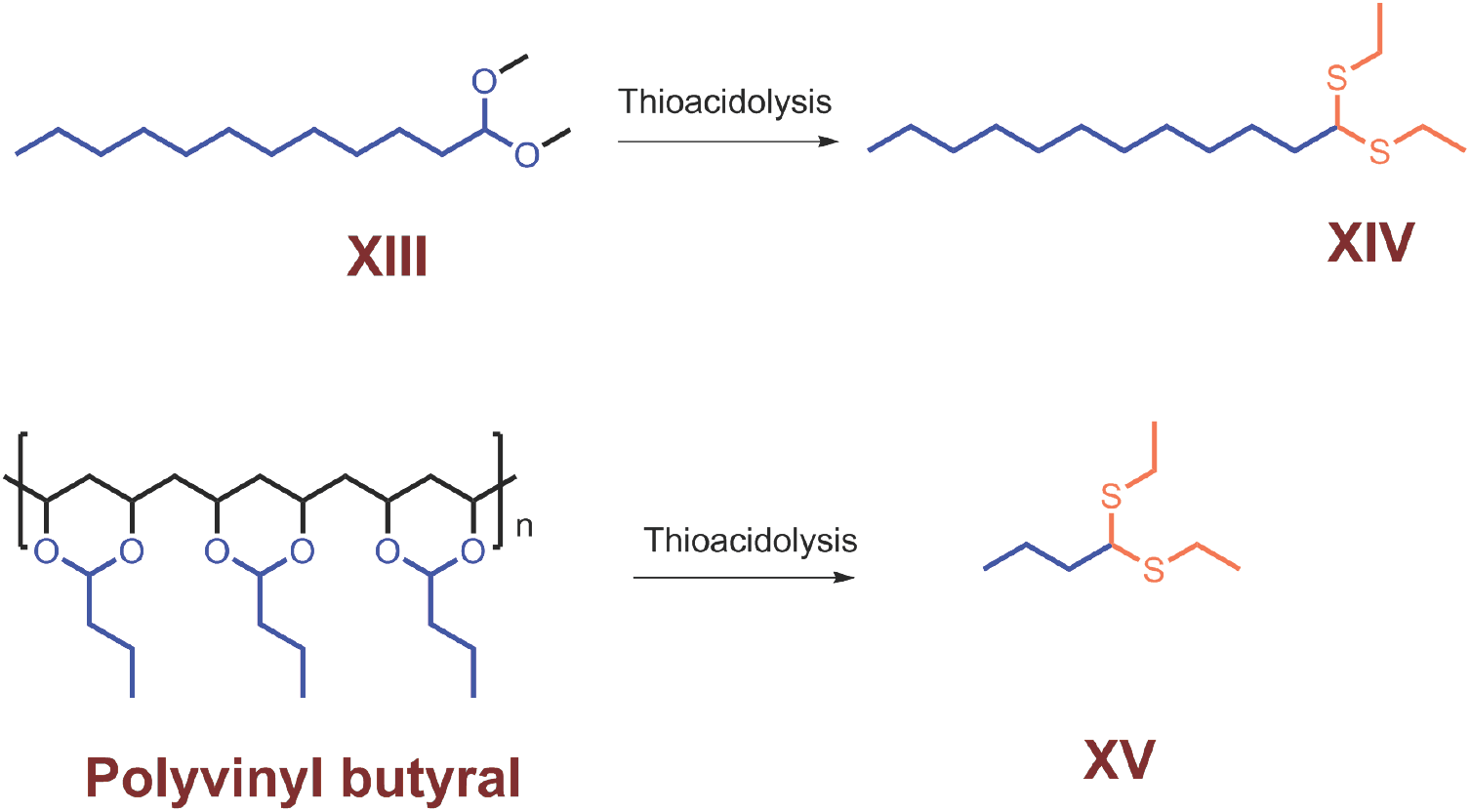
Thioacidolysis of 1, 1-dimethoxydodecane (**XIII**) and Polyvinyl butyral produced **XIV** and **XV** respectively.

**Fig. S5.**
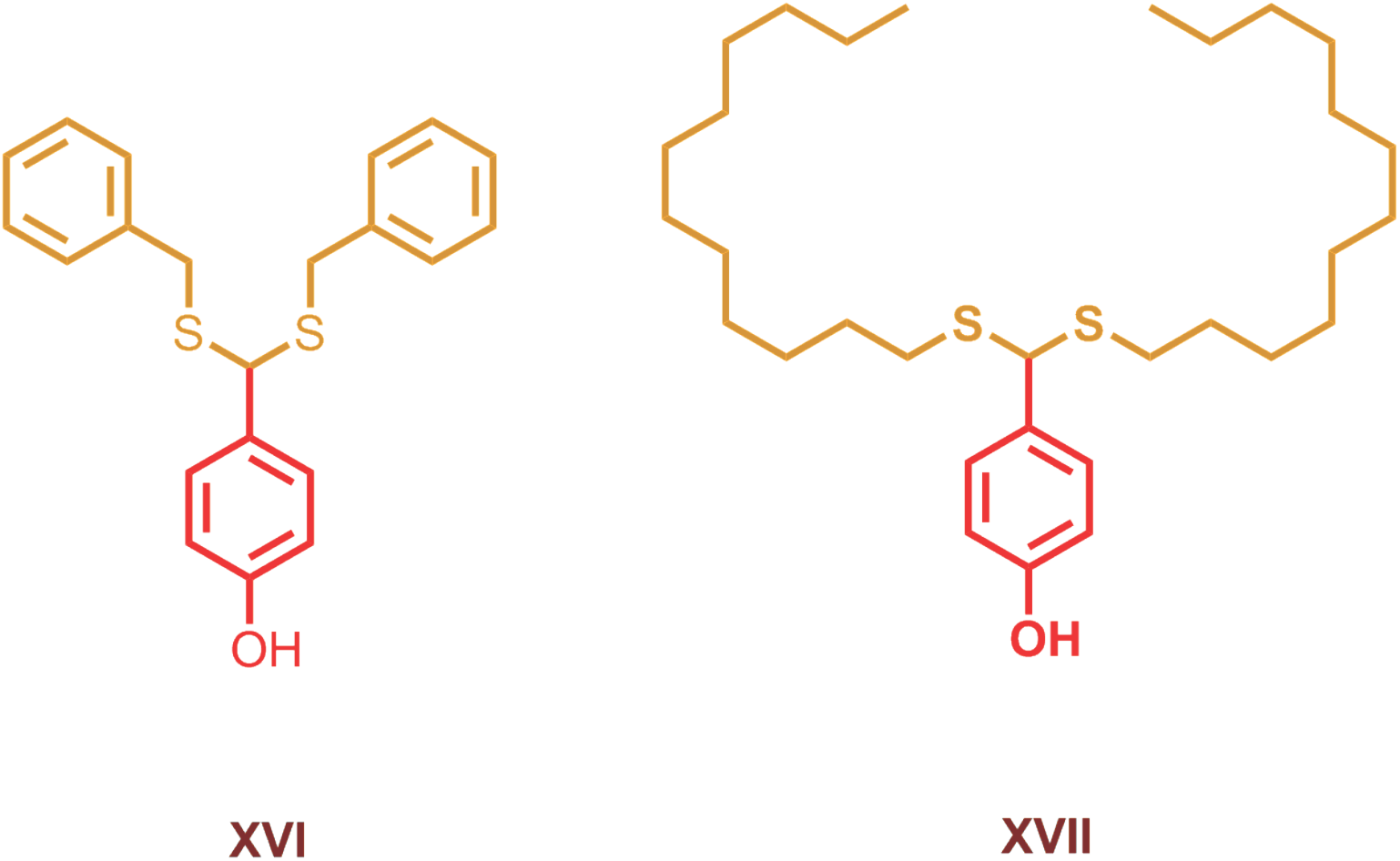
Degradative products of *P. rigida* sporopollenin benzyl mercaptan thioacidolysis (**XVI** and n-dodecyl mercaptan thioacidolysis (**XVII**).

**Fig. S6.**
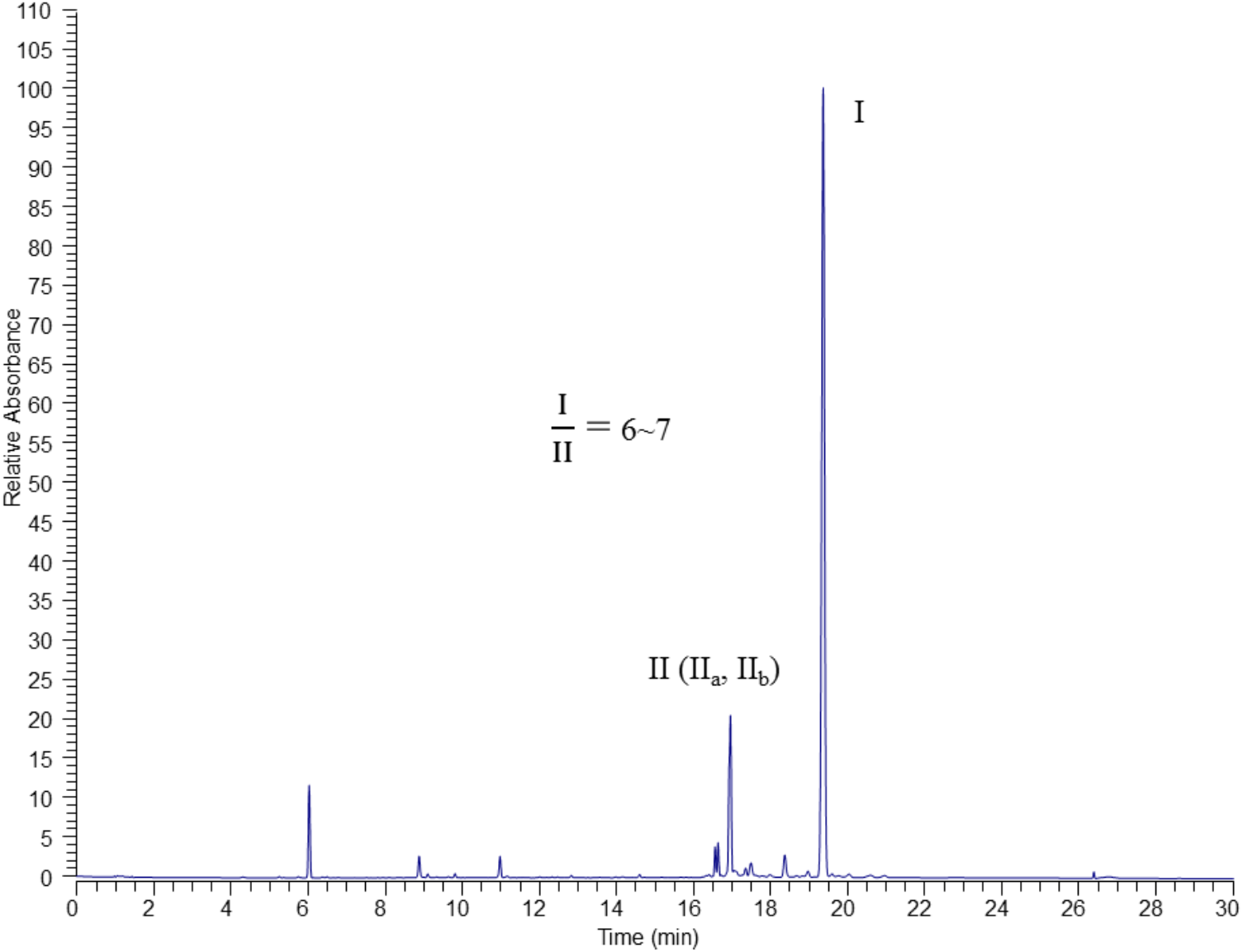
HPLC-UV (310nm) chromatogram of *P. rigida* sporopollenin thioacidolysis, the ratio of peak areas of **I** and **II** is about 6∼7. A subfraction (∼15%) of the C16 aliphatic units (**ii**) are crosslinked only on one end, with the other end existing as free hydroxy group.

**Fig. S7.**
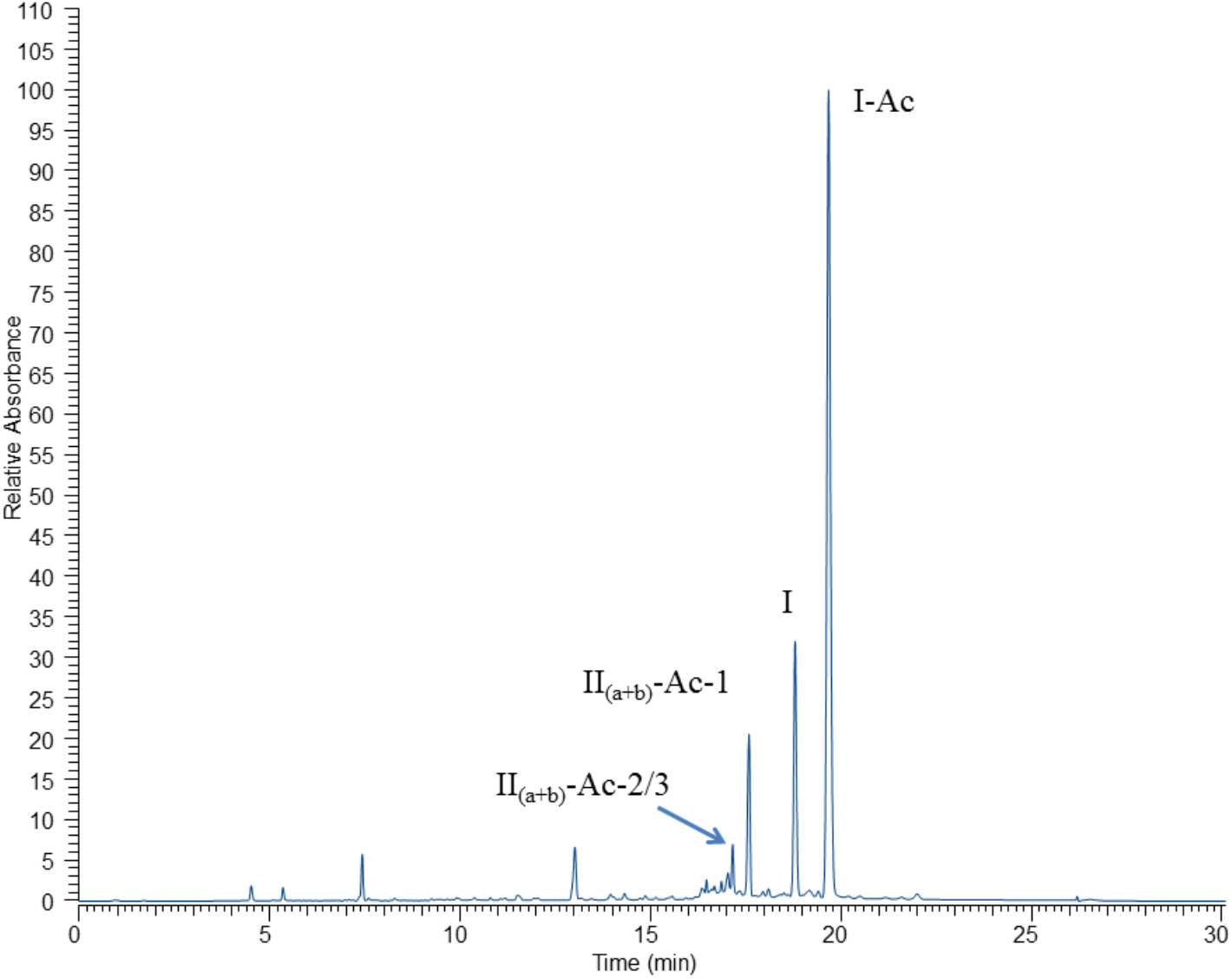
HPLC-UV (280nm) chromatogram of acetylated *P. rigida* sporopollenin thioacidolysis.

**Fig. S8.**
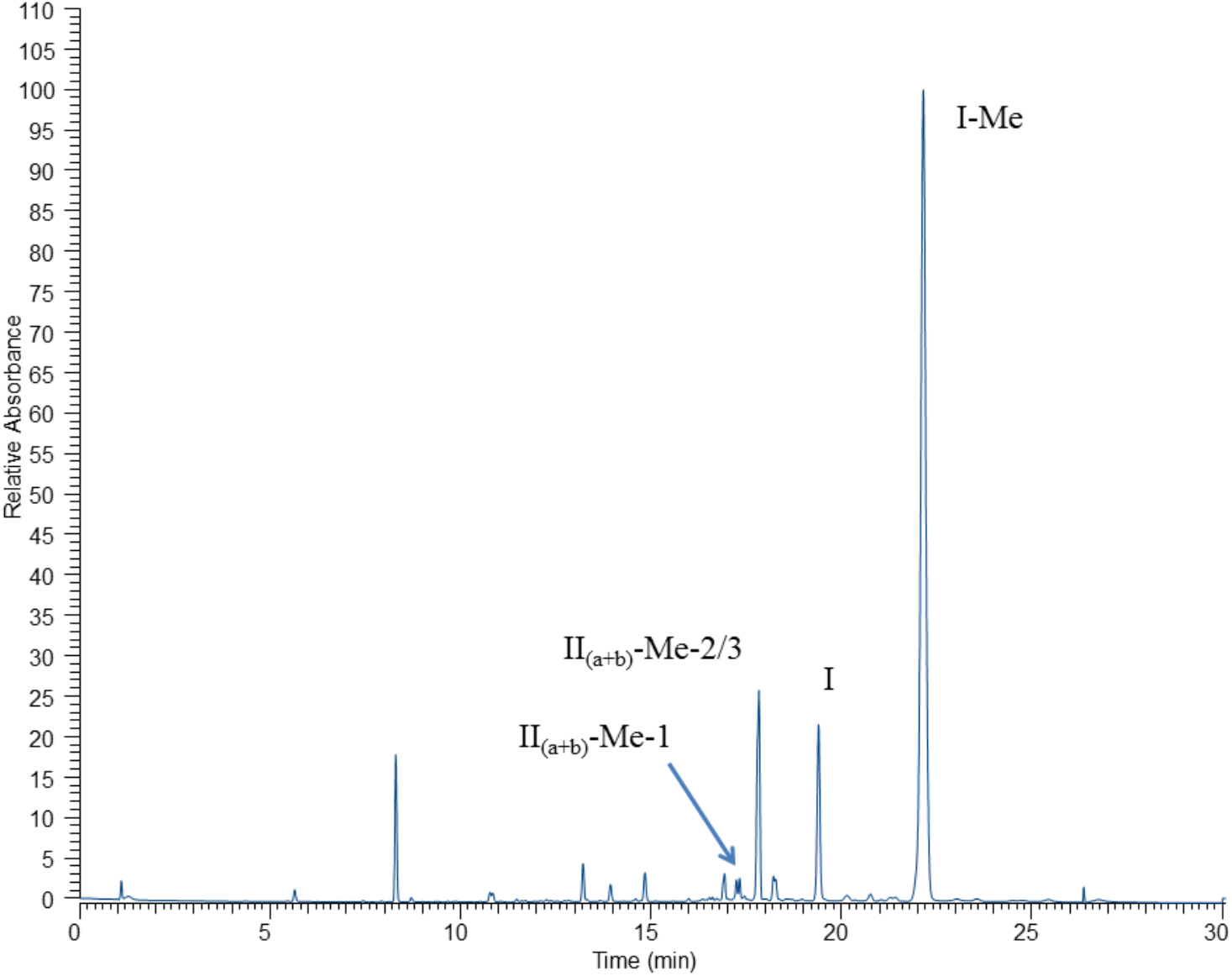
HPLC-UV (310nm) chromatogram of methylated *P. rigida* sporopollenin thioacidolysis.

**Fig. S9.**
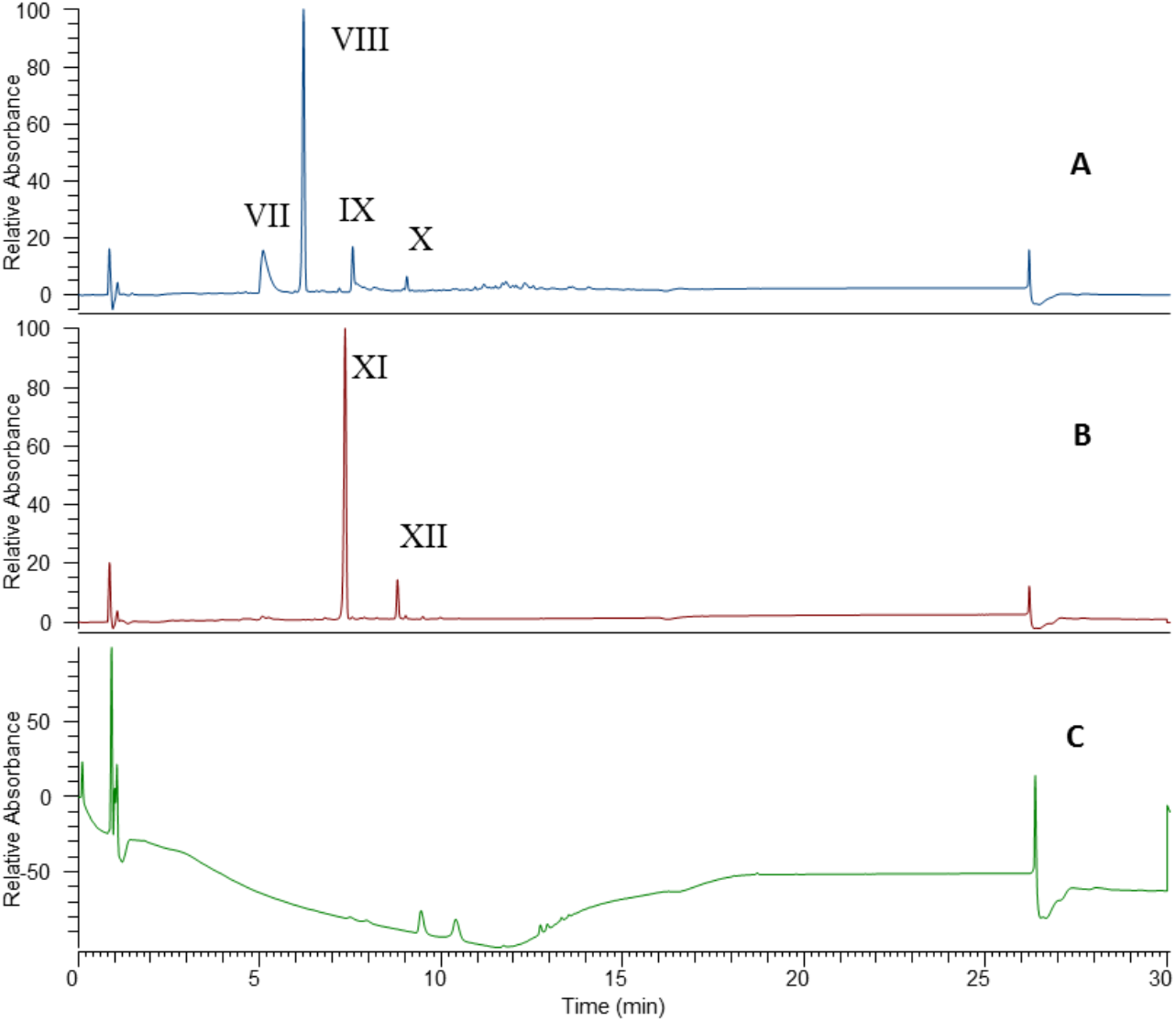
HPLC-UV (PDA) chromatogram of *P. rigida* sporopollenin and methylated *P. rigida* sporopollenin treated with nitric acid. **(A)** Degradative products of *P. rigida* sporopollenin treated with nitric acid, **(B)** Degradative products of methylated *P. rigida* sporopollenin treated with nitric acid, **(C)** Nitric acid treatment of the residue from methylated *P. rigida* sporopollenin thioacidolysis did not produce detectable degradative products.

**Fig. S10.**
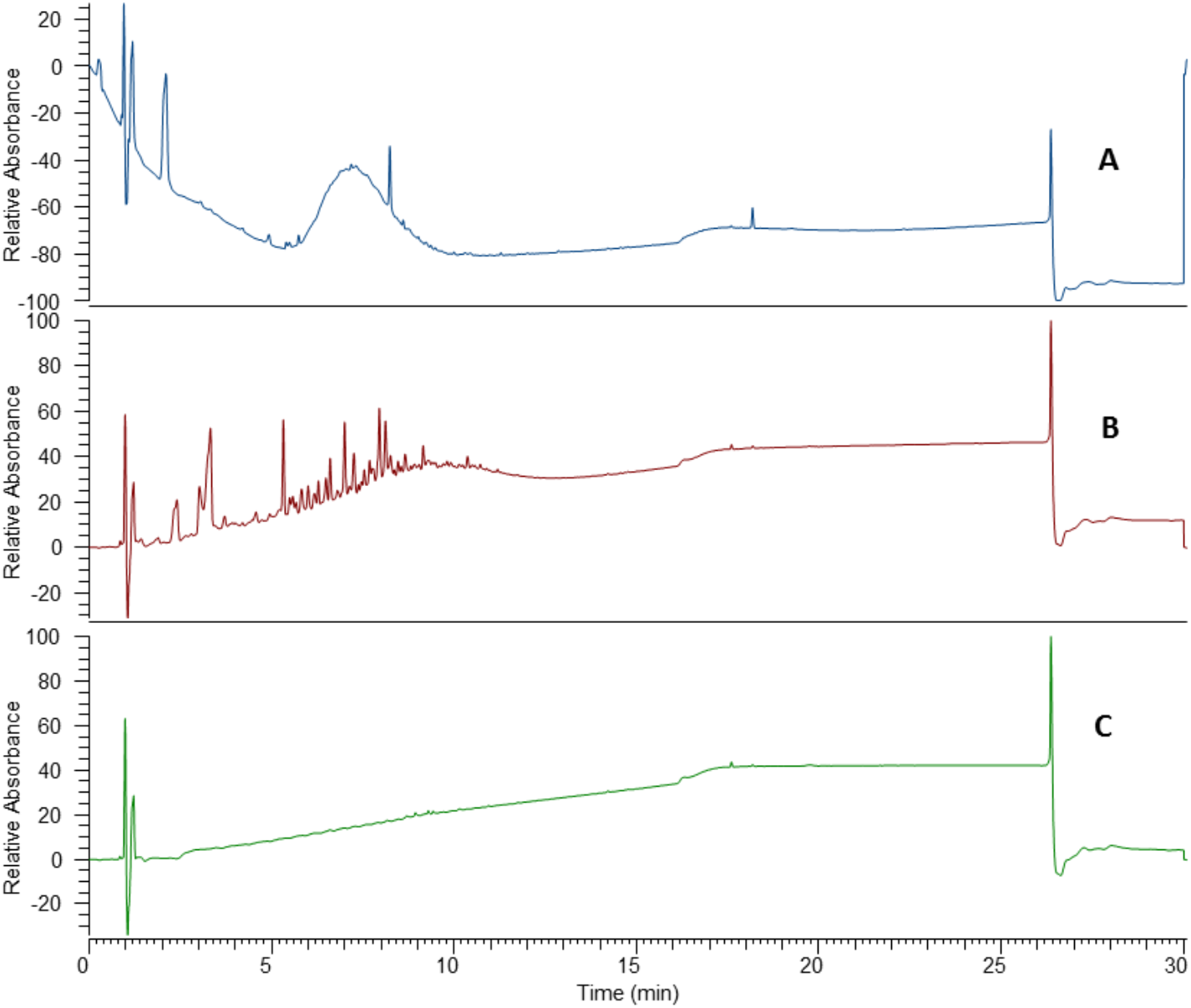
HPLC-UV (PDA) chromatogram of *P. rigida* sporopollenin and cellulose treated with 80% sulfuric acid. **(A)** Degradative products of *P. rigida* sporopollenin treated with 80% sulfuric acid, **(B)** Degradative products of cellulose treated with 80% sulfuric acid **(C)** Blank control chromatogram of 80% sulfuric acid reaction.

**Fig. S11-A.**
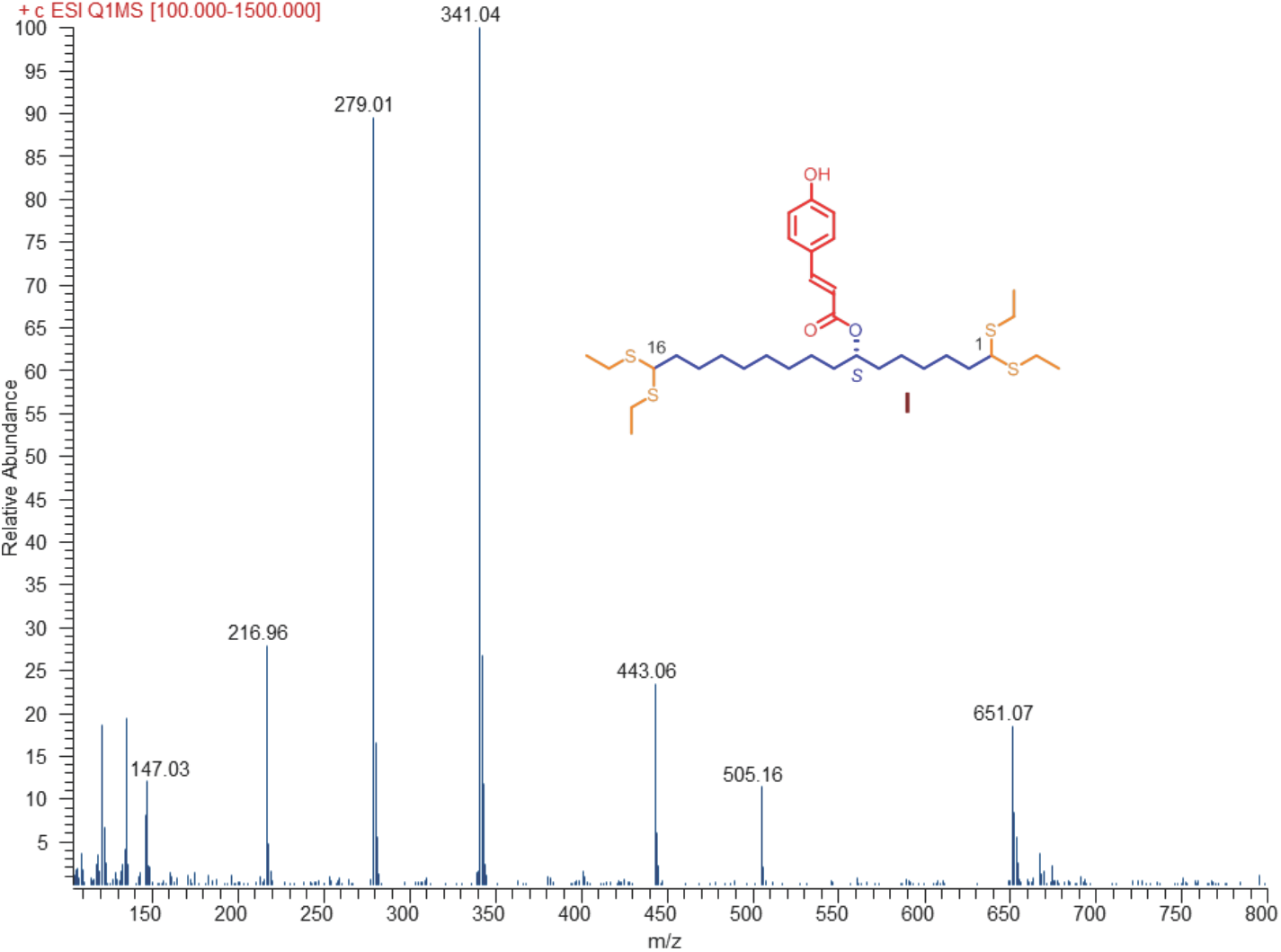
Characterization of compound I, MS/MS spectrum of compound **I**. MS/MS spectrum of compound **I**.

**Fig. S11-B.**
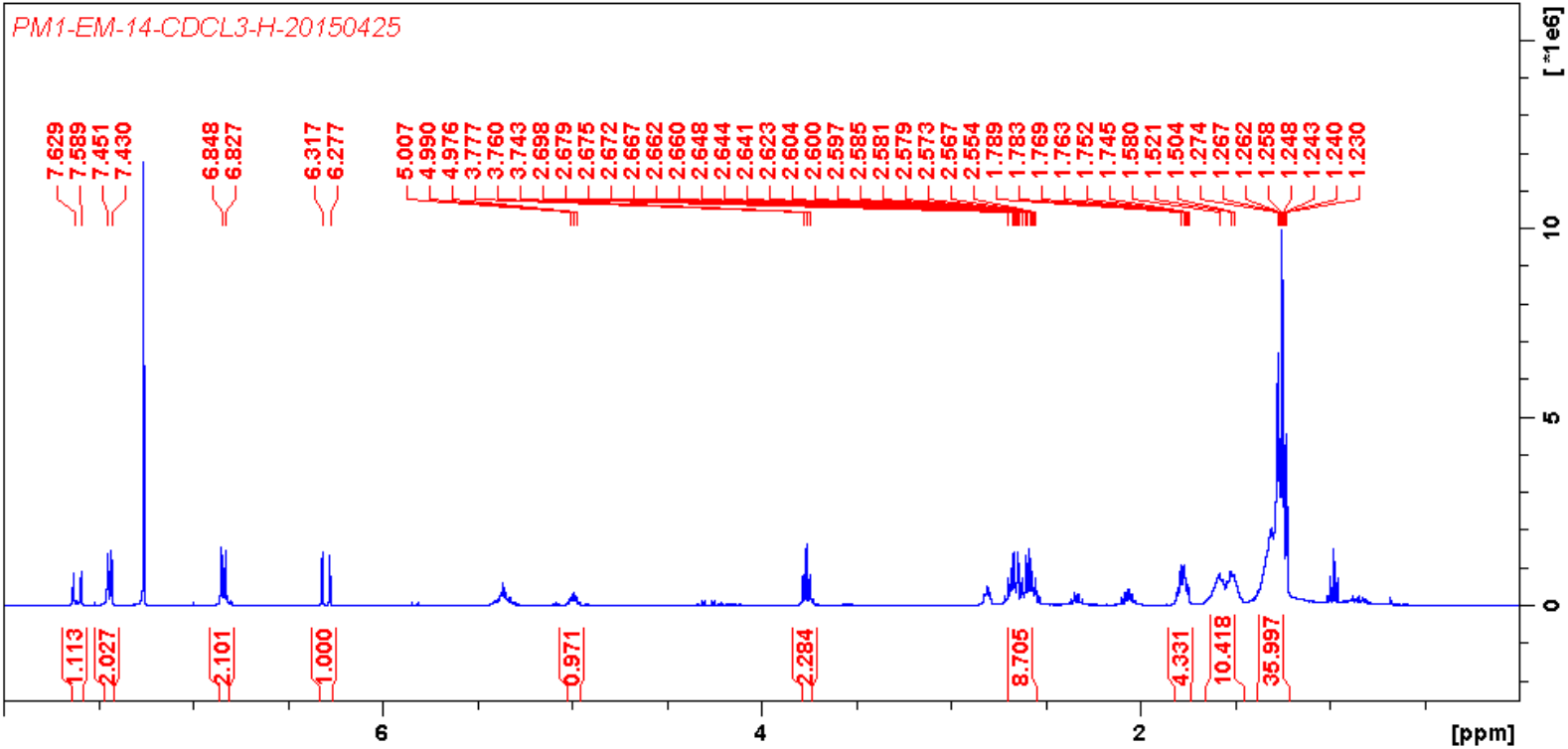
Characterization of compound I, ^1^H NMR (400 MHz, CDCl_3_) spectrum of compound **I**.

**Fig. S11-C.**
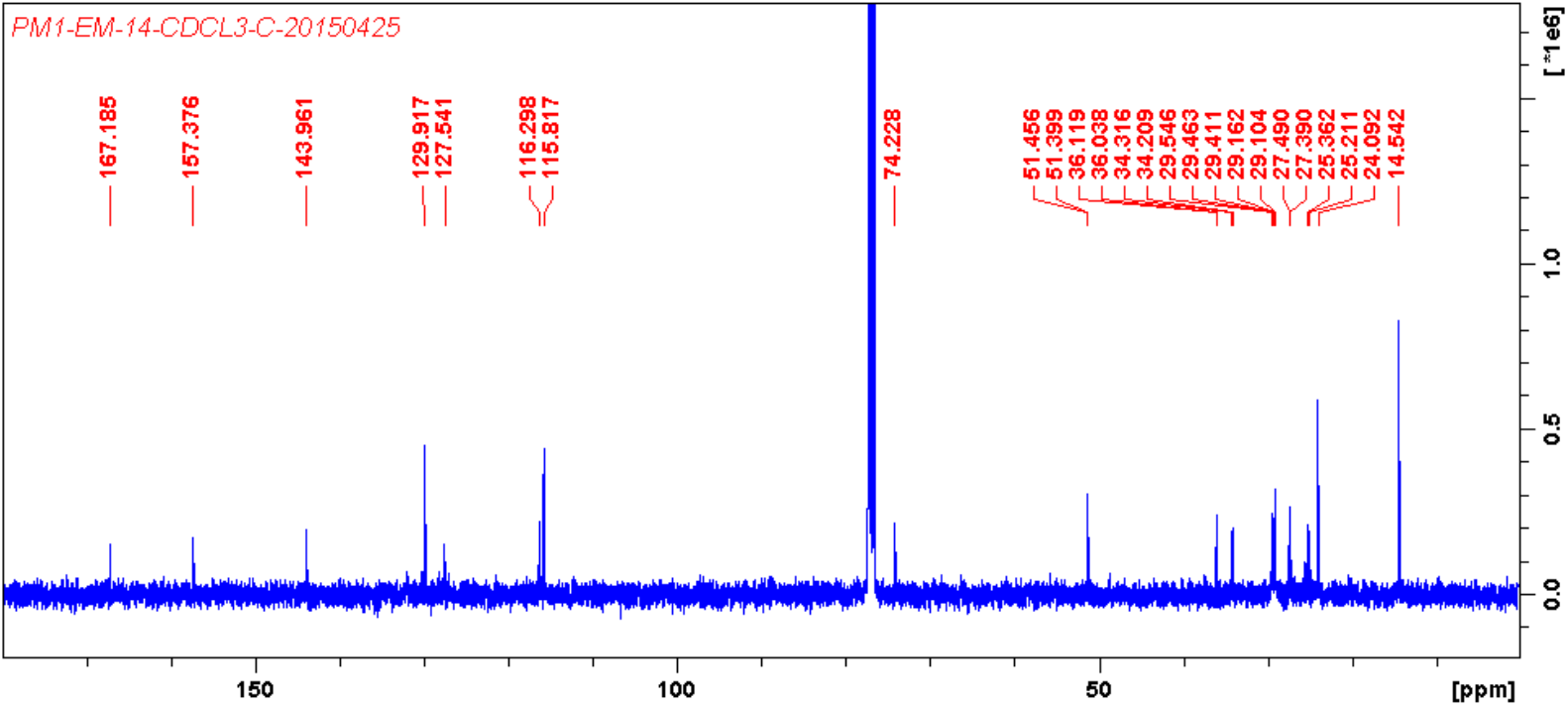
Characterization of compound I, ^13^C NMR (100 MHz, CDCl_3_) spectrum of compound **I**

**Fig. S11-D.**
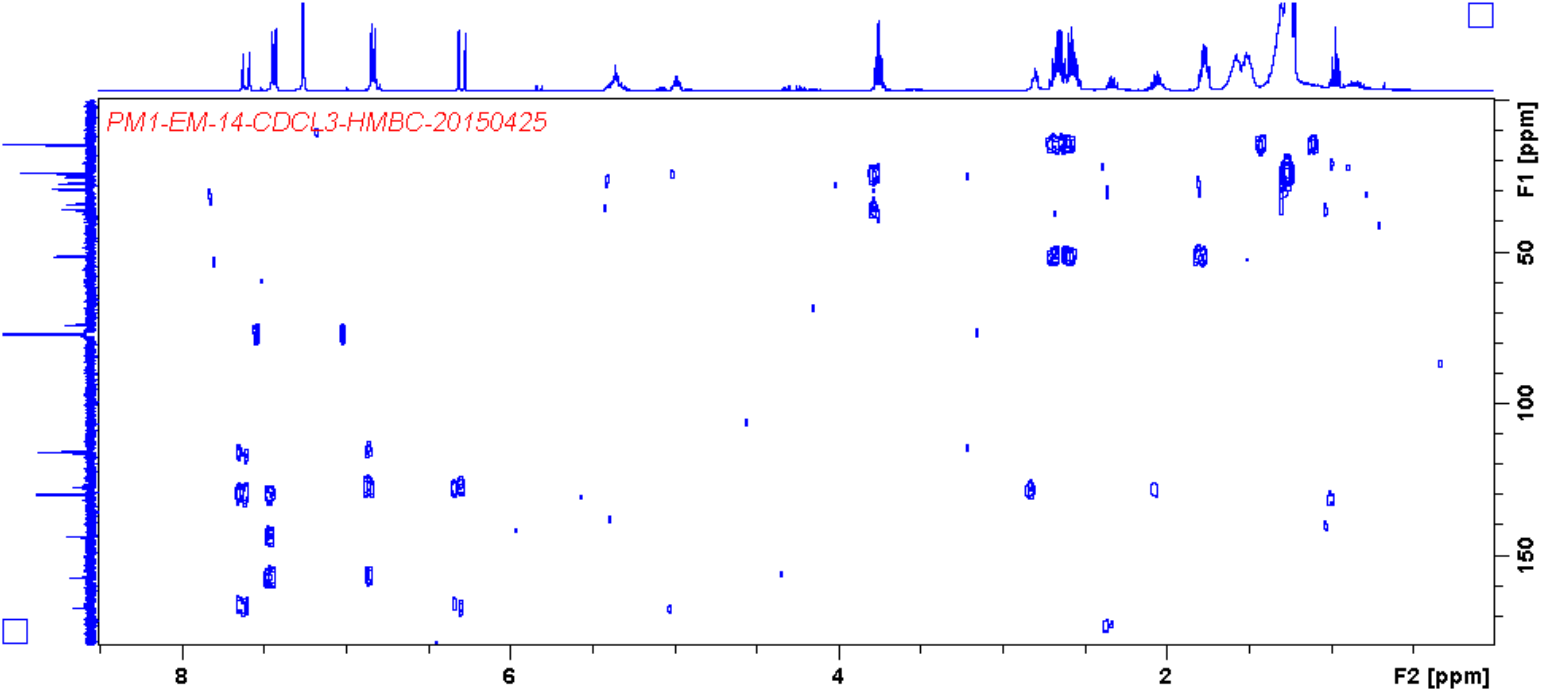
Characterization of compound I, HMBC spectrum of compound **I**.

**Fig. S12-A.**
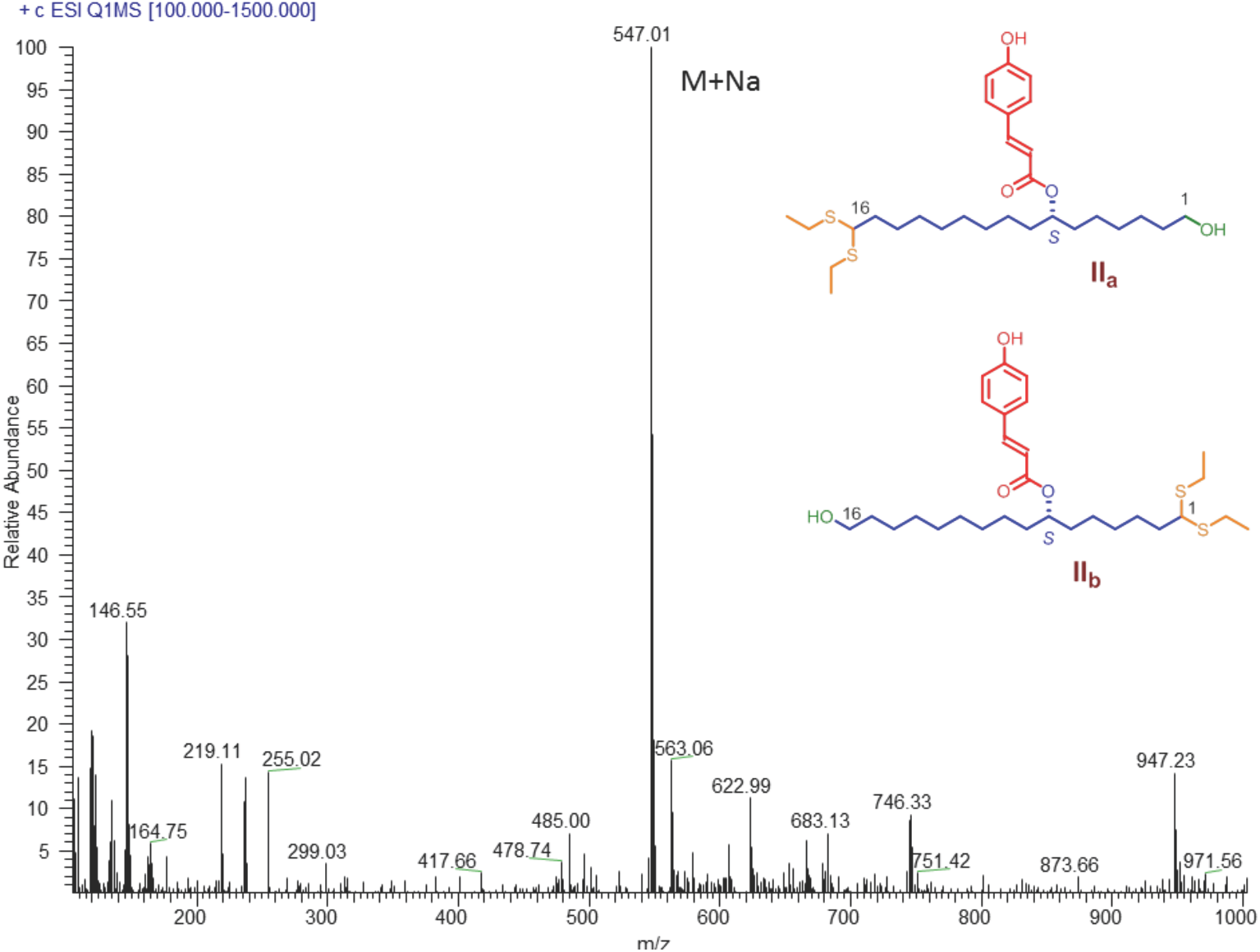
Characterization of compound II, MS/MS spectrum of compound **II**.

**Fig. S12-B.**
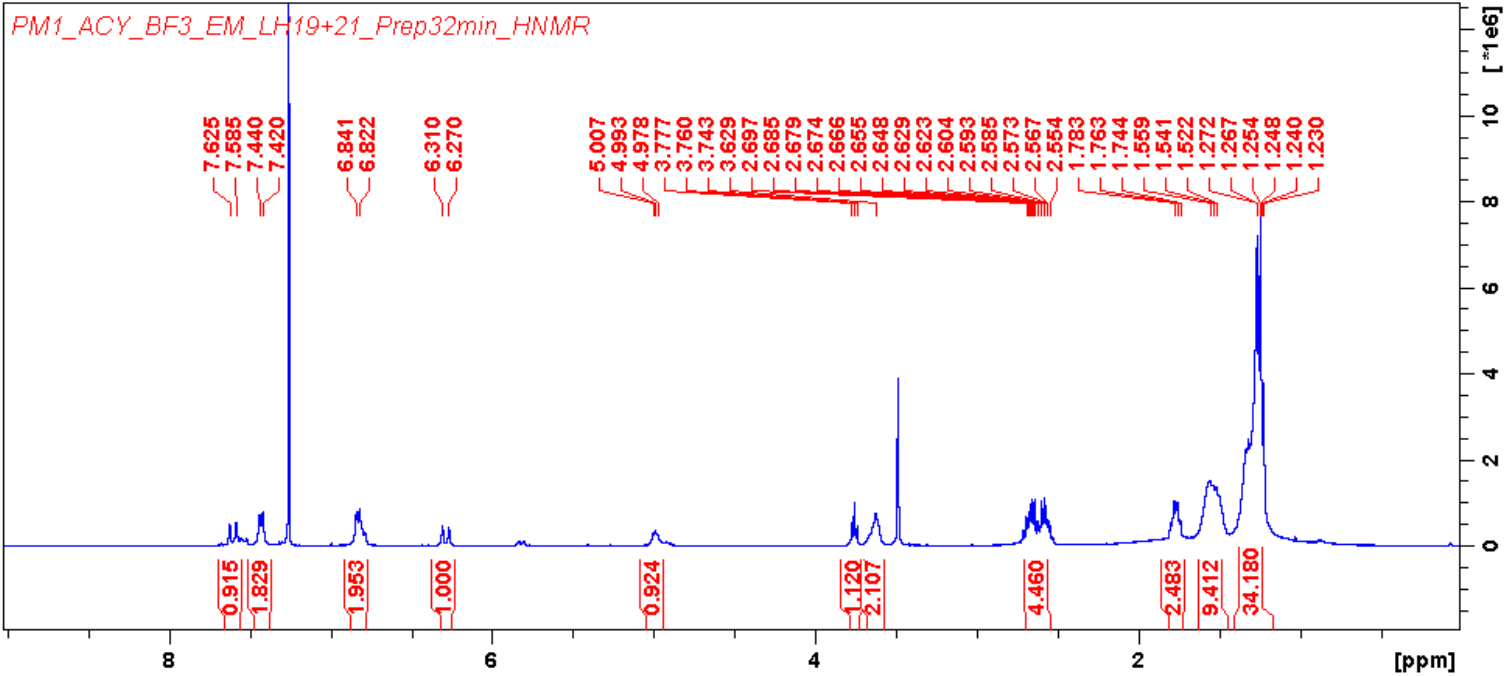
Characterization of compound II, ^1^H NMR (400 MHz, CDCl_3_) spectrum of compound **II**.

**Fig. S13.**
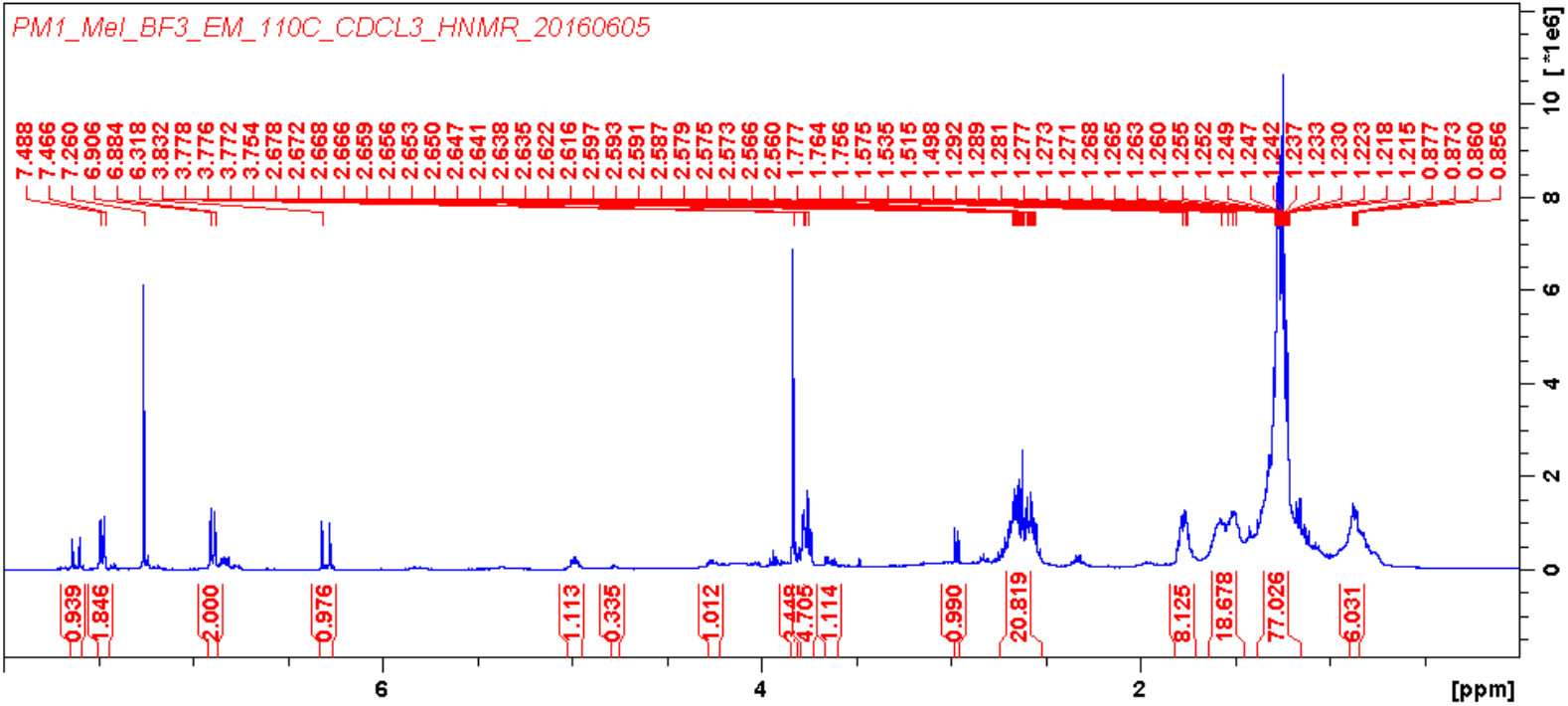
^1^H NMR (400 MHz, CDCl_3_) spectrum of total soluble products of methylated *P. rigida* sporopollenin thioacidolysis.

**Fig. S14-A.**
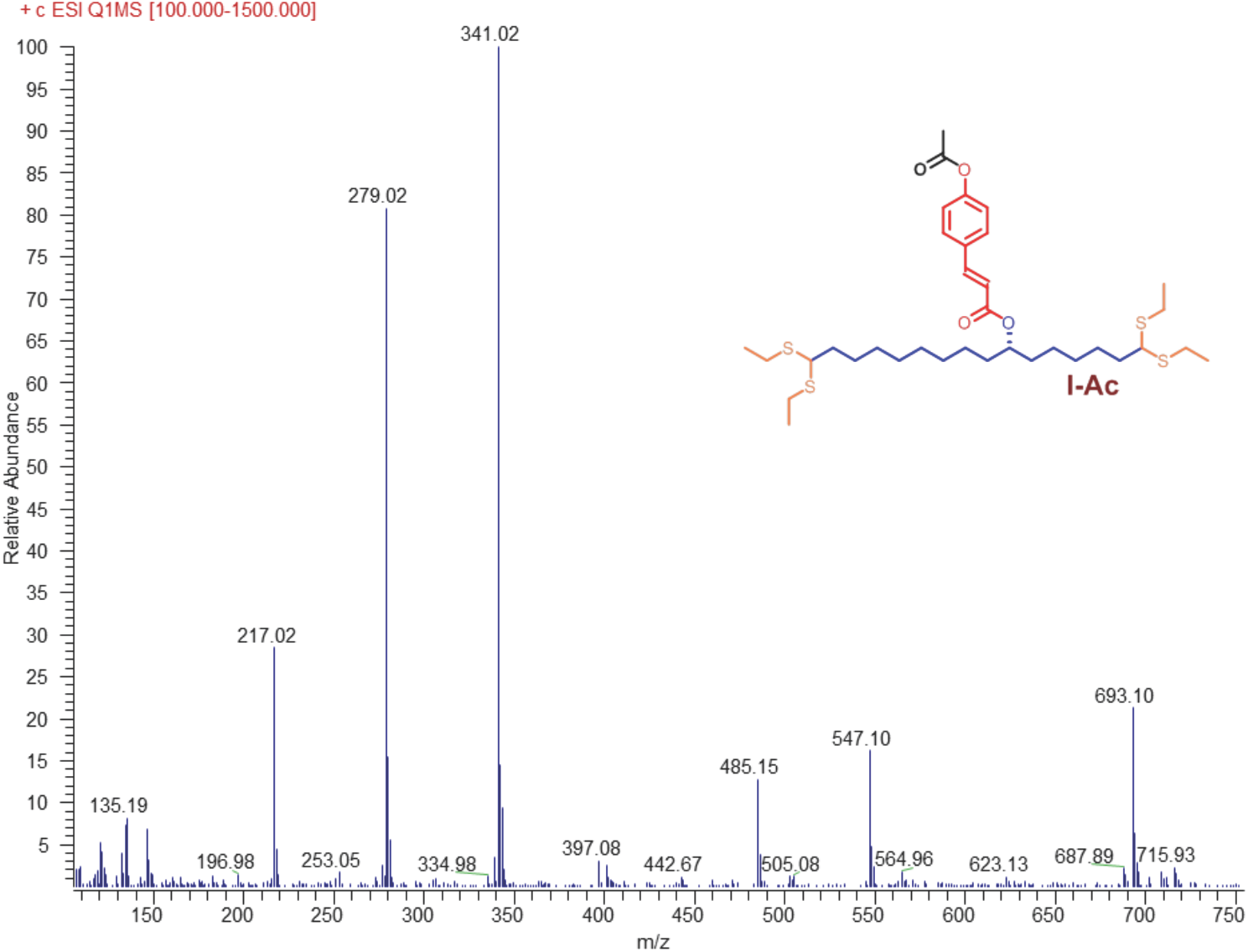
Characterization of compound I-Ac, MS/MS spectrum of compound **I-Ac**.

**Fig. S14-B.**
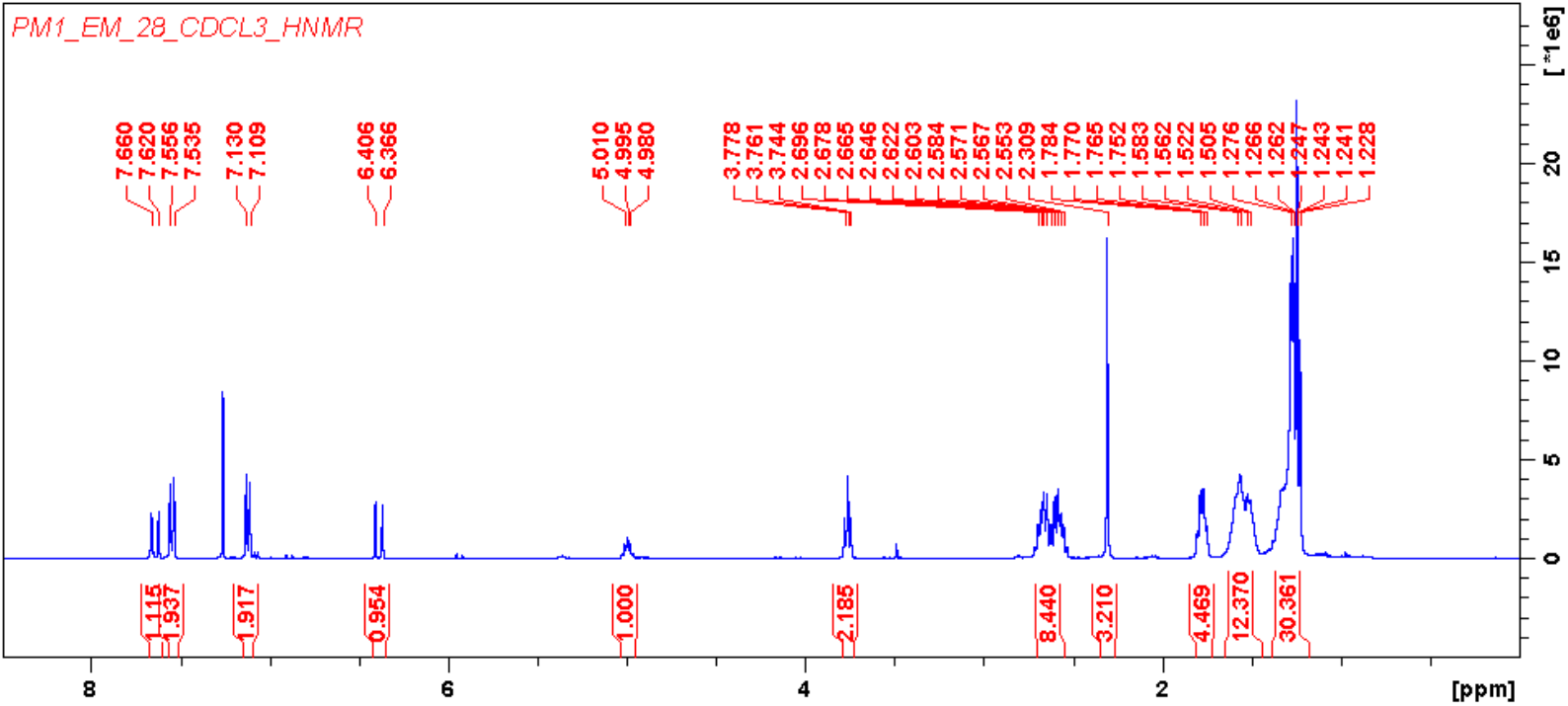
Characterization of compound I-Ac, ^1^H NMR (400 MHz, CDCl_3_) spectrum of compound **I-Ac**.

**Fig. S14-C.**
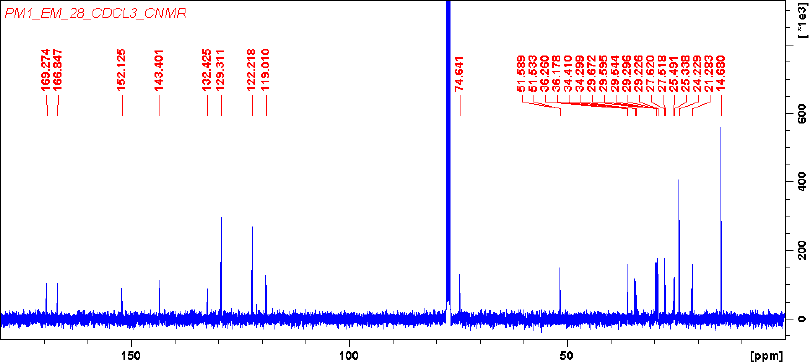
Characterization of compound I-Ac, ^13^C NMR (100 MHz, CDCl_3_) spectrum of compound **I-Ac**.

**Fig. S15-A.**
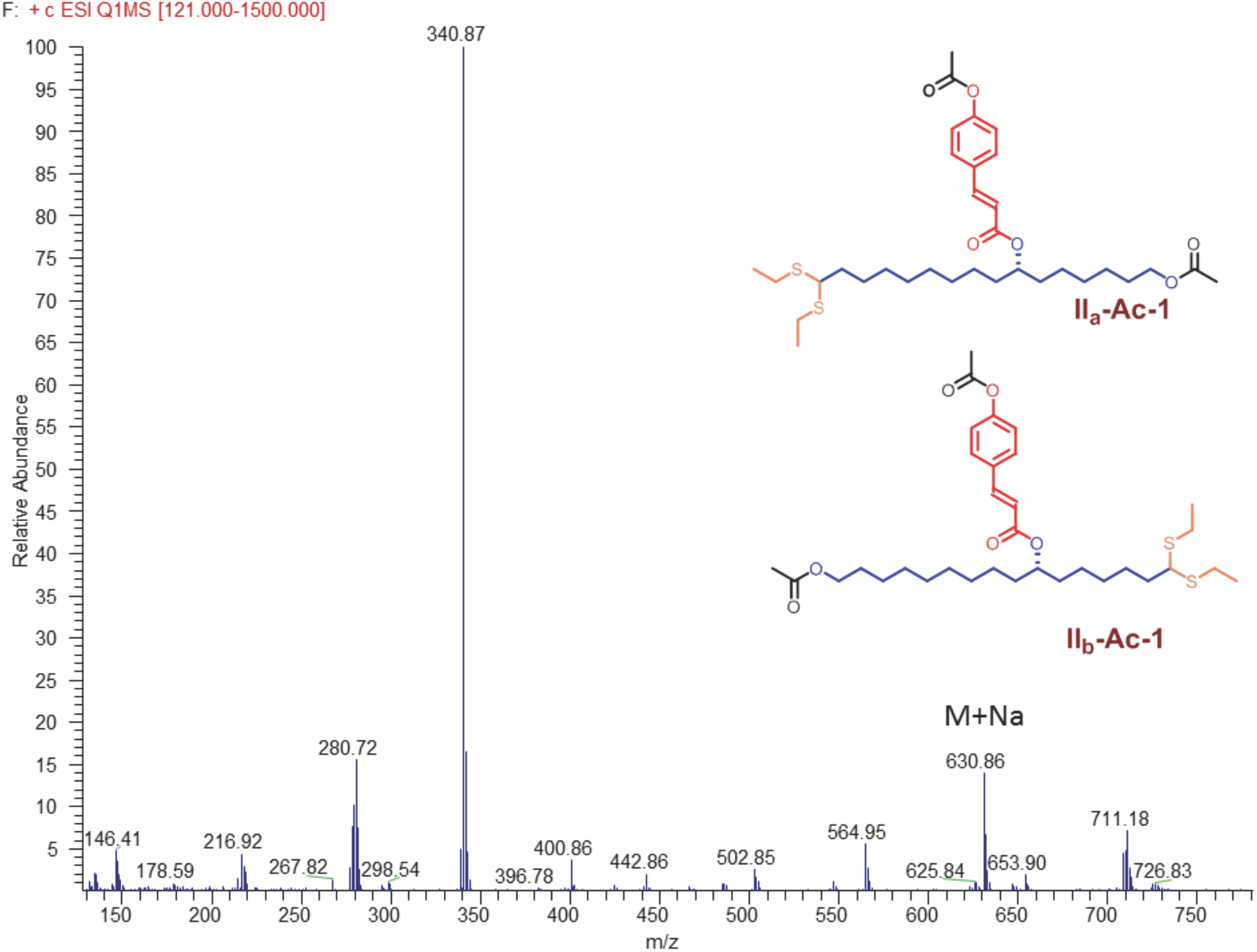
Characterization of compound II_(a+b)_-Ac-1, MS/MS spectrum of compound **II_(a+b)_-Ac-1**.

**Fig. S15-B.**
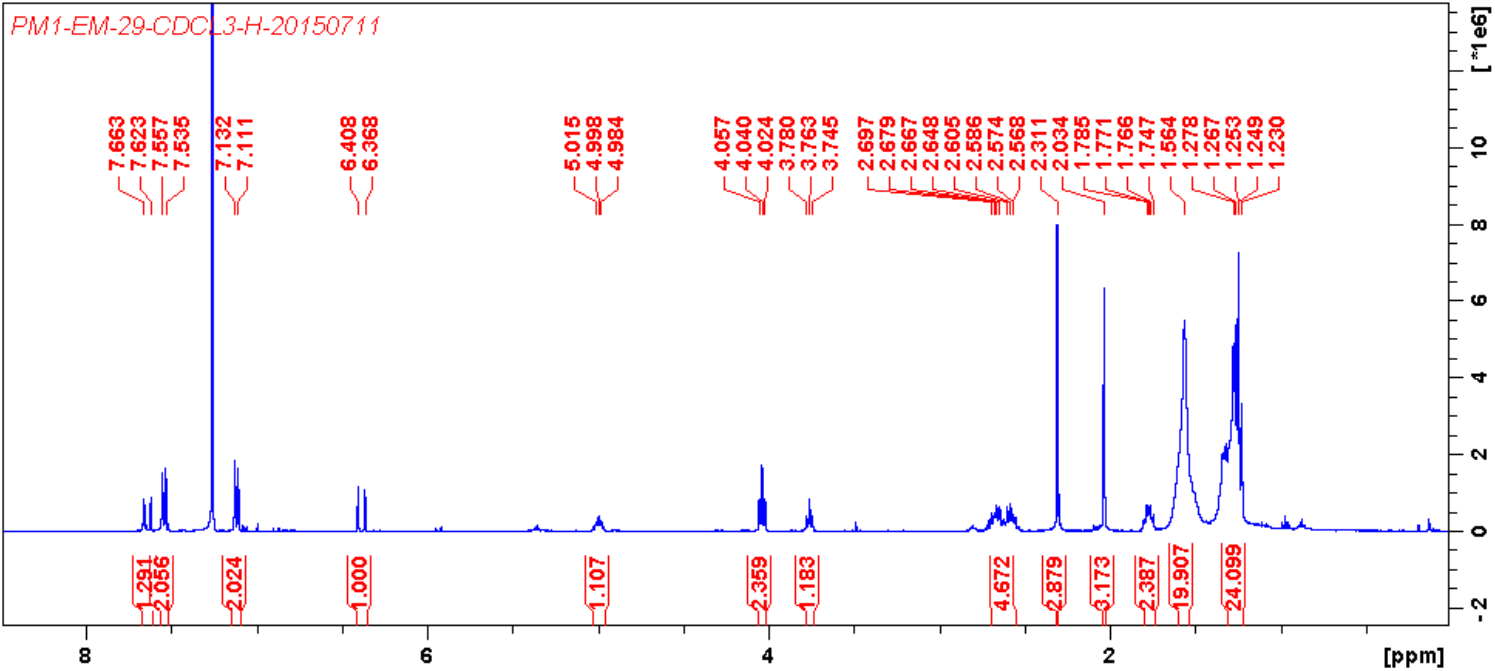
Characterization of compound II_(a+b)_-Ac-1, ^1^H NMR (400 MHz, CDCl_3_) spectrum of compound **II_(a+b)_-Ac-1**.

**Fig. S16.**
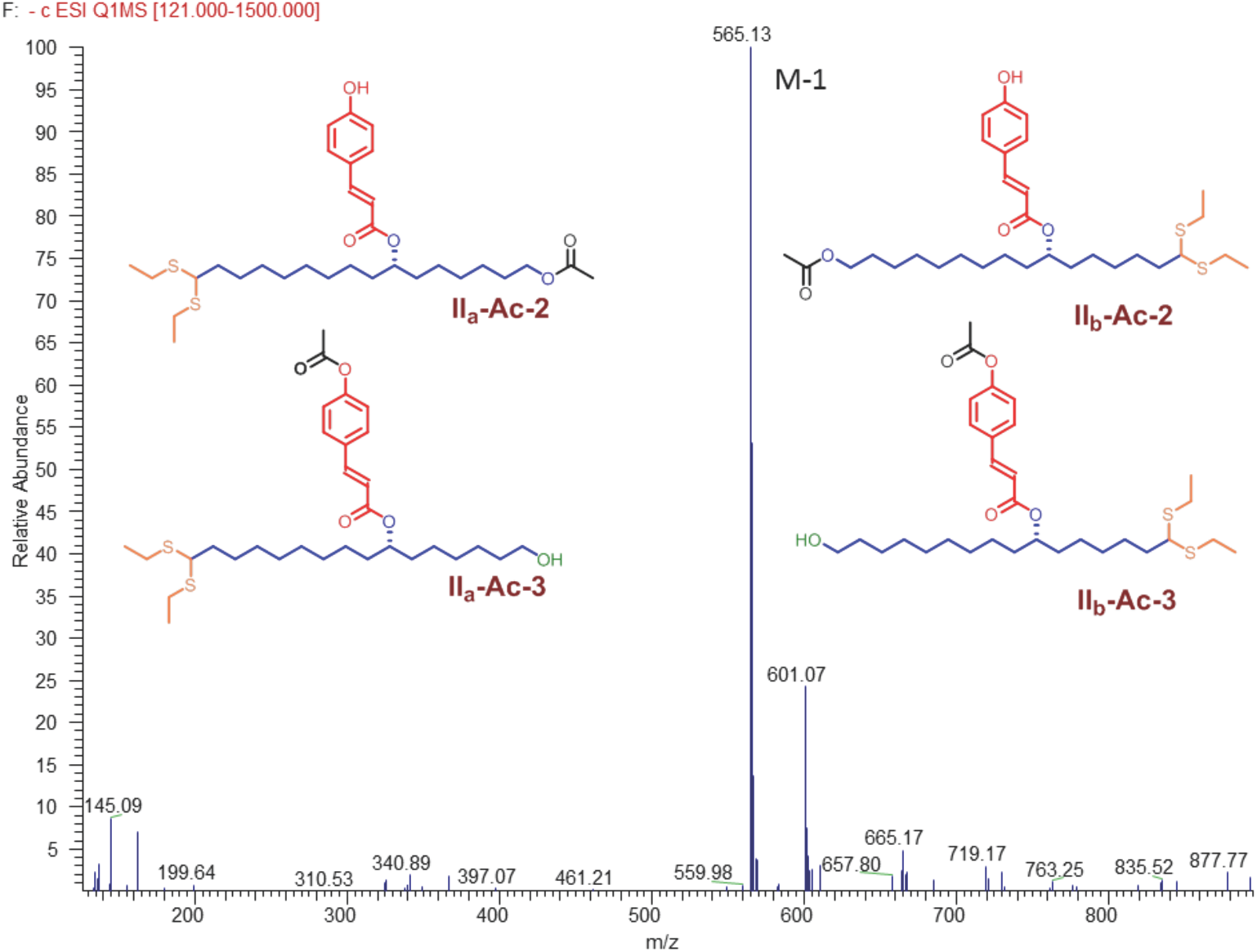
Characterization of compound II_(a+b)_-Ac-2 and II_(a+b)_-Ac-3, MS/MS spectrum of compound **II_(a+b)_-Ac-2 and II_(a+b)_-Ac-3**.

**Fig. S17.**
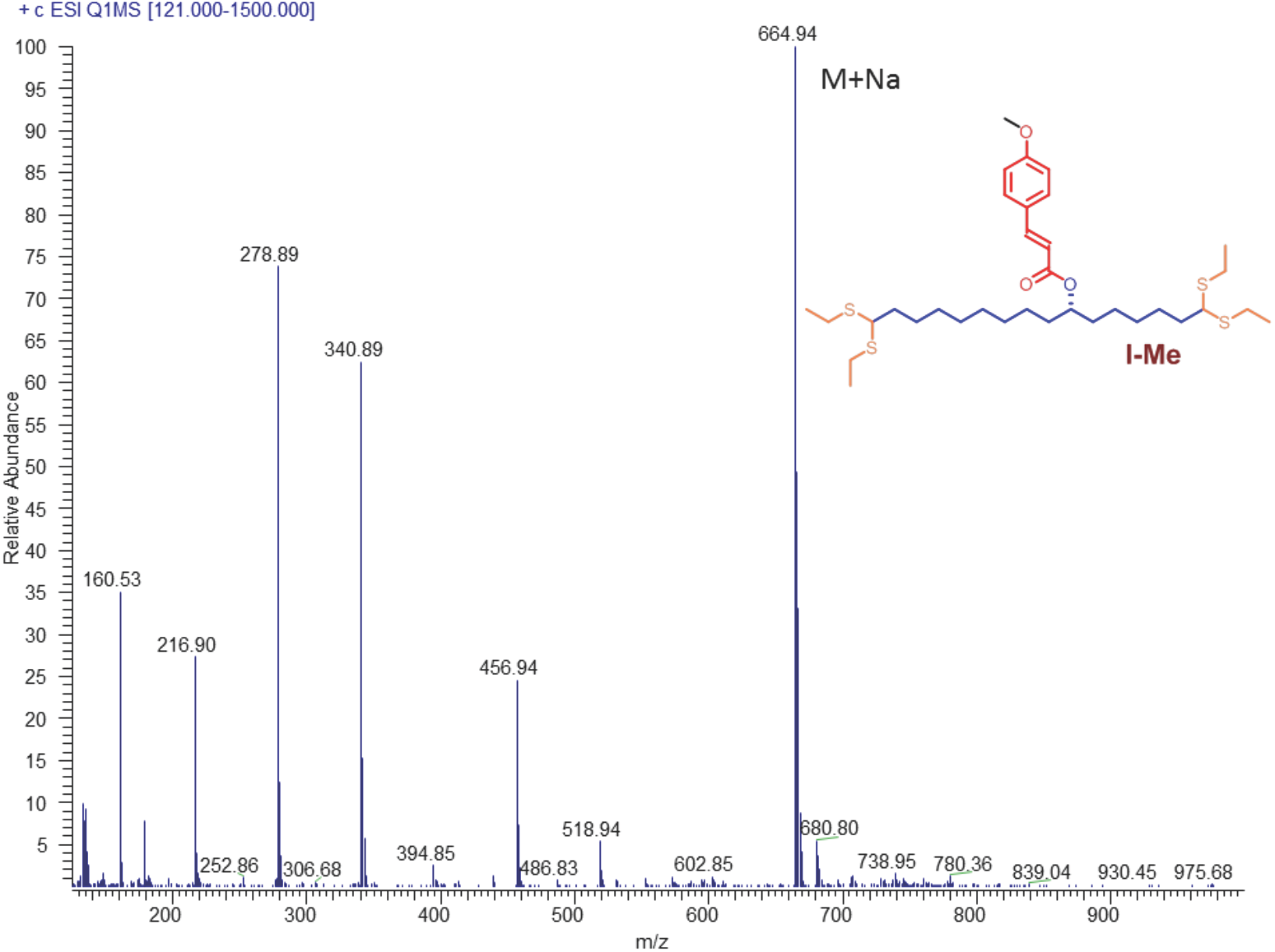
Characterization of compound I-Me, MS/MS spectrum of compound **I-Me.**

**Fig. S18.**
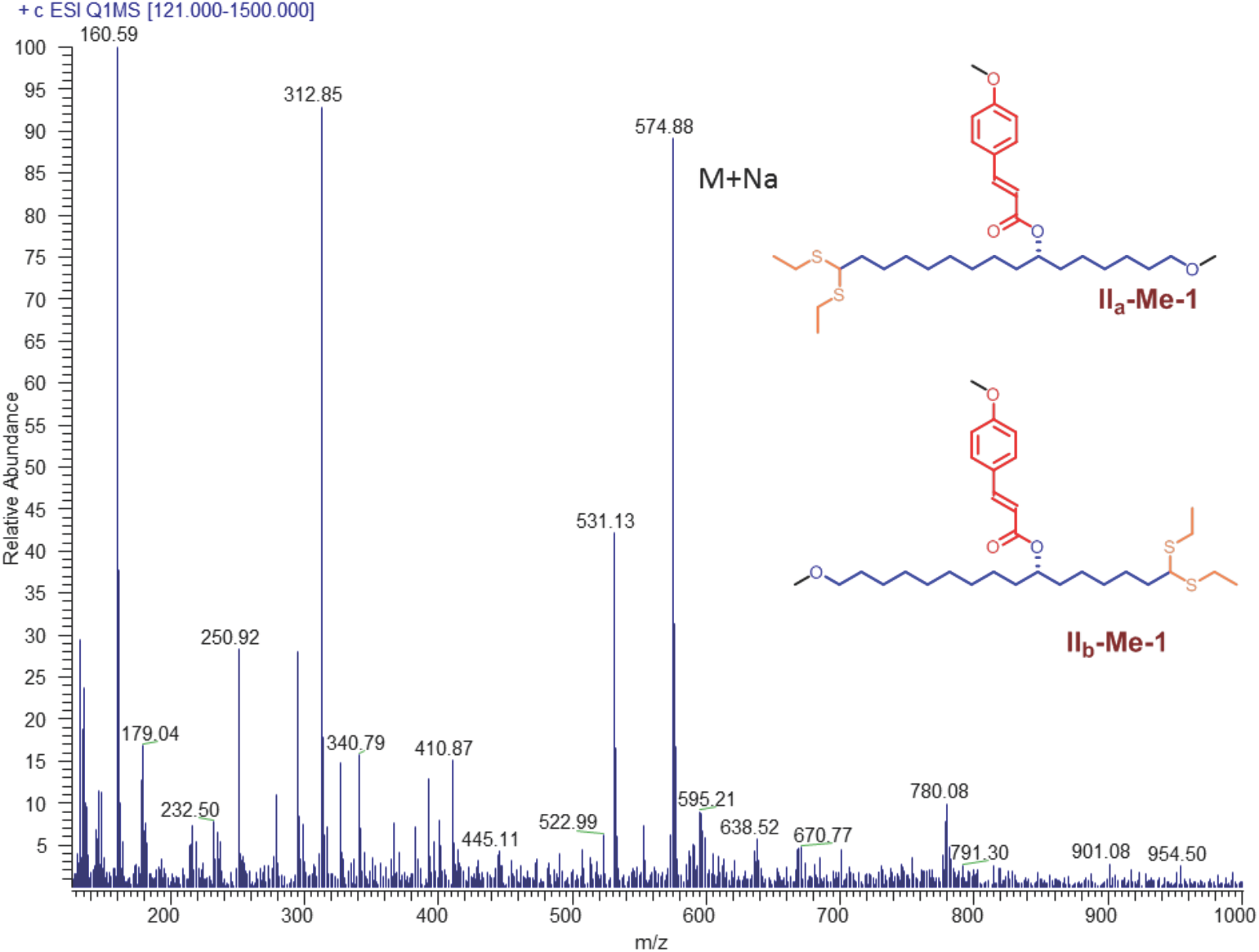
Characterization of compound II_(a+b)_-Me-1, MS/MS spectrum of compound **II_(a+b)_-Me-1.**

**Fig. S19.**
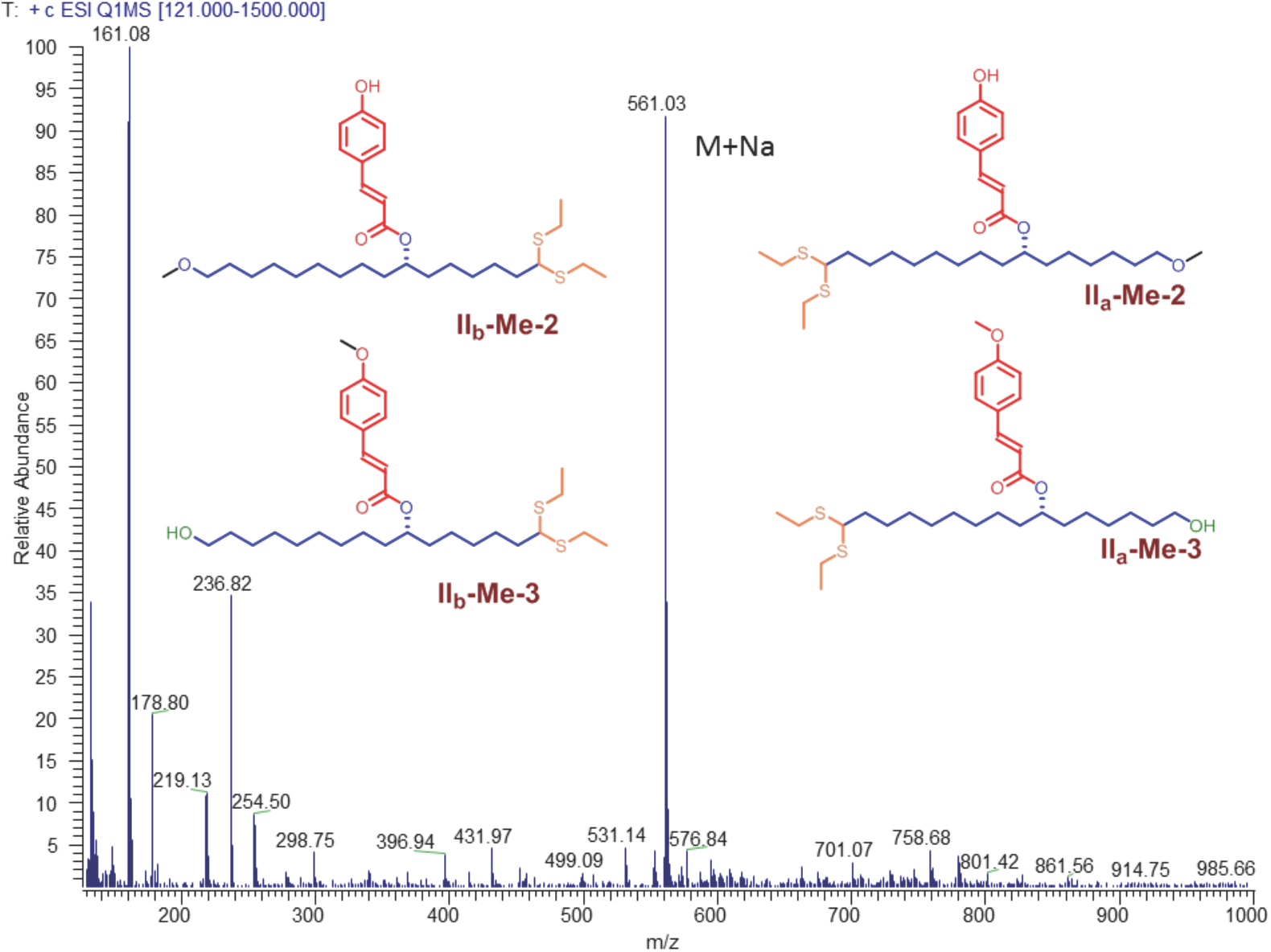
Characterization of compound II_(a+b)_-Me-2 and II_(a+b)_-Me-3, MS/MS spectrum of compound **II_(a+b)_-Me-2 and II_(a+b)_-Me-3**.

**Fig. S20-A.**
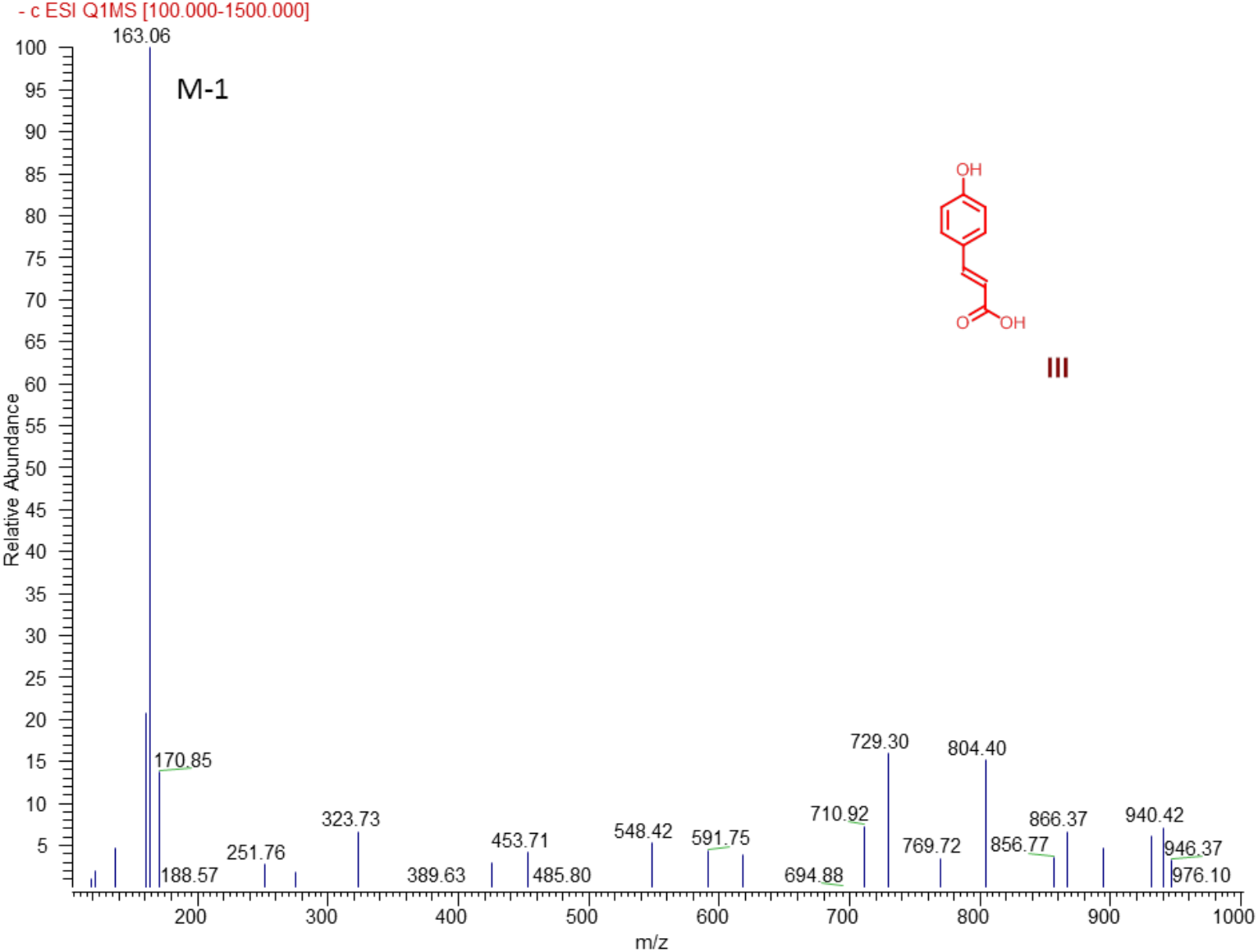
MS/MS spectrum of compound III.

**Fig. S20-B.**
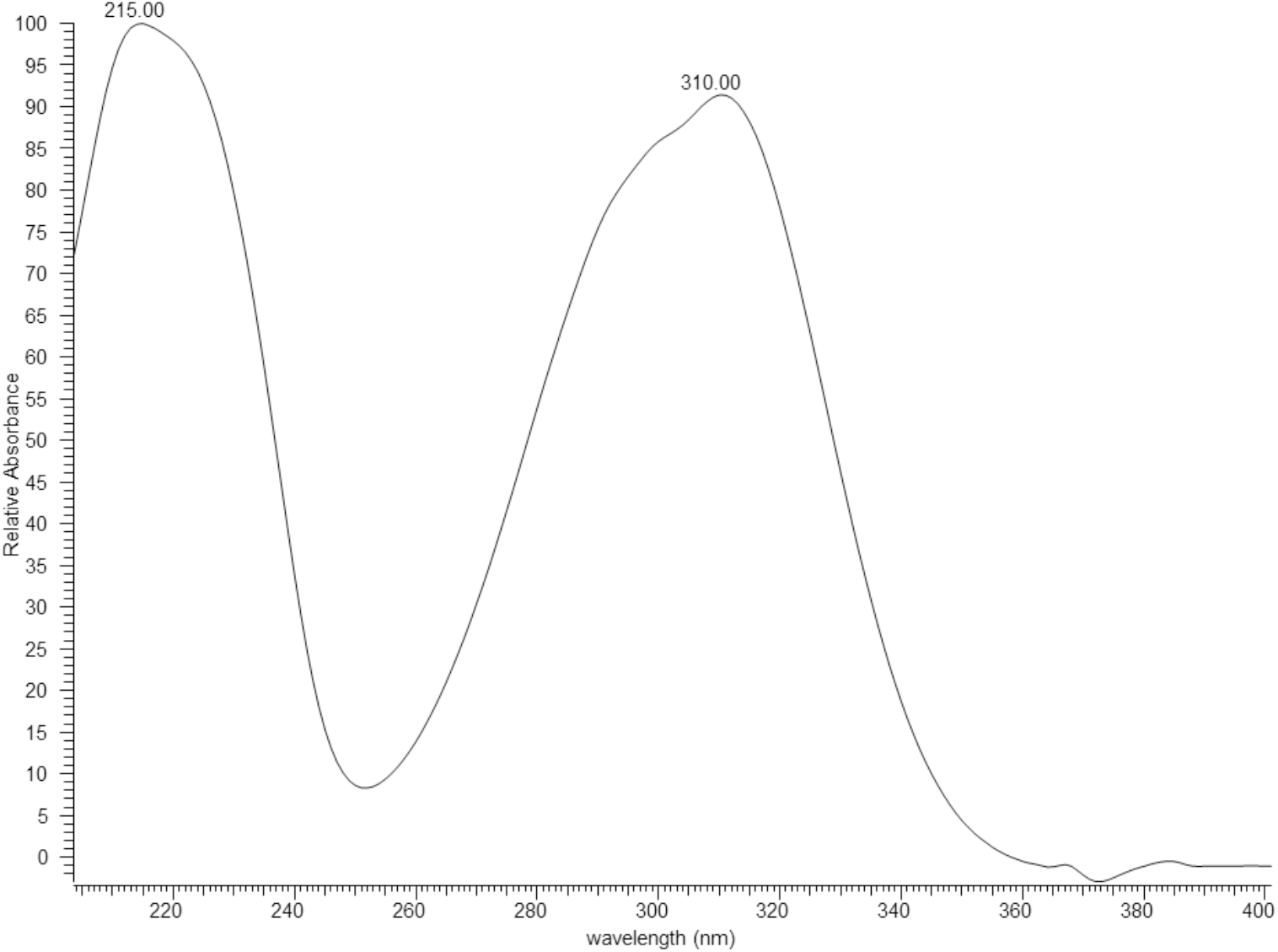
UV spectrum of compound III, Compound **III** was confirmed as *p*- coumaric acid by comparing its retention time, UV and MS/MS spectra with that of the reference standard.

**Fig. S21-A.**
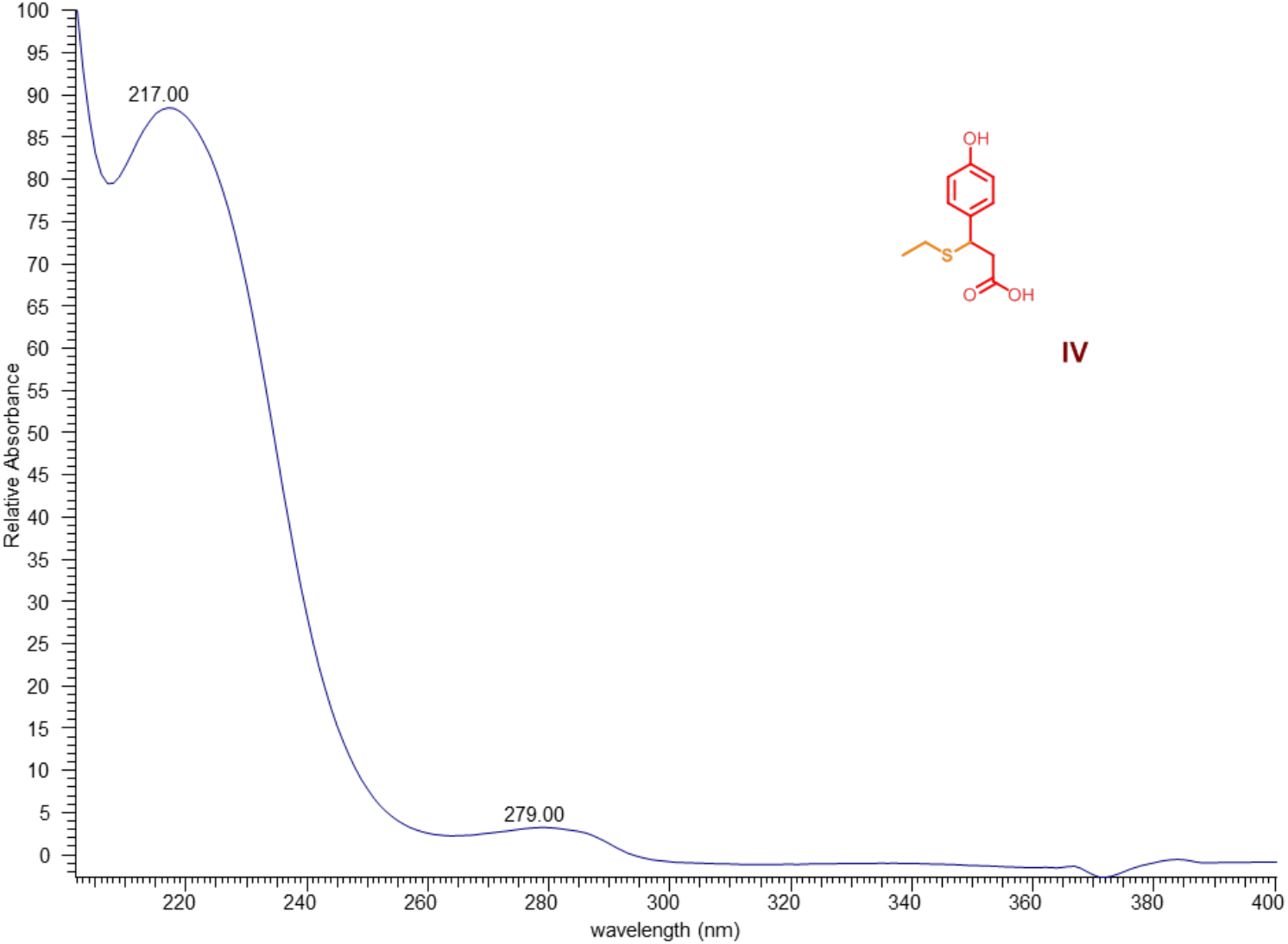
Characterization of compound IV, UV spectrum of compound **IV.**

**Fig. S21-B.**
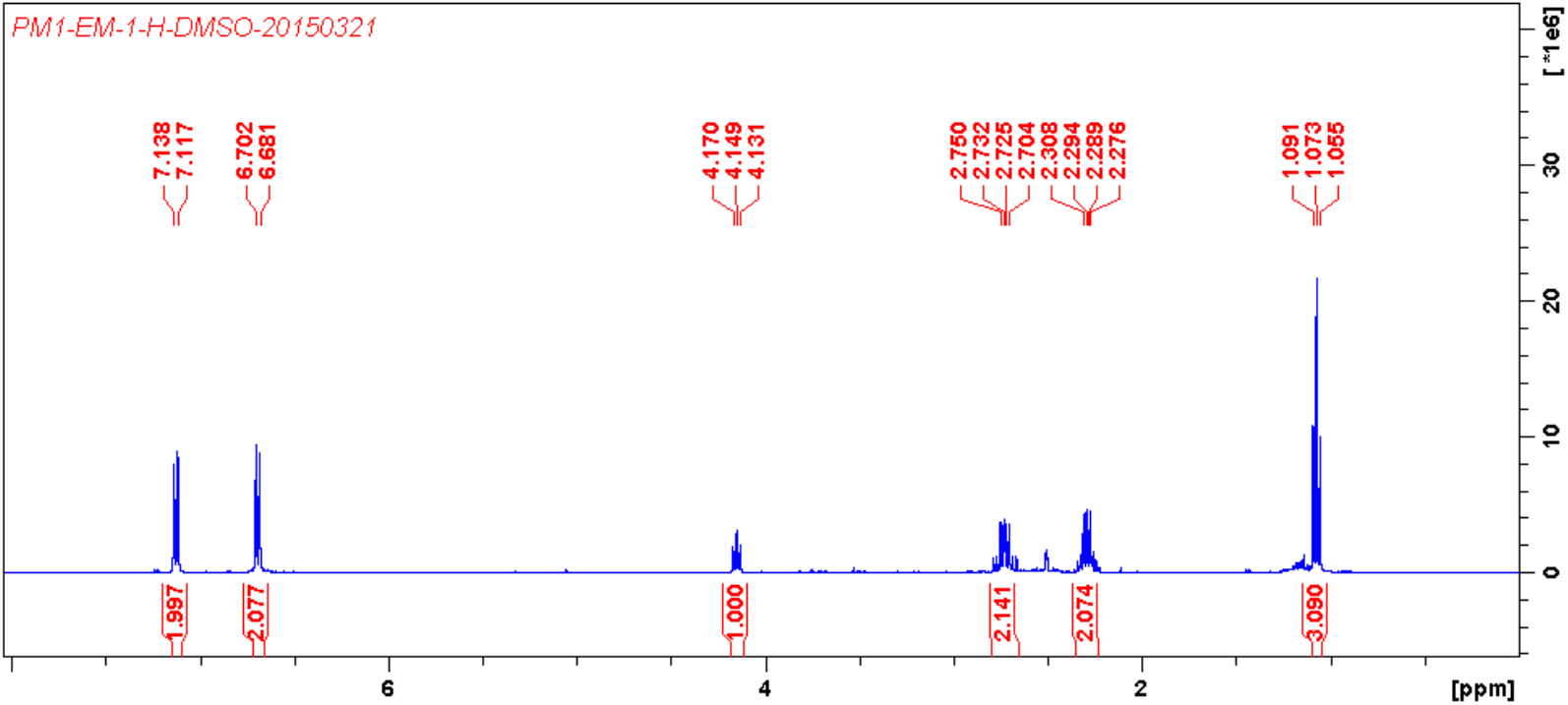
Characterization of compound IV, ^1^H NMR (400 MHz, DMSO-*d*_*6*_) spectrum of compound **IV.**

**Fig. S21-C.**
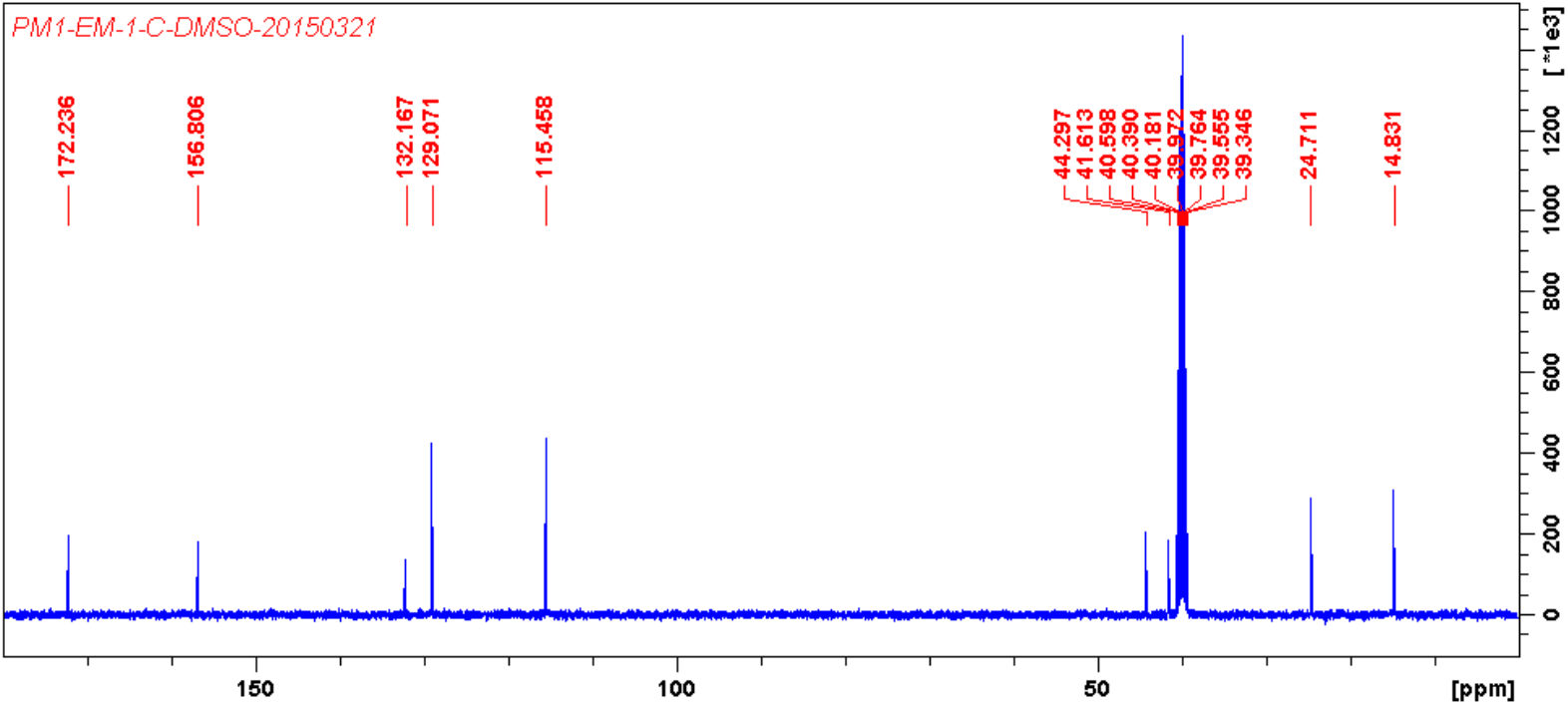
Characterization of compound IV, ^13^C NMR (100 MHz, DMSO-*d*_*6*_) spectrum of compound **IV.**

**Fig. S22-A.**
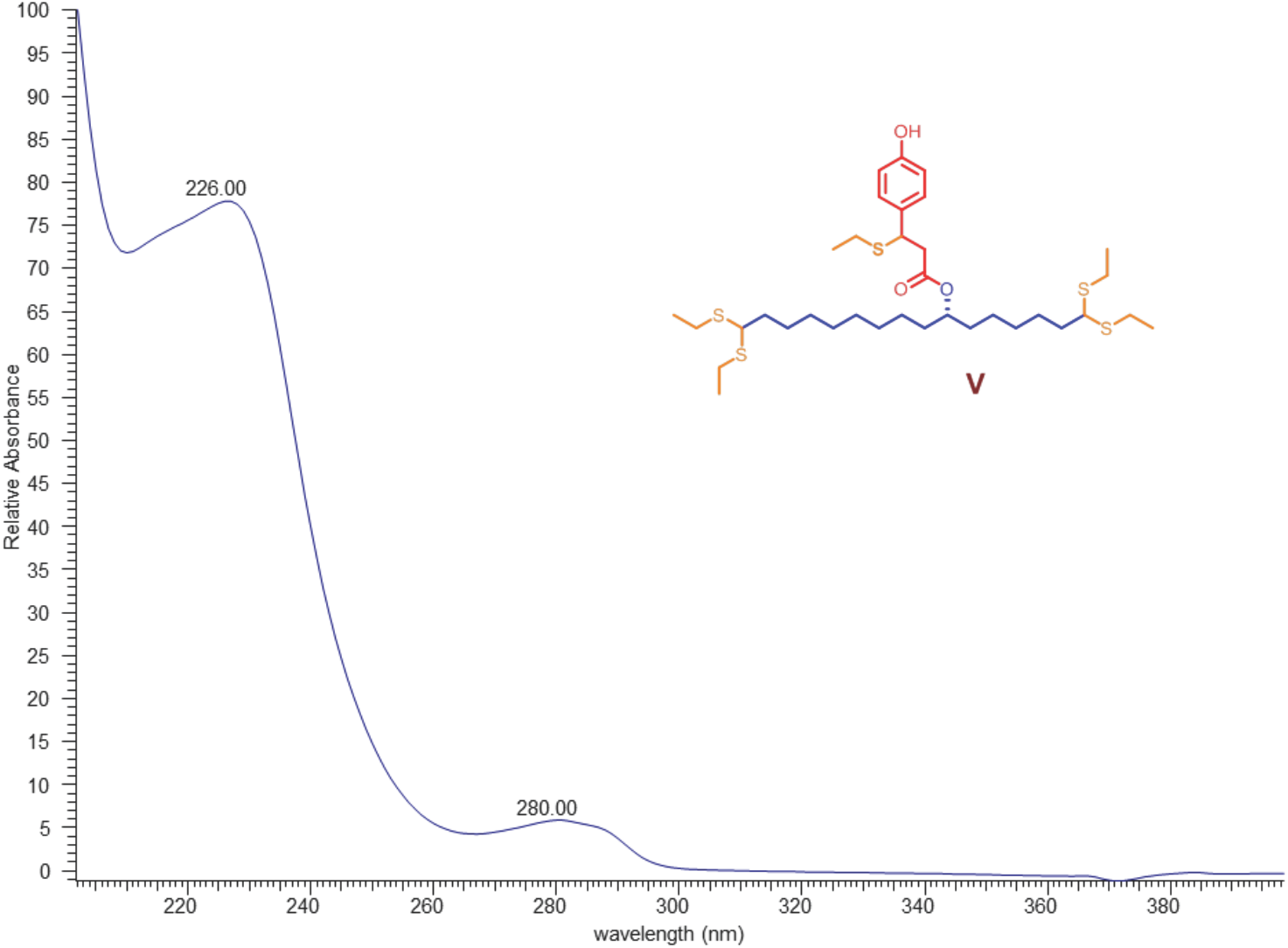
Characterization of compound V, UV spectrum of compound **V.**

**Fig. S22-B.**
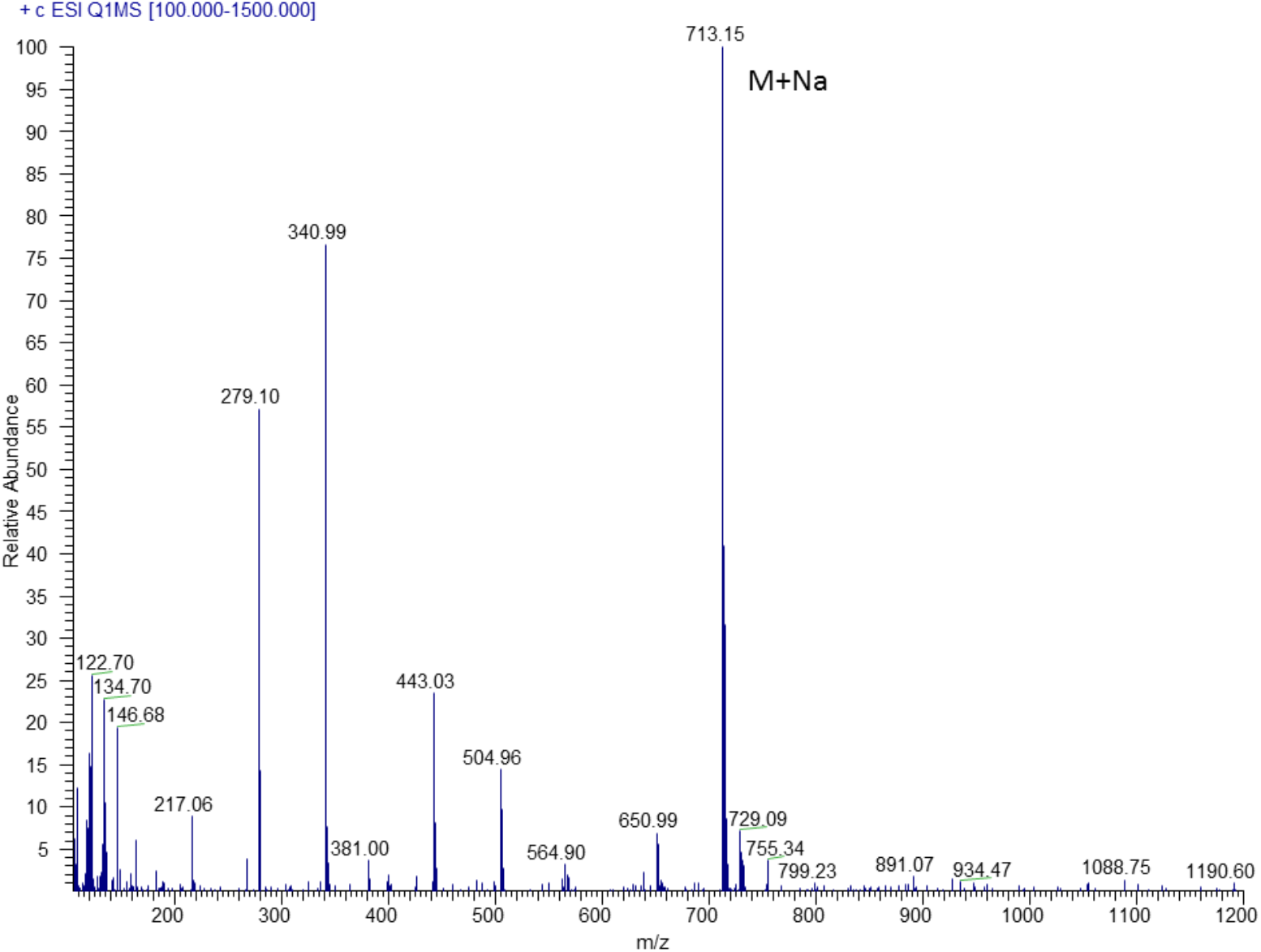
Characterization of compound V, MS/MS spectrum of compound **V.**

**Fig. S22-C.**
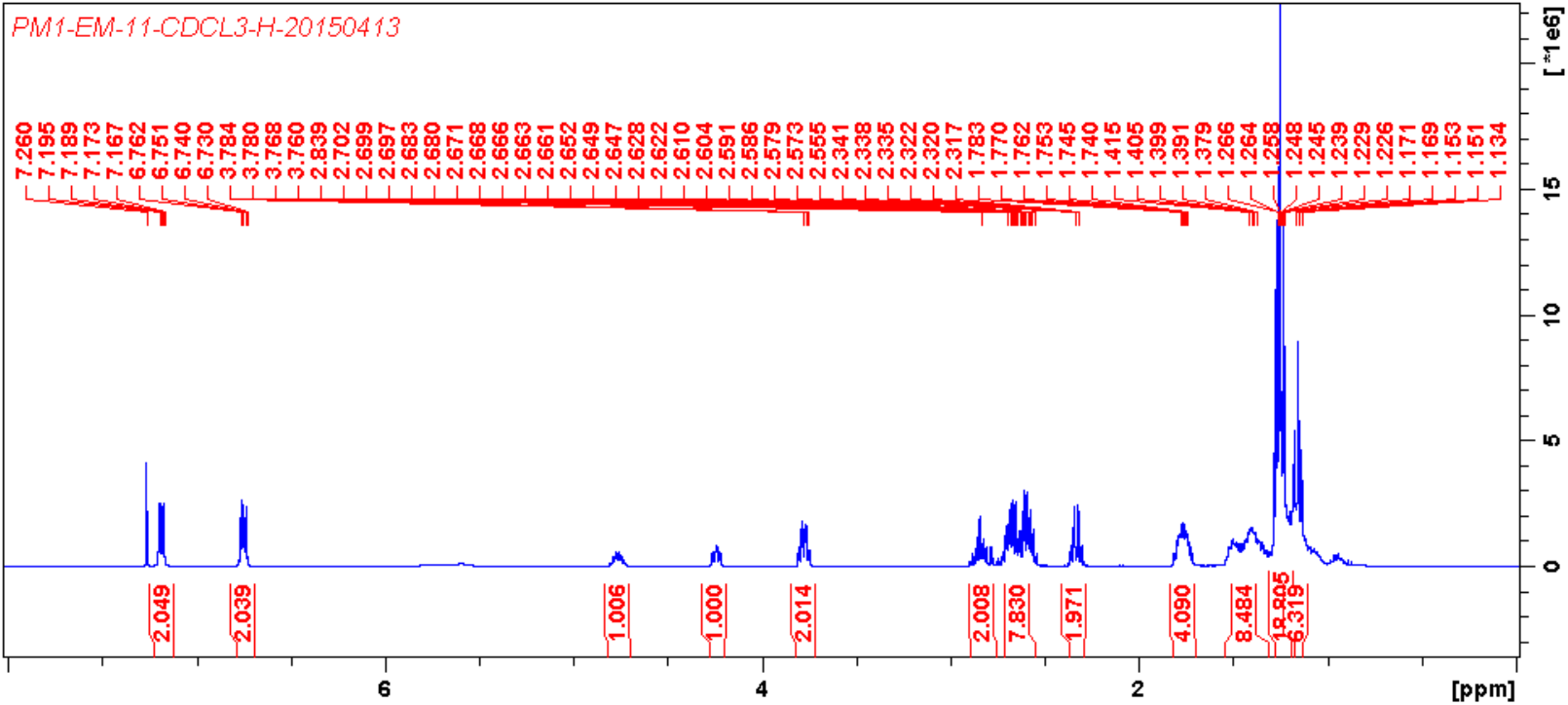
Characterization of compound V, ^1^H NMR (400 MHz, CDCl_3_) spectrum of compound **V.**

**Fig. S22-D.**
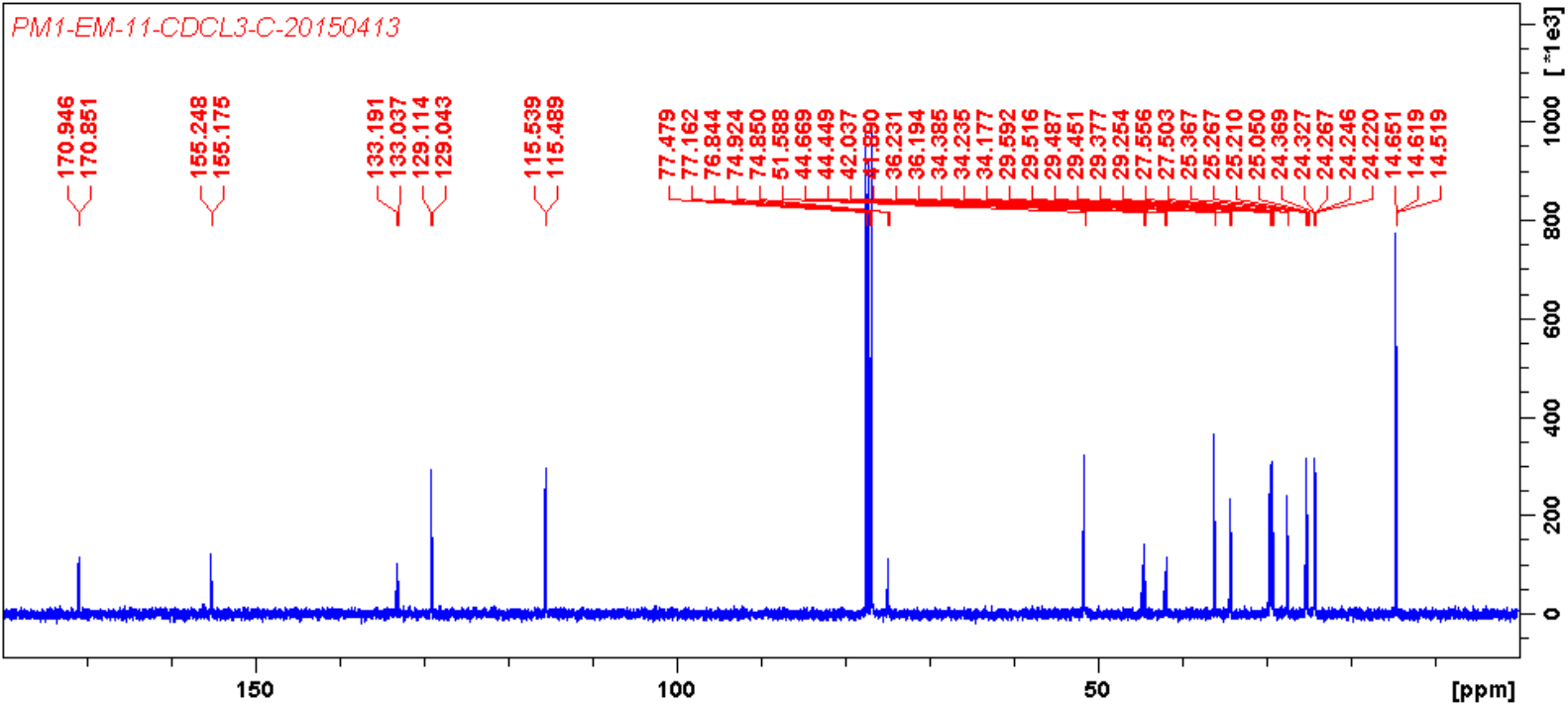
Characterization of compound V, ^13^C NMR (100 MHz, CDCl_3_) spectrum of compound **V.**

**Fig. S22-E.**
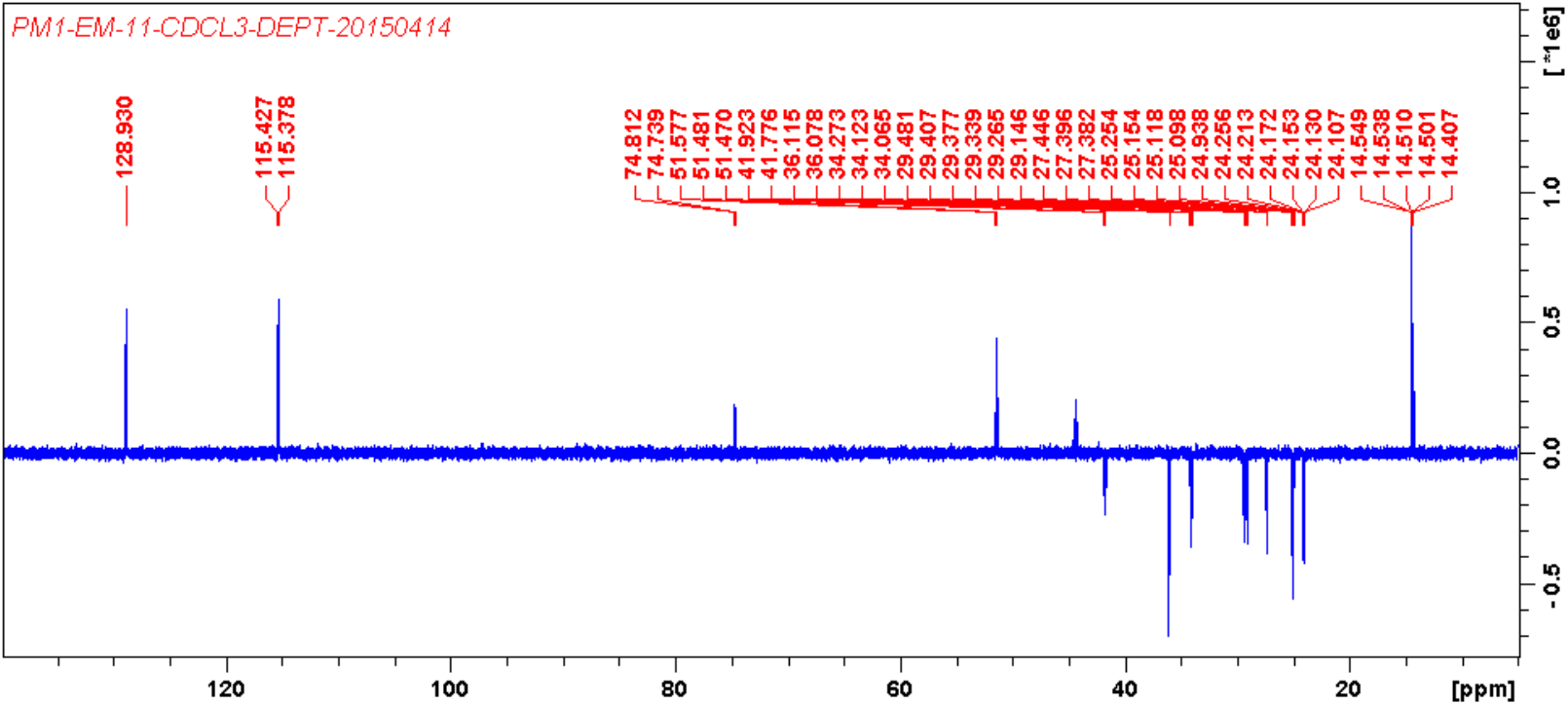
Characterization of compound V, DEPT spectrum of compound **V.**

**Fig. S22-F.**
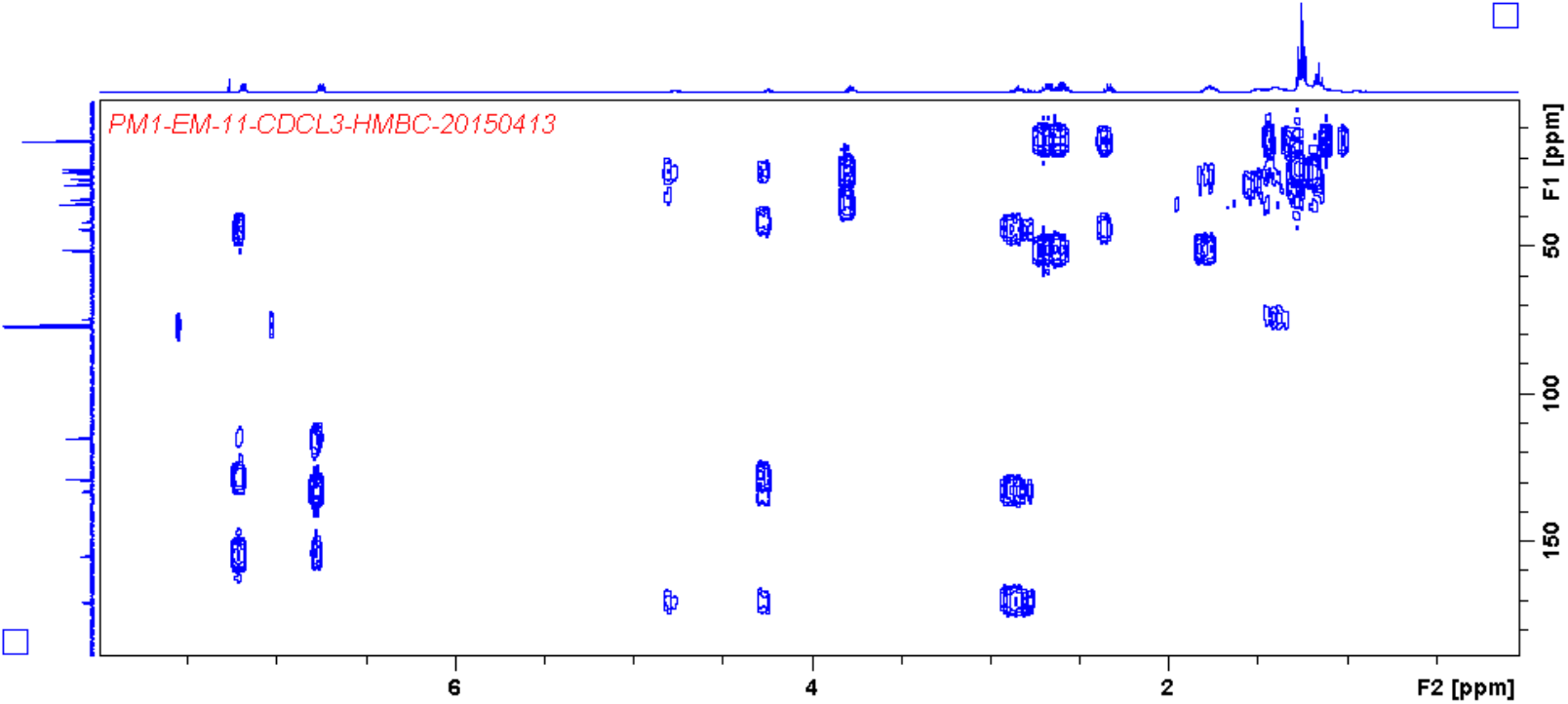
Characterization of compound V, HMBC spectrum of compound **V.**

**Fig. S22-G.**
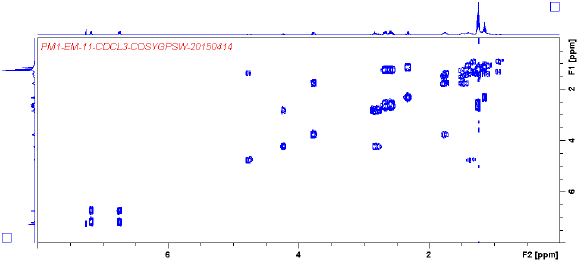
Characterization of compound V, HH-COSY spectrum of compound **V.**

**Fig. S23-A.**
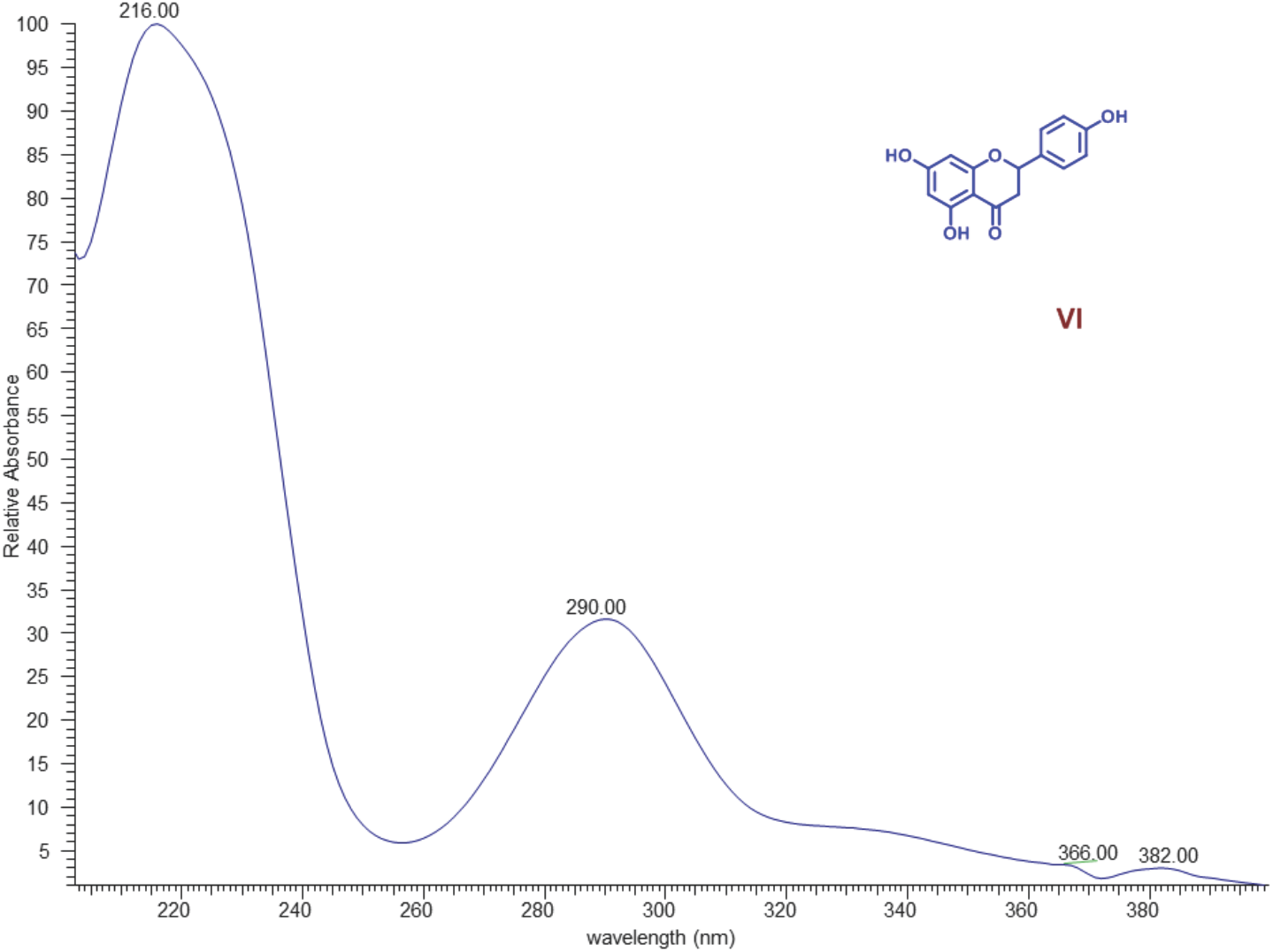
UV spectrum of compound VI.

**Fig. S23-B.**
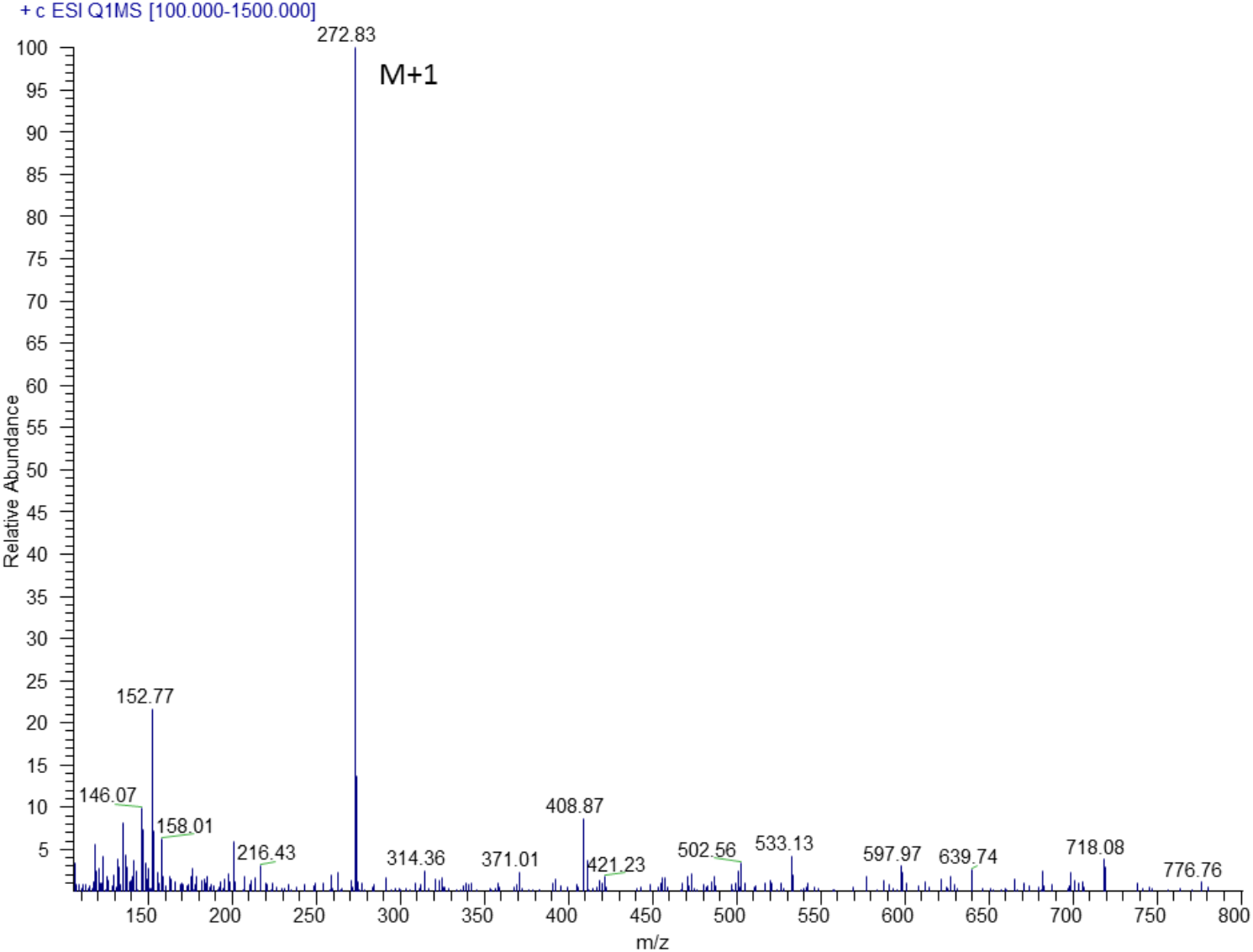
MS/MS spectrum of compound VI, Compound **VI** was confirmed as naringenin by comparing its retention time, UV and MS/MS spectra with that of the reference standard.

**Fig. S24-A.**
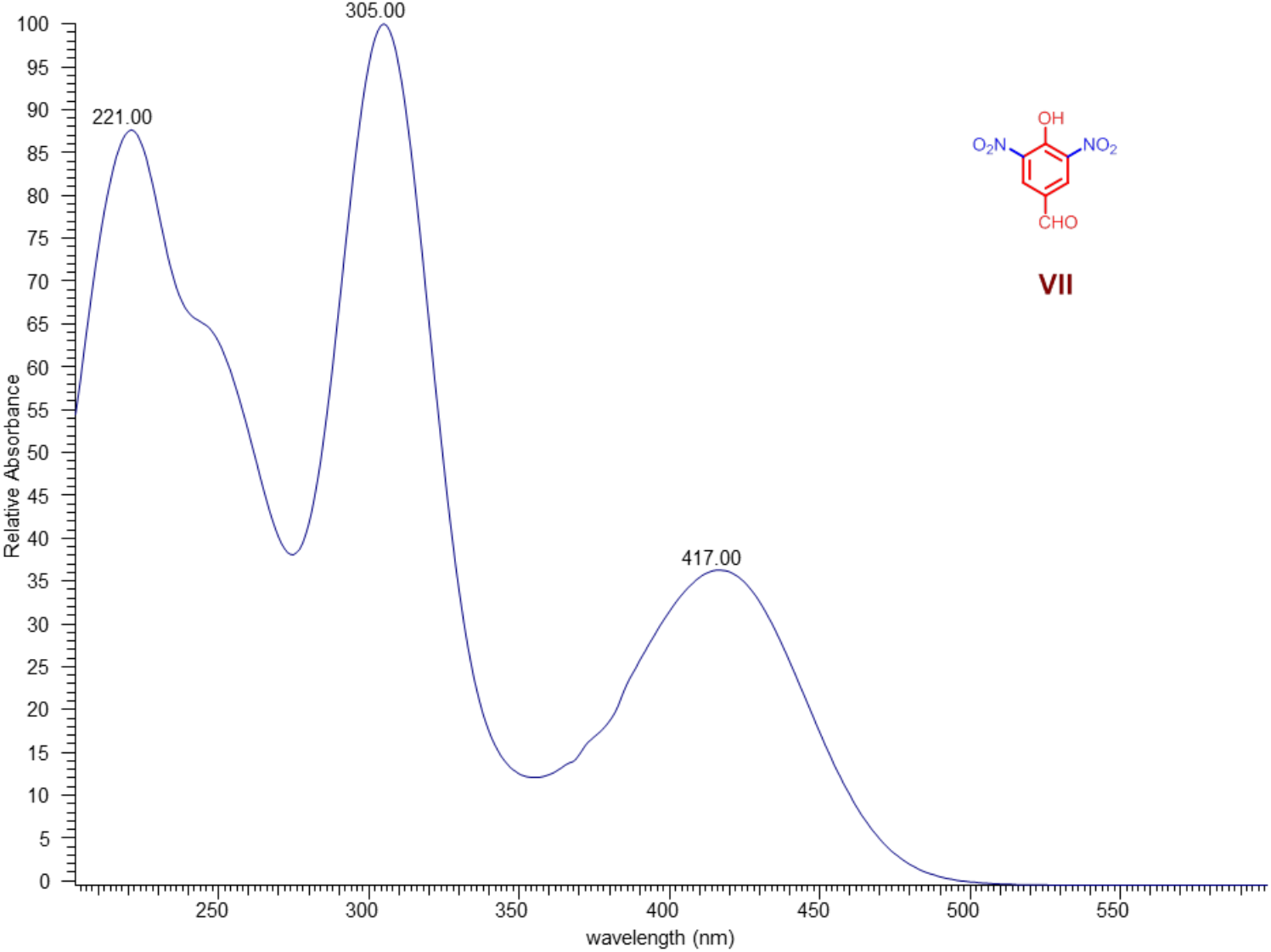
Characterization of compound VII, UV spectrum of compound **VII.**

**Fig. S24-B.**
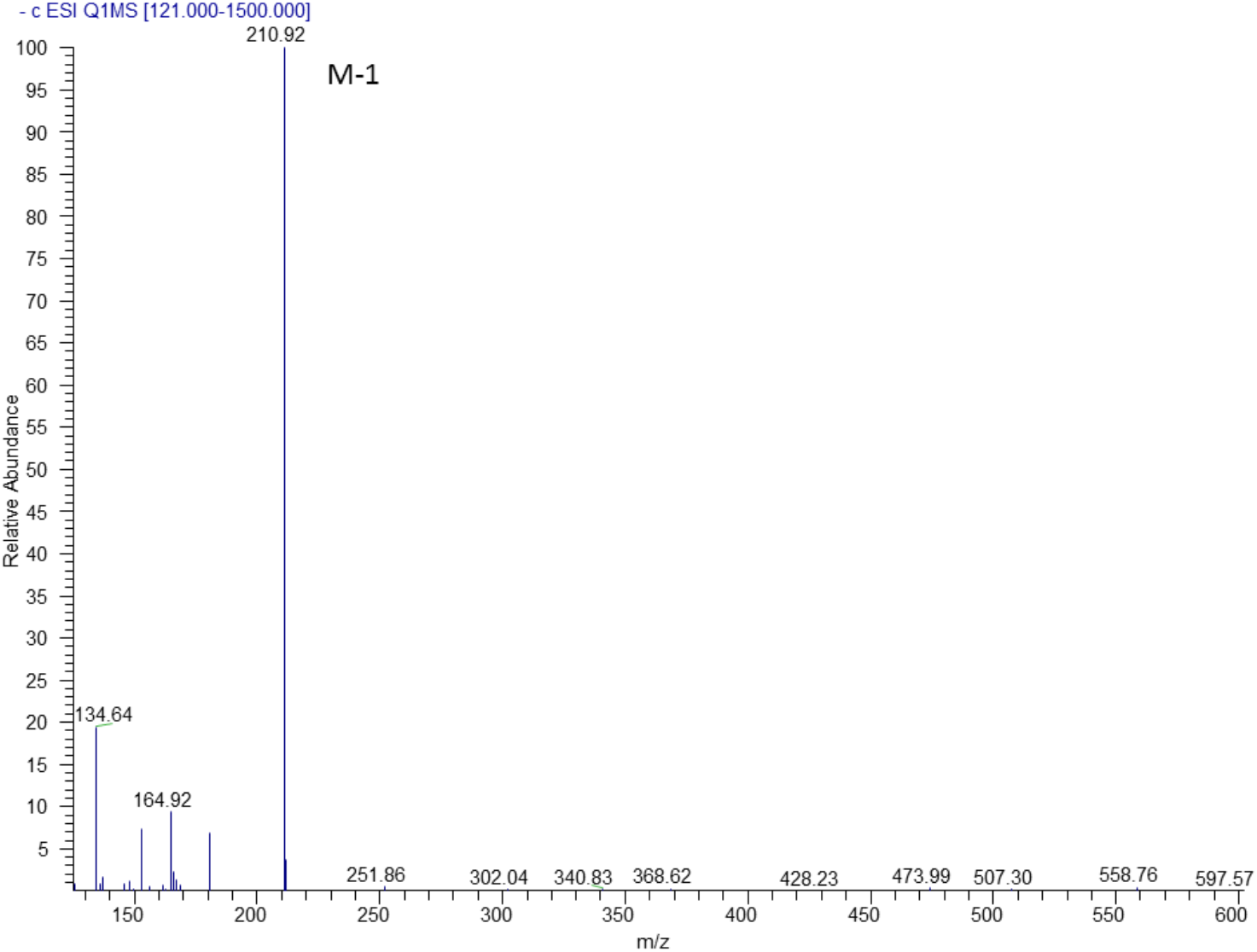
Characterization of compound VII, MS/MS spectrum of compound **VII.**

**Fig. S25-A.**
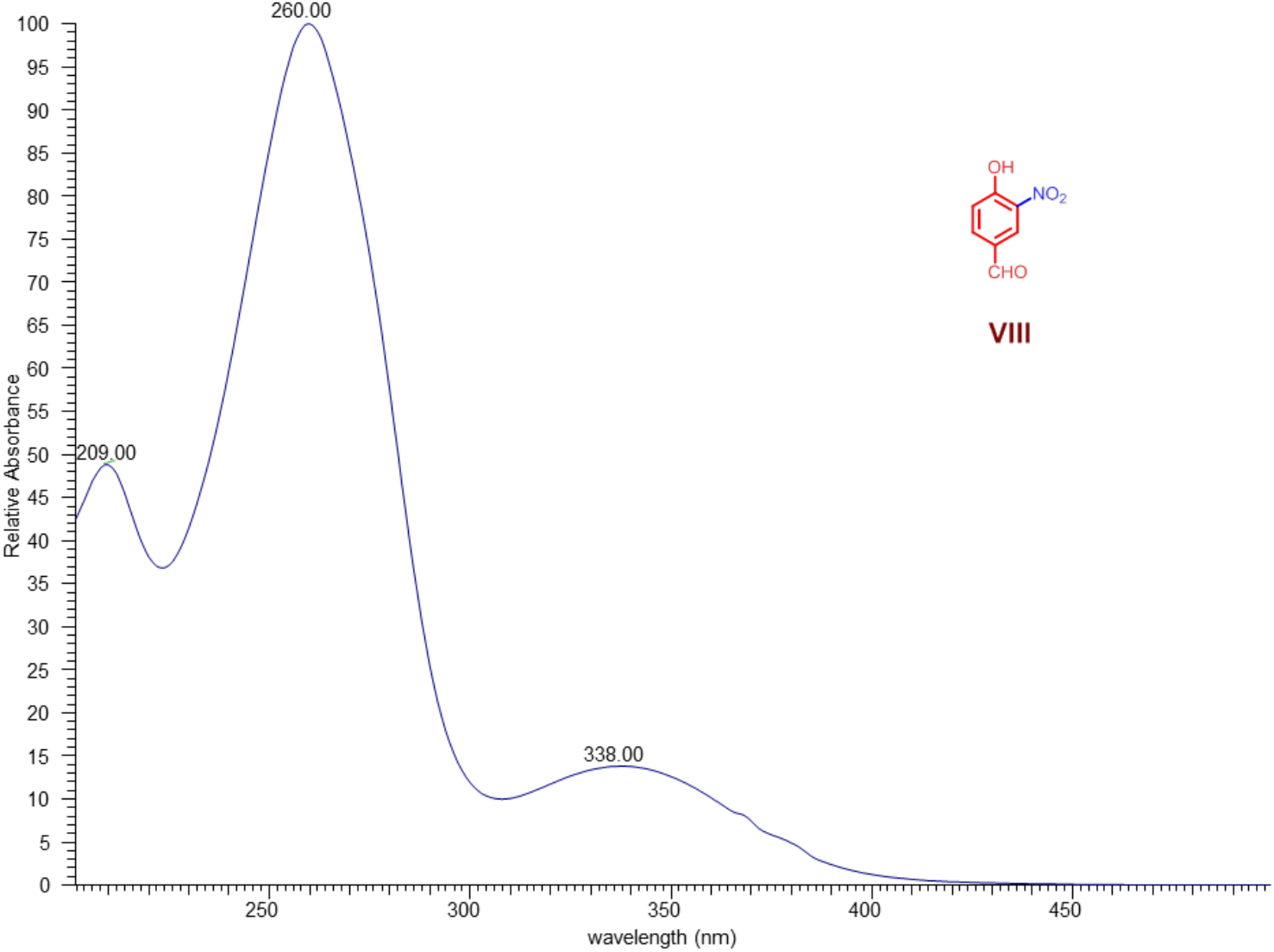
Characterization of compound VIII, UV spectrum of compound **VIII.**

**Fig. S25-B.**
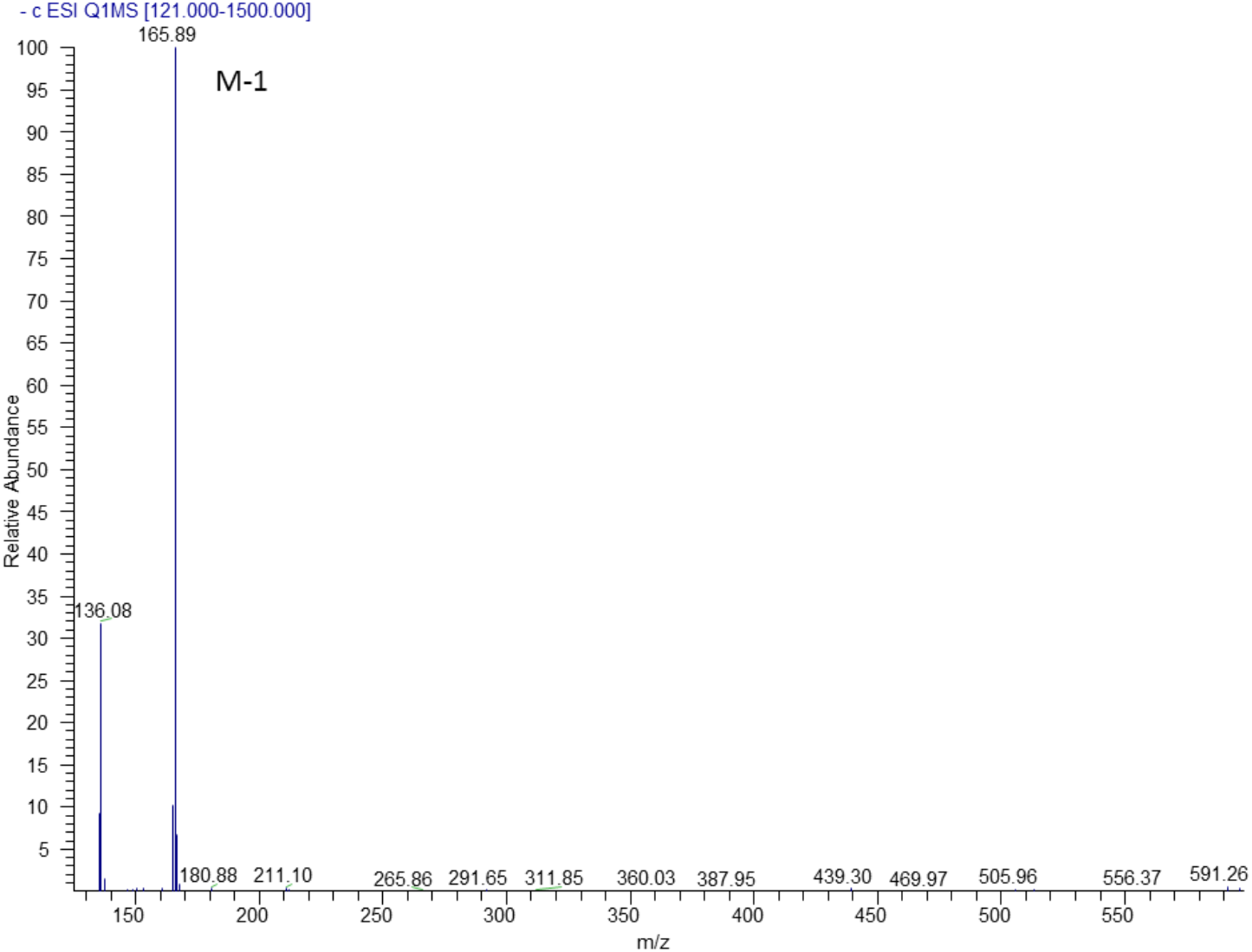
Characterization of compound VIII, MS/MS spectrum of compound **VIII.**

**Fig. S26-A.**
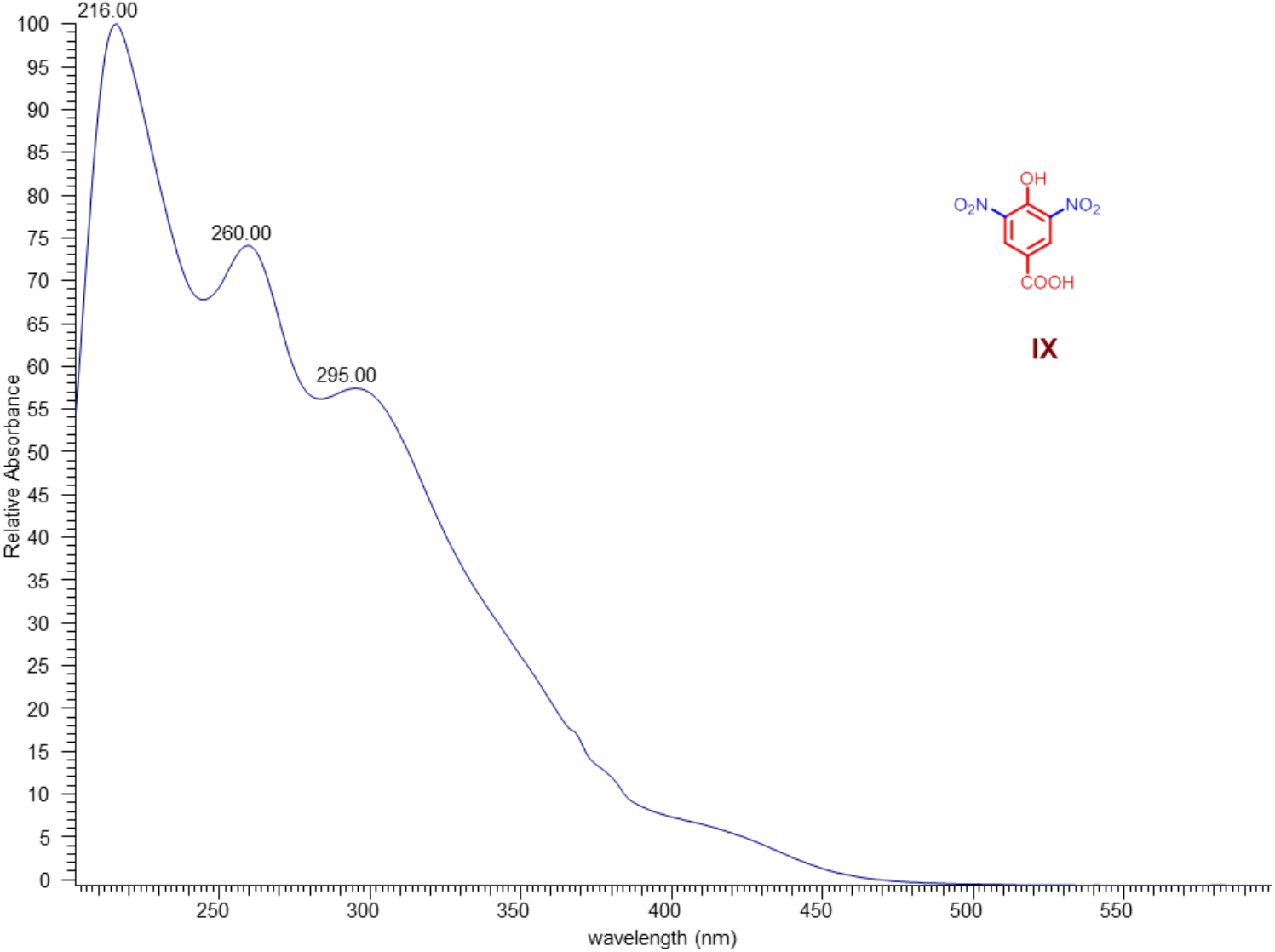
Characterization of compound IX, UV spectrum of compound **IX.**

**Fig. S26-B.**
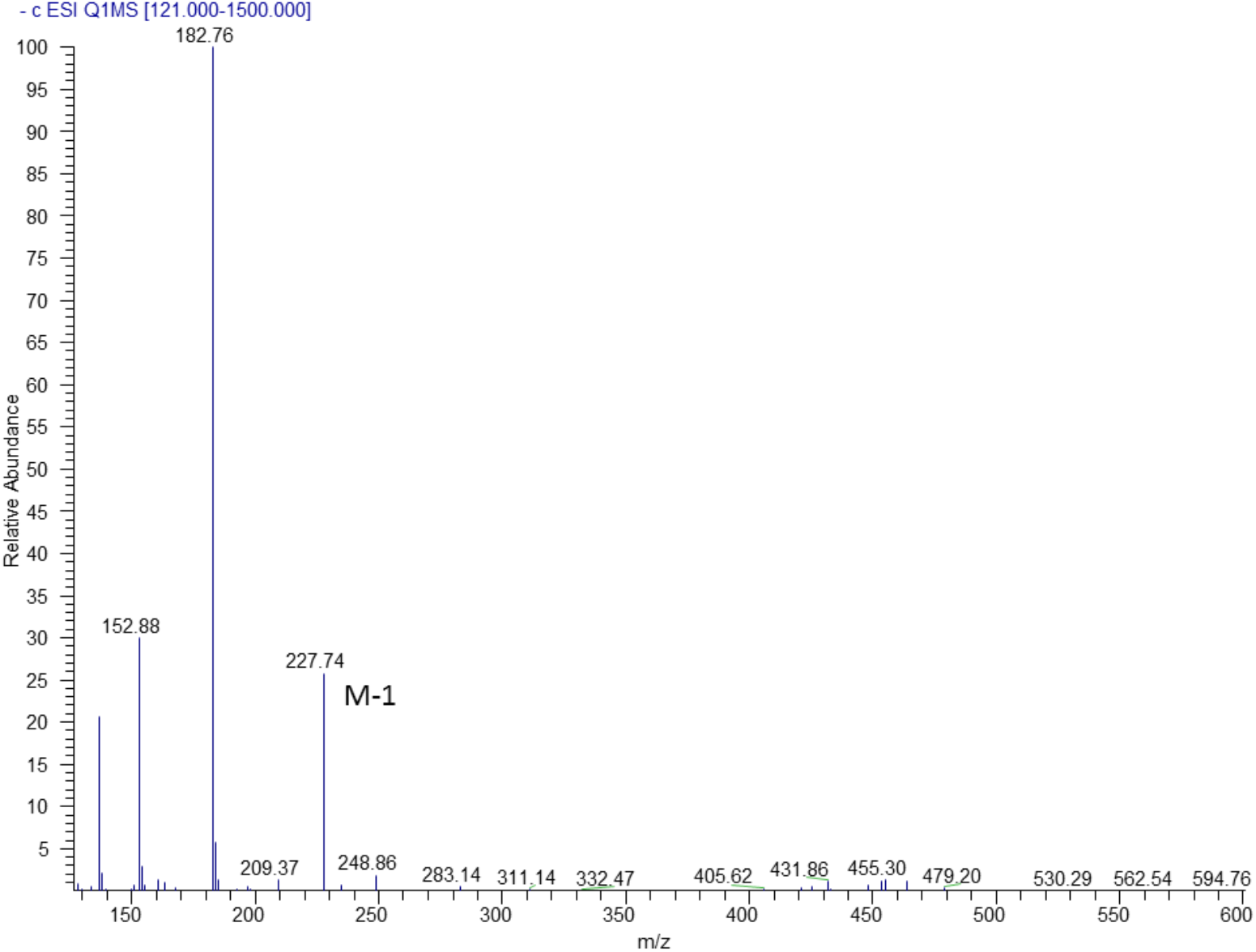
Characterization of compound IX, MS/MS spectrum of compound **IX.**

**Fig. S27-A.**
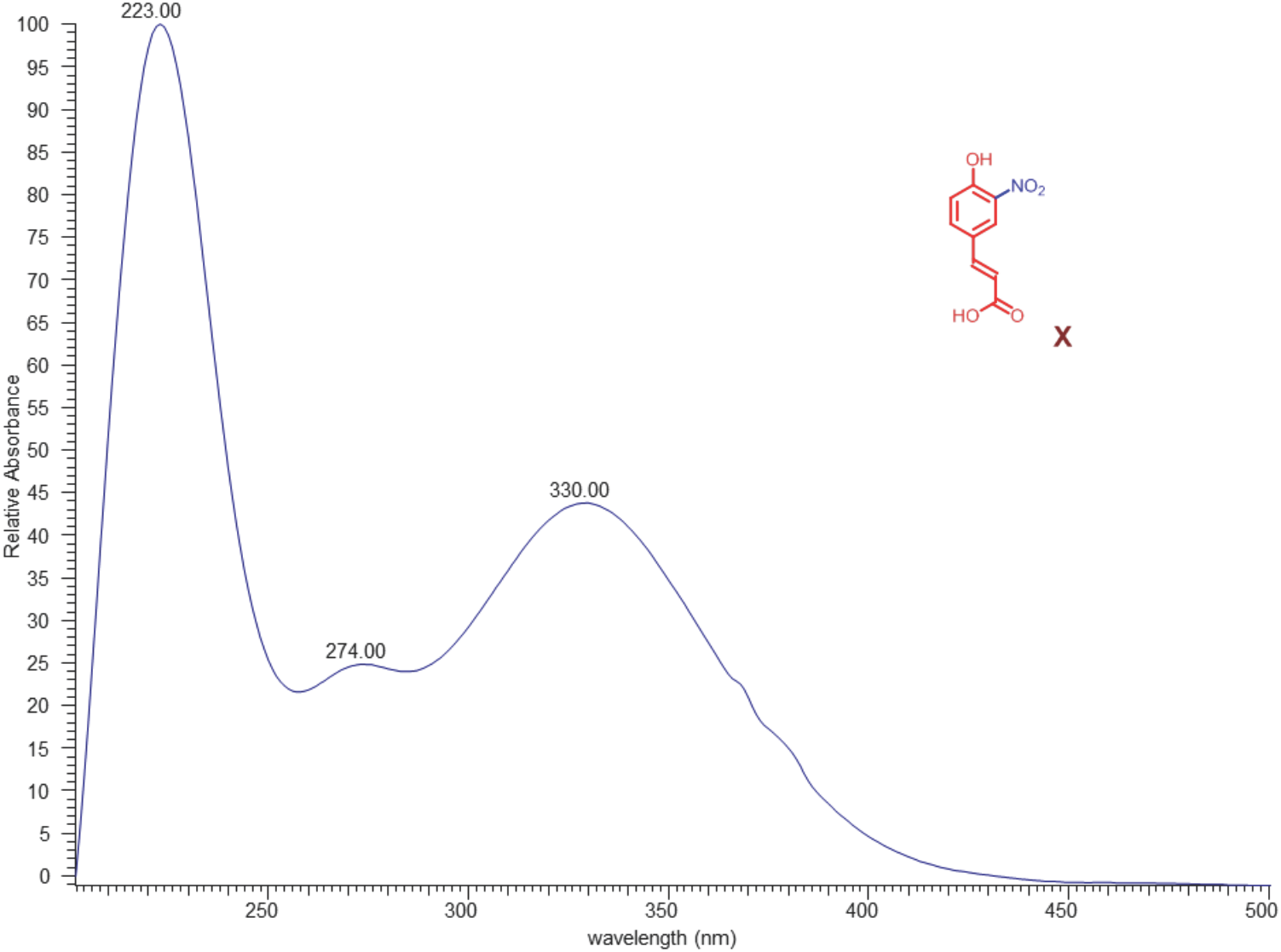
Characterization of compound X, UV spectrum of compound **X**.

**Fig. S27-B.**
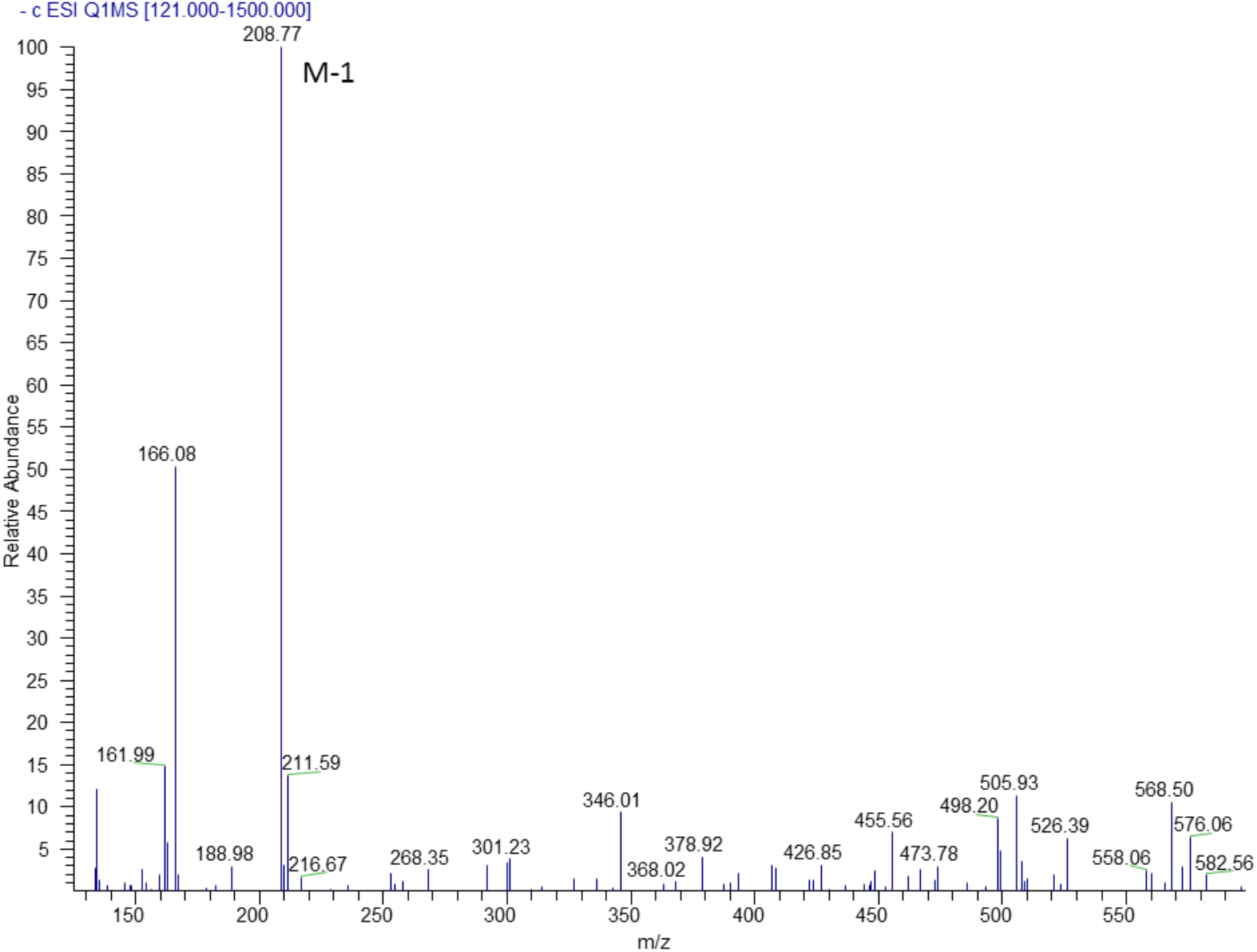
Characterization of compound X, MS/MS spectrum of compound **X**.

**Fig. S28-A.**
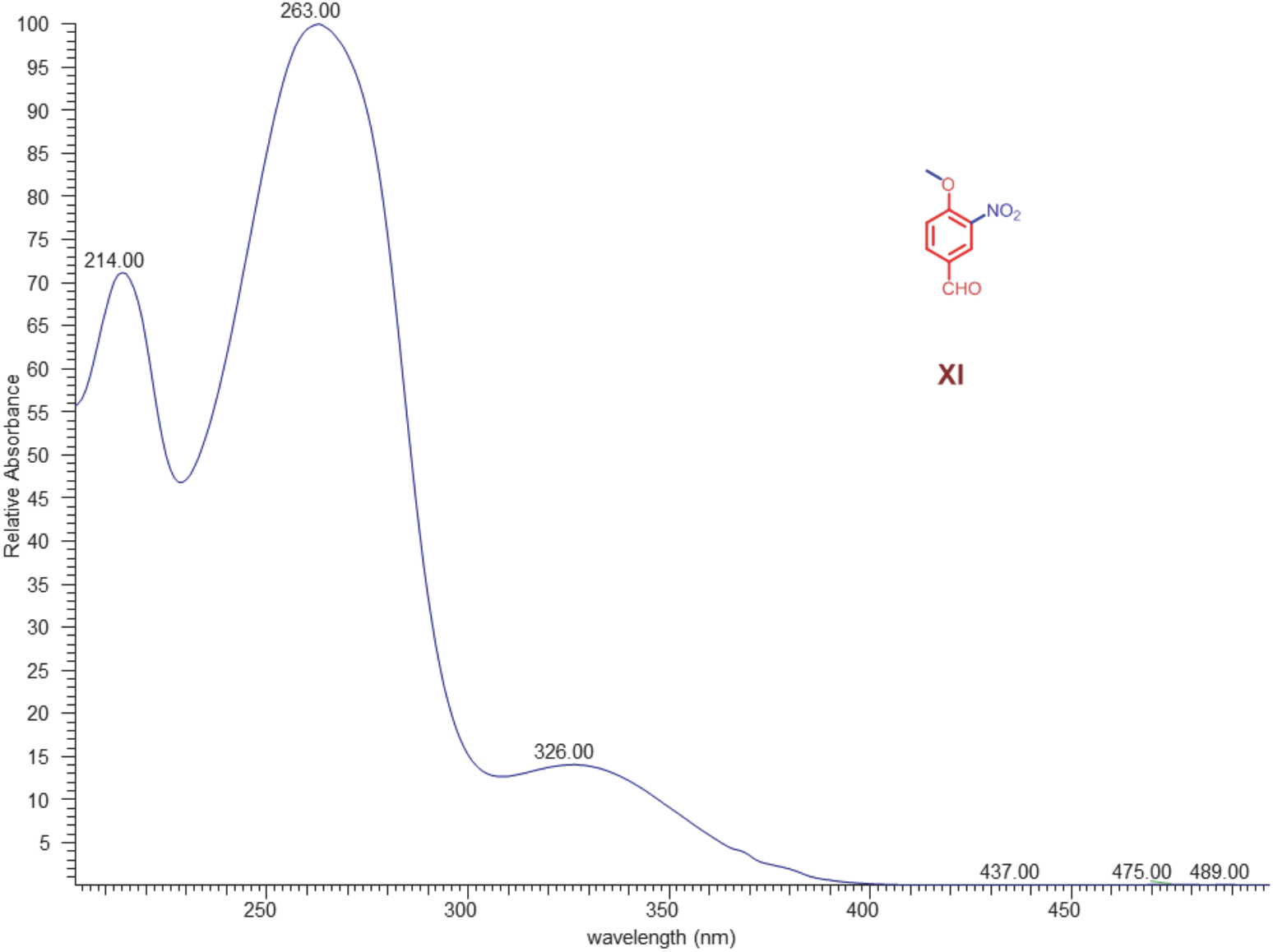
Characterization of compound XI, UV spectrum of compound **XI**.

**Fig. S28-B.**
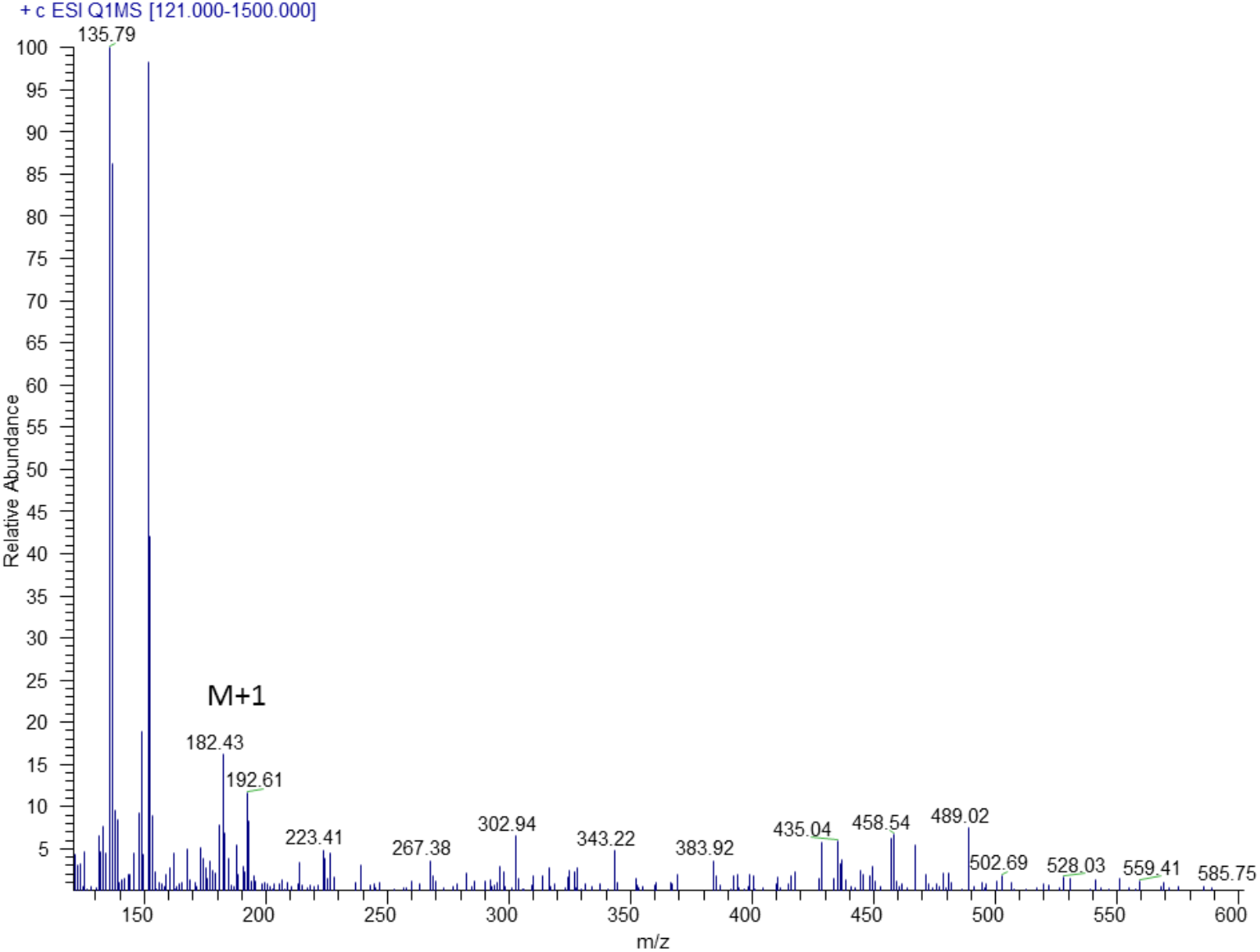
Characterization of compound XI, MS/MS spectrum of compound **XI**.

**Fig. S28-C.**
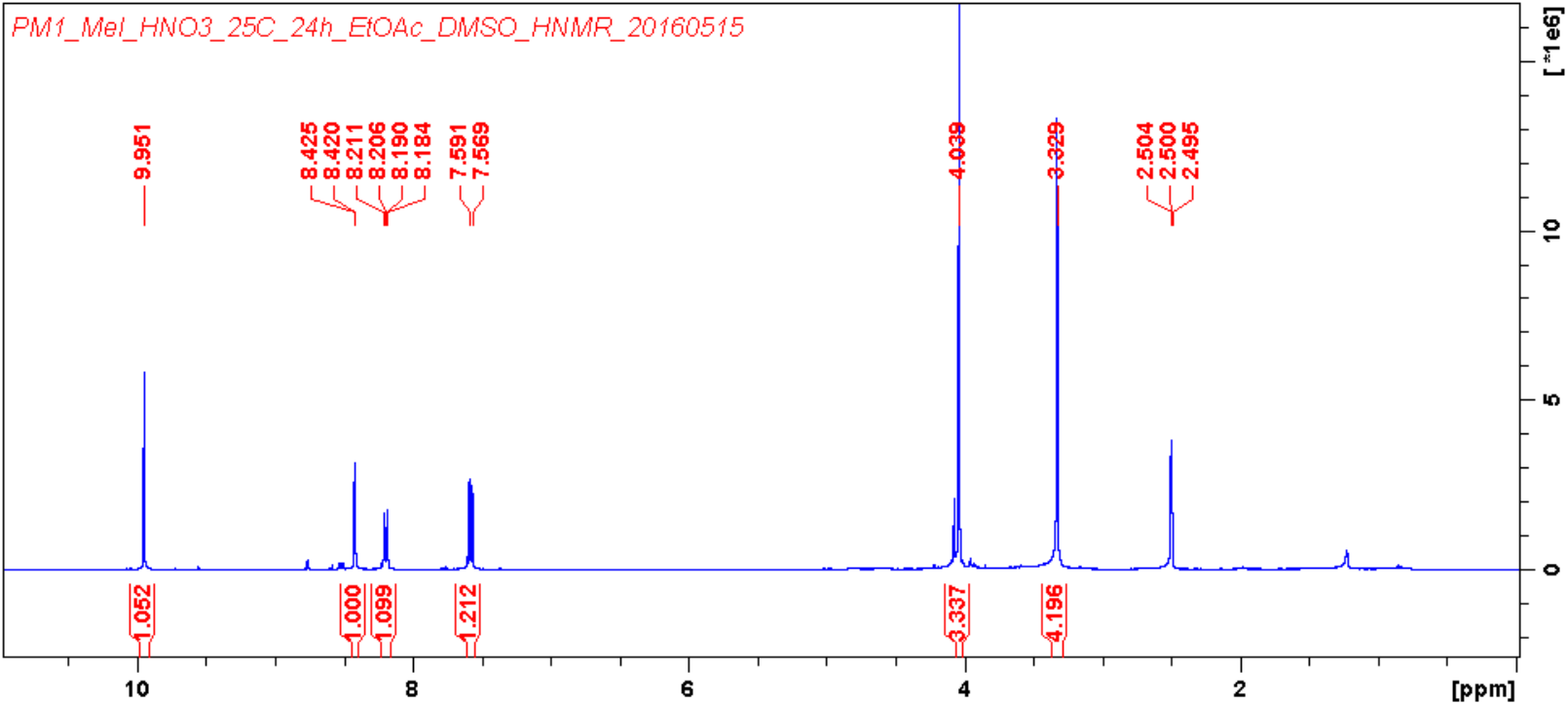
Characterization of compound XI, ^1^H NMR (400 MHz, CDCl_3_) spectrum of compound **XI**.

**Fig. S28-D.**
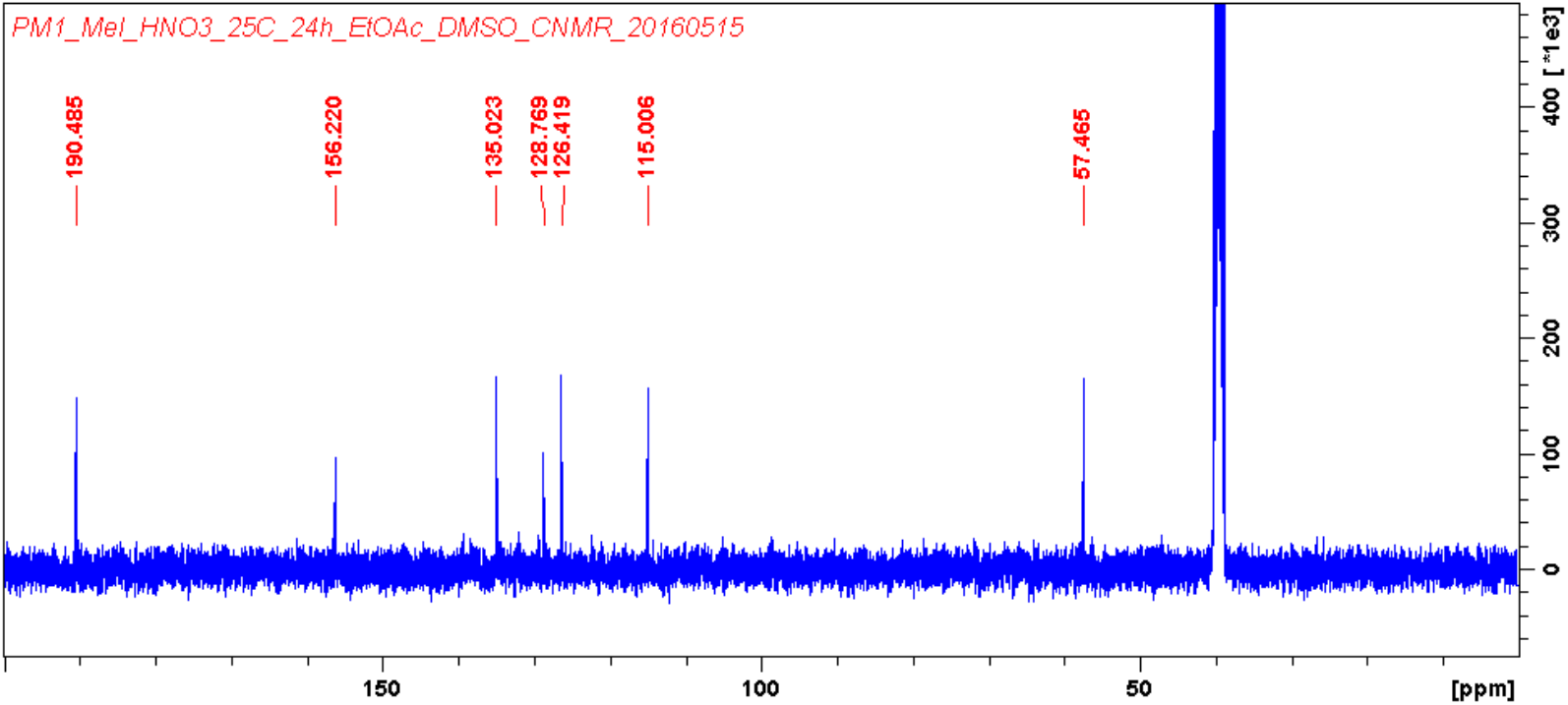
Characterization of compound XI, ^13^C NMR (100 MHz, CDCl_3_) spectrum of compound **XI**.

**Fig. S28-E.**
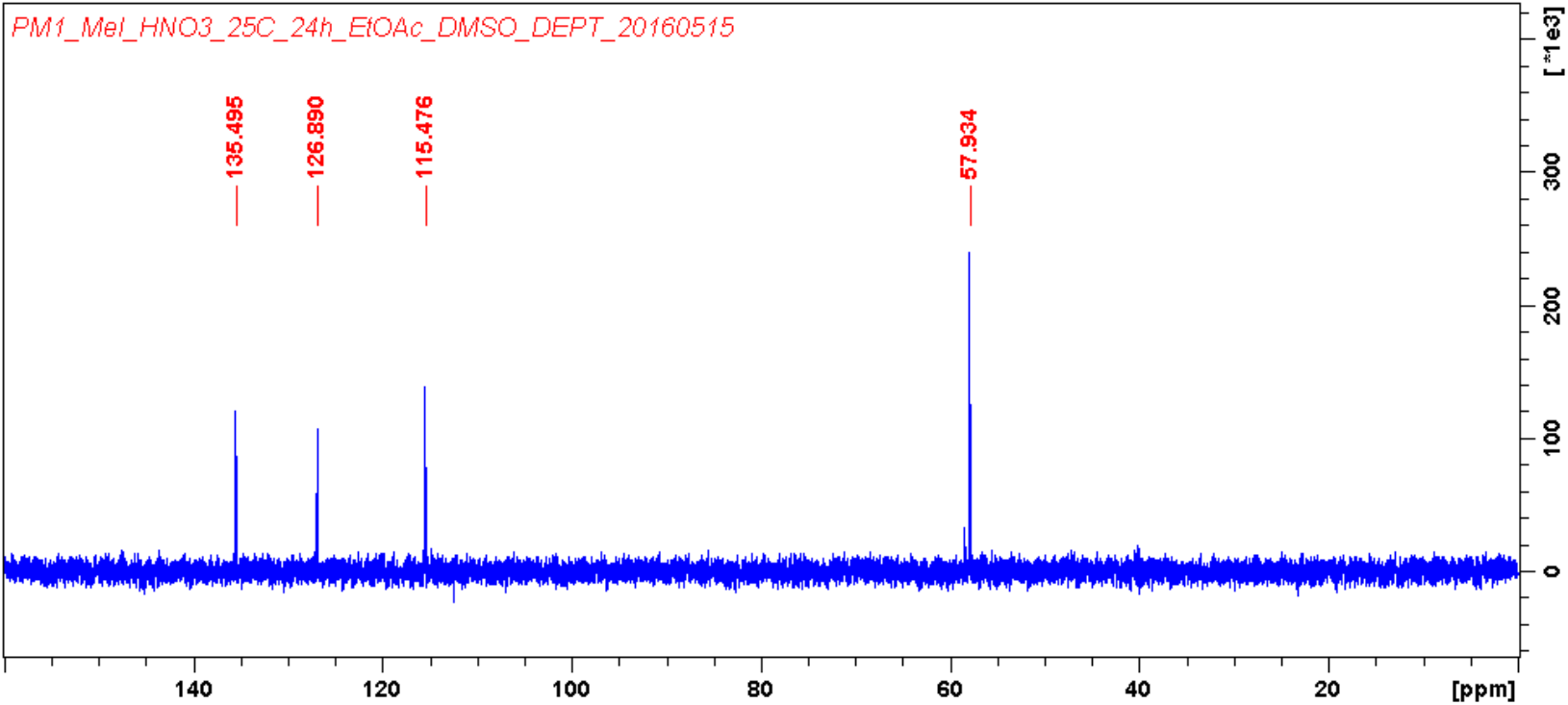
Characterization of compound XI, DEPT spectrum of compound **XI**.

**Fig. S28-F.**
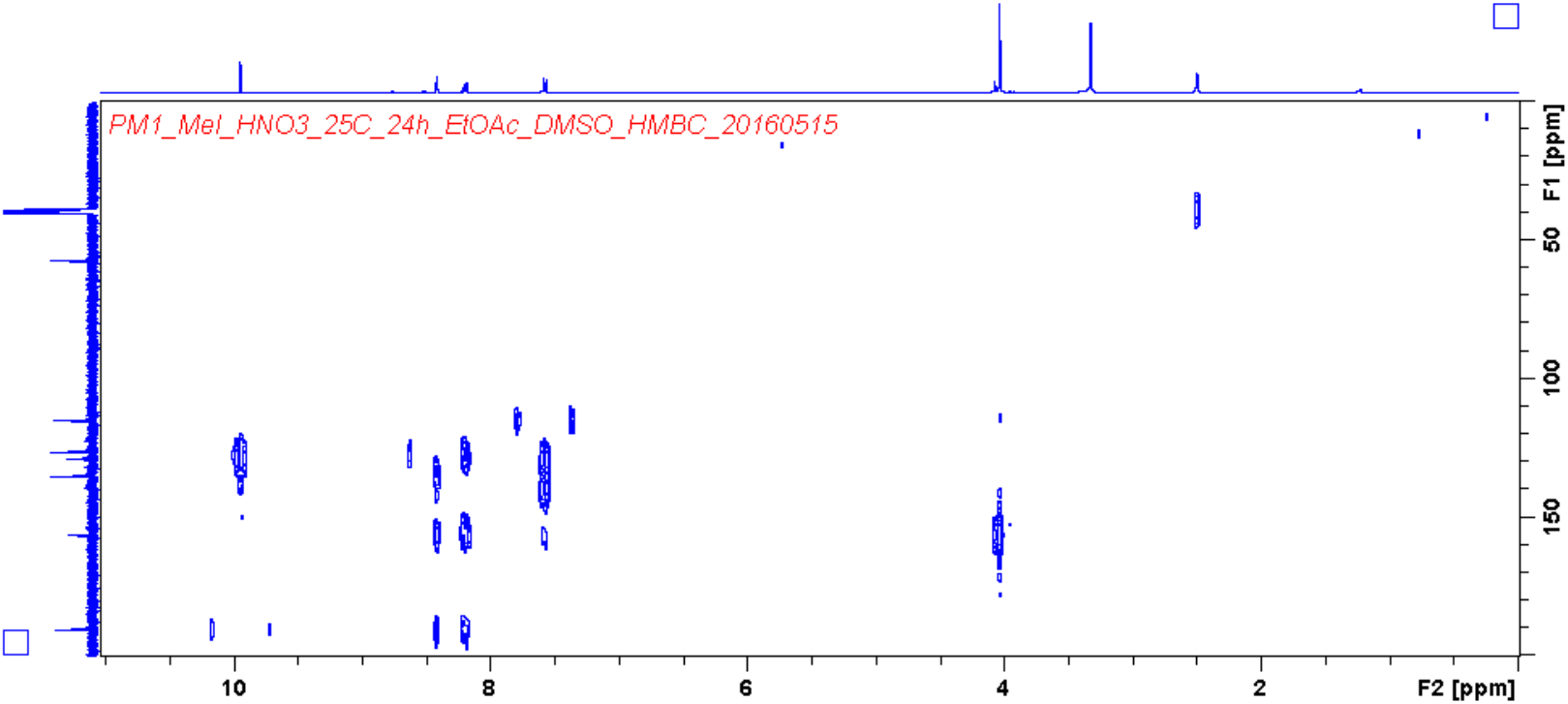
Characterization of compound XI, HMBC spectrum of compound **XI**.

**Fig. S28-G.**
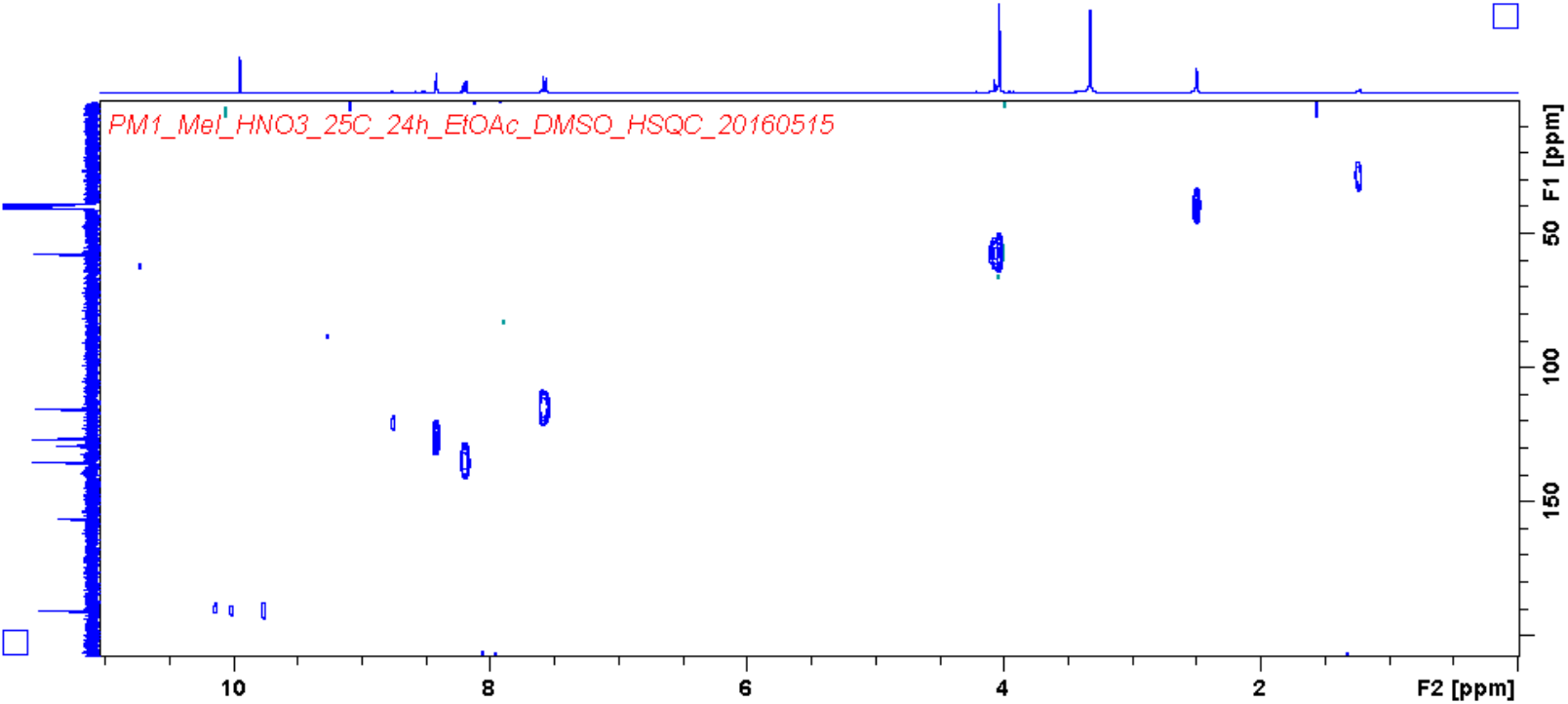
Characterization of compound XI, HSQC spectrum of compound **XI**.

**Fig. S29-A.**
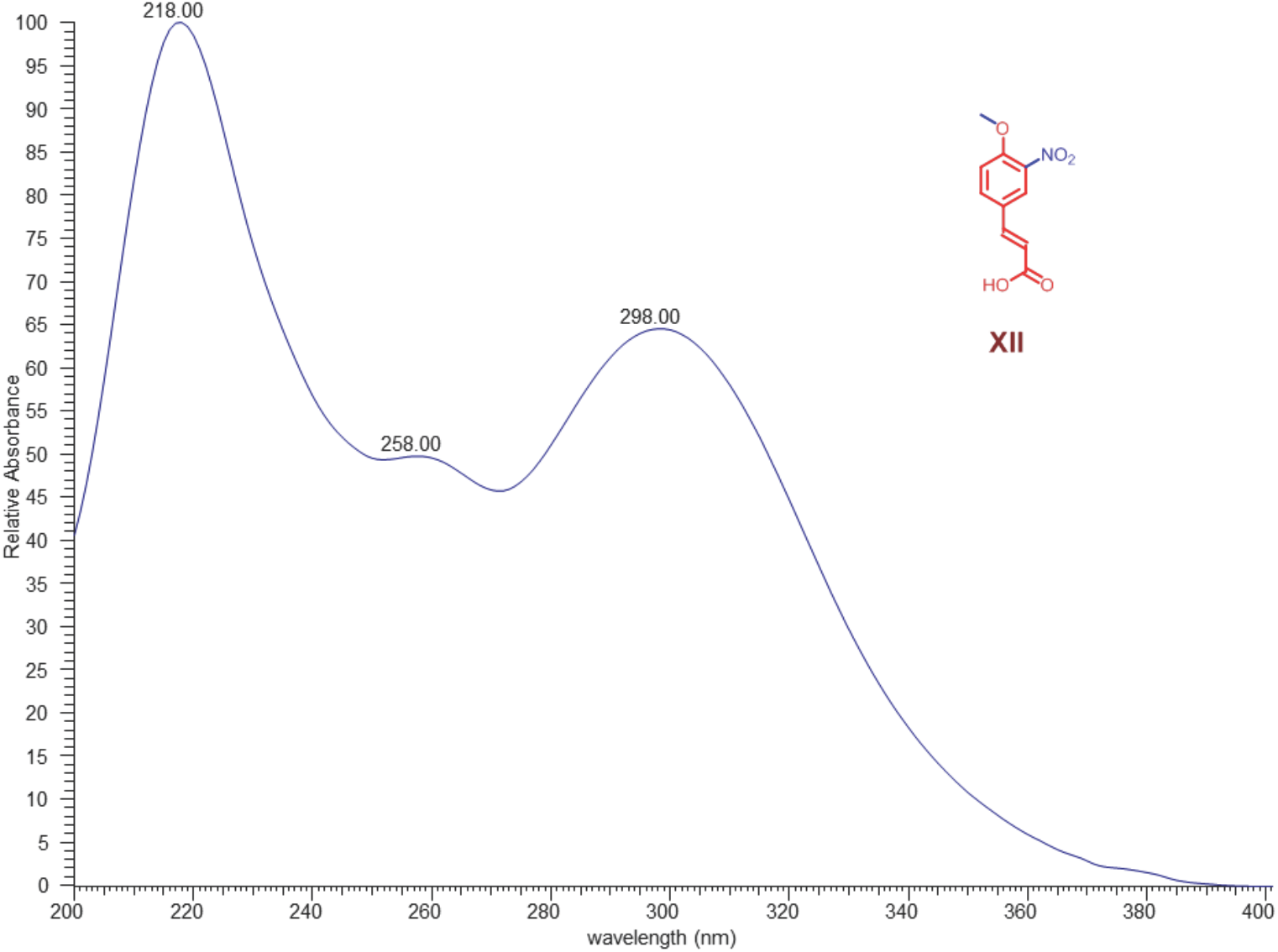
Characterization of compound XII, UV spectrum of compound **XII**.

**Fig. S29-B.**
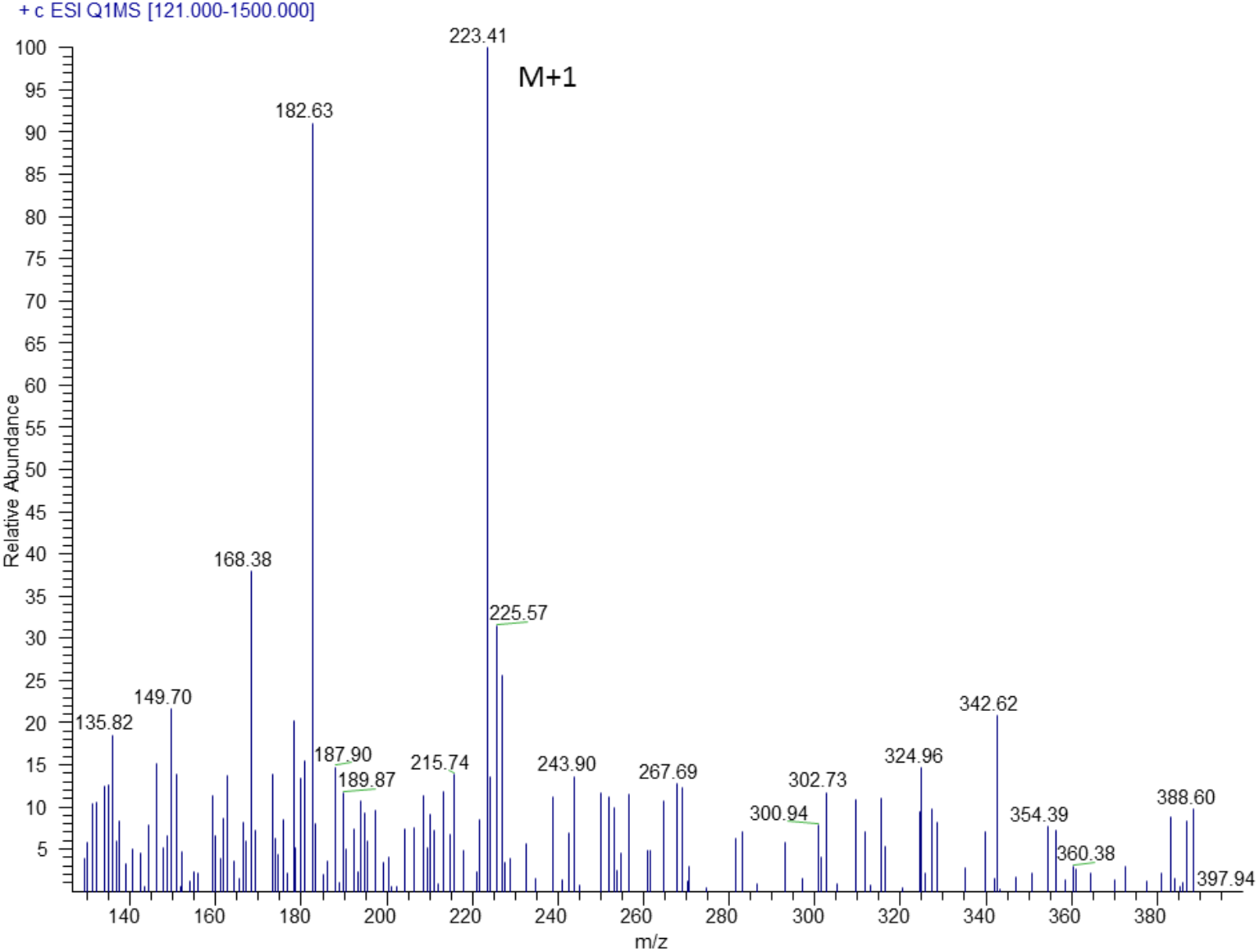
Characterization of compound XII, MS/MS spectrum of compound **XII**.

**Fig. S30-A.**
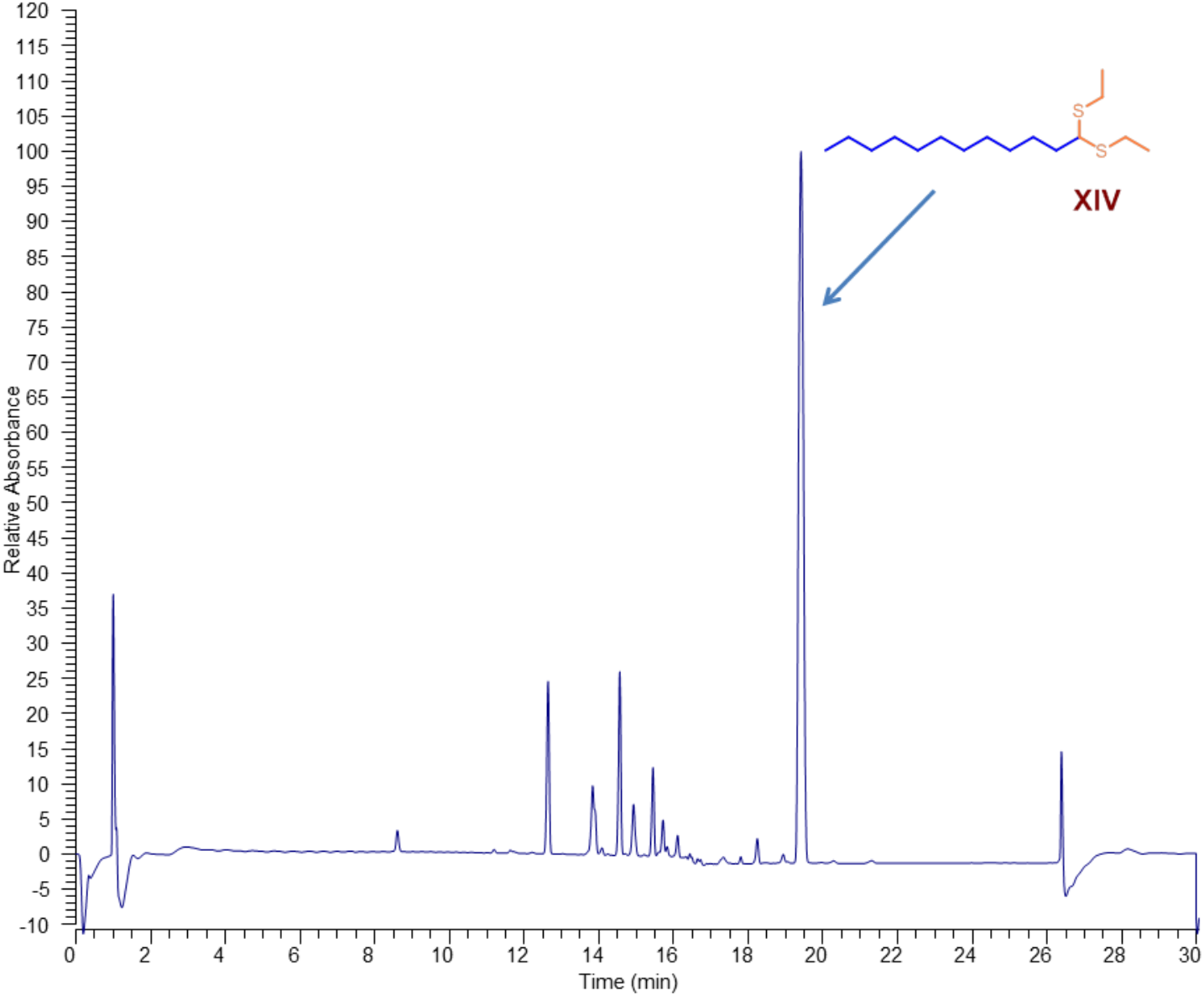
Characterization of compound XIV, HPLC-UV (PDA) spectrum of compound **XIV**.

**Fig. S30-B.**
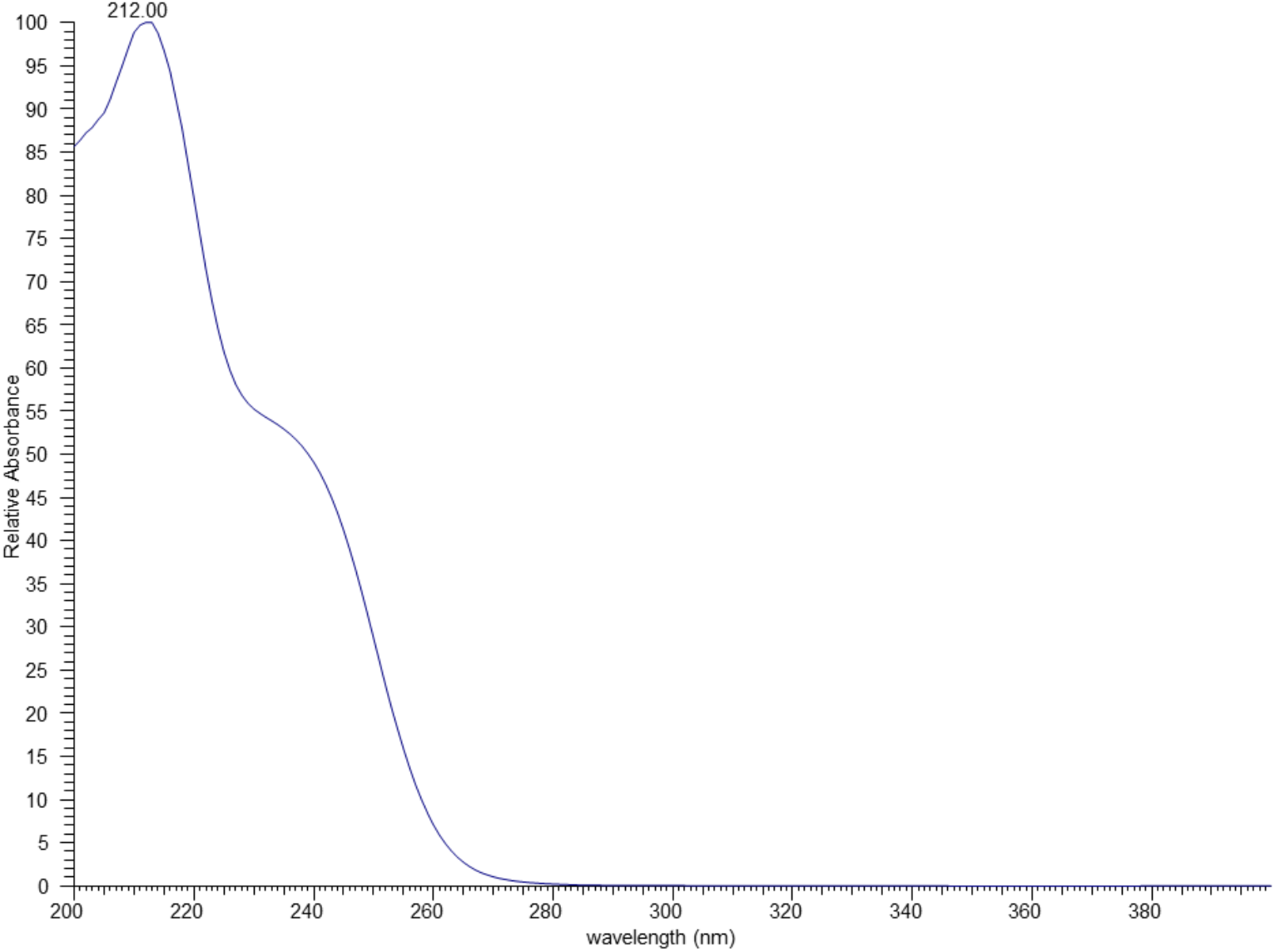
Characterization of compound XIV, UV spectrum of compound **XIV**.

**Fig. S30-C.**
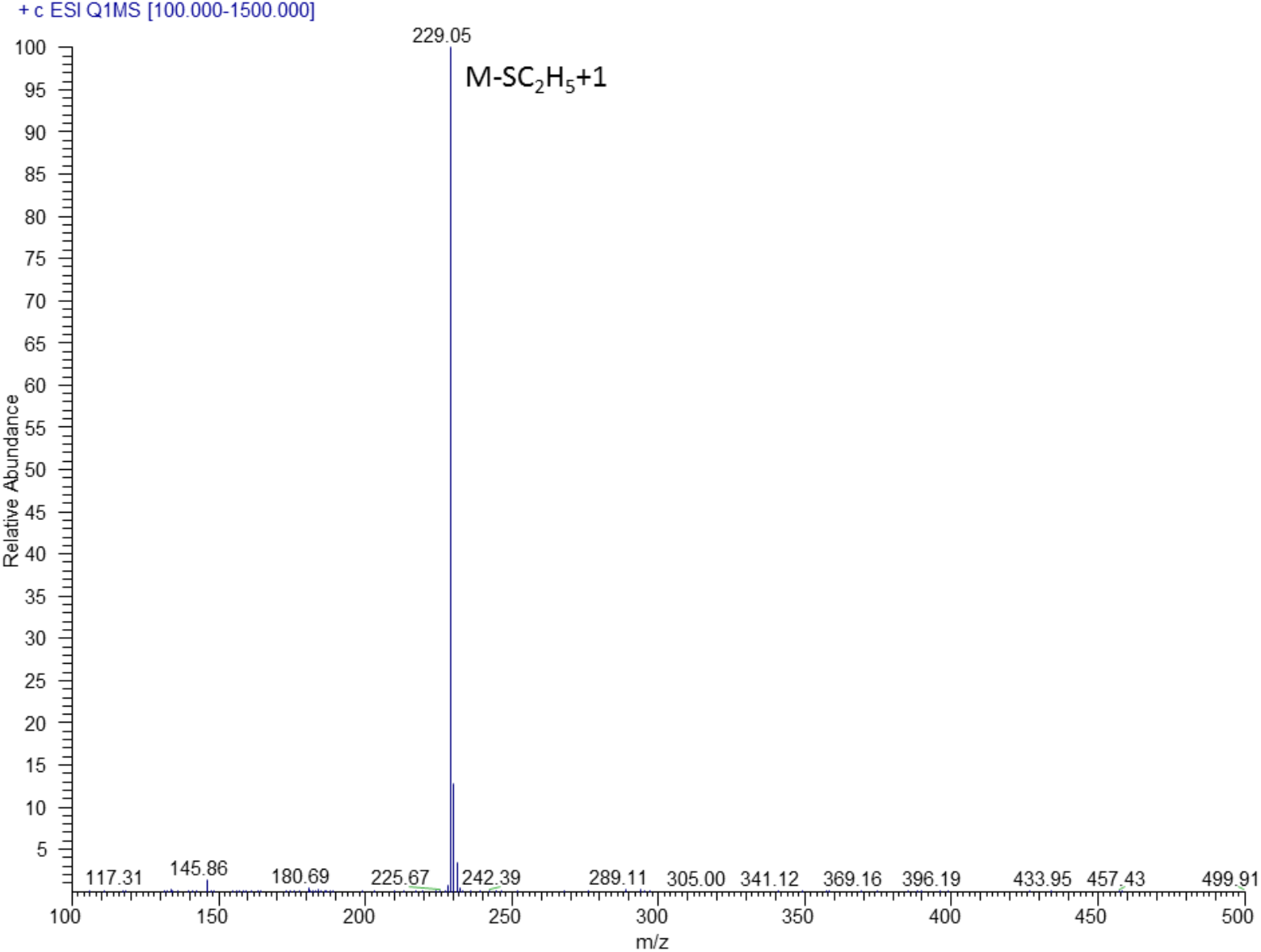
Characterization of compound XIV, MS/MS spectrum of compound **XIV**.

**Fig. S30-D.**
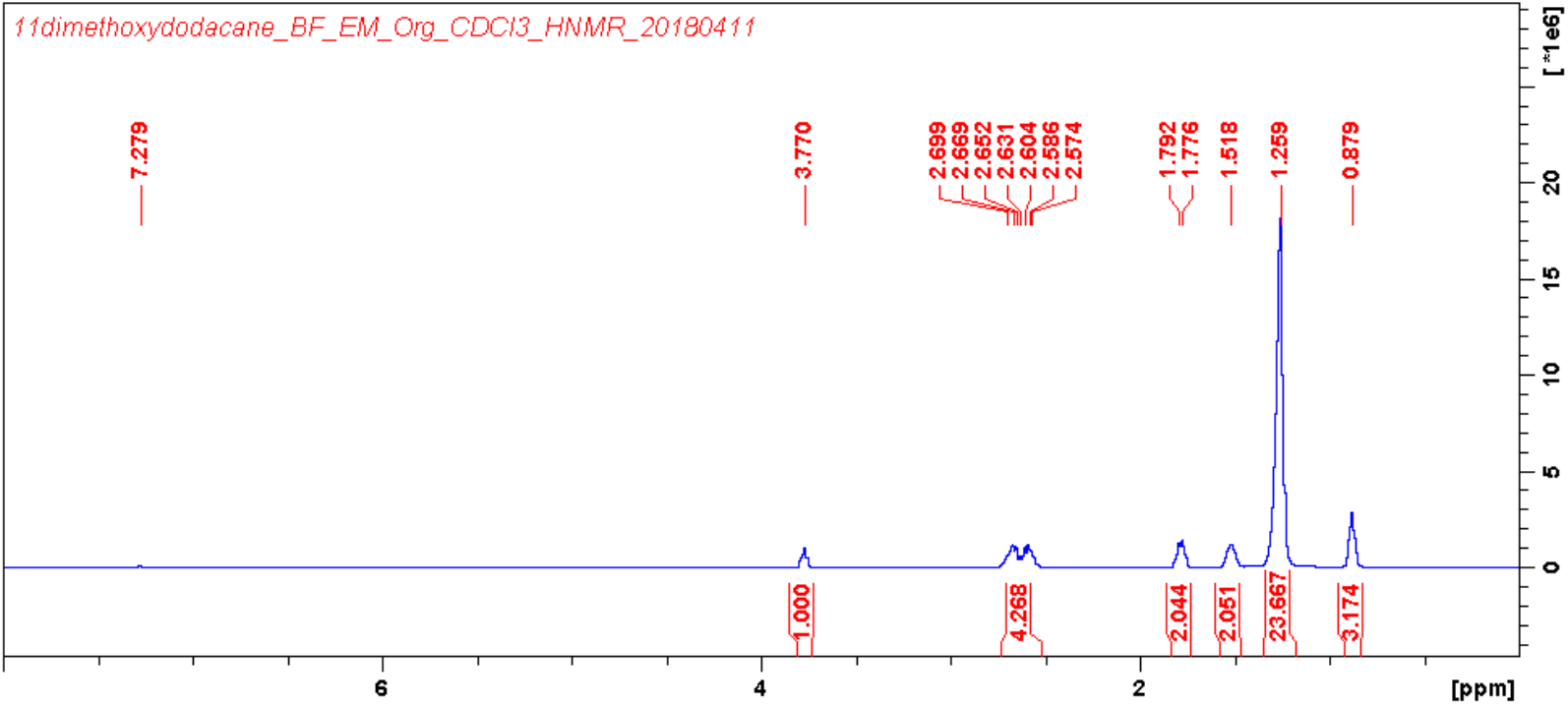
Characterization of compound XIV, ^1^H NMR (400 MHz, CDCl_3_) spectrum of compound **XIV**.

**Fig. S30-E.**
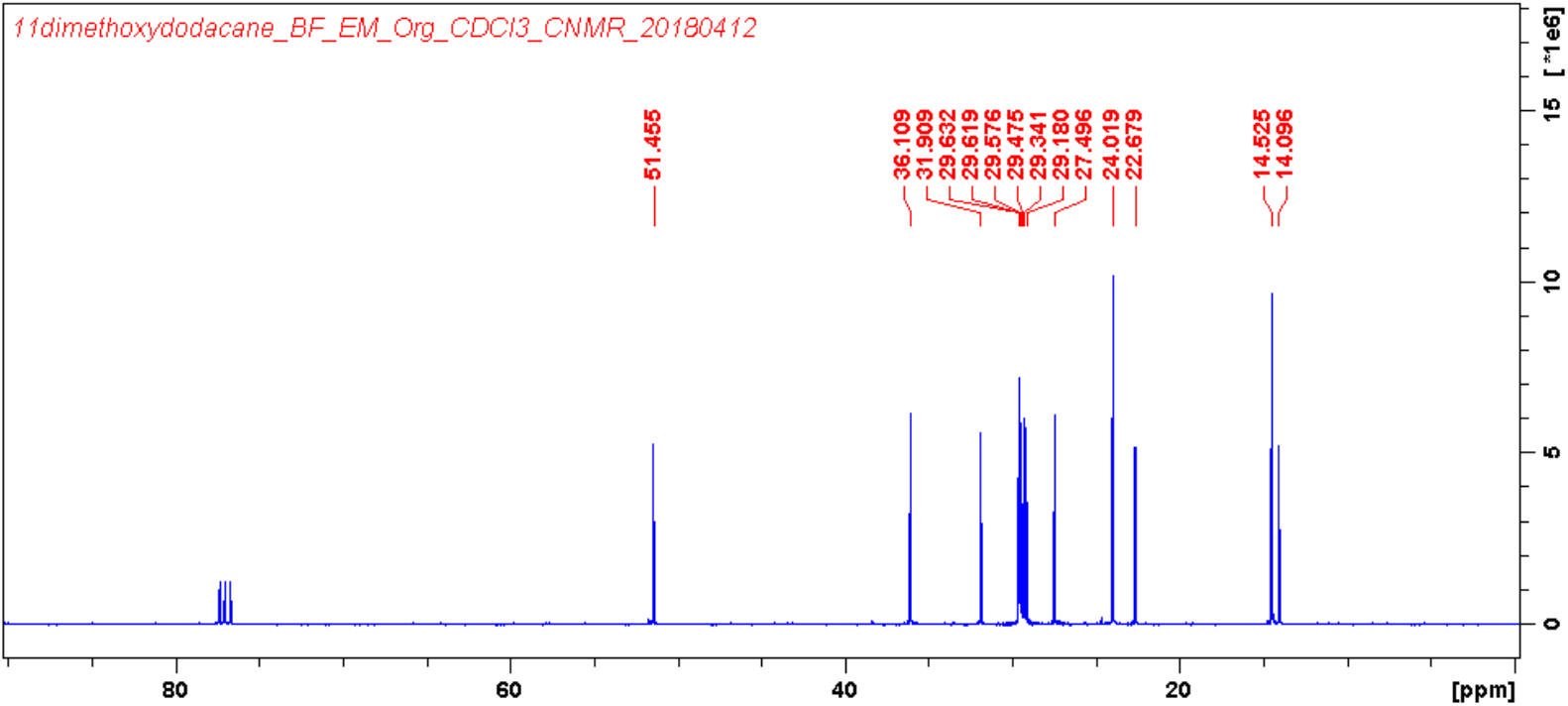
Characterization of compound XIV, ^13^C NMR (100 MHz, CDCl_3_) spectrum of compound **XIV**.

**Fig. S31.**
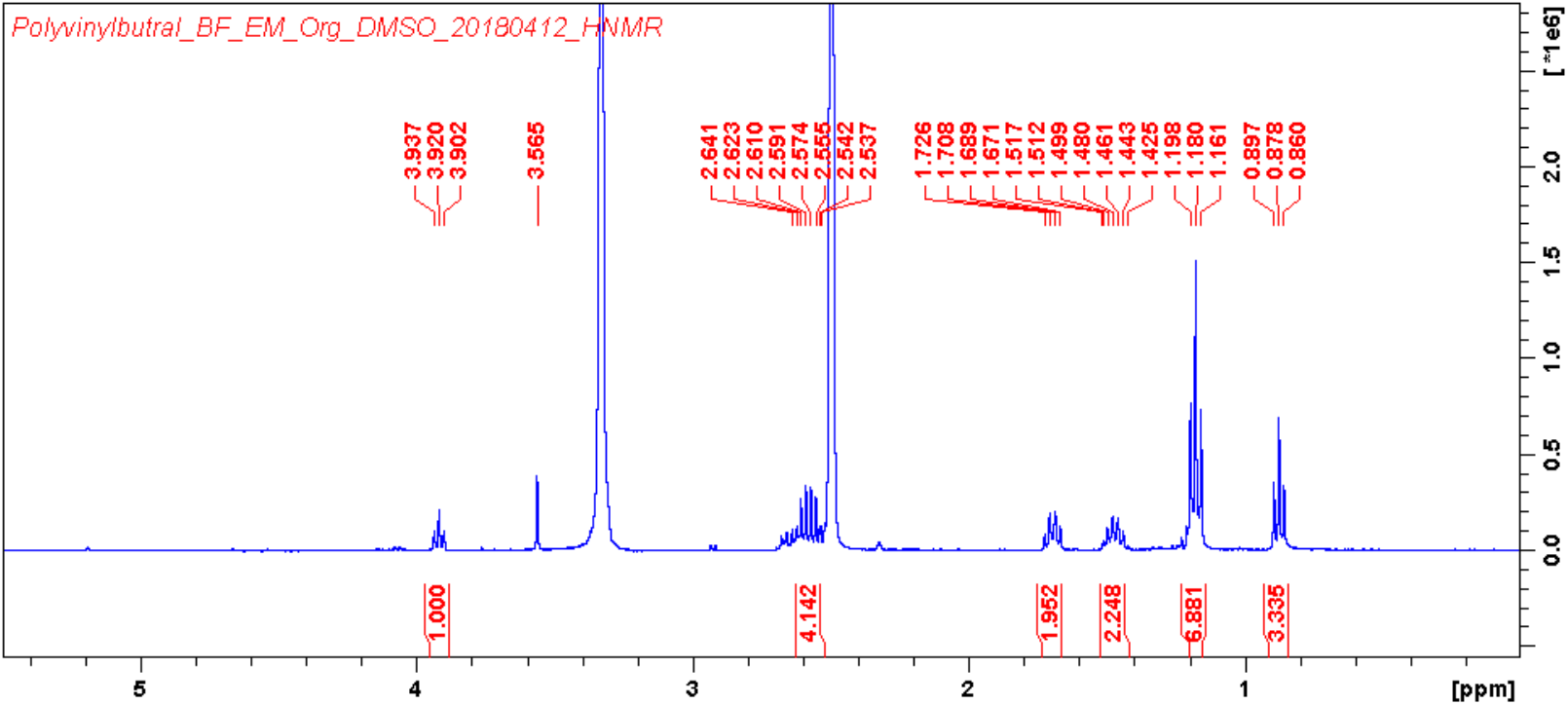
Characterization of compound XV, ^1^H NMR (400 MHz, DMSO-*d*_*6*_) spectrum of compound **XV**.

**Fig. S32.**
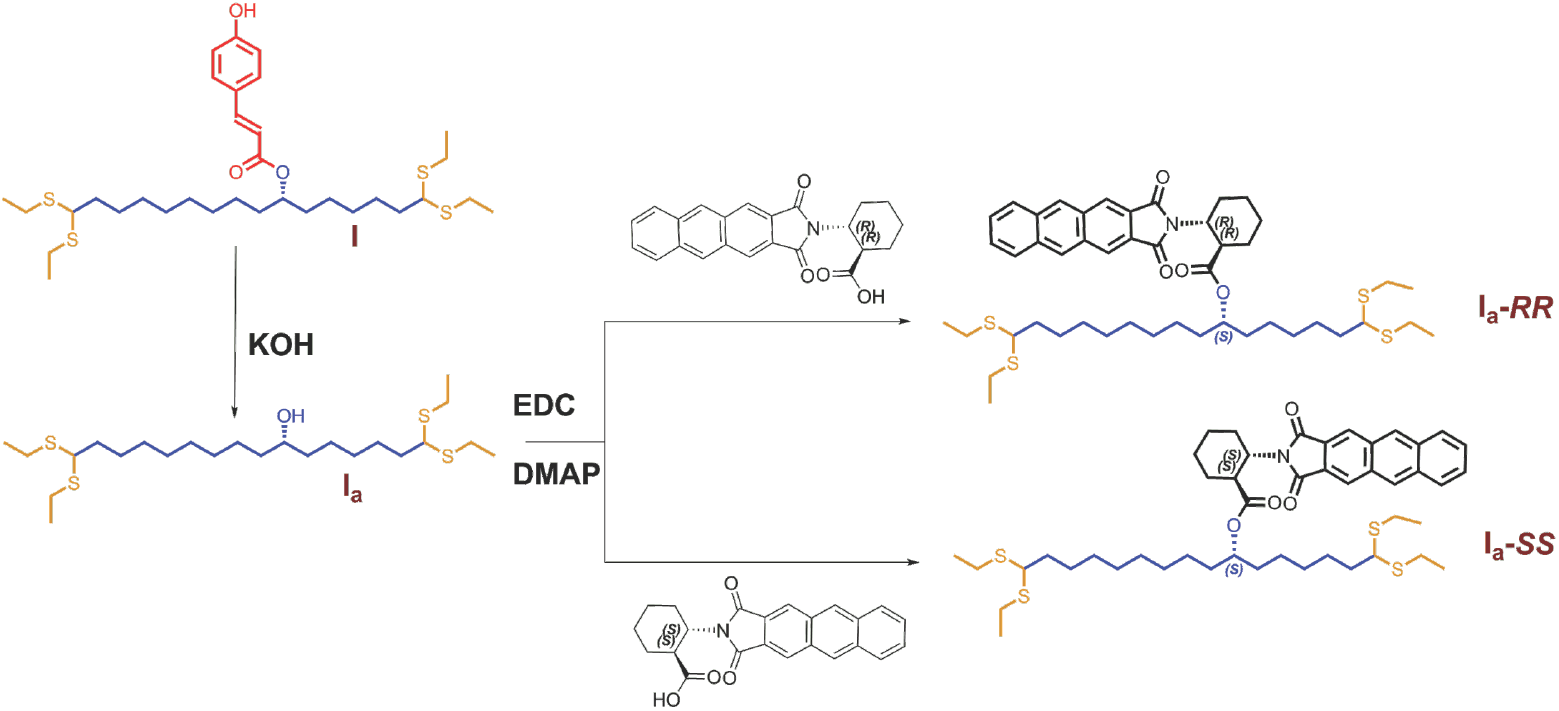
Derivatization compound I_a_ with (1R,2R)-(-)-2-(2,3-anthracenedicarboximide) cyclohexane carboxylic acids and (1S,2S)-(+)-2-(2,3-anthracenedicarboximide) cyclohexane carboxylic acids.

**Fig. S33.**
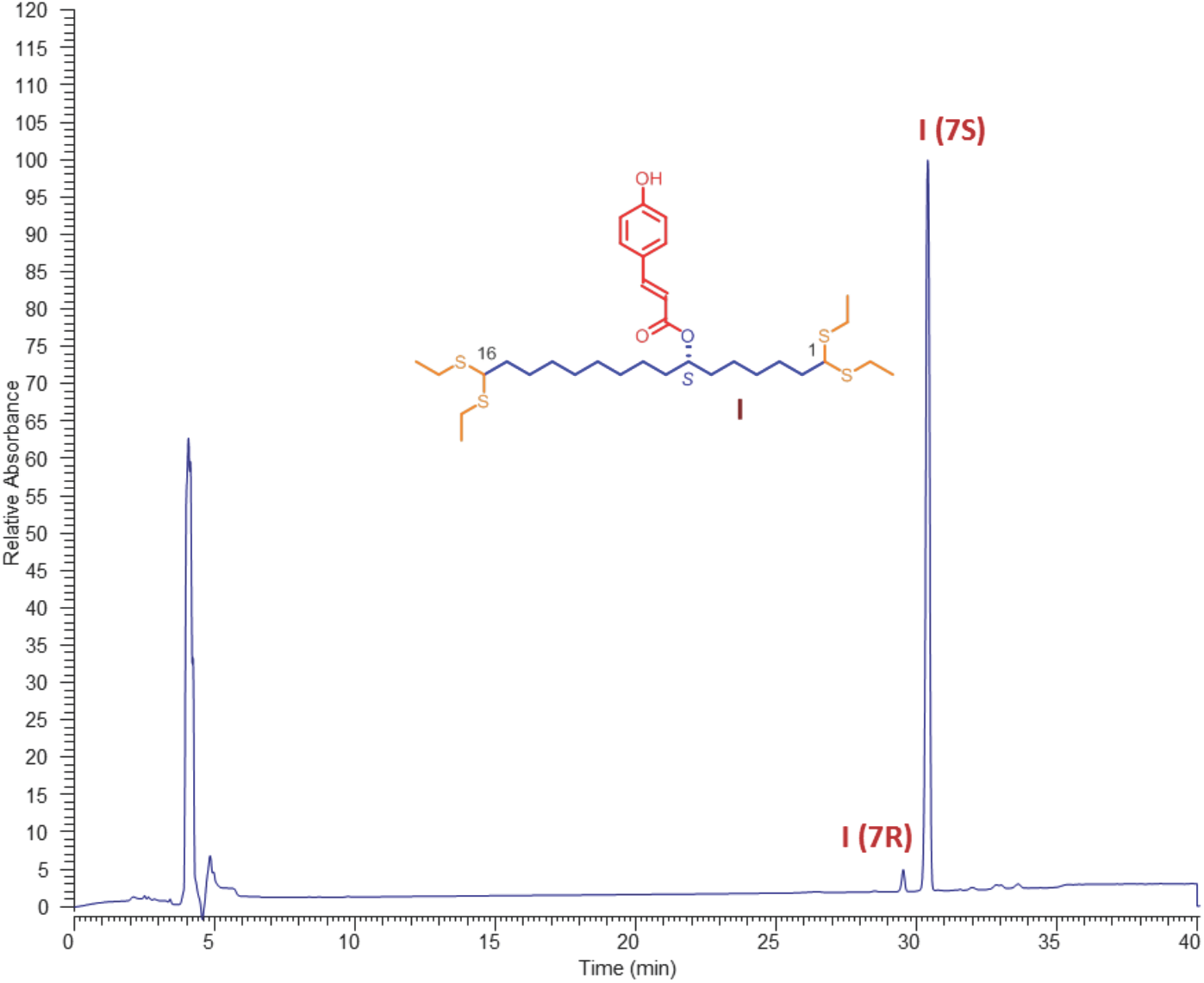
HPLC-UV (PDA) chromatogram of enantiomeric resolution of I on a Lux 3 μ Cellulose-4, 250 × 4.6 mm chiral column.

**Fig. S34.**
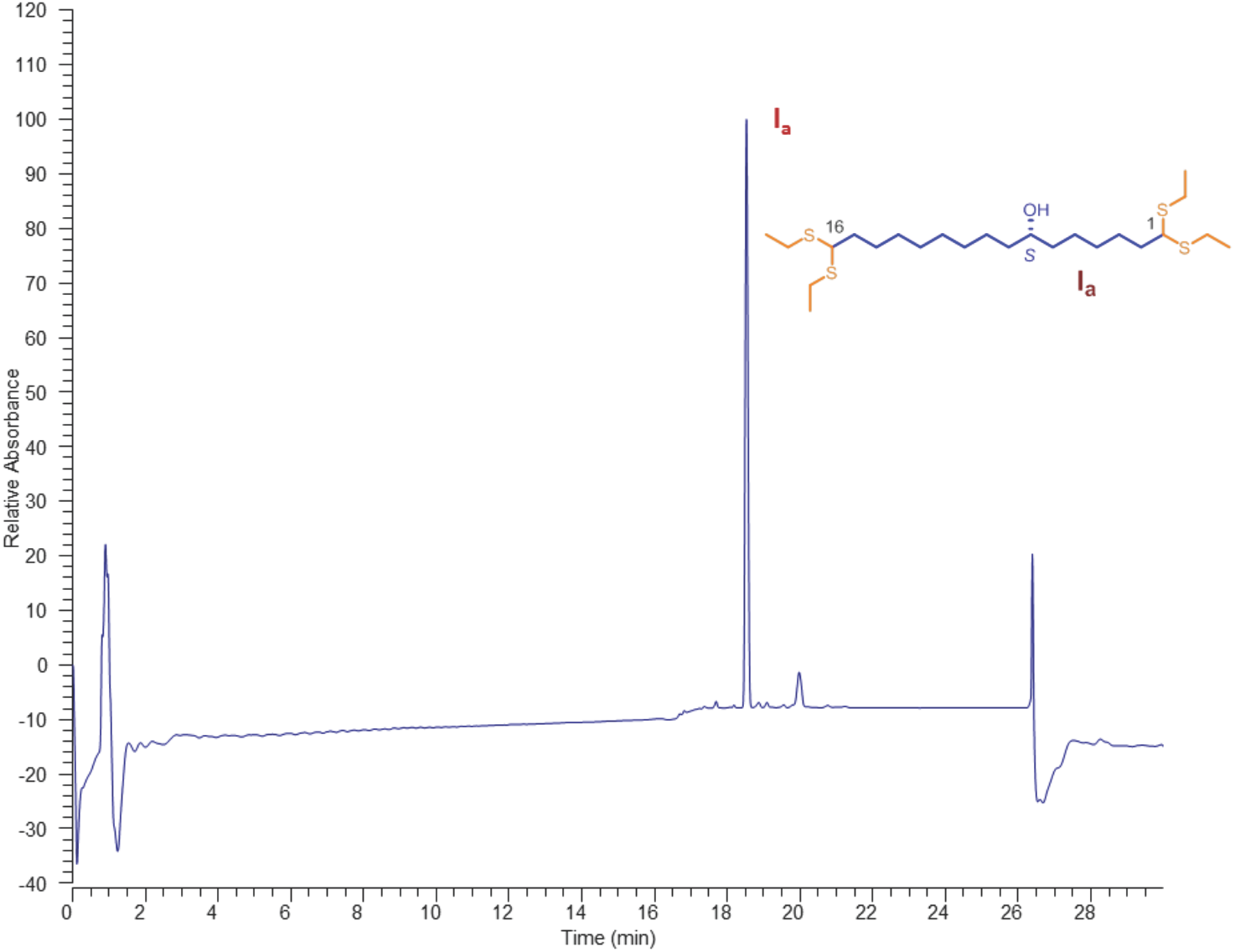
HPLC-UV (PDA) chromatogram of compound I_a_.

**Fig. S35-A.**
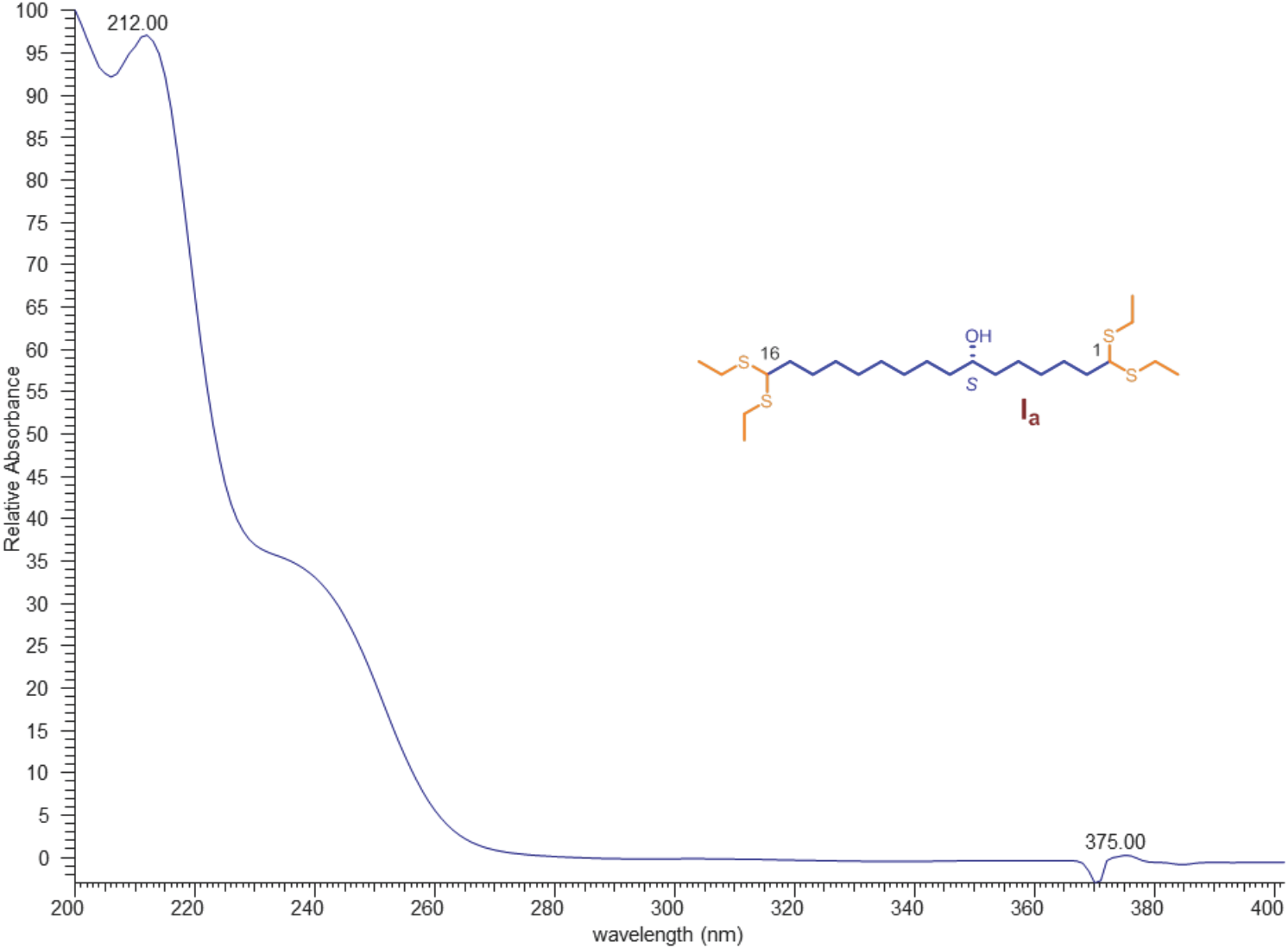
Characterization of compound I_a_, UV spectrum of compound **I_a_**.

**Fig. S35-B.**
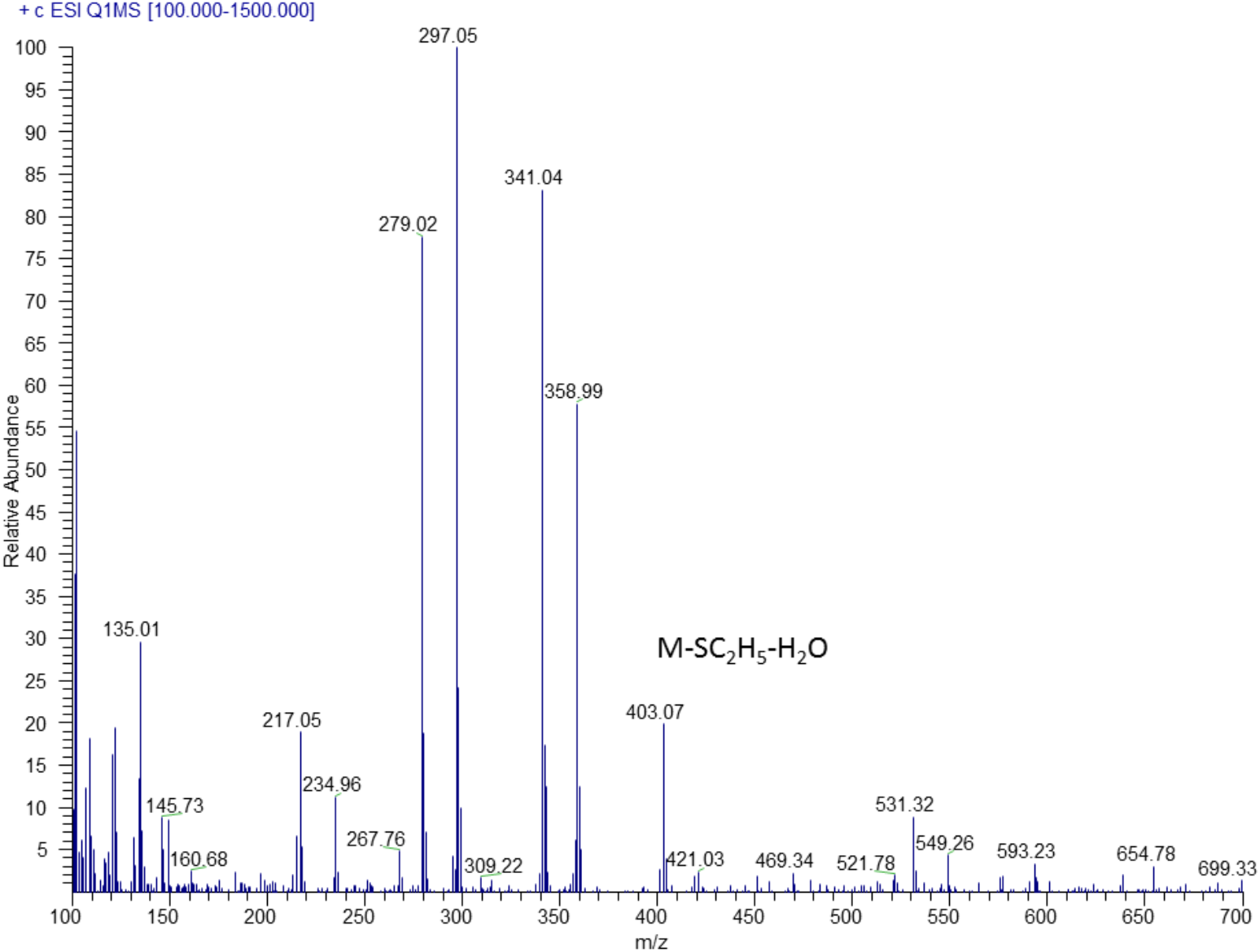
Characterization of compound I_a_, MS/MS spectrum of compound **I_a_**.

**Fig. S35-C.**
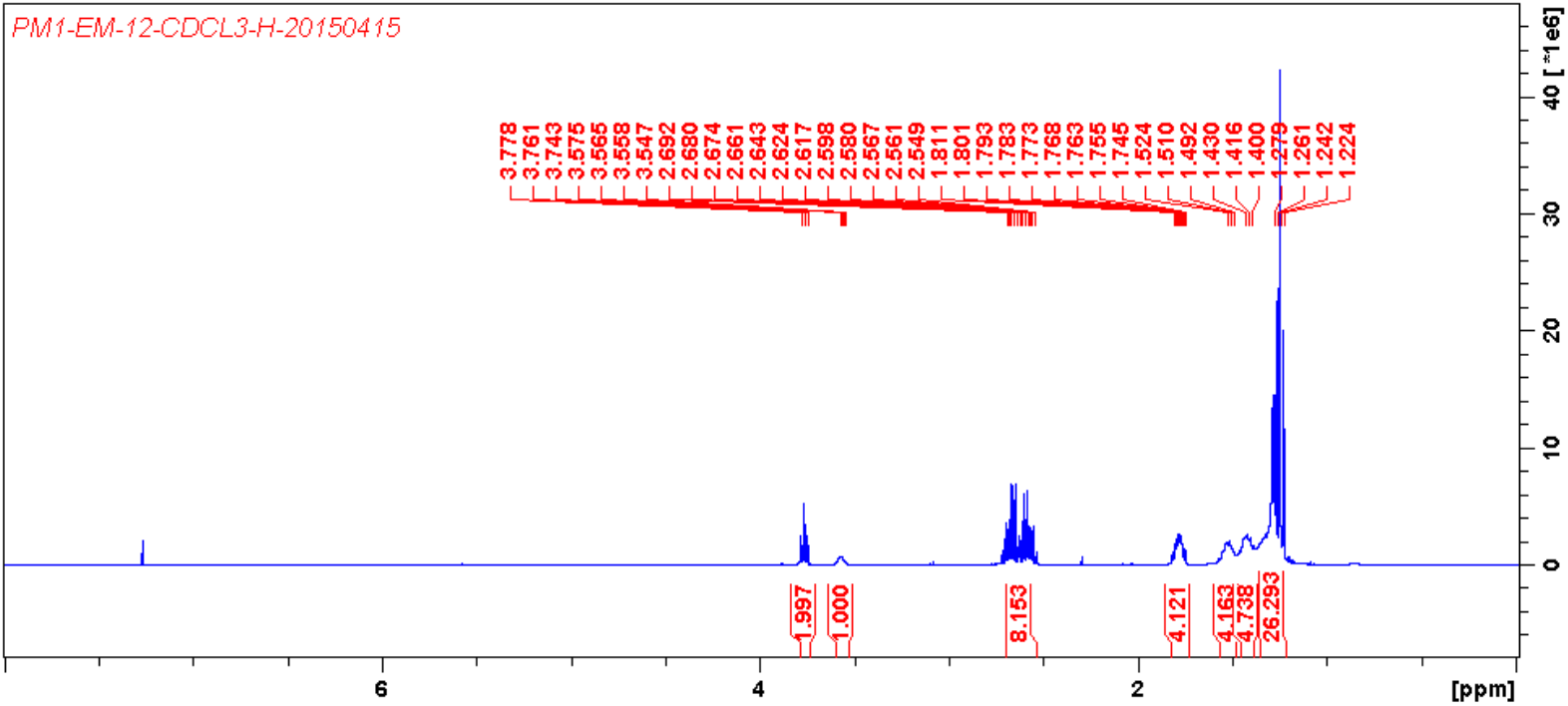
Characterization of compound I_a_, ^1^H NMR (400 MHz, CDCl_3_) spectrum of compound **I_a_**.

**Fig. S35-D.**
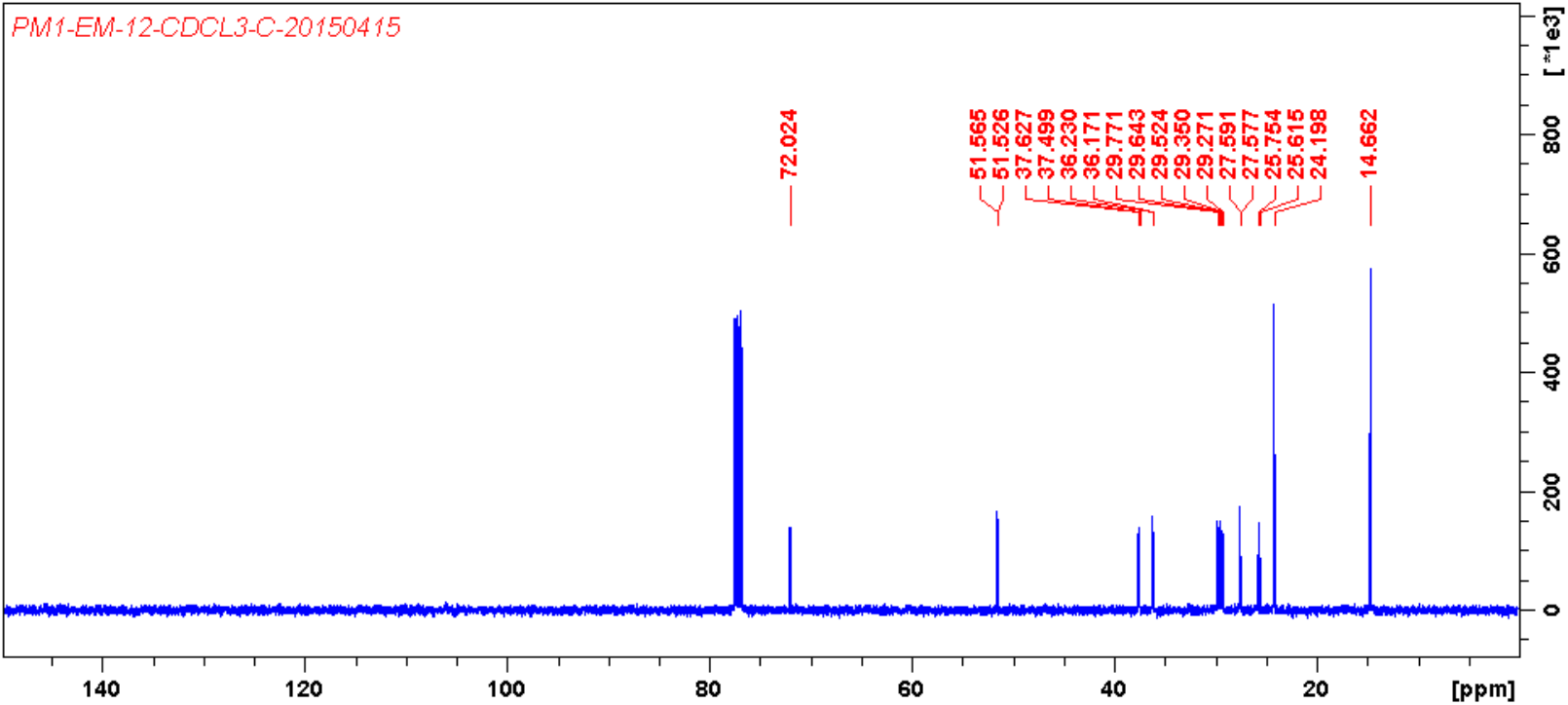
Characterization of compound I_a_, ^13^C NMR (100 MHz, CDCl_3_) spectrum of compound **I_a_**.

**Fig. S35-E.**
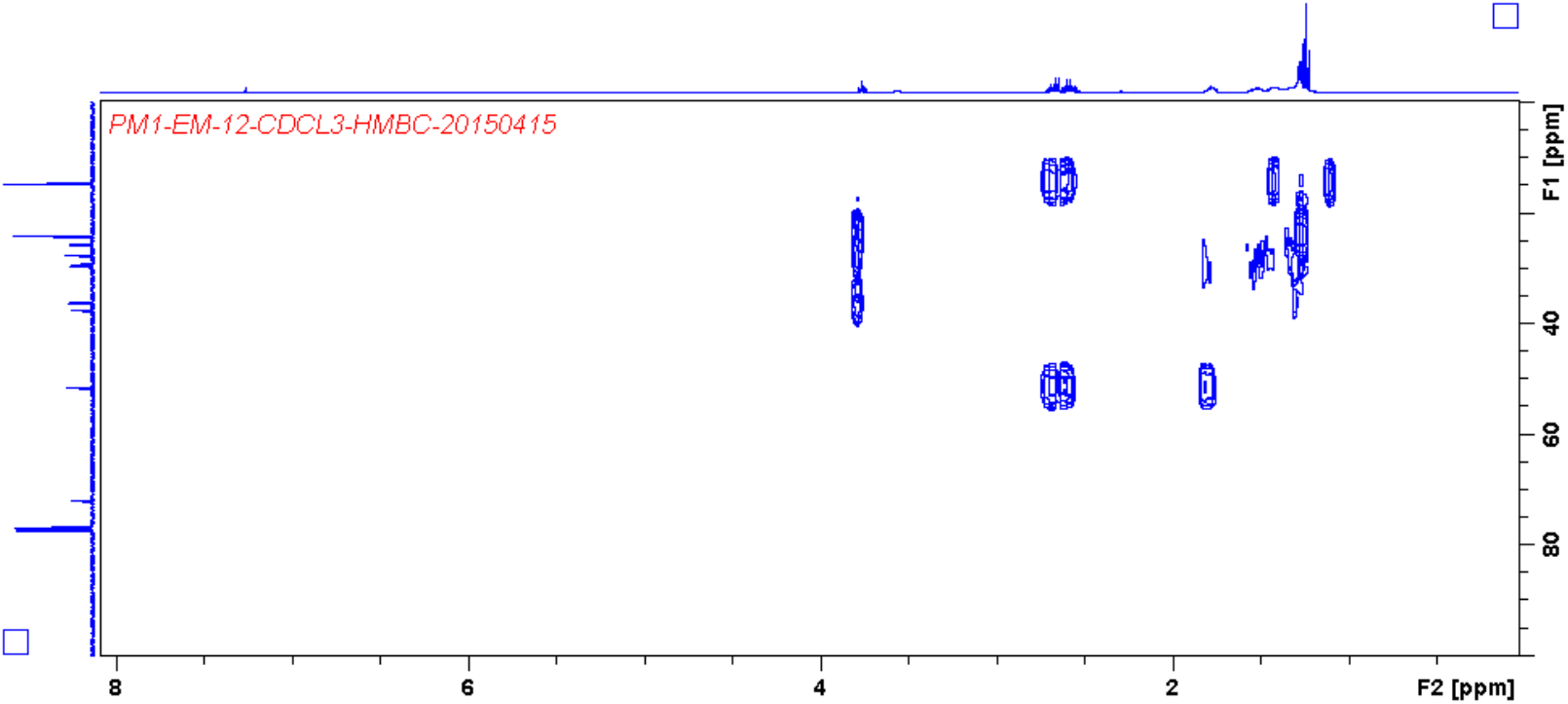
Characterization of compound I_a_, HMBC spectrum of compound **I_a_**.

**Fig. S35-F.**
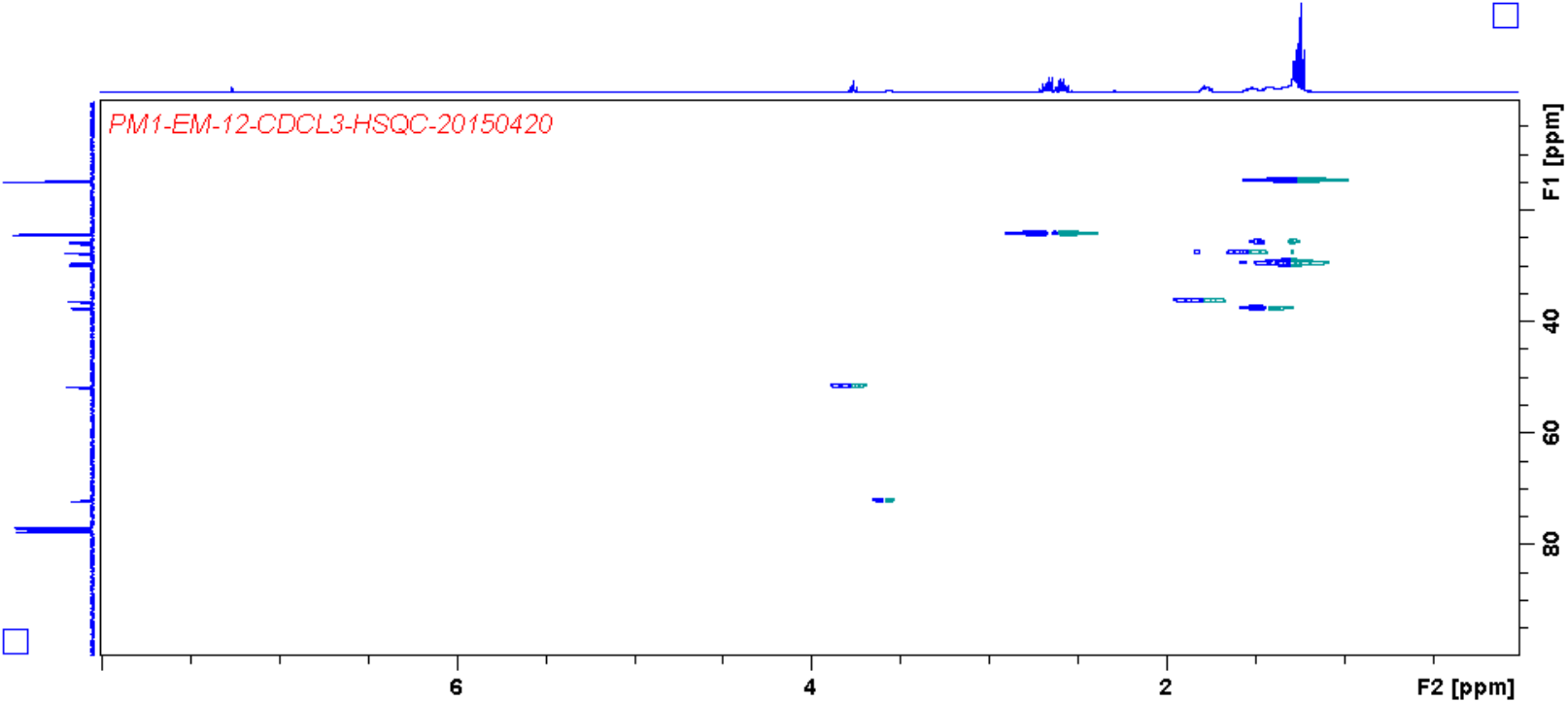
Characterization of compound I_a_, HSQC spectrum of compound **I_a_**.

**Fig. S36-A.**
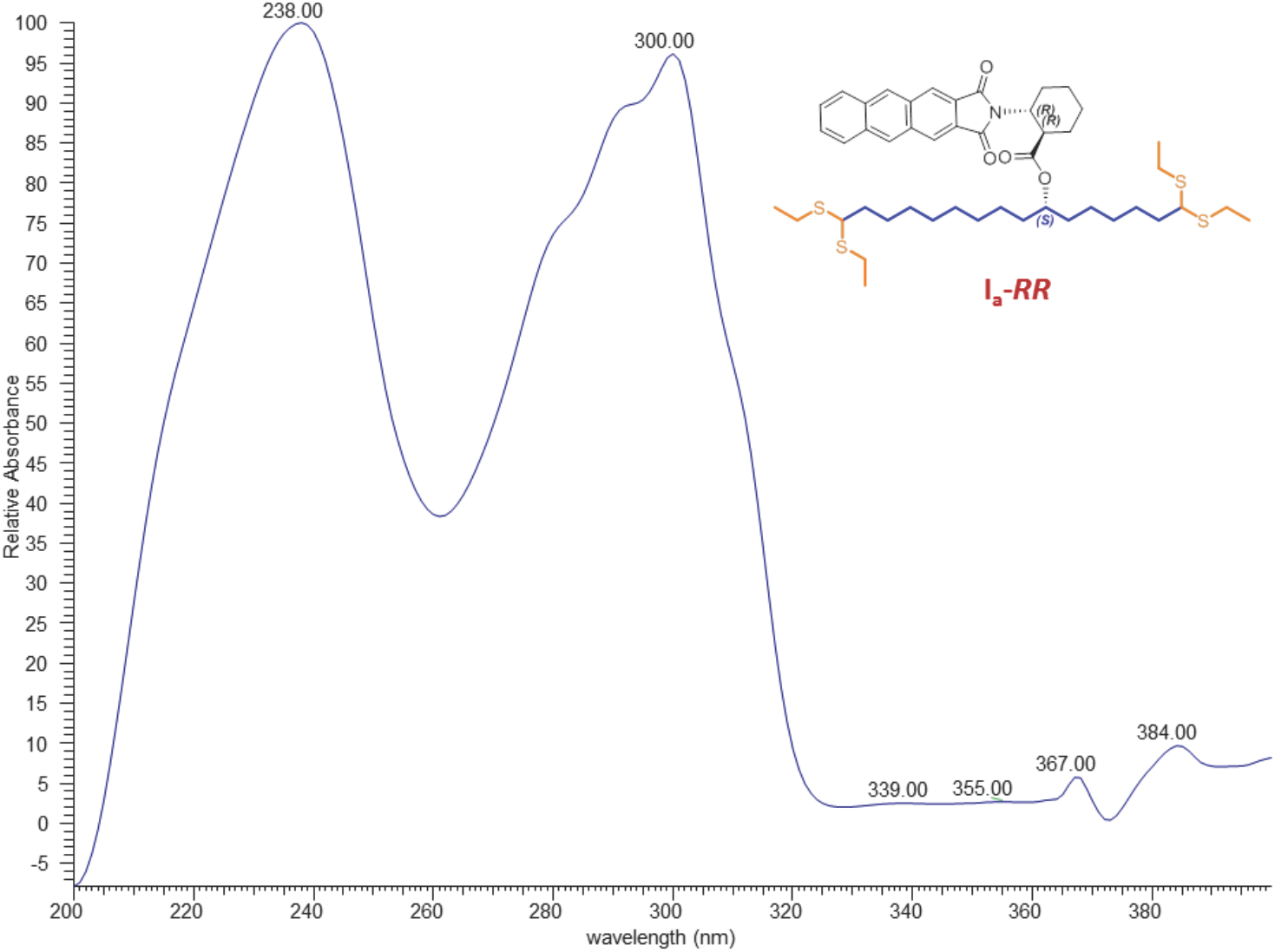
Characterization of compound I_a_-*RR*, UV spectrum of compound **I_a_-*RR***.

**Fig. S36-B.**
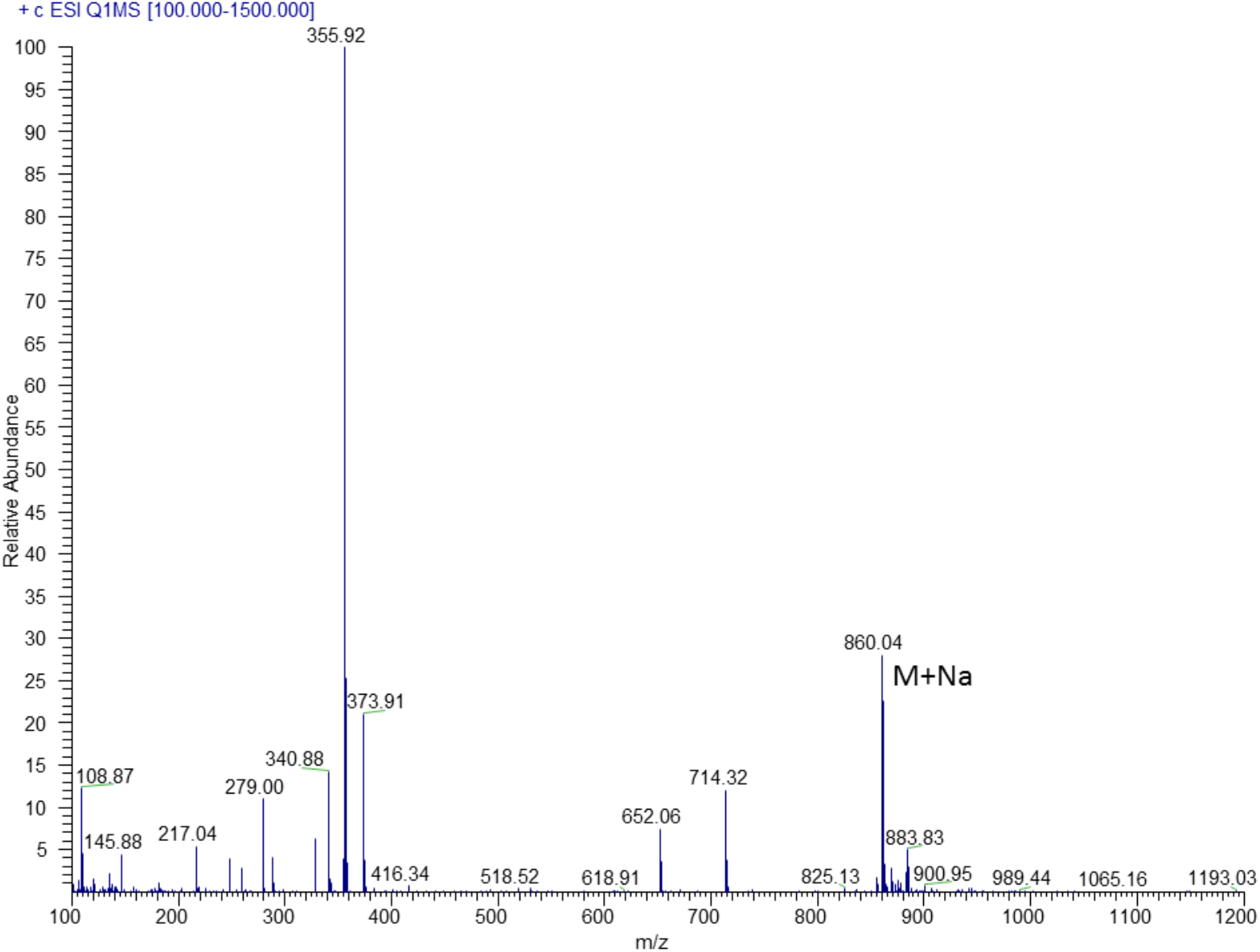
Characterization of compound I_a_-*RR*, MS/MS spectrum of compound **I_a_- *RR***.

**Fig. S36-C.**
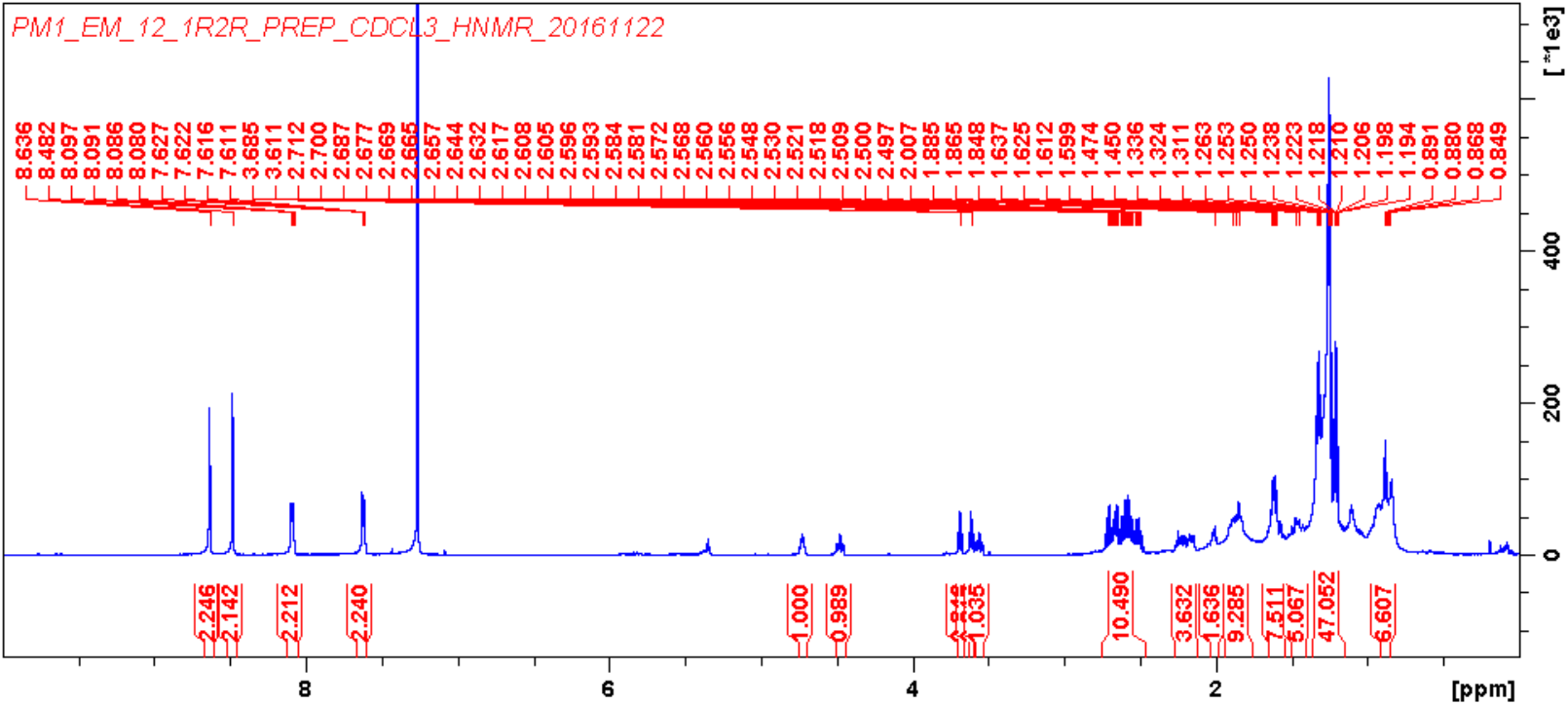
Characterization of compound I_a_-*RR*, ^1^H NMR (600 MHz, CDCl_3_) spectrum of compound **I_a_-*RR***.

**Fig. S36-D.**
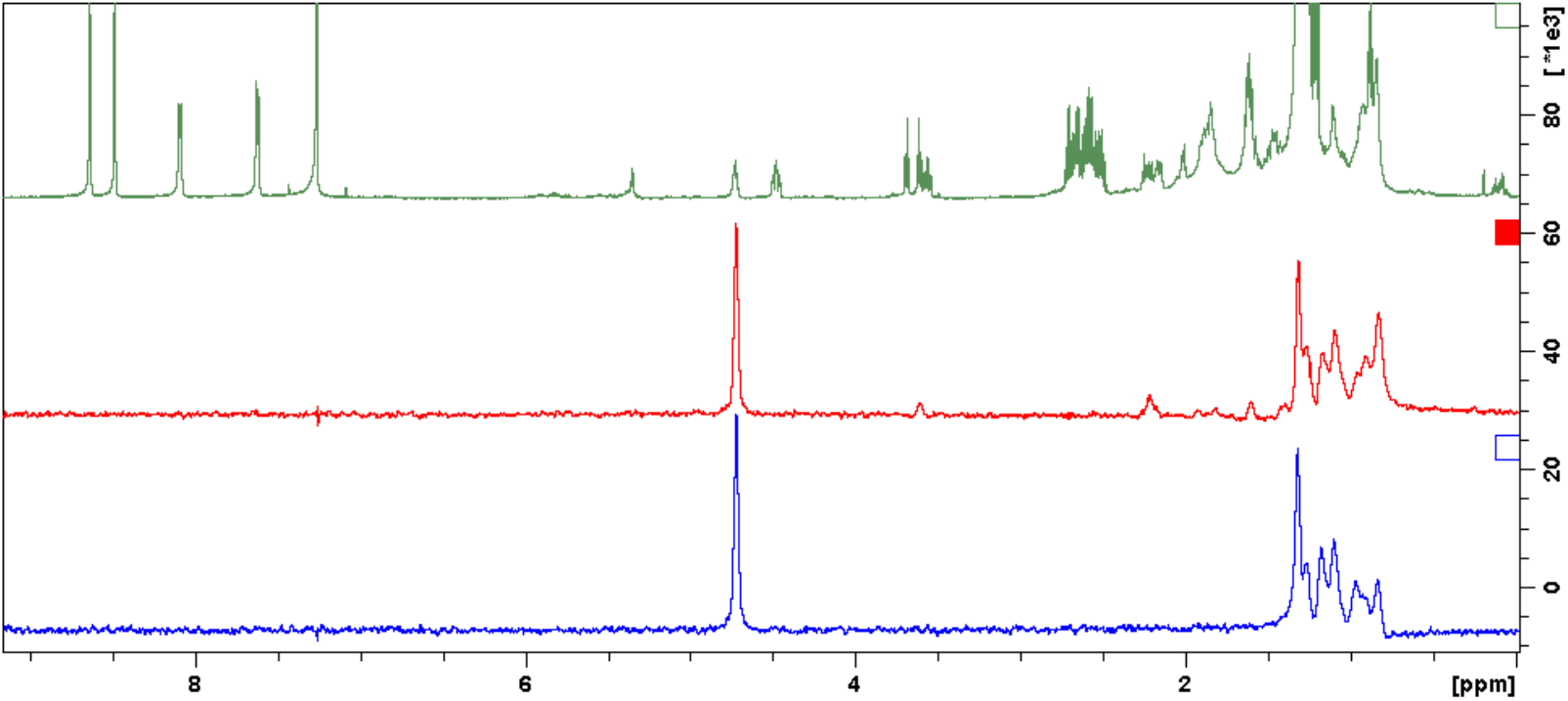
Characterization of compound I_a_-*RR*, Select 1D-TOCSY (600 MHz, CDCl_3_) spectrum of compound **I_a_-*RR***.

**Fig. S36-E.**
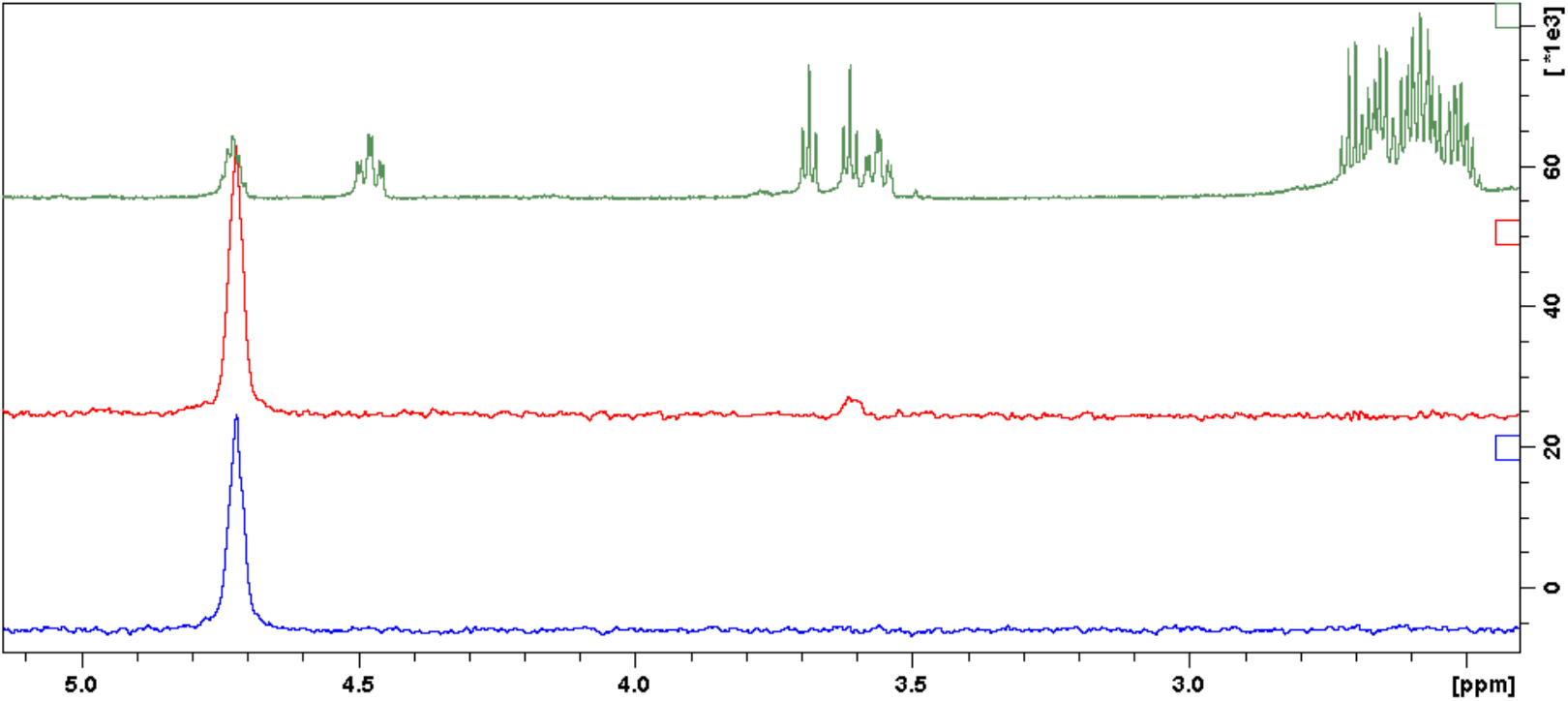
Characterization of compound I_a_-*RR*, Enlarged select 1D-TOCSY (600 MHz, CDCl_3_) spectrum of compound **I_a_-*RR***.

**Fig. S37-A.**
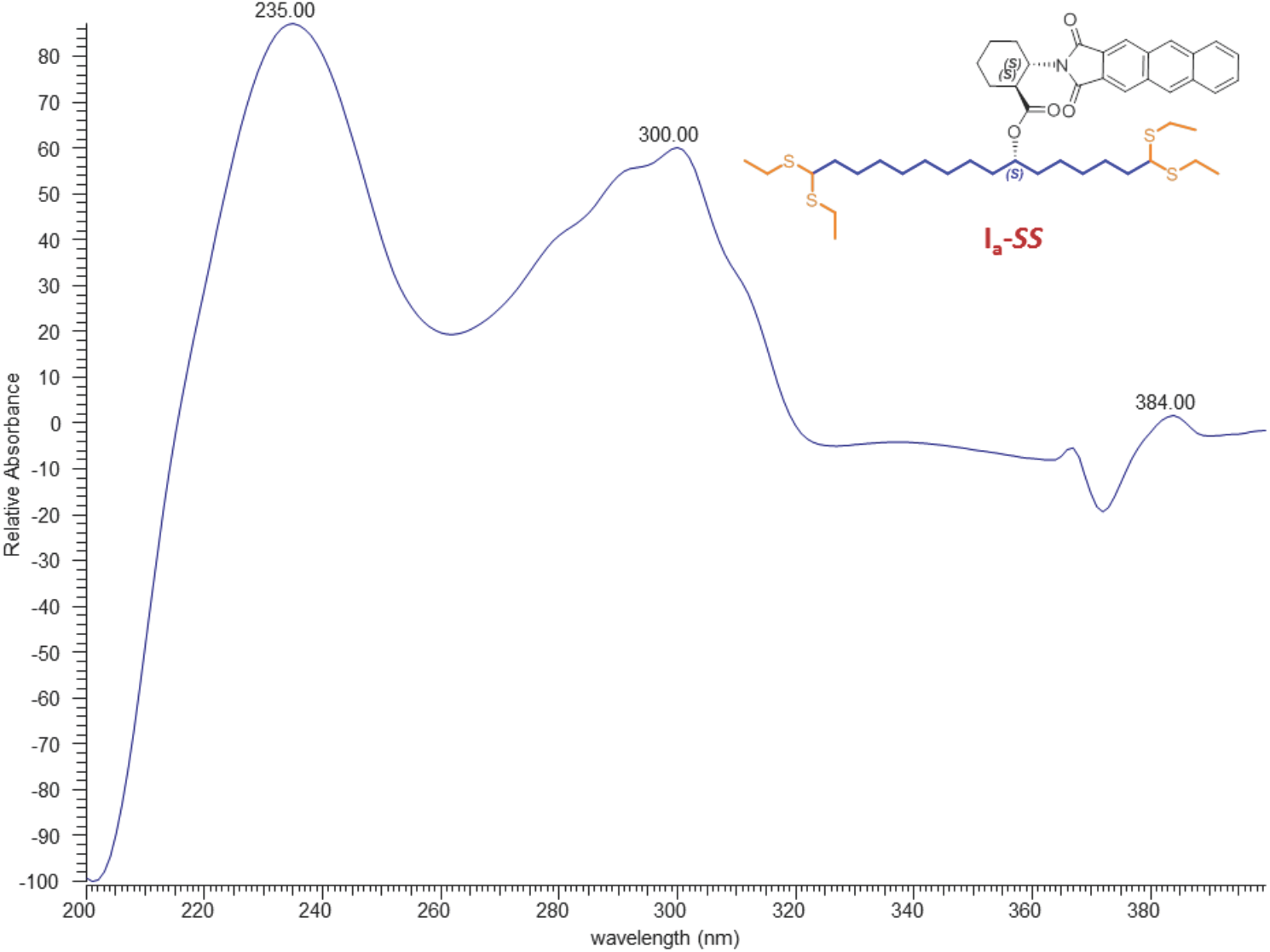
Characterization of compound I_a_-*SS*, UV spectrum of compound **I_a_-*SS***.

**Fig. S37-B.**
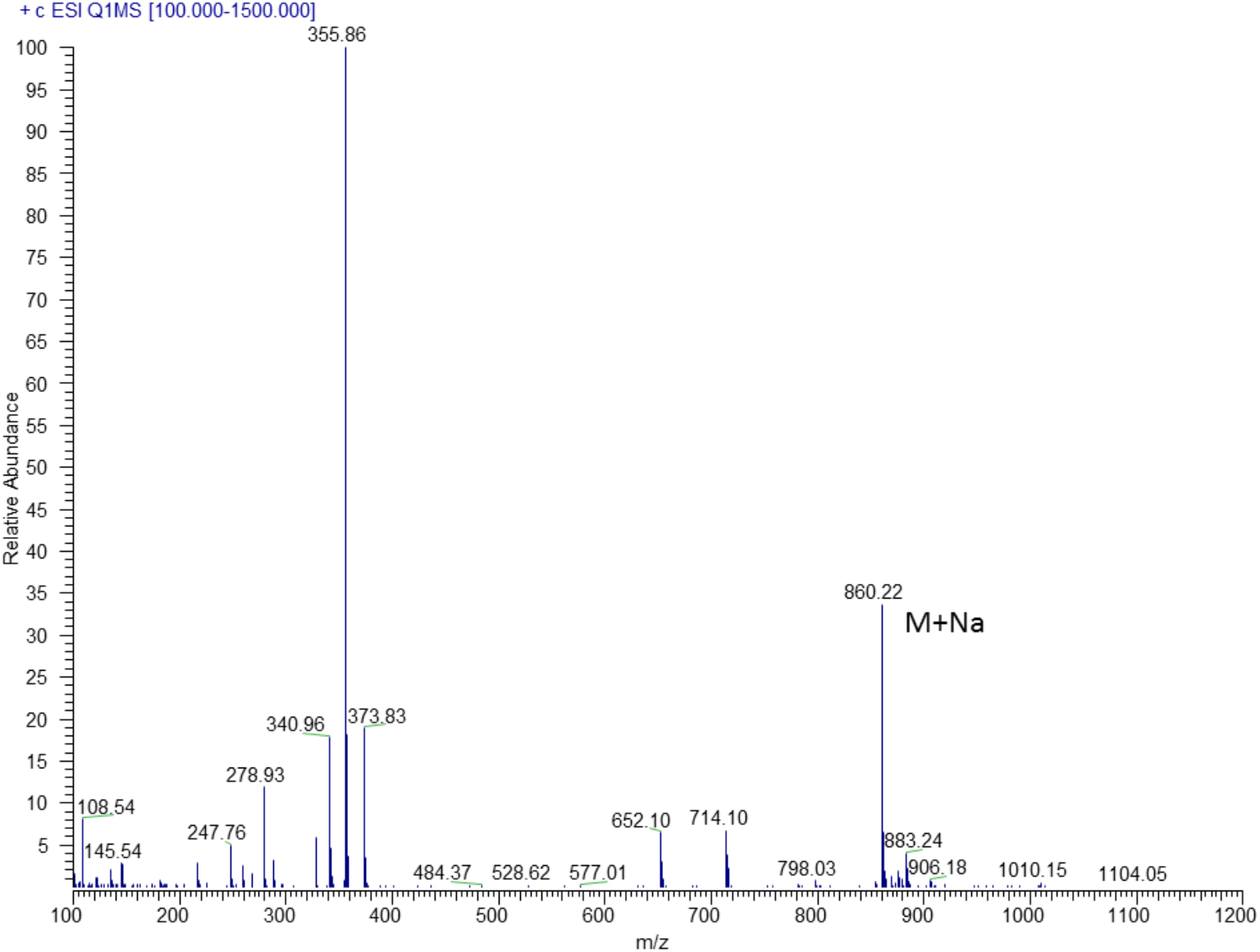
Characterization of compound I_a_-*SS*, MS/MS spectrum of compound **I_a_- *SS***.

**Fig. S37-C.**
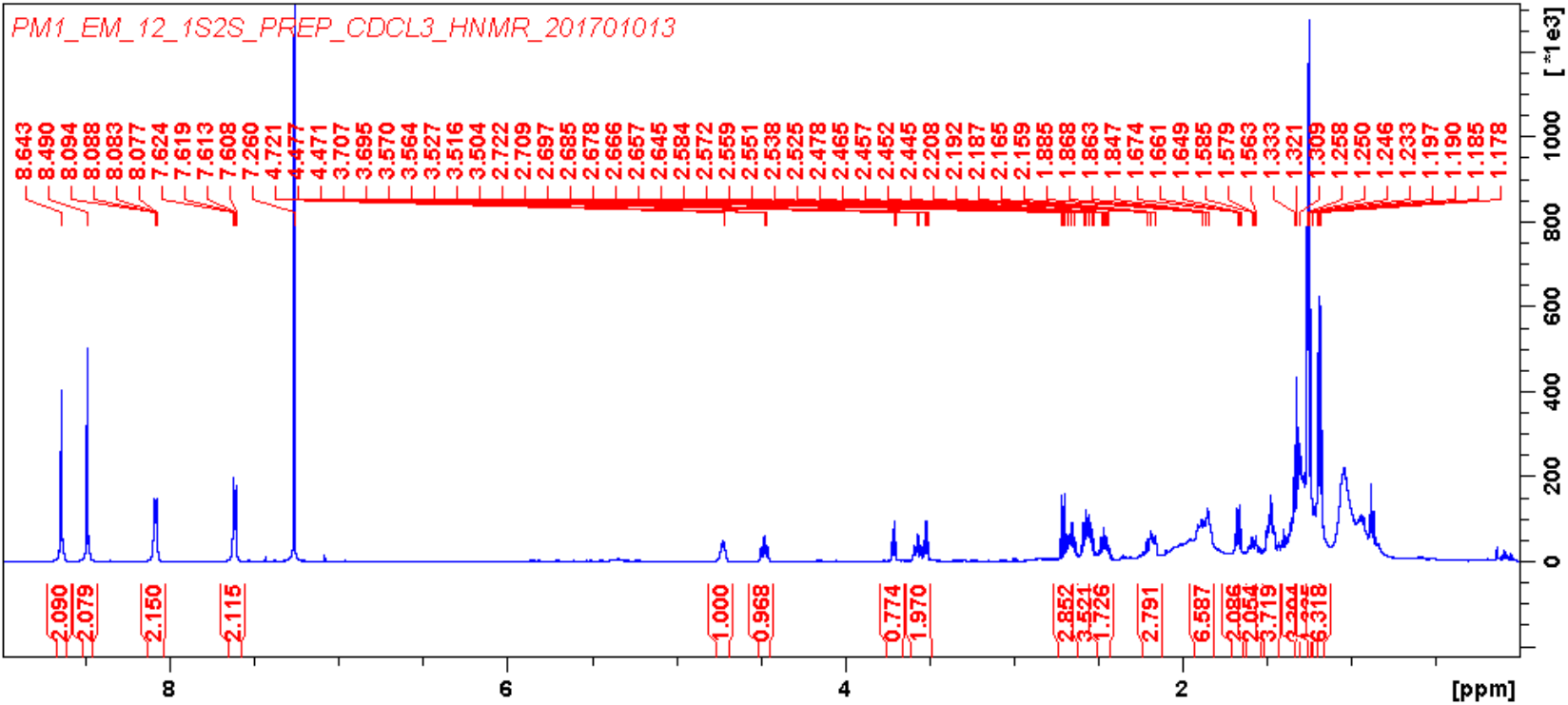
Characterization of compound I_a_-*SS*, ^1^H NMR (600 MHz, CDCl_3_) spectrum of compound **I_a_-*SS***.

**Fig. S37-D.**
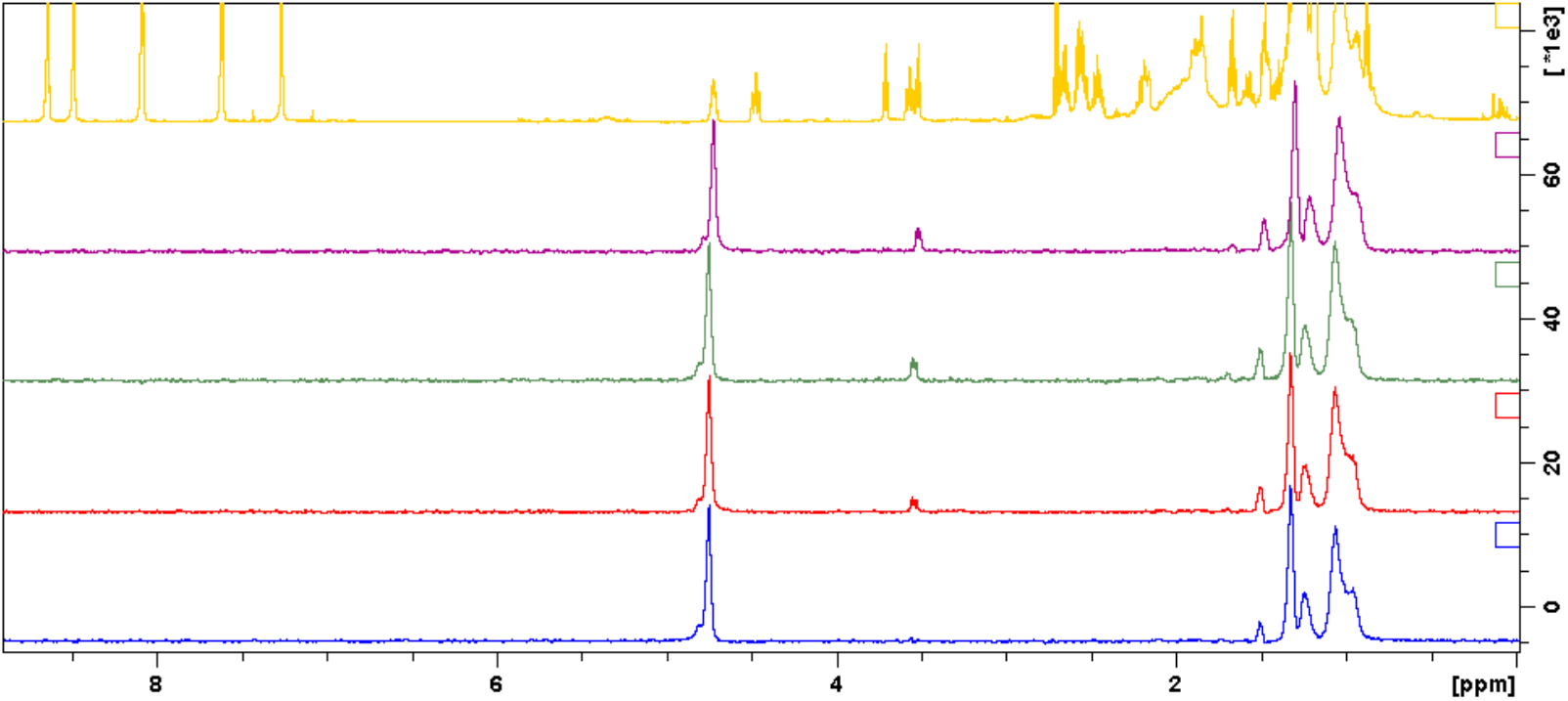
Characterization of compound I_a_-*SS*, Select 1D-TOCSY (600 MHz, CDCl_3_) spectrum of compound **I_a_-*SS***.

**Fig. S37-E.**
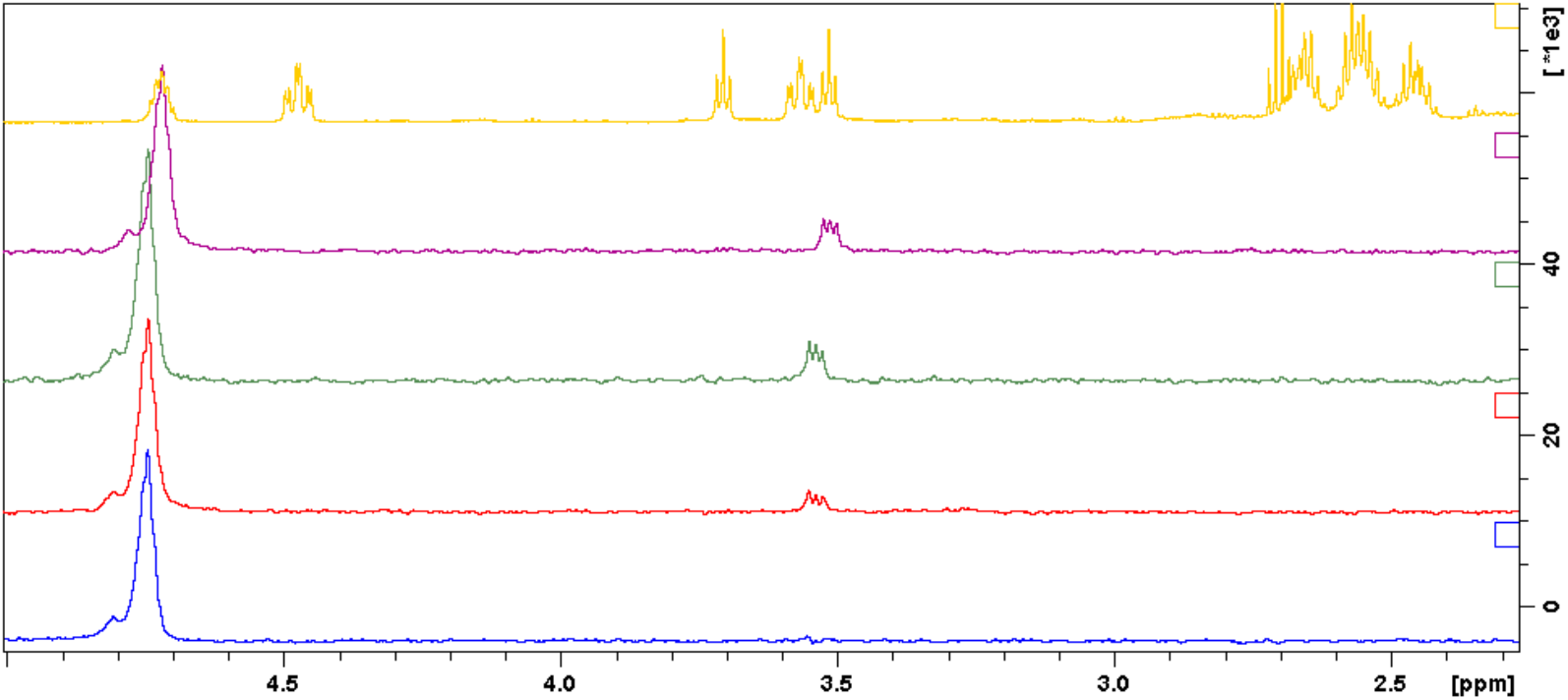
Characterization of compound I_a_-*SS*, Enlarged select 1D-TOCSY (600 MHz, CDCl_3_) spectrum of compound **I_a_-*SS***.

**Fig. S38.**
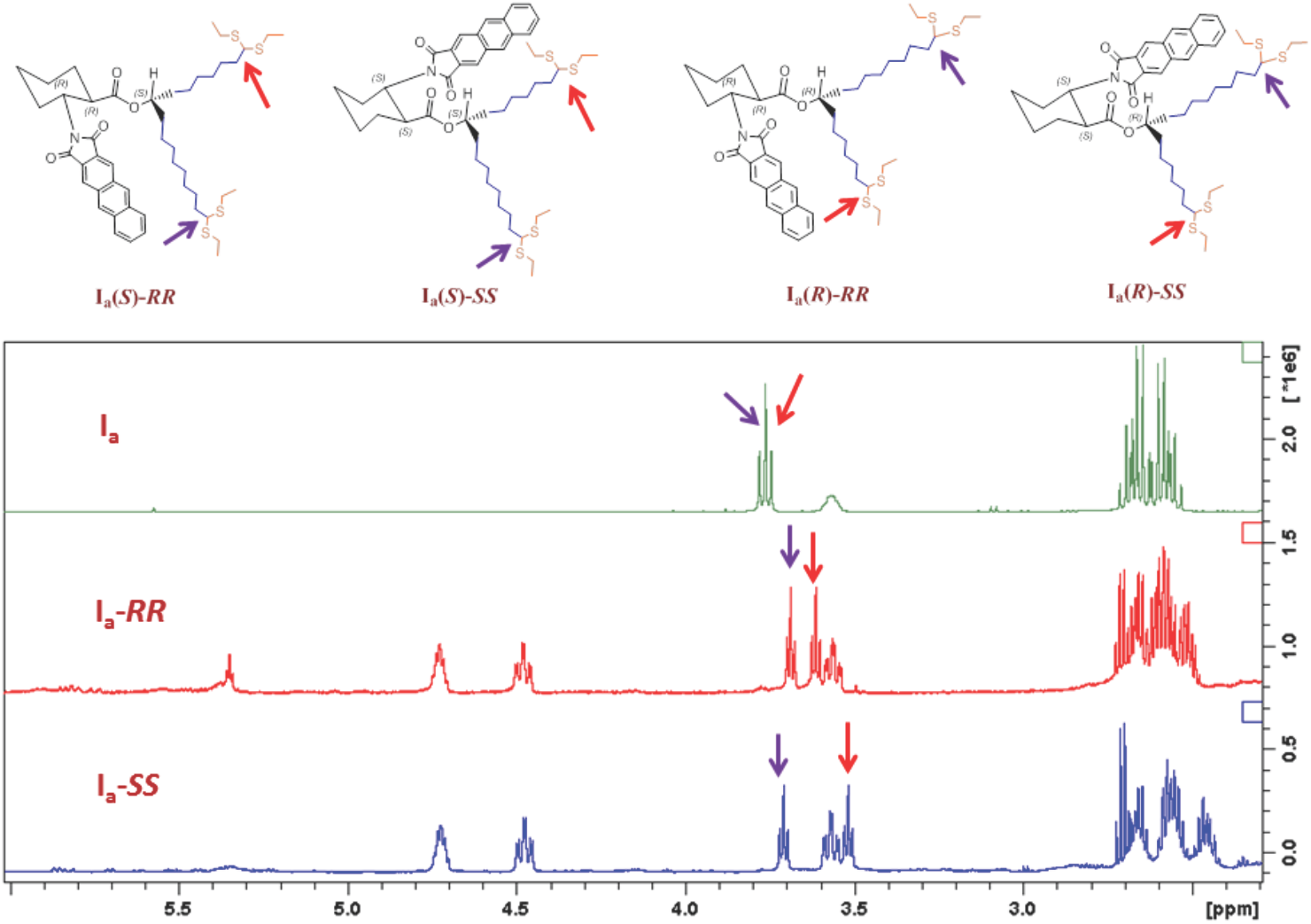
Absolute configuration identification of compound I_a_, Possible enantiomer derivatives of **I_a_** and enlarged select 1D-TOCSY (600 MHz, CDCl_3_) of compound **I_a_-*RR***, **I_a_-*SS***.

**Fig. S39-A.**
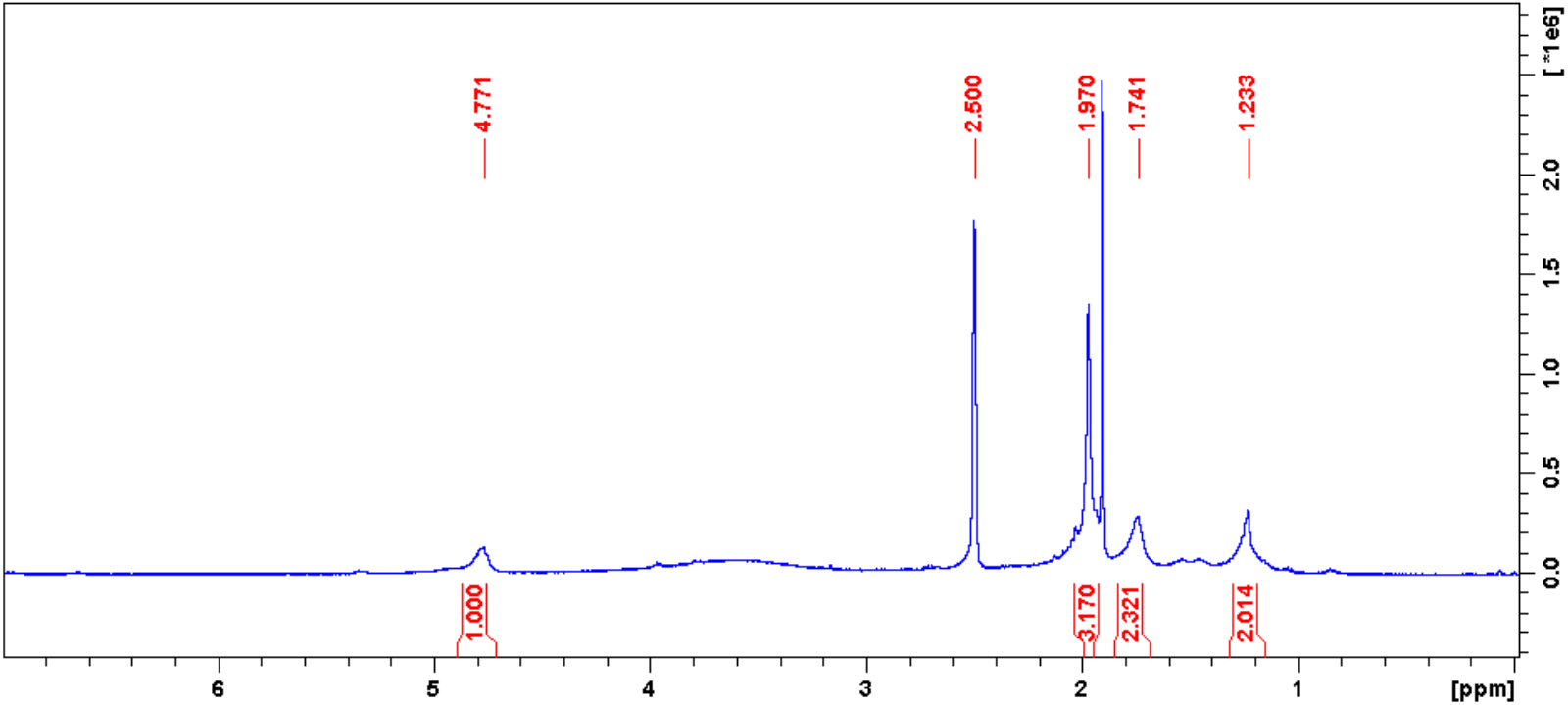
Characterization of R-Ac, ^1^H NMR (400 MHz, CDCl_3_) spectrum of **R-Ac**.

**Fig. S39-B.**
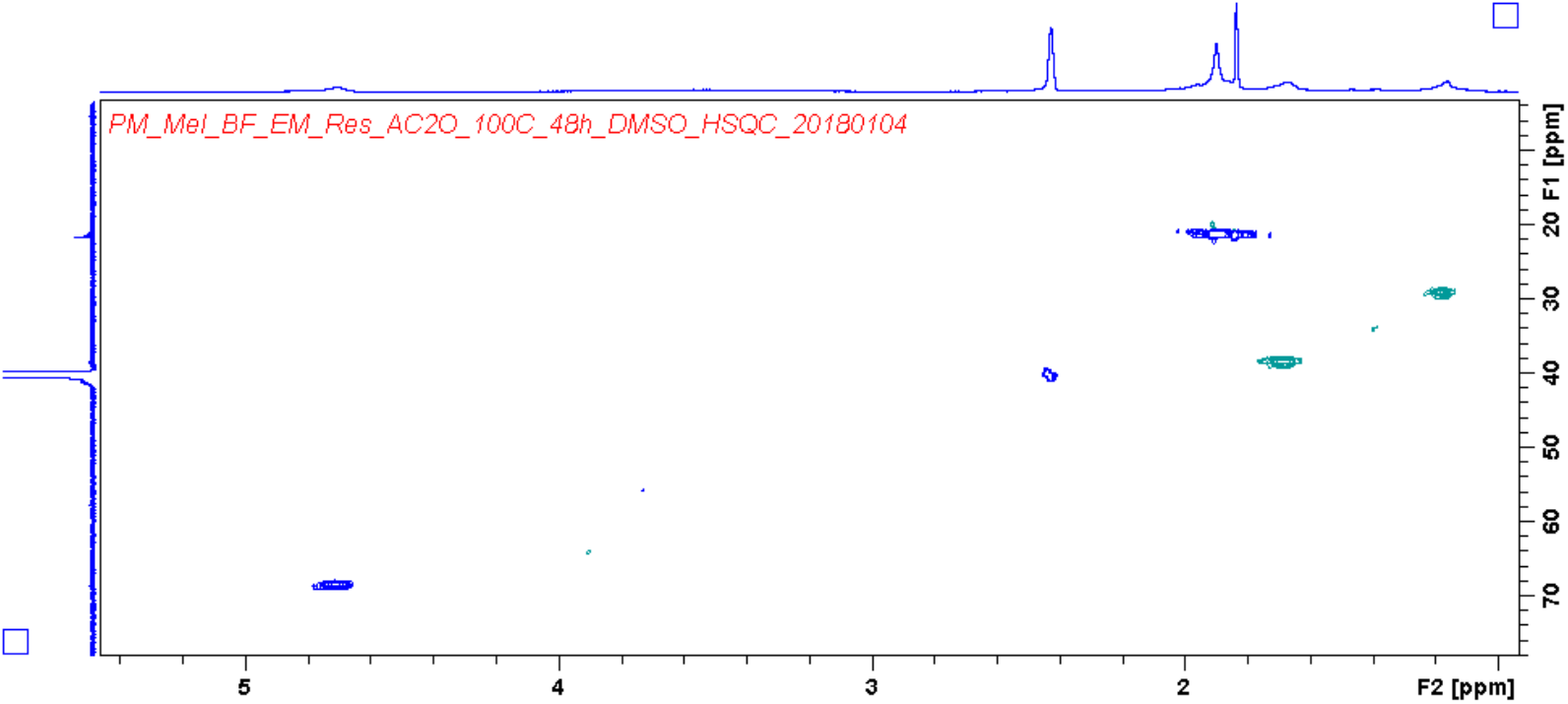
Characterization of R-Ac, HSQC spectrum of **R-Ac**.

**Fig. S39-C.**
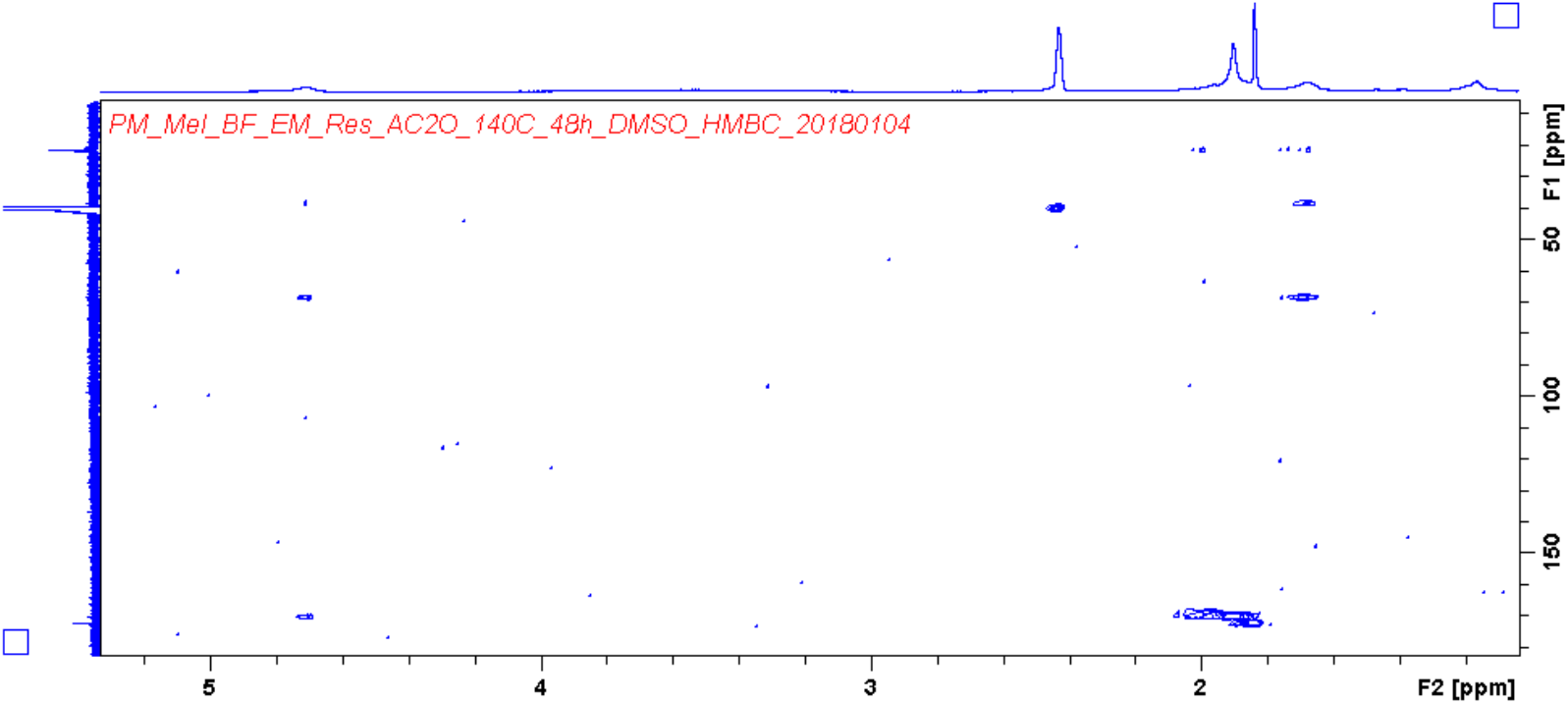
Characterization of R-Ac, HMBC spectrum of **R-Ac**.

**Fig. S39-D.**
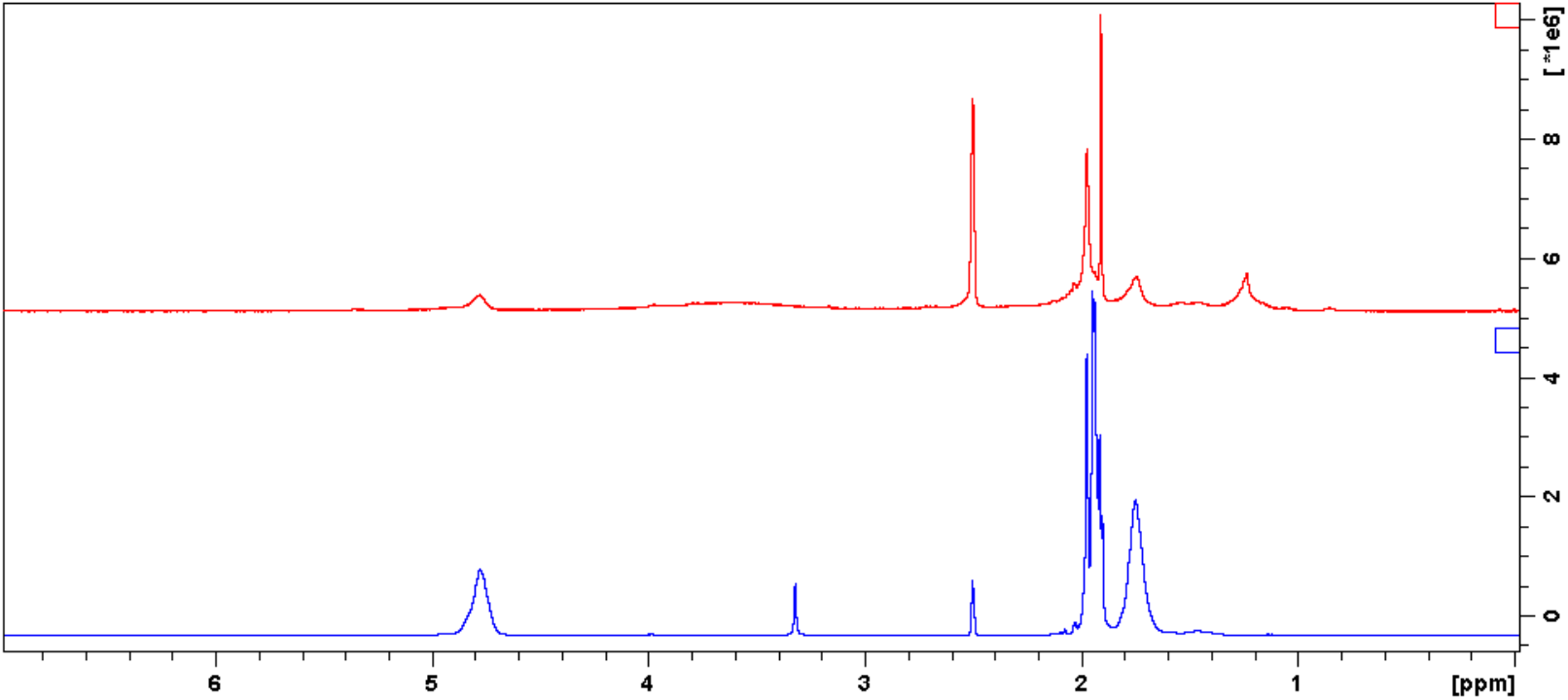
Characterization of R-Ac, Comparison ^1^H NMR (400 MHz, CDCl_3_) of **R-Ac** (red) and polyvinyl acetate (blue).

**Fig. S40-A.**
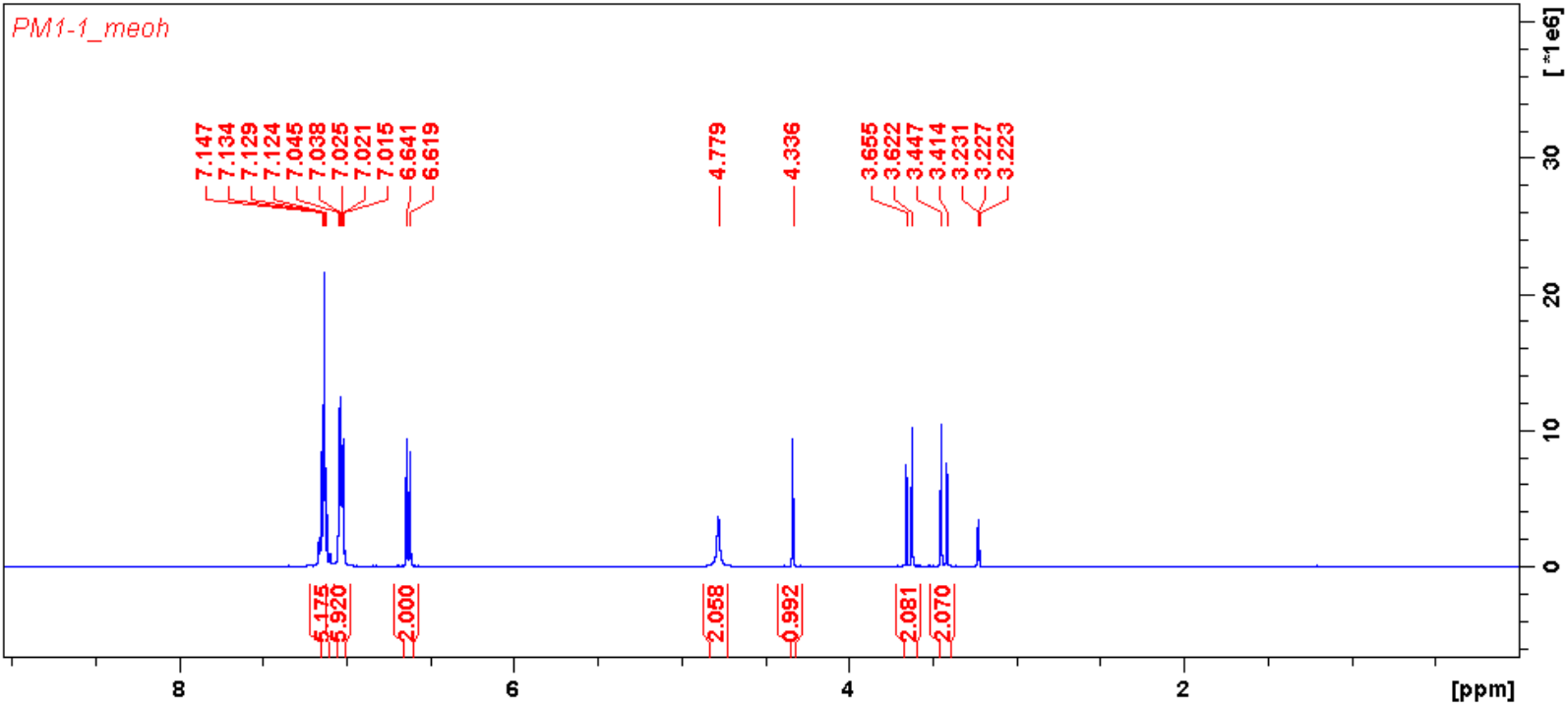
Characterization of compound XVI, ^1^H NMR (400 MHz, MeOD) spectrum of compound XVI.

**Fig. S40-B.**
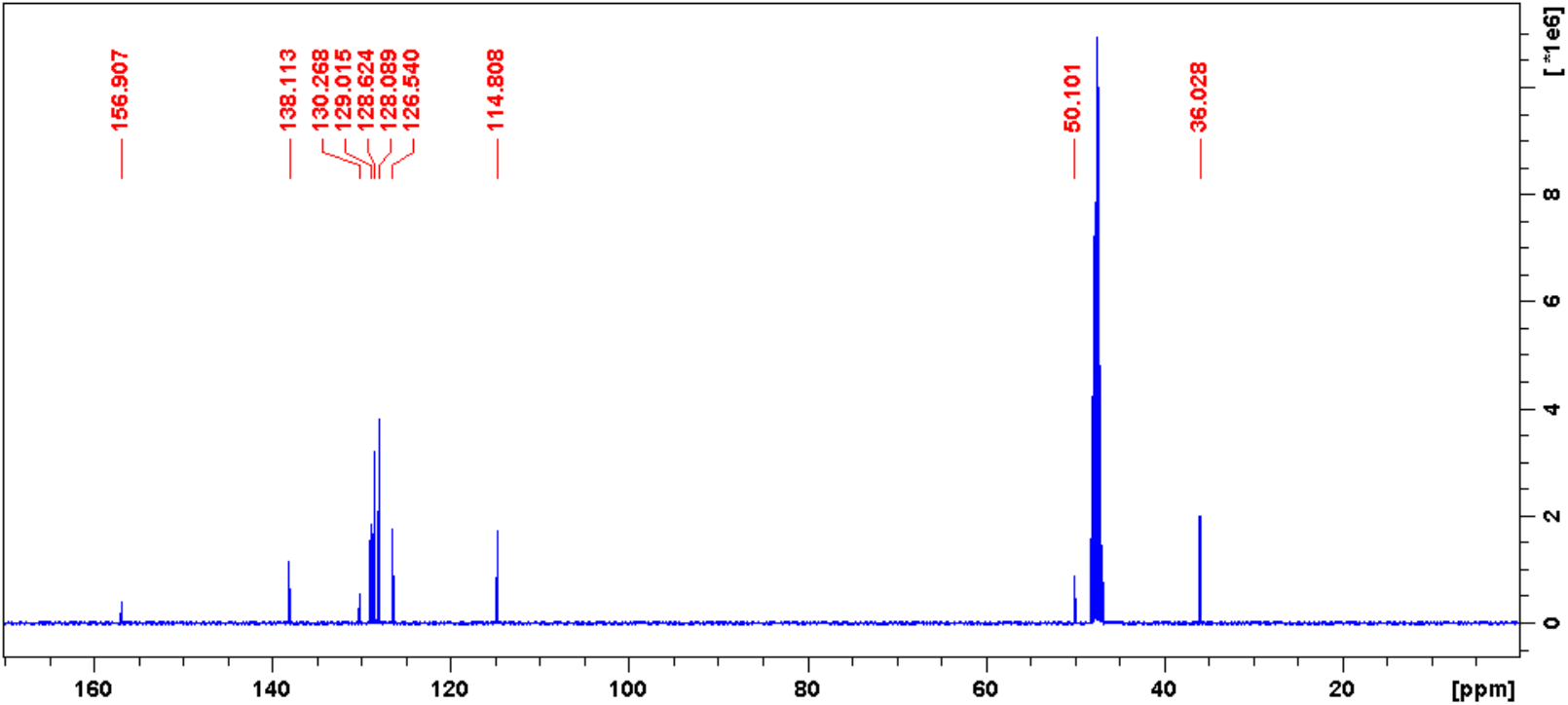
Characterization of compound XVI, ^13^C NMR (100 MHz, MeOD) spectrum of compound XVI.

**Fig. S40-C.**
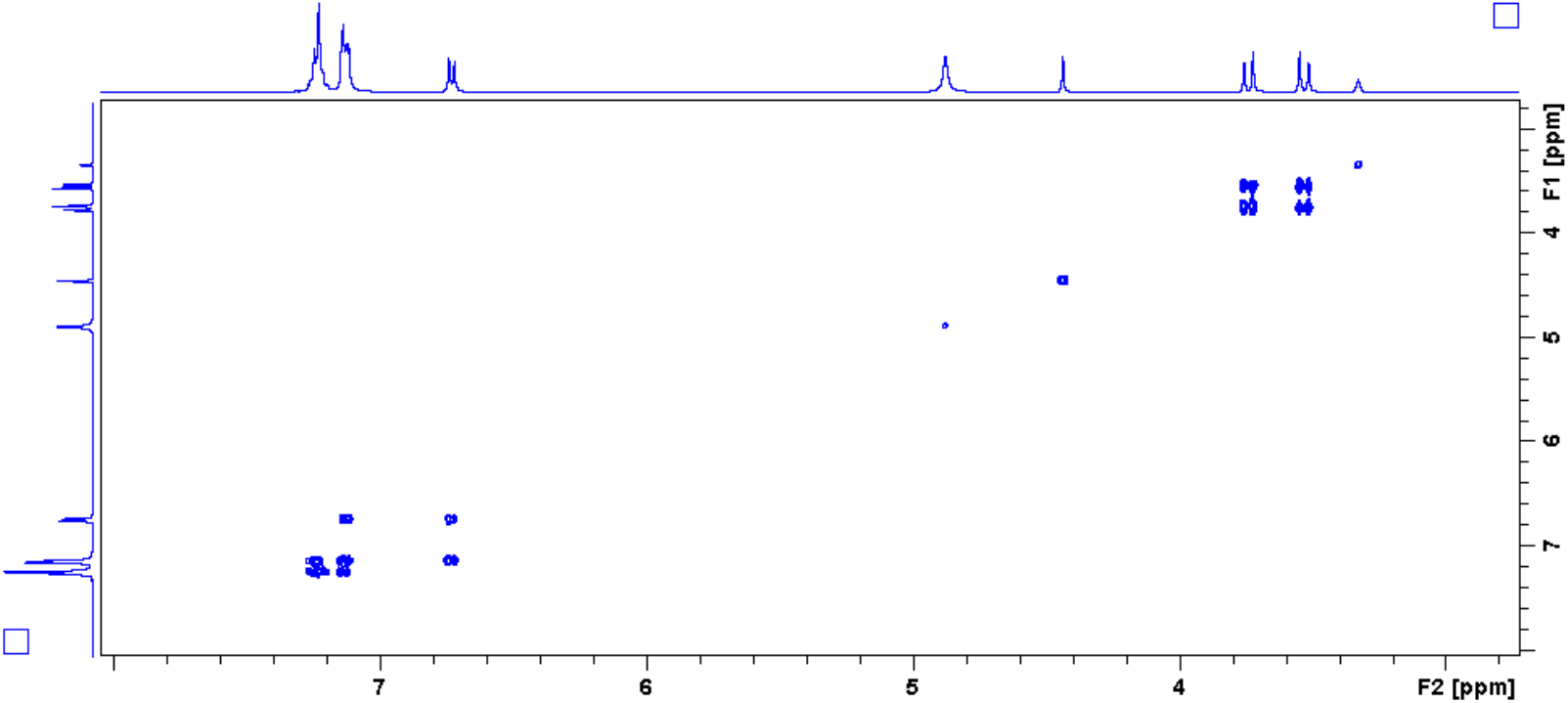
Characterization of compound XVI, HH-COSY spectrum of compound XVI.

**Fig. S40-D.**
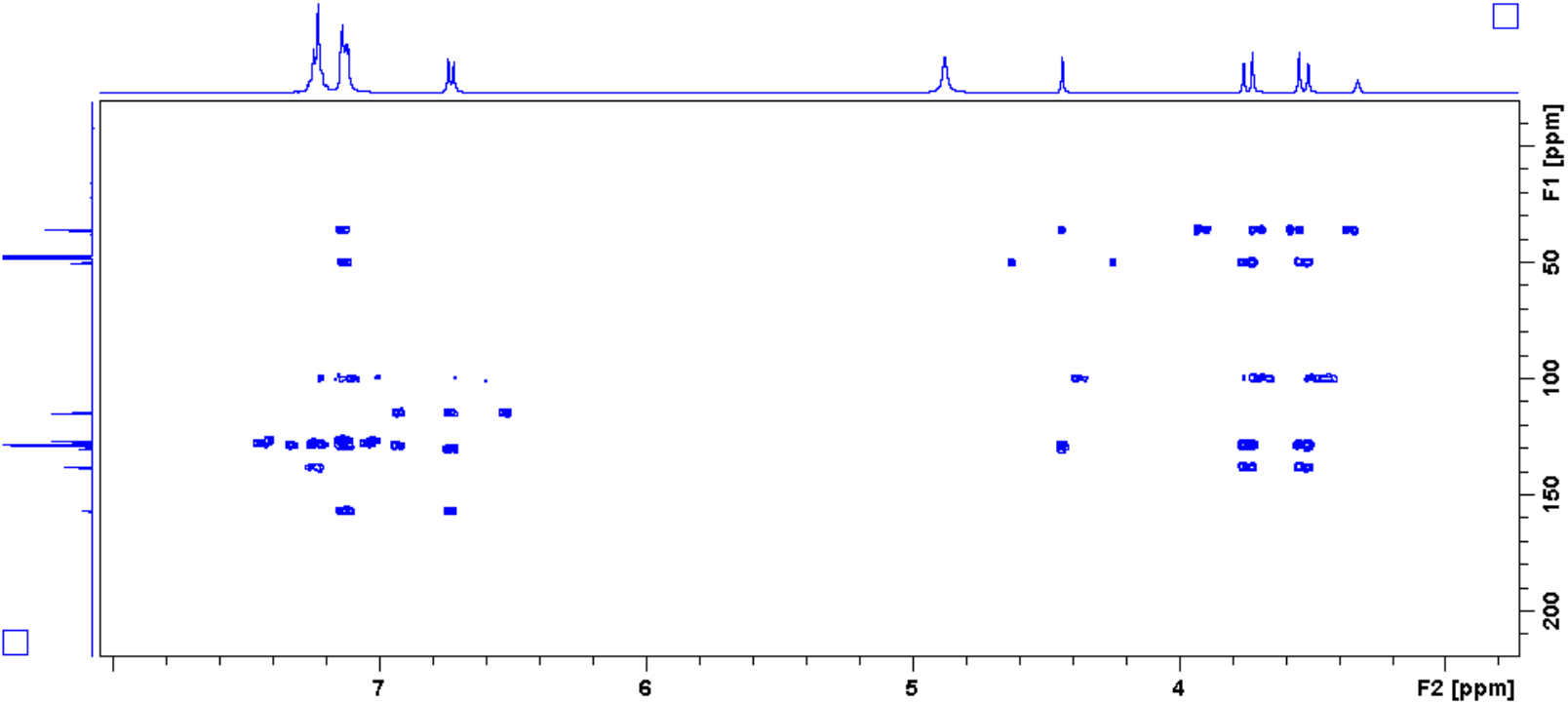
Characterization of compound XVI, HMBC spectrum of compound XVI.

**Fig. S41-A.**
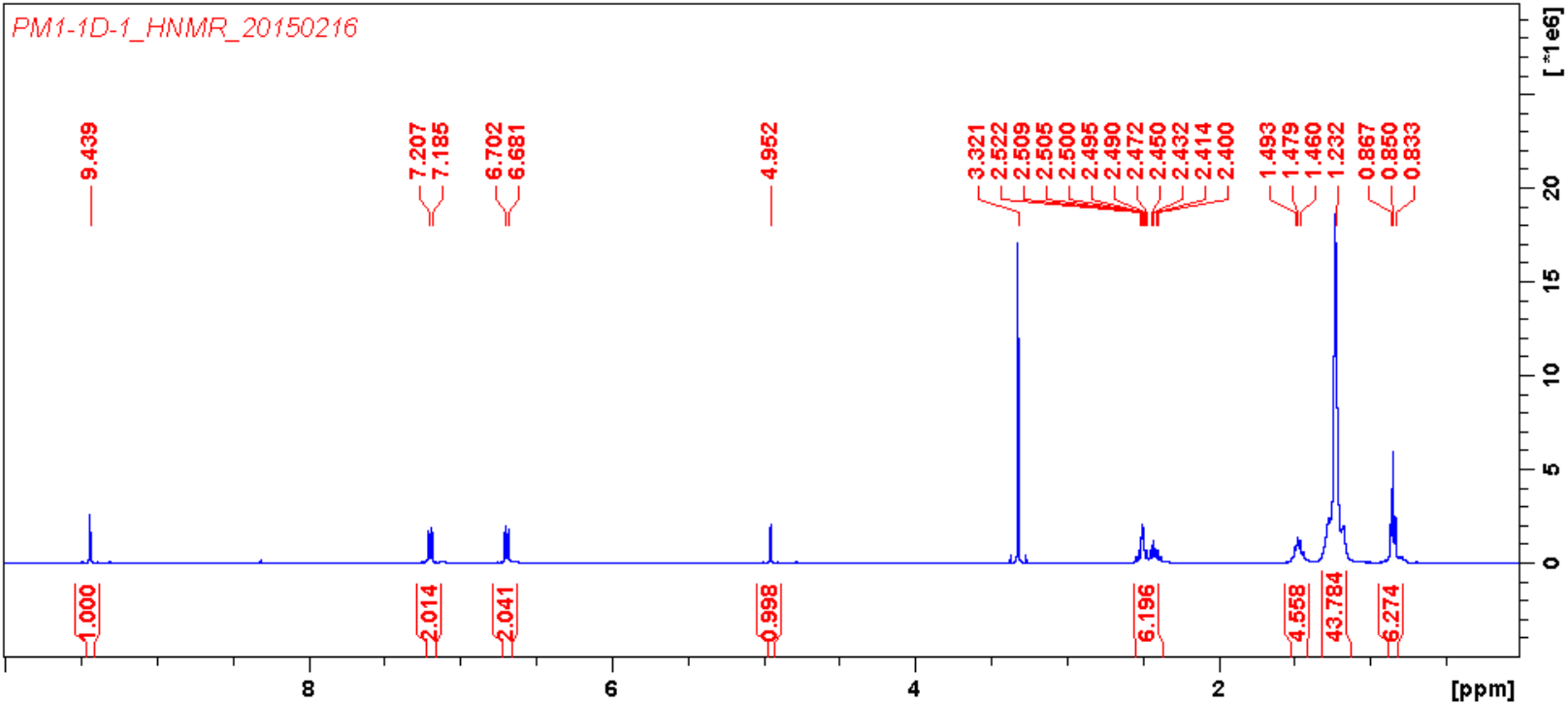
Characterization of compound XVII, ^1^H NMR (400 MHz, DMSO-*d*_*6*_) spectrum of compound XVII.

**Fig. S41-B.**
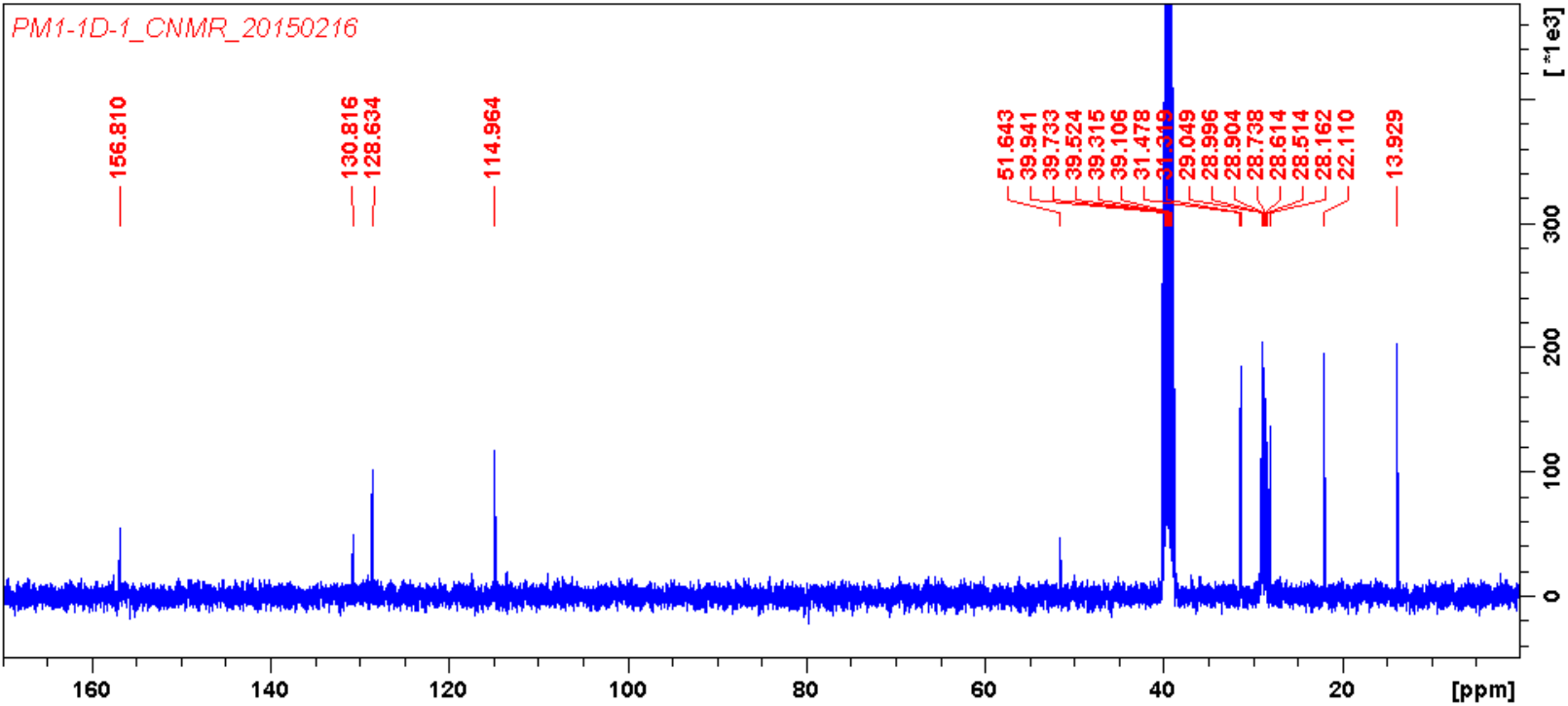
Characterization of compound XVII, ^13^C NMR (100 MHz, DMSO-*d*_*6*_) spectrum of compound XVII.

**Fig. S41-C.**
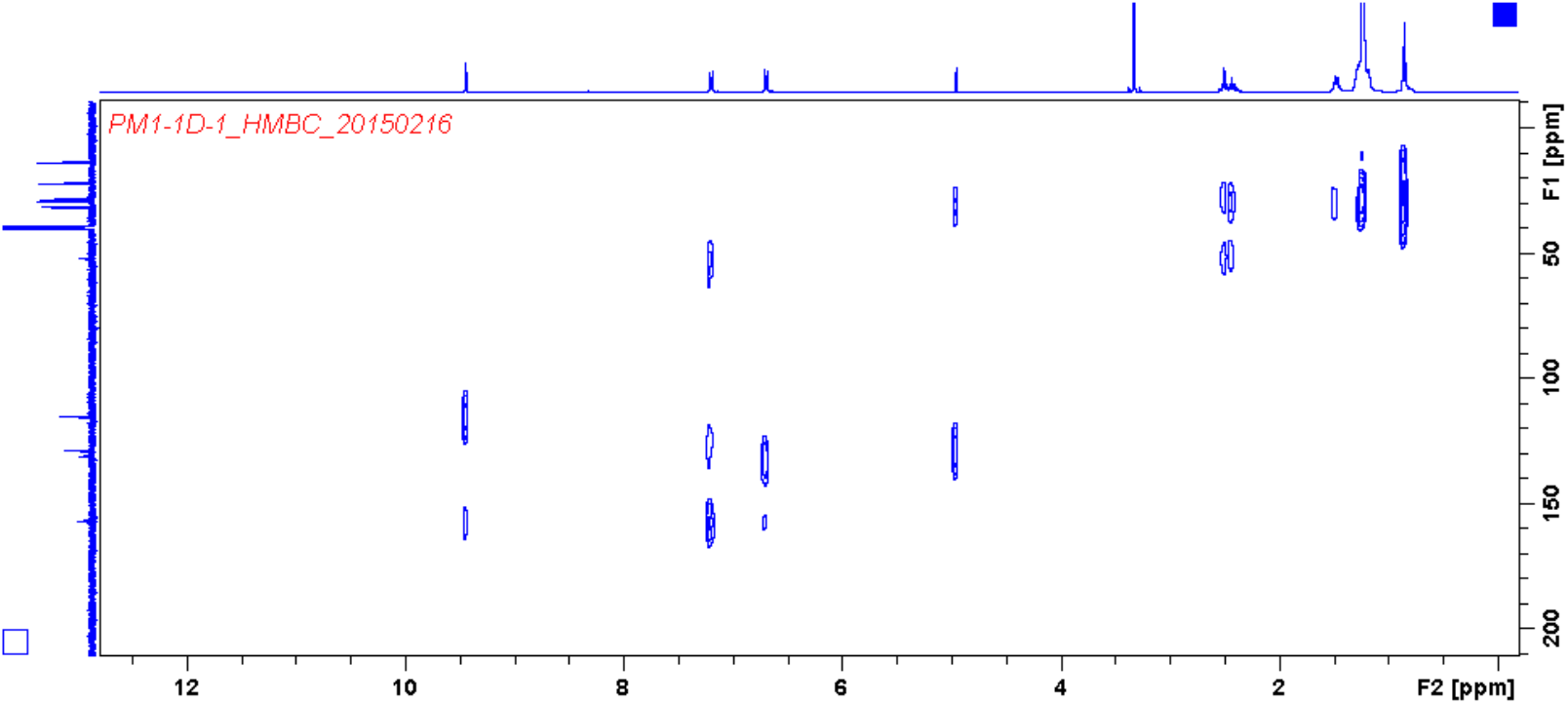
Characterization of compound XVII, HMBC spectrum of compound XVII.

**Fig. S42.**
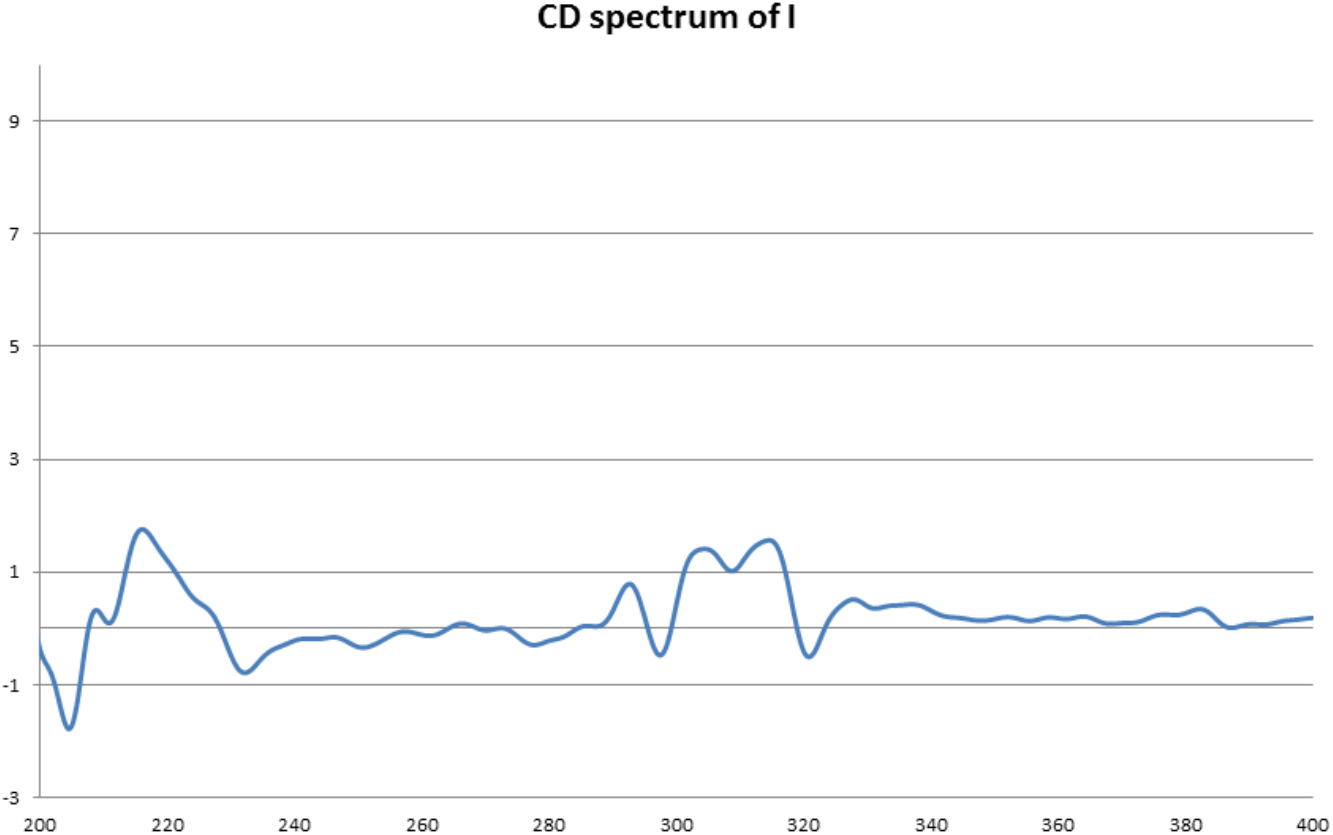
CD spectrum of I.

**Fig. S43.**
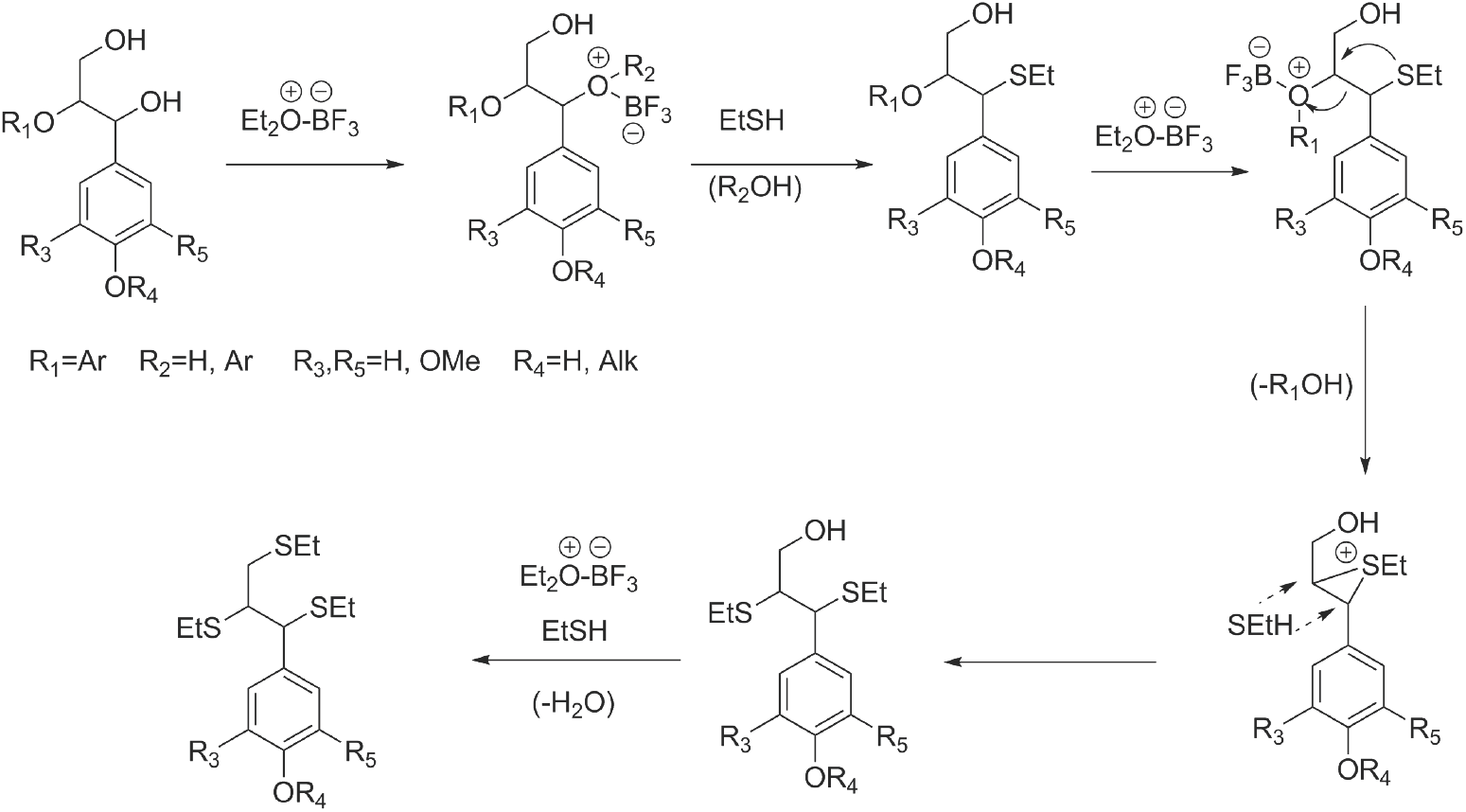
The proposed mechanism ^10^ of lignin thioacidolysis.

**Fig. S44.**
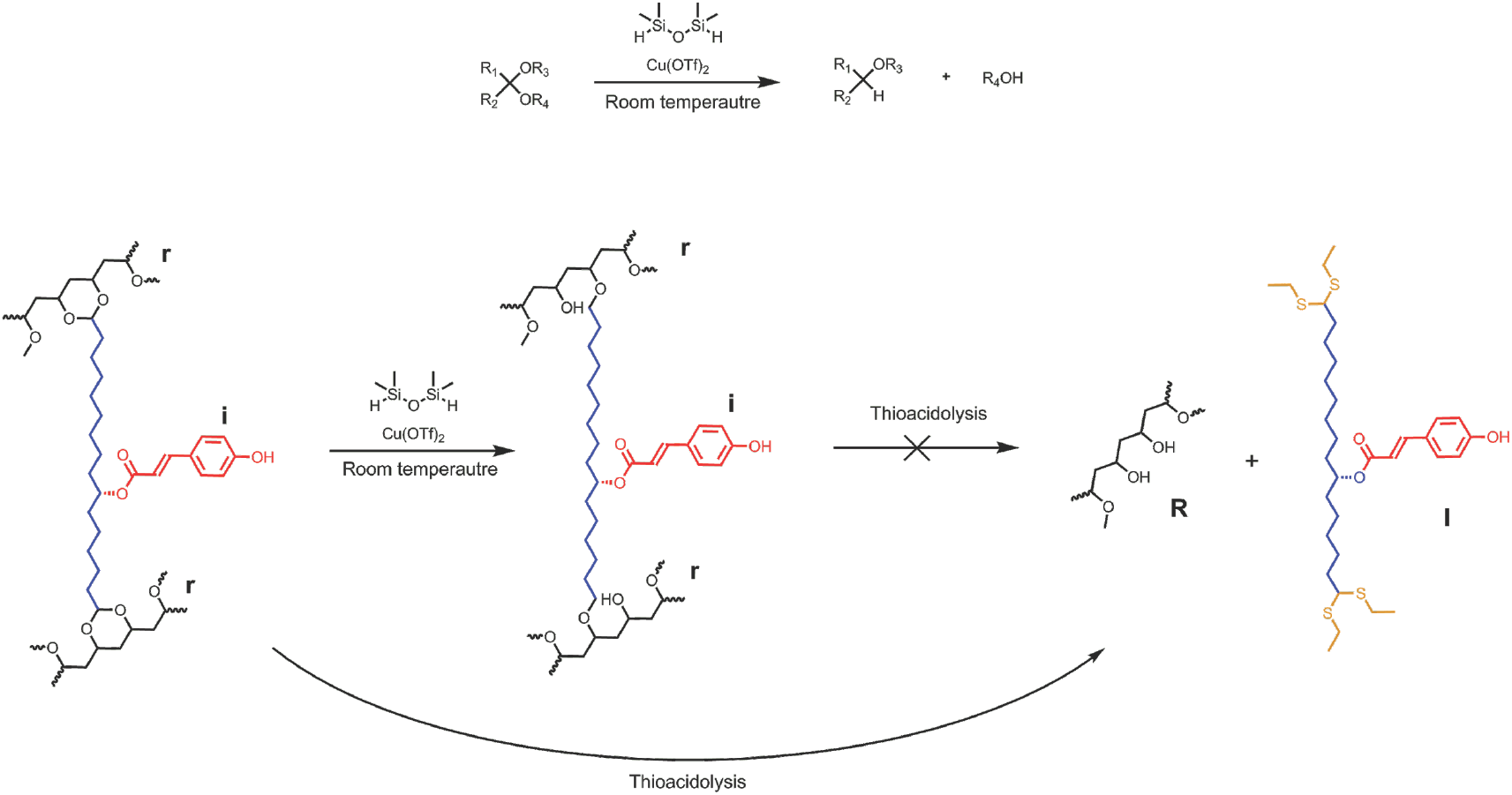
Reduction of acetals to ethers with TMDS catalyzed by Cu(OTf)_2_^25^. Sporopollenin pretreated with TMDS/Cu(OTf)_2_ becomes completely inert to thioacidolysis degradation.

**Fig. S45.**
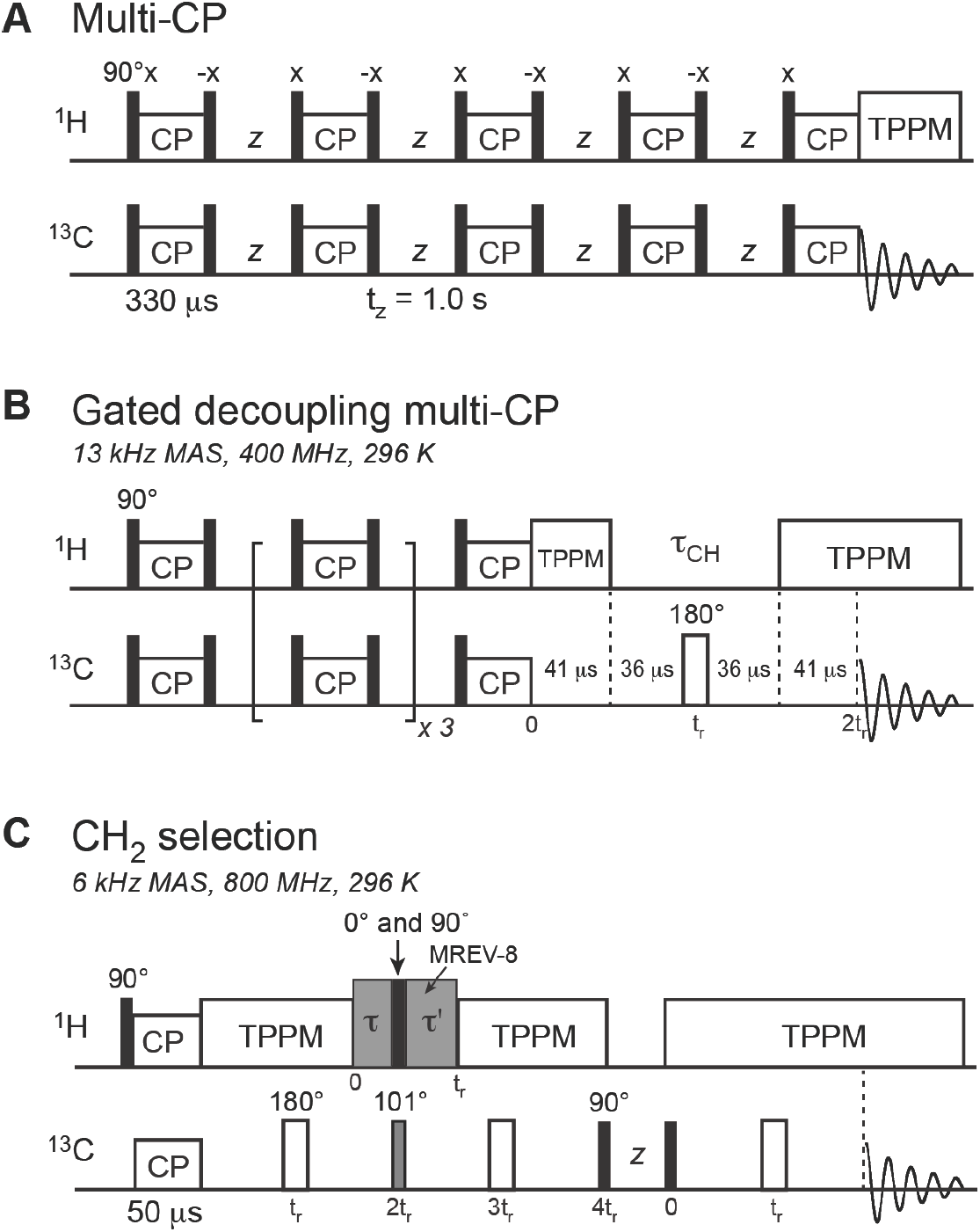
^13^C SSNMR pulse sequences. **A**, Multi-CP sequence. Five CP periods alternated with ^1^H spin–lattice relaxation periods (t_z_) to give quantitative ^13^C spectra. The CP contact time was 330 μs and the t_z_ period was 1.0 s. These values were optimized using the model tripeptide, formyl-MLF, by comparing its multi-CP spectrum with a quantitative direct polarization spectrum measured with a recycle delay of 35 s. **B**, Gated-decoupling multi-CP sequence, where ^1^H-^13^C dipolar coupling during the window t_CH_ suppressed the signals of protonated carbons and selected the signals of quaternary carbons. **C**, CH_2_ selection sequence. MREV-8 was used for ^1^H homonuclear decoupling, and t and t’ were 35 μs and 123 μs, respectively, which were optimized using L-Leucine as the model compound.

**Fig. S46.**
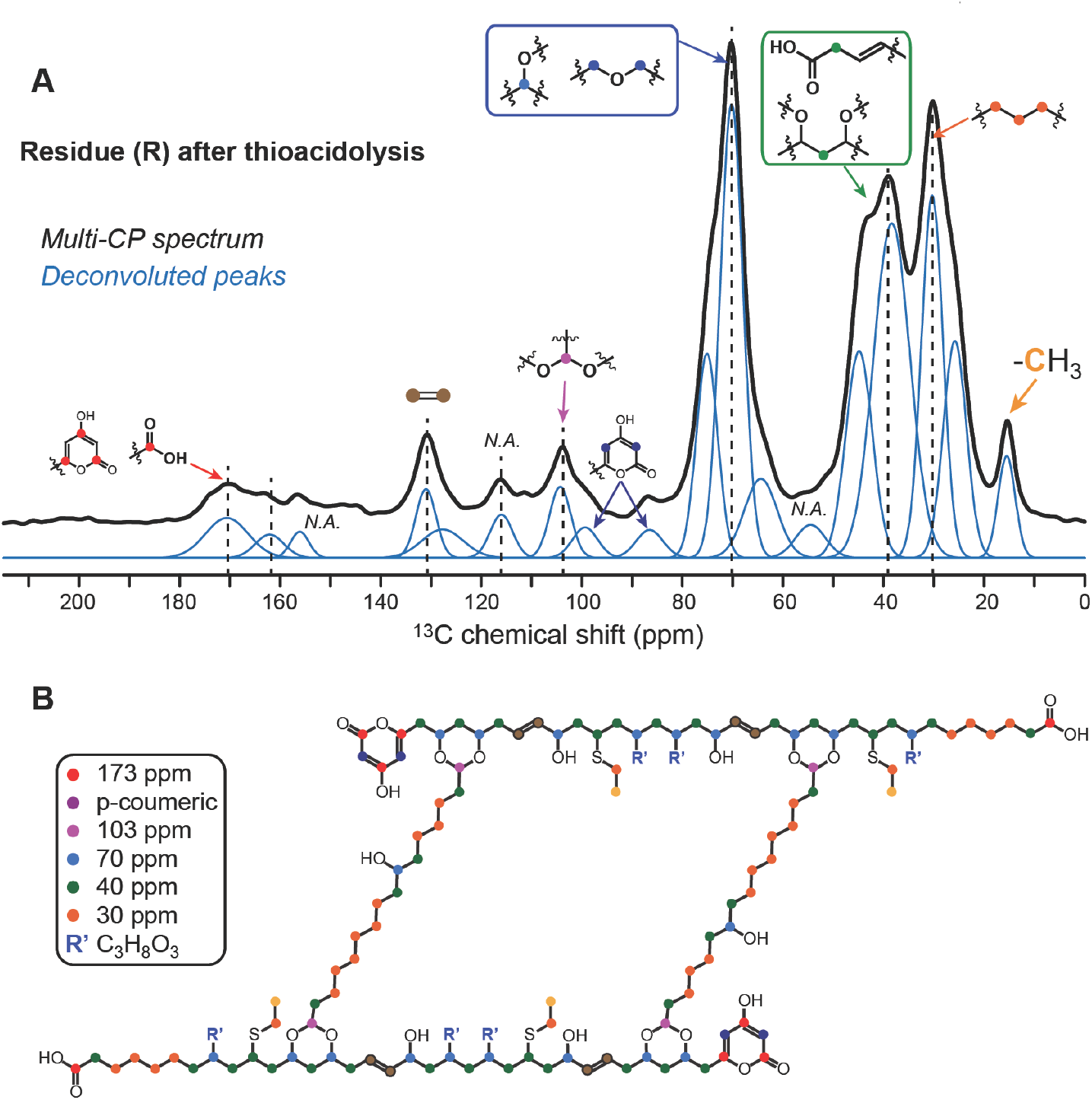
^13^C solid-state NMR spectrum and proposed structure of residue (**R**) after thioacidolysis. **A**, Multi-CP spectrum of **R**, measured on the 400 MHz spectrometer at 296 K under 13 kHz MAS. Spectral deconvolution gave the relative number of carbons for each peak. **B**, Proposed structure of **R**. The functional group R’ is proposed to be glycerol (C_3_H_8_O_3_) or glycerol like moiety, and some of these groups were removed and replaced with ethylsulfanyl groups during thioacidolysis.

**Fig. S47.**
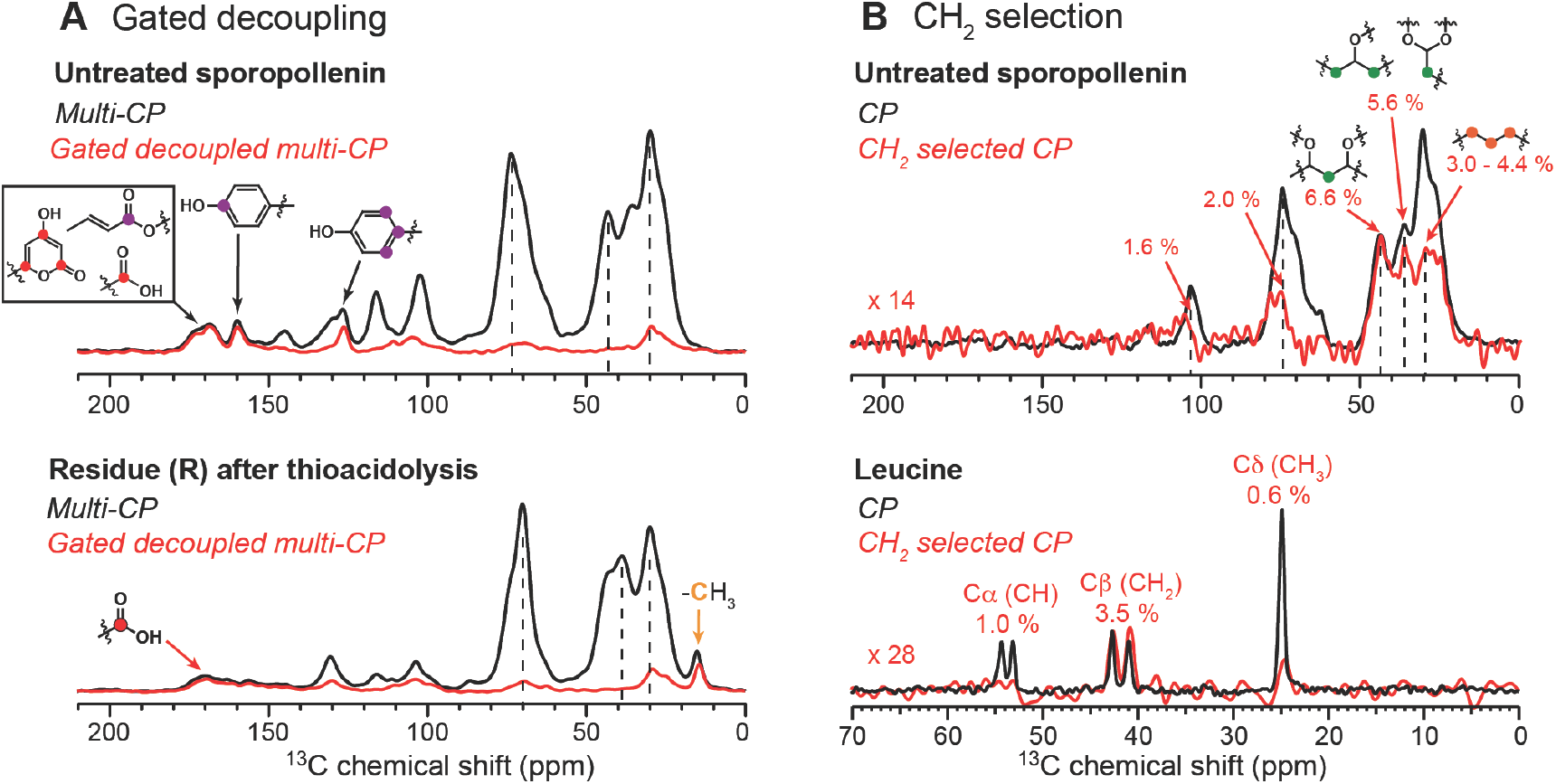
Spectrally edited ^13^C solid-state NMR spectra of sporopollenin and residue (**R**). **A**, Comparison between regular multi-CP and gated ^1^H-decoupled multi-CP spectra. The latter suppresses the signals of protonated carbons. **B,** Comparison of regular single-CP spectrum and CH_2_-selected single-step CP spectrum. The non-CH_2_ signals are suppressed to 1-2% of its original intensity, as shown by the spectra of unlabeled L-Leucine.

**Fig. S48.**
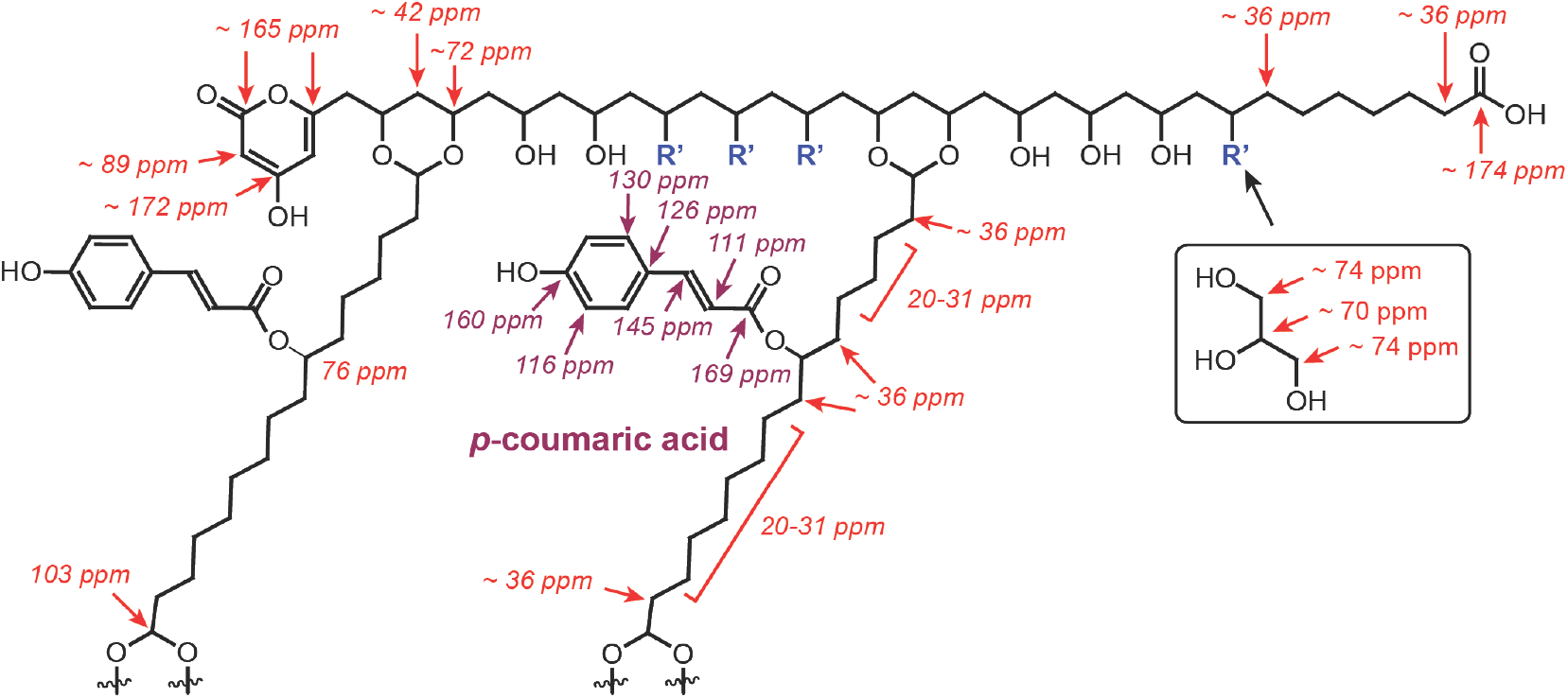
Approximate ^13^C chemical shifts obtained from ChemDraw and from measured spectra for the proposed structure of pine sporopollenin. For simplicity the second backbone is not shown.

## Supplementary Tables

**Table S1.**
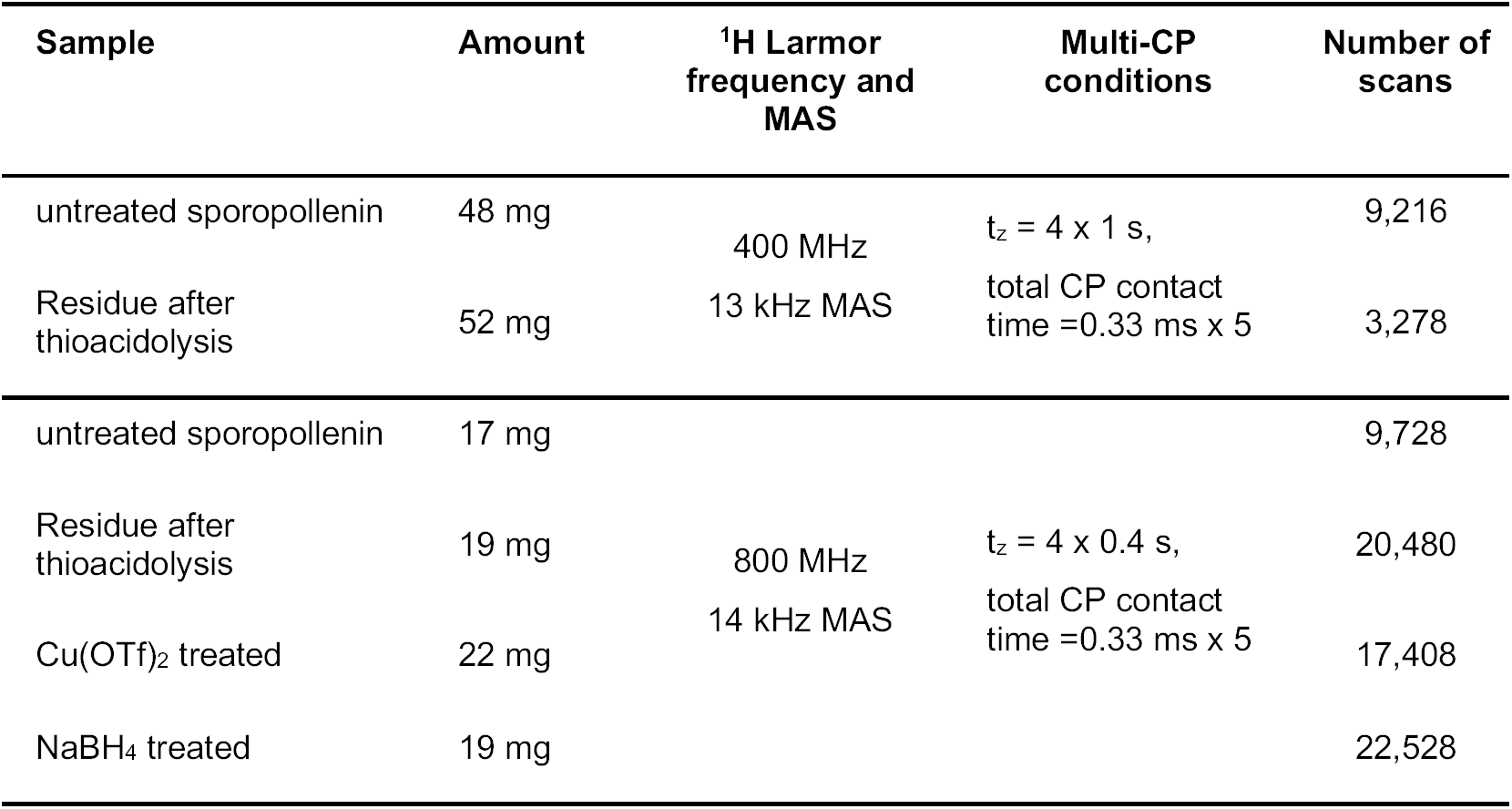
Conditions for ^13^C multi-CP solid-state NMR experiments on untreated and treated sporopollenin samples.

**Table S2.** Quantification of carbon numbers in untreated sporopollenin from ^13^C solid-state NMR spectra. The ^13^C chemical shifts, percent integrated intensity of each peak, ratio of centerband intensity to total intensity under 13 kHz MAS on the 400 MHz spectrometer, integrated intensities after correcting for sideband intensity loss, and the final relative carbon numbers, are shown. The intensity ratios of the multi-CP spectrum to the quantitative spectrum of formyl-MLF are also given and are used to calculate the relative carbon numbers. The intensity of the resolved 160-ppm peak (*) is taken to represent 2 carbons, which correspond to two sets of *p*-coumaric acids.

| $^{13}\text{C}$ chemical shifts (ppm) | Integrated intensity (%) | Centerband / Total intensity ratio | Multi-CP/ quantitative intensity ratio | Corrected integrated intensity (%) | No. of carbons | Assignment | No. carbons per group |
| --- | --- | --- | --- | --- | --- | --- | --- |
| 173.0 | 1.33 | 0.90 | 0.80 | 1.85 | 1 & 1 | Ester & α-pyrone | 5 |
| 96.4 | 1.06 | 1.00 | 1.00 | 1.06 | 1 | α-pyrone | |
| 87.5 | 1.18 | 1.00 | 1.00 | 1.18 | 1 | α-pyrone | |
| 168.4 | 2.23 | 0.90 | 0.80 | 3.10 | 1 & 2 | α-pyrone & *p*-coumaric | 18 |
| **160.0\*** | 1.50 | 0.90 | 0.75 | 2.22 | 2 | *p*-coumaric |  |
| 145.3 | 1.68 | 0.90 | 0.75 | 2.49 | 2 | *p*-coumaric |  |
| 130.0 | 3.05 | 0.87 | 0.75 | 4.67 | 4 | *p*-coumaric |  |
| 126.4 | 1.28 | 0.87 | 0.75 | 1.96 | 2 | *p*-coumaric |  |
| 116.5 | 3.13 | 0.89 | 0.75 | 4.69 | 4 | *p*-coumaric |  |
| 111.4 | 1.68 | 0.89 | 0.75 | 2.52 | 2 | *p*-coumaric |  |
| 102.8 | 5.30 | 1.00 | 1.00 | 5.30 | 4-5 | Acetal | 4-5 |
| 74.3 | 8.61 | 1.00 | 1.00 | 8.61 | 8 | O-bearing | 26 |
| 69.8 | 2.26 | 1.00 | 1.00 | 2.26 | 2 |  |  |
| 71.0 | 16.58 | 1.00 | 1.00 | 16.58 | 15 |  |  |
| 64.0 | 1.01 | 1.00 | 1.00 | 1.01 | 1 |  |  |
| 43.8 | 13.23 | 1.00 | 1.00 | 13.23 | 12 | C-(C-O) | 21 |
| 36.4 | 9.97 | 1.00 | 1.00 | 9.97 | 9 | C-(C-O) |  |
| 30.9 | 10.79 | 1.00 | 1.00 | 10.79 | 10 | $\text{CH}_2$ | 21 |
| 26.3 | 11.92 | 1.00 | 1.00 | 11.92 | 11 |  |  |
| 153.6 | 0.80 | 0.90 | 0.75 | 1.19 | 1 | - |  |
| 53.2 | 1.47 | 1.00 | 1.00 | 1.47 | 1 | - |  |

**Table S3.** Comparison of the carbon numbers for each major functional group determined from ^13^C solid-state NMR spectra and the carbon numbers in the proposed structure of pine sporopollenin.

| $^{13}\text{C}$ chemical shift ranges | Functional groups | Assigned No. of carbons in 2 units | No. of carbons in the proposed structure |
| --- | --- | --- | --- |
| 173, 168, 96-87 ppm | Ester & α-pyrone | $5 \times 2 = 10$ | 12 |
| 168-111 ppm | p-coumaric acid | $18 \times 2 = 36$ | 36 |
| 103 ppm | Acetal | $4-5 \times 2 = 8-10$ | 8 |
| 74-63 ppm | O-bearing | $26 \times 2 = 52$ | 54 |
| 44-36 ppm | C-(CO) | $21 \times 2 = 42$ | 46 |
| 31-26 ppm | Aliphatic $\text{CH}_2$ | $21 \times 2 = 42$ | 44 |
| | Unassigned | $2 \times 2 = 4$ | |
|  | **Total** | **194 - 196** | **200** |

**Table S4.**
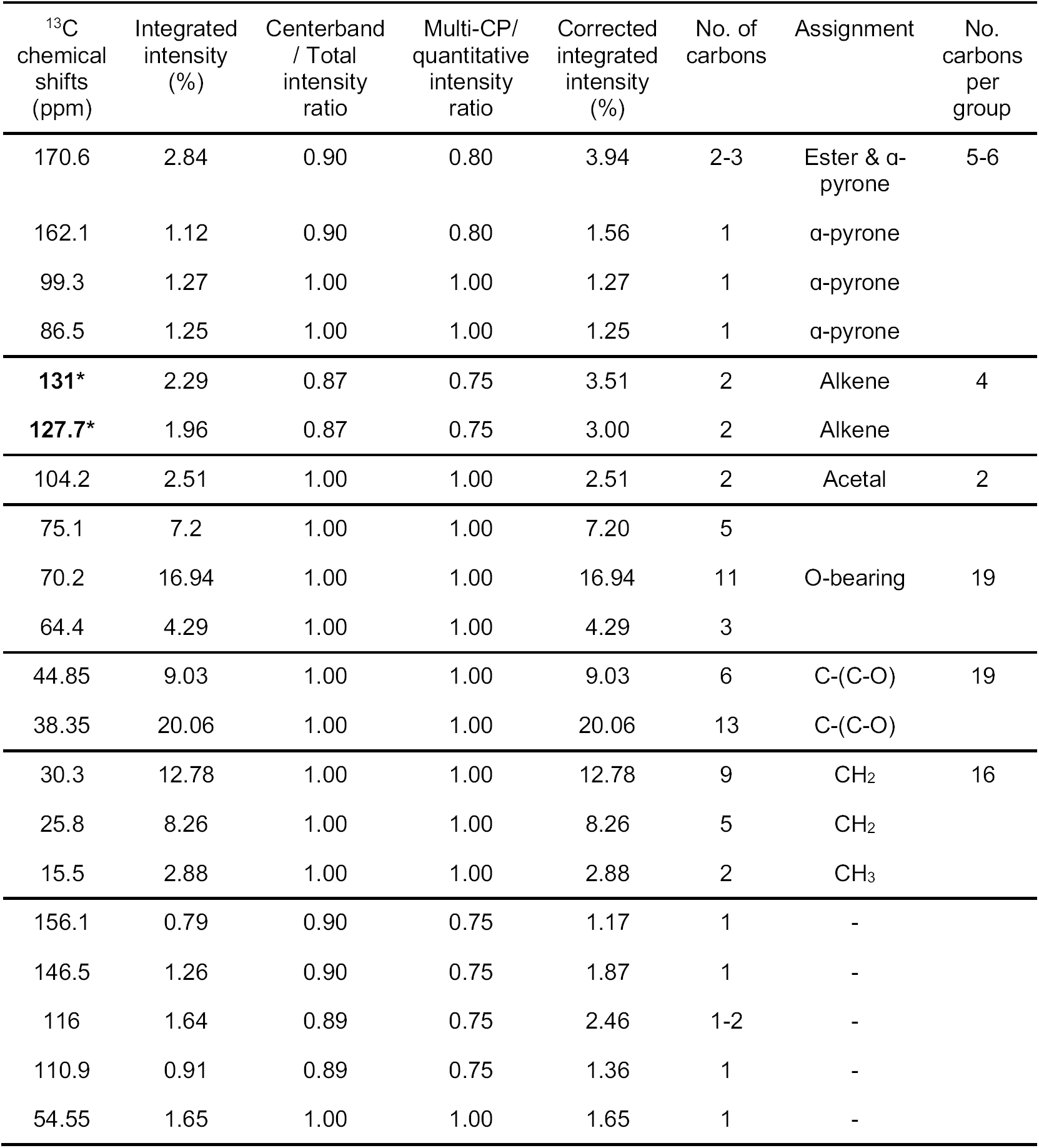
Quantification of carbon numbers in insoluble residue (**R**) after thioacidolysis from ^13^C NMR spectra. The ^13^C chemical shifts, percent integrated intensity of each peak, ratio of centerband intensity to total intensity under 13 kHz MAS on a 400 MHz spectrometer, integrated intensities after correcting for sideband intensity loss, and the final relative carbon numbers, are shown. The intensity ratios of the multi-CP spectrum to the quantitative spectrum of formyl-MLF are also shown and are used to calculate the relative carbon numbers. The integrated intensities of the resolved 131-ppm and 128-ppm peaks (*) are taken to represent 2 carbons, which correspond to two sets of alkene groups.

**Table S5.**
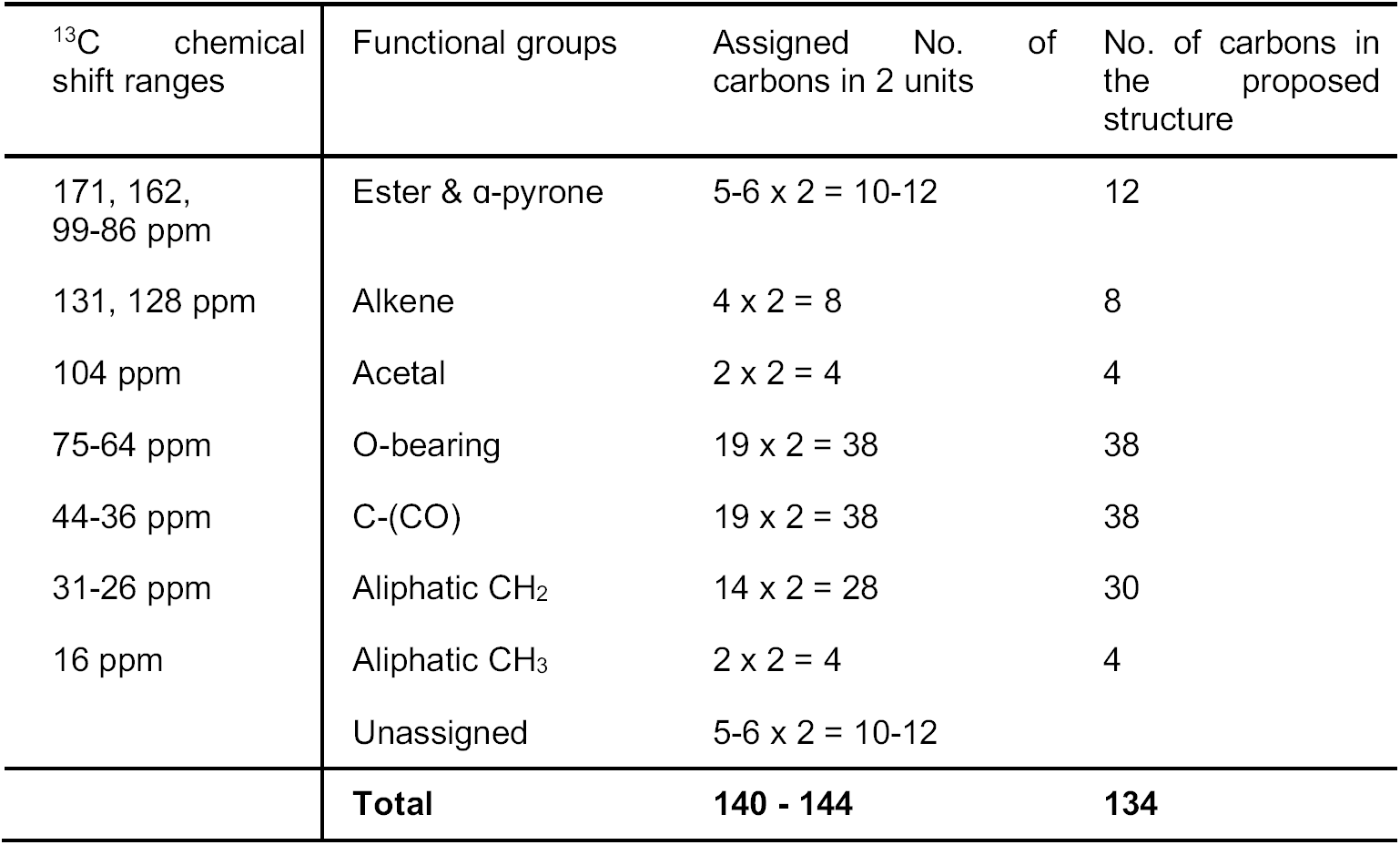
Comparison of the carbon numbers for each major functional group determined from ^13^C solid-state NMR spectra and the carbon numbers in the proposed residue **R** structure.

## NMR data for chemical degradative products described in this study

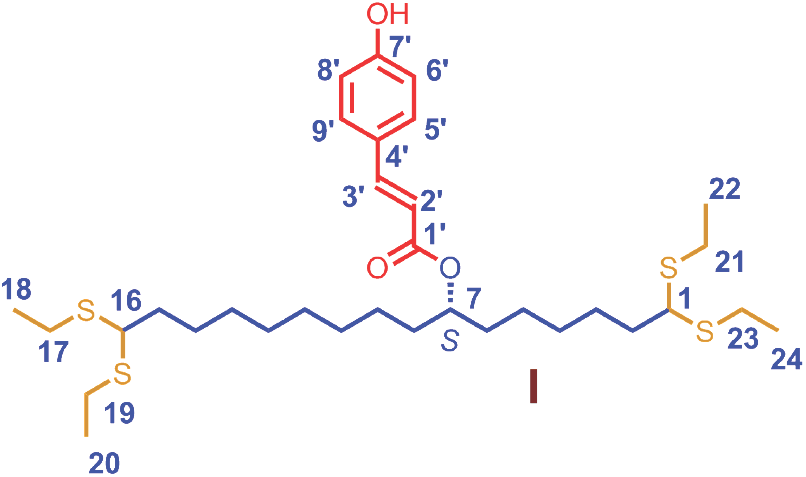

^1^H NMR (CDCl_3_, 400 MHz), *δ* 7.59 (1H, d, *J* = 16.0 Hz, H-3’), 7.42 (2H, d, *J* = 8.4 Hz, H-5’,9’), 6.82 (2H, d, *J* = 8.4 Hz, H-6’, 8’), 6.28 (1H, d, *J* = 16.0 Hz, H-2’), 4.97 (1H, t, *J* = 5.6 Hz, H-7), 3.74 (2H, t, *J* = 6.8 Hz, H-1, 16), 2.60 (8H, m, H-17, 19, 21, 23), 1.76 (4H, m, H-2,15), 1.53 (12H, m, H-3, 5, 6, 8, 9, 14), 1.29 (10H, m, H-4,10, 11, 12, 13), 1.23 (12H, t, *J* = 7.2 Hz, H-18, 20, 22, 24).

^13^C NMR (CDCl_3_, 100 MHz), *δ* 167.2 (C-1’), 157.4 (C-7’), 144.0 (C-3’), 129.9 (C-5’, 9’), 127.5 (C-4’), 116.3 (C-2’), 115.8 (C-6’, 8’), 74.2 (C-7), 51.4, 51.5 (C-1, 16), 36.0, 36.1 (C-2, 15), 34.2, 34.3 (C-6, 8), 29.1, 29.2, 29.4, 29.5, 29.6 (C-4, 10, 11, 12, 13), 27.4, 27.5 (C-3, 14), 25.2, 25.4 (C-5, 9), 24.1 (C-17, 19, 21, 23), 14.5 (C-18, 20, 22, 24).

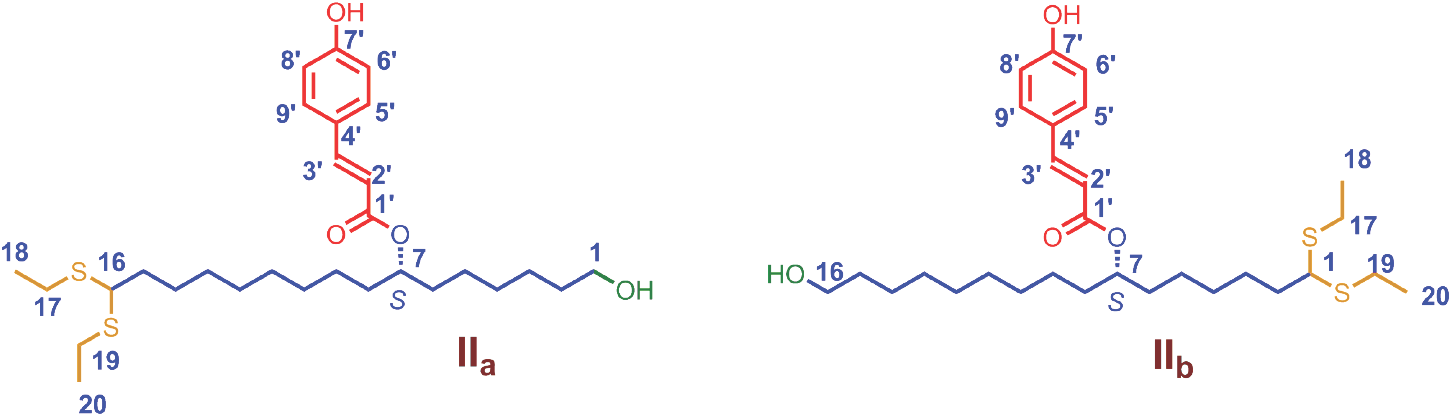

^1^H NMR (**II_a_**, CDCl_3_, 400 MHz), *δ* 7.61 (1H, d, *J* = 16.0 Hz, H-3’), 7.43 (2H, d, *J* = 8.0 Hz, H-5’,9’), 6.84 (2H, d, *J* = 8.0 Hz, H-6’, 8’), 6.30 (1H, d, *J* = 16.0 Hz, H-2’), 4.99 (1H, t, *J* = 5.6 Hz, H-7), 3.76 (1H, t, *J* = 6.8 Hz, H-16), 3.63 (2H, m, H-1), 2.63 (4H, m, H-17, 19), 1.78 (2H, m, H-15), 1.56 (10H, m, H-5, 6, 8, 9, 14), 1.31 (14H, m, H-2, 3, 4, 10, 11, 12, 13), 1.25 (6H, t, *J* = 7.2 Hz, H-18, 20).

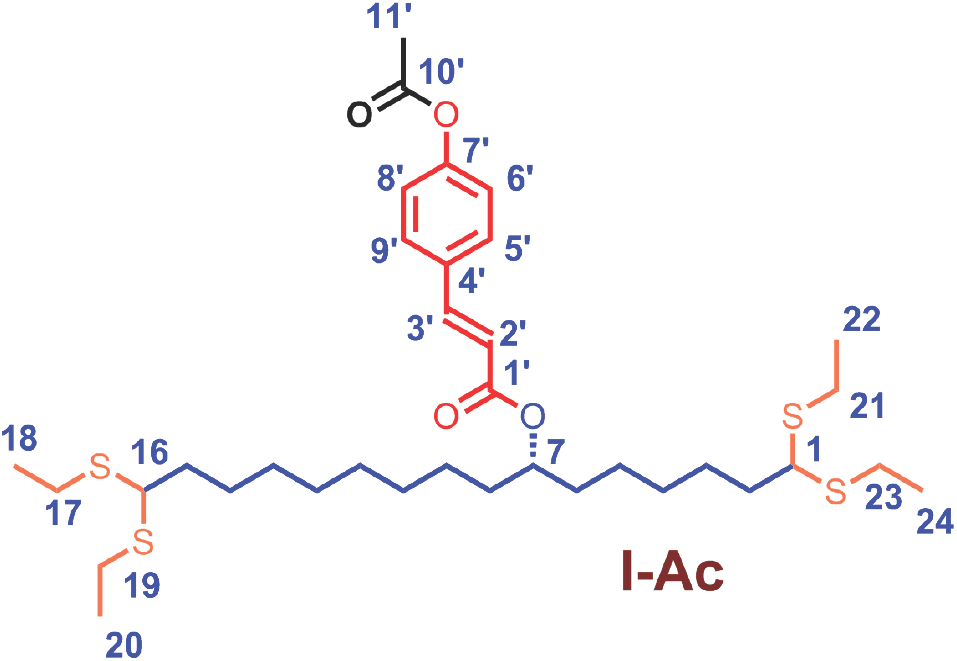

^1^H NMR (CDCl_3_, 400 MHz), *δ* 7.64 (1H, d, *J* = 16.0 Hz, H-3’), 7.54 (2H, d, *J* = 8.4 Hz, H-5’,9’), 7.12 (2H, d, *J* = 8.4 Hz, H-6’, 8’), 6.38 (1H, d, *J* = 16.0 Hz, H-2’), 4.99 (1H, m, H-7), 3.76 (2H, t, *J* = 6.8 Hz, H-1, 16), 2.61 (8H, m, H-17, 19, 21, 23), 2.31 (3H, s, H-11’), 1.77 (4H, m, H-2,15), 1.55 (12H, m, H-3, 5, 6, 8, 9, 14), 1.26 (10H, m, H-4,10, 11, 12, 13), 1.23 (12H, t, *J* = 7.2 Hz, H-18, 20, 22, 24).

^13^C NMR (CDCl_3_, 100 MHz), *δ* 169.3 (C-10’), 166.9 (C-1’), 152.1 (C-7’), 143.4 (C-3’), 132.4 (C-4’), 129.3 (C-5’, 9’), 122.2 (C-6’, 8’), 119.0 (C-2’), 74.6 (C-7), 51.6, 51.5 (C-1, 16), 36.3, 36.2 (C-2, 15), 34.4, 34.3 (C-6, 8), 29.2, 29.3, 29.5, 29.6, 29.7 (C-4, 10, 11, 12, 13), 27.6, 27.5 (C-3, 14), 25.5, 25.3 (C-5, 9), 24.2 (C-17, 19, 21, 23), 14.7 (C-18, 20, 22, 24).

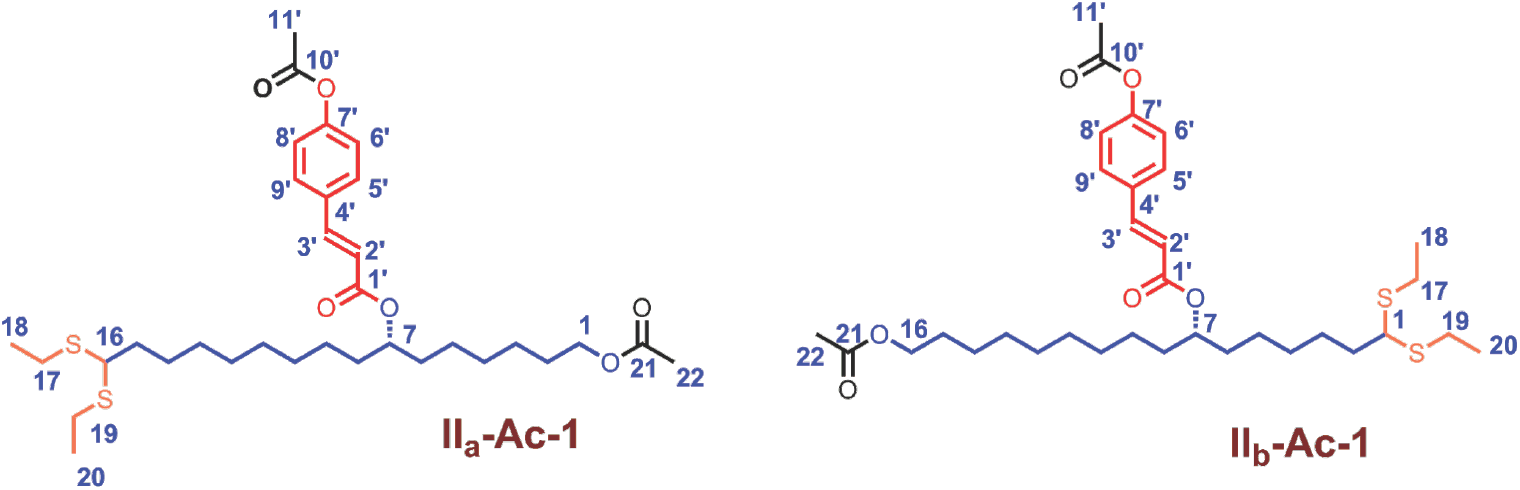

^1^H NMR (**II_a_-Ac-1**, CDCl_3_, 400 MHz), *δ* 7.65 (1H, d, *J* = 16.0 Hz, H-3’), 7.55 (2H, d, *J* = 8.4 Hz, H-5’,9’), 7.12 (2H, d, *J* = 8.4 Hz, H-6’, 8’), 6.39 (1H, d, *J* = 16.0 Hz, H-2’), 5.00 (1H, t, *J* = 5.6 Hz, H-7), 4.04 (2H, t, *J* = 6.8 Hz, H-1), 3.76 (1H, t, *J* = 6.8 Hz, H-16), 2.63 (4H, m, H-17, 19), 2.31 (3H, s, H-11’), 2.03 (3H, s, H-22), 1.78 (2H, m, H-15), 1.55 (10H, m, H-5, 6, 8, 9, 14), 1.26 (14H, m, H-2, 3, 4, 10, 11, 12, 13), 1.25 (6H, t, *J* = 7.2 Hz, H-18, 20).

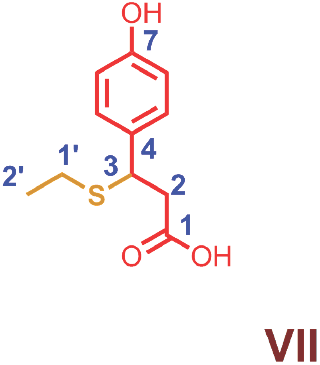

^1^H NMR (CDCl_3_, 400 MHz), *δ* 7.13 (2H, d, *J* = 8.4 Hz, H-5, 9), 6.69 (2H, d, *J* = 8.4 Hz, H-6, 8), 4.14 (1H, t, *J* = 7.2 Hz, H-3), 2.73 (2H, m, H-2), 2.29 (2H, m, H-1’), 1.07 (3H, t, *J* = 7.2 Hz, H-2’).

^13^C NMR (CDCl_3_, 100 MHz), *δ* 172.2 (C-1), 156.8 (C-7), 132.2 (C-4), 129.1 (C-5, 9), 115.5 (C-6, 8), 44.3 (C-2), 41.6 (C-3), 24.7 (C-1’), 14.8 (C-2’).

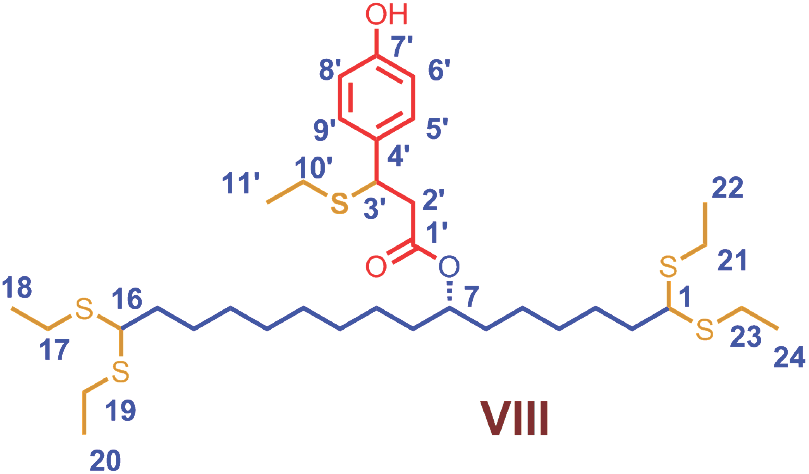

^1^H NMR (CDCl_3_, 400 MHz), *δ* 7.18 (2H, d, *J* = 8.4 Hz, H-5’, 9’), 6.75 (2H, d, *J* = 8.4 Hz, H-6’, 8’), 4.76 (1H, m, H-7), 4.25 (1H, m, H-3’), 3.77 (2H, t, *J* = 6.8 Hz, H-1, 16), 2.84 (2H, m, H-2’), 2.64 (8H, m, H-17, 19, 21, 23), 2.33 (2H, m, H-10’), 1.76 (4H, m, H-2,15), 1.54-1.31 (8H, m, H-3, 6, 8, 14), 1.26 (14H, m, H-4, 5, 9, 10, 11, 12, 13), 1.25 (12H, m, H-18, 20, 22, 24), 1.15 (3H, t, *J* = 7.6 Hz, H-11’).

^13^C NMR (CDCl_3_, 100 MHz), *δ* 170.8, 170.9 (C-10’), 155.1, 155.2 (C-7’), 133.0, 133.2 (C-4’), 129.0, 129.1 (C-5’, 9’), 115.4, 115.5 (C-6’, 8’), 74.8, 74.9 (C-7), 51.5, 51.7 (C-1, 16), 44.7, 44.4 (C-2’), 42.0, 41.9 (C-3’), 36.2, 36.1 (C-2, 15), 34.4, 34.2, 34.1 (C-6, 8), 29.6, 29.5, 29.4, 29.4, 29.3, 29.2 (C-4, 10, 11, 12, 13), 27.6, 27.5 (C-3, 14), 25.4, 25.3, 25.2, 25.1 (C-5, 9), 24.4, 24.3, 24.3, 24.2, 24.2 (C-17, 19, 21, 23, 10’), 14.7, 14.6 (C-18, 20, 22, 24), 14.5 (C-11’).

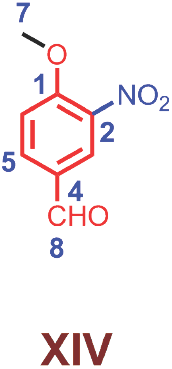

^1^H NMR (CDCl_3_, 400 MHz), *δ* 9.95 (1H, s, H-8), 8.42 (1H, d, *J* = 2.0 Hz, H-3), 8.20 (1H, dd, *J* = 8.8, 2.0 Hz, H-5), 7.58 (1H, d, *J* = 8.8 Hz, H-6), 4.04 (3H, s, H-7).

^13^C NMR (CDCl_3_, 100 MHz), *δ* 190.5 (C-8), 156.2 (C-1), 135.0 (C-2), 128.8 (C-5), 126.4 (C-3, 4), 115.0 (C-6), 57.5 (C-7).

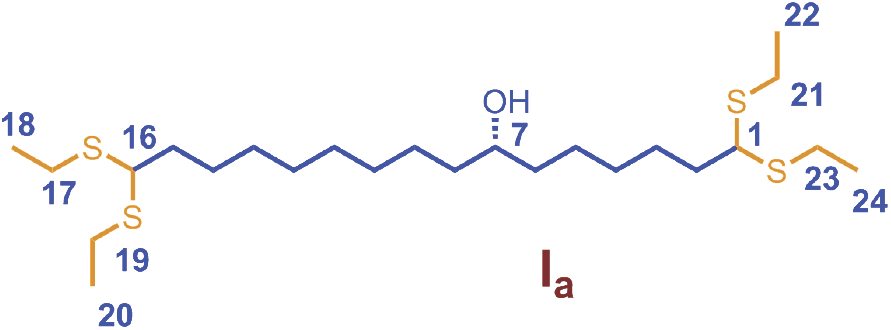

^1^H NMR (CDCl_3_, 400 MHz), *δ* 3.79 (2H, t, *J* = 6.8 Hz, H-1, 16), 3.57 (1H, m, H-7), 2.62 (8H, m, H-17, 19, 21, 23), 1.78 (4H, m, H-2,15), 1.52 (4H, m, H-3, 14), 1.42 (4H, m, H-6, 8), 1.26 (14H, m, H-4, 5, 9, 10, 11, 12, 13), 1.24 (12H, t, *J* = 7.2 Hz, H-18, 20, 22, 24).

^13^C NMR (CDCl_3_, 100 MHz), *δ* 72.0 (C-7), 51.6, 51.5 (C-1, 16), 37.5, 37.6 (C-2, 15), 36.3, 36.2 (C-6, 8), 29.8, 29.6, 29.5, 29.4, 29.3 (C-4, 10, 11, 12, 13), 27.6, 27.5 (C-3, 14), 25.8, 25.6 (C-5, 9), 24.2 (C-17, 19, 21, 23), 14.7 7 (C-18, 20, 22, 24).

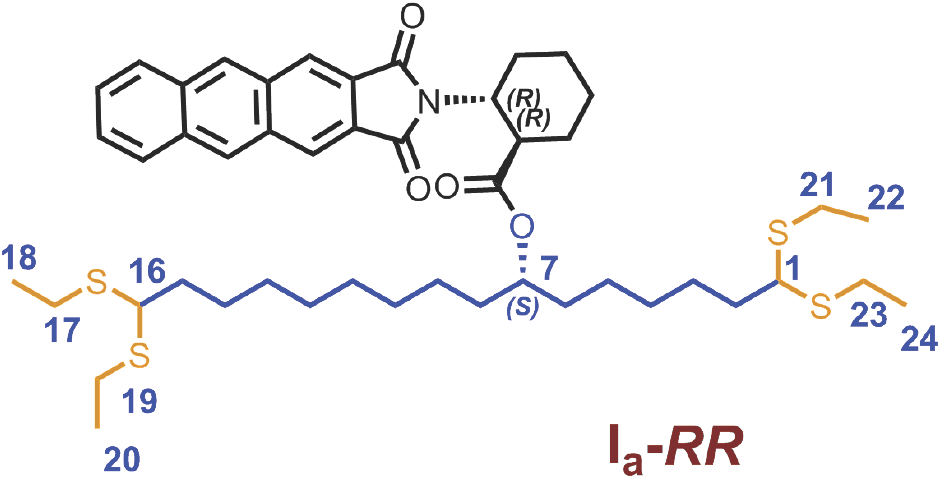

^1^H NMR (CDCl_3_, 600 MHz), *δ* 3.73 (1H, t, *J* = 7.2 Hz, H-7), 3.69 (1H, t, *J* = 6.6 Hz, H-16), 3.61 (1H, t, *J* = 6.6 Hz, H-1), 2.62 (8H, m, H-17, 19, 21, 23), 1.86 (4H, m, H-2,15), 1.62 (4H, m, H-3, 14), 1.33 (4H, m, H-6, 8), 1.27 (14H, m, H-4, 5, 9, 10, 11, 12, 13), 1.21 (6H, t, *J* = 7.8 Hz, H-18, 20), 1.20 (6H, t, *J* = 7.8 Hz, H-22, 24).

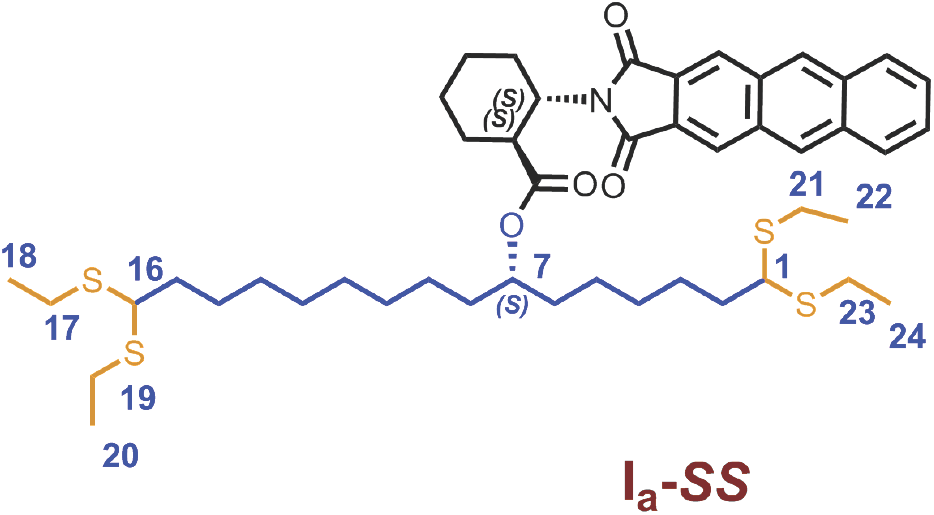

^1^H NMR (CDCl_3_, 600 MHz), *δ* 3.72 (1H, t, *J* = 6.6 Hz, H-7), 3.71 (1H, t, *J* = 7.2 Hz, H-16), 3.52 (1H, t, *J* = 7.2 Hz, H-1), 2.56 (8H, m, H-17, 19, 21, 23), 1.87 (4H, m, H-2,15), 1.67 (4H, m, H-3, 14), 1.47 (4H, m, H-6, 8), 1.25 (14H, m, H-4, 5, 9, 10, 11, 12, 13), 1.19 (6H, t, *J* = 7.2 Hz, H-18, 20), 1.18 (6H, t, *J* = 7.2 Hz, H-22, 24).

## References

1. Weng, J.-K., Philippe, R. N. & Noel, J. P. The rise of chemodiversity in plants. Science 336, 1667–1670 (2012).

2. Kim, S. S. & Douglas, C. J. Sporopollenin monomer biosynthesis in arabidopsis. J. Plant Biol. 56, 1–6 (2013).

3. Lu, F. & Ralph, J. Non-degradative dissolution and acetylation of ball-milled plant cell walls: high-resolution solution-state NMR. Plant J. 35, 535–544 (2003).

4. Ahlers, F., Thom, I., Lambert, J., Kuckuk, R. & Rolf Wiermann. 1H NMR analysis of sporopollenin from Typha Angustifolia. Phytochemistry 50, 1095–1098 (1999).

5. Rolando, C., Monties, B. & Lapierre, C. Thioacidolysis. in Methods in Lignin Chemistry 334–349 (Springer, Berlin, Heidelberg, 1992).

6. Seco, J. M., Quiñoá, E. & Riguera, R. The Assignment of Absolute Configuration by NMR. Chem. Rev. 104, 17–118 (2004).

7. Imaizumi, K., Terasima, H., Akasaka, K. & Ohrui, H. Highly Potent Chiral Labeling Reagents for the Discrimination of Chiral Alcohols. Anal. Sci. 19, 1243–1249 (2003).

8. Ohrui, H. Development of highly potent chiral discrimination methods that solve the problems of diastereomer method. Anal. Sci. 24, 31–38 (2008).

9. Ohtaki, T., Akasaka, K., Kabuto, C. & Ohrui, H. Chiral discrimination of secondary alcohols by both 1H-NMR and HPLC after labeling with a chiral derivatization reagent, 2-(2, 3-anthracenedicarboximide)cyclohexane carboxylic acid. Chirality 17, 171–176 (2005).

10. Zhang, Y.-J., Dayoub, W., Chen, G.-R. & Lemaire, M. Environmentally benign metal triflate-catalyzed reductive cleavage of the C–O bond of acetals to ethers. Green Chem. 13, 2737–2742 (2011).

11. Zhang, Y. J., Dayoub, W. & Chen, G. R. TMDS as a Dual-Purpose Reductant in the Regioselective Ring Cleavage of Hexopyranosyl Acetals to Ethers. European Journal of (2012).

12. Dick-Pérez, M. et al. Structure and Interactions of Plant Cell-Wall Polysaccharides by Two- and Three-Dimensional Magic-Angle-Spinning Solid-State NMR. Biochemistry 50, 989–1000 (2011).

13. Wang, T., Phyo, P. & Hong, M. Multidimensional solid-state NMR spectroscopy of plant cell walls. Solid State Nucl. Magn. Reson. 78, 56–63 (2016).

14. Phyo, P., Wang, T., Yang, Y., O’Neill, H. & Hong, M. Direct Determination of Hydroxymethyl Conformations of Plant Cell Wall Cellulose Using 1H Polarization Transfer Solid-State NMR. Biomacromolecules 19, 1485–1497 (2018).

15. Phyo, P. et al. Gradients in Wall Mechanics and Polysaccharides along Growing Inflorescence Stems. Plant Physiol. 175, 1593–1607 (2017).

16. Guilford, W. J., Schneider, D. M., Labovitz, J. & Opella, S. J. High resolution solid state C NMR spectroscopy of sporopollenins from different plant taxa. Plant Physiol. 86, 134–136 (1988).

17. Mao, J., Cory, R. M., McKnight, D. M. & Schmidt-Rohr, K. Characterization of a nitrogen-rich fulvic acid and its precursor algae from solid state NMR. Org. Geochem. 38, 1277–1292 (2007).

18. Mao, J.-D. et al. Abundant and Stable Char Residues in Soils: Implications for Soil Fertility and Carbon Sequestration. Environ. Sci. Technol. 46, 9571–9576 (2012).

19. Johnson, R. L. & Schmidt-Rohr, K. Quantitative solid-state 13C NMR with signal enhancement by multiple cross polarization. J. Magn. Reson. 239, 44–49 (2014).

20. Kim, S. S., Grienenberger, E. & Lallemand, B. LAP6/POLYKETIDE SYNTHASE A and LAP5/POLYKETIDE SYNTHASE B encode hydroxyalkyl α-pyrone synthases required for pollen development and …. The Plant (2010).

21. Grienenberger, E. et al. Analysis of TETRAKETIDE α-PYRONE REDUCTASE function in Arabidopsis thaliana reveals a previously unknown, but conserved, biochemical pathway in sporopollenin monomer biosynthesis. Plant Cell 22, 4067–4083 (2010).

22. Rasouli, M., Ostovar-Ravari, A. & Shokri-Afra, H. Characterization and improvement of phenol-sulfuric acid microassay for glucose-based glycogen. Eur. Rev. Med. Pharmacol. Sci. 18, 2020–2024 (2014).

23. Mao, J. D. & Schmidt-Rohr, K. Accurate quantification of aromaticity and nonprotonated aromatic carbon fraction in natural organic matter by 13C solid-state nuclear magnetic resonance. Environ. Sci. Technol. 38, 2680–2684 (2004).

24. Mao, J.-D. & Schmidt-Rohr, K. Methylene spectral editing in solid-state 13C NMR by three-spin coherence selection. J. Magn. Reson. 176, 1–6 (2005).

25. Liu, R., He, B. & Chen, X. Degradation of poly(vinyl butyral) and its stabilization by bases. Polym. Degrad. Stab. 93, 846–853 (2008).

26. Weng, J.-K. & Chapple, C. The origin and evolution of lignin biosynthesis. New Phytol. 187, 273–285 (2010).

27. Hayatsu, R. E. Botto, R. L. McBeth, R. G. Scott, R. & Winans, R. Chemical structure of a sporinite from a lignite: Comparison with a synthetic sporinite transformed from sporopollenin. Prepr. Pap., Am. Chem. Soc., Div. Fuel Chem 32, 1–8 (1987).

28. Dobritsa, A. A. et al. CYP704B1 is a long-chain fatty acid omega-hydroxylase essential for sporopollenin synthesis in pollen of Arabidopsis. Plant Physiol. 151, 574–589 (2009).

29. Morant, M. et al. CYP703 is an ancient cytochrome P450 in land plants catalyzing in-chain hydroxylation of lauric acid to provide building blocks for sporopollenin synthesis in pollen. Plant Cell 19, 1473–1487 (2007).

30. Austin, M. B. & Noel, J. P. The Chalcone Synthase Superfamily of Type III Polyketide Synthases. ChemInform 34, (2003).

31. de Azevedo Souza, C. et al. A novel fatty Acyl-CoA Synthetase is required for pollen development and sporopollenin biosynthesis in Arabidopsis. Plant Cell 21, 507–525 (2009).

## Supplementary References

1. Shaw, G. & Yeadon, A. Chemical studies on the constitution of some pollen and spore membranes. J. Chem. Soc. Perkin 1 1, 16–22 (1966).

2. Schulze Osthoff, K. & Wiermann, R. Phenols as Integrated Compounds of Sporopollenin from Pinus Pollen. J. Plant Physiol. 131, 5–15 (1987).

3. Wehling, K., Niester, C., Boon, J. J., Willemse, M. T. & Wiermann, R. p-Coumaric acid - a monomer in the sporopollenin skeleton. Planta 179, 376–380 (1989).

4. Gubatz, S., Rittscher, M., Meuter, A., Nagler, A. & Wiermann, R. Tracer Experiments on Sporopollenin Biosynthesis. Grana 32, 12–17 (1993).

5. Meuter-Gerhards, A., Riegert, S. & Wiermann, R. Studies on Sporopollenin Biosynthesis in Cucurbita maxima (DUCH.) — II. The Involvement of Aliphatic Metabolism. J. Plant Physiol. 154, 431–436 (1999).

6. Couderchet, M., Schmalfuß, J. & Böger, P. Incorporation of Oleic Acid into Sporopollenin and Its Inhibition by the Chloroacetamide Herbicide Metazachlor. Pestic. Biochem. Physiol. 55, 189–199 (1996).

7. Ahlers, F., Thom, I., Lambert, J., Kuckuk, R. & Rolf Wiermann. 1H NMR analysis of sporopollenin from Typha Angustifolia. Phytochemistry 50, 1095–1098 (1999).

8. Ahlers, F., Lambert, J. & Wiermann, R. Acetylation and silylation of piperidine solubilized sporopollenin from pollen of Typha angustifolia L. Z. Naturforsch. C 58, 807–811 (2003).

9. Kettley, S. J. Novel derivatives of sporopollenin for potential applications in solid phase organic synthesis and drug delivery. (2001).

10. Rolando, C., Monties, B. & Lapierre, C. Thioacidolysis. in Methods in Lignin Chemistry 334–349 (Springer, Berlin, Heidelberg, 1992).

11. Sawamura, K., Tobimatsu, Y., Kamitakahara, H. & Takano, T. Lignin Functionalization through Chemical Demethylation: Preparation and Tannin-Like Properties of Demethylated Guaiacyl-Type Synthetic Lignins. ACS Sustainable Chemistry & Engineering 5, 5424–5431 (2017).

12. Thompson, R. S., Jacques, D., Haslam, E. & Tanner, R. J. N. Plant proanthocyanidins. Part Introduction; the isolation, structure, and distribution in nature of plant procyanidins. J. Chem. Soc. Perkin 1 0, 1387–1399 (1972).

13. Osman, S. F., Irwin, P., Fett, W. F., O’Connor, J. V. & Parris, N. Preparation, isolation, and characterization of cutin monomers and oligomers from tomato peels. J. Agric. Food Chem. 47, 799–802 (1999).

14. Lu, F. & Ralph, J. Non-degradative dissolution and acetylation of ball-milled plant cell walls: high-resolution solution-state NMR. Plant J. 35, 535–544 (2003).

15. Rasouli, M., Ostovar-Ravari, A. & Shokri-Afra, H. Characterization and improvement of phenol-sulfuric acid microassay for glucose-based glycogen. Eur. Rev. Med. Pharmacol. Sci. 18, 2020–2024 (2014).

16. Zhang, Y.-J., Dayoub, W., Chen, G.-R. & Lemaire, M. TMDS as a Dual-Purpose Reductant in the Regioselective Ring Cleavage of Hexopyranosyl Acetals to Ethers. Eur. J. Org. Chem. 2012, 1960–1966 (2012).

17. Ohtaki, T., Akasaka, K., Kabuto, C. & Ohrui, H. Chiral discrimination of secondary alcohols by both 1H-NMR and HPLC after labeling with a chiral derivatization reagent, 2-(2, 3-anthracenedicarboximide) cyclohexane carboxylic acid. Chirality 17, (2005).

18. Johnson, R. L. & Schmidt-Rohr, K. Quantitative solid-state 13C NMR with signal enhancement by multiple cross polarization. J. Magn. Reson. 239, 44–49 (2014).

19. Mao, J. D. & Schmidt-Rohr, K. Accurate quantification of aromaticity and nonprotonated aromatic carbon fraction in natural organic matter by 13C solid-state nuclear magnetic resonance. Environ. Sci. Technol. 38, 2680–2684 (2004).

20. Guilford, W. J., Schneider, D. M., Labovitz, J. & Opella, S. J. High resolution solid state C NMR spectroscopy of sporopollenins from different plant taxa. Plant Physiol. 86, 134–136 (1988).

21. Mao, J.-D. & Schmidt-Rohr, K. Methylene spectral editing in solid-state 13C NMR by three-spin coherence selection. J. Magn. Reson. 176, 1–6 (2005).

22. Shibuya, T., Funamizu, M. & Kitahara, Y. Novel p-coumaric acid esters from Pinus densiflora pollen. Phytochemistry 17, 979–981 (1978).

23. Tian, S. et al. Isolation and identification of oligomers from partial degradation of lime fruit cutin. J. Agric. Food Chem. 56, 10318–10325 (2008).

24. Massiot, D. et al. Modelling one- and two-dimensional solid-state NMR spectra. Magn. Reson. Chem. 40, 70–76 (2001).

25. Zhang, Y.-J., Dayoub, W., Chen, G.-R. & Lemaire, M. Environmentally benign metal triflate-catalyzed reductive cleavage of the C–O bond of acetals to ethers. Green Chem. 13, 2737 (2011).

